# Contriving a chimeric polyvalent vaccine to prevent infections caused by Herpes Simplex Virus (Type-1 and Type-2): an exploratory immunoinformatic approach

**DOI:** 10.1101/679639

**Authors:** Mahmudul Hasan, Md Shiful Islam, Sourav Chakraborty, Abu Hasnat Mustafa, Kazi Faizul Azim, Ziaul Faruque Joy, Md Nazmul Hossain, Shakhawat Hossain Foysal, Md Nazmul Hasan

## Abstract

Herpes simplex virus type 1 (HSV-1) and 2 (HSV-2) cause a variety of infections including oral-facial infections, genital herpes, herpes keratitis, cutaneous infection and so on. To date, FDA-approved licensed HSV vaccine is not available yet. Hence, the study was conducted to identify and characterize an effective epitope based polyvalent vaccine against both types of Herpes Simplex Virus through targeting six viral proteins. The selected proteins were retrieved from viralzone and assessed to design highly antigenic epitopes by binding analyses of the peptides with MHC class-I and class-II molecules, antigenicity screening, transmembrane topology screening, allergenicity and toxicity assessment, population coverage analysis and molecular docking approach. The final vaccine was constructed by the combination of top CTL, HTL and BCL epitopes from each protein along with suitable adjuvant and linkers. Physicochemical and secondary structure analysis, disulfide engineering, molecular dynamic simulation and codon adaptation were further employed to develop a unique multi-epitope peptide vaccine. Docking analysis of the refined vaccine structure with different MHC molecules and human immune TLR-2 receptor demonstrated higher interaction. Complexed structure of the modeled vaccine and TLR-2 showed minimal deformability at molecular level. Moreover, translational potency and microbial expression of the modeled vaccine was analyzed with pET28a(+) vector for *E. coli* strain strain K12. The study enabled design of a novel chimeric polyvalent vaccine to confer broad range immunity against both HSV serotypes. However, further wet lab based research using model animals are highly recommended to experimentally validate our findings.

## Introduction

Herpes simplex virus 1 (HSV-1) and herpes simplex virus2 (HSV-2) are the two members of the Herpesviridae family. Oro-labial infection, mostly caused by HSV-1 is found in children while HSV-2 has been associated with genital disorders (Nahmias et al., 1990; Lafferty et al., 1987). Symptomatic HSV-1 infections are usually manifested as recurrent oro-labial and facial lesions. On the other hand, HSV-2 infections are usually symptomatic and are typically characterized by recurrent, painful vesicular and ulcerative lesions in the genital and anal areas (Corey & Wald, 1999; Stanberry et al., 1997; Kimberlin & Rouse, 2004). In addition, genital herpes may lead substantial psychological morbidity (Patel et al., 2001). Moreover, HSV-2 infection has received renewed attention in recent years as the epidemiological synergy between HSV-2 and HIV has been revealed. It has been claimed that HSV-2 infection increases the risk of HIV acquisition by approximately three-fold (Freeman et al., 2006; Brown et al., 2007; Reynolds et al., 2003). Some serious diseases such as blindness, encephalitis and neonatal infections are also being reported for both HSV-1 and HSV-2 infections (Stanberry et al., 1997).

Both of HSV-1 and HSV-2 are ubiquitous and contagious in nature (Ryan & Ray, 2004). Recent data have claimed that HSV-1 is involved in substantial proportion of genital herpes infections, particularly in Europe (Lowhagen et al., 2002; Lafferty et al., 2017; Strutt et al., 2003; Scoular et al., 2002; Christie et al., 1997). In the United States, HSV-1 infections have been found surprisingly in college-students and other selected populations (Lafferty et al., 2000; Ribes et al., 2001; Roberts et al., 2003; Langenberg et al., 1999). Besides, 50% of the adults were seropositive for HSV-1 and about 15% of those who were sexually active were infected by HSV-2 (McQuillan et al., 2018; Xu et al., 2018; Roberts et al., 2003; Looker et al., 2017). Literature review suggests that 536 million people exhibited existing (prevalent) and 23.6 million people had new (incident) HSV-2 infection all over the world (Looker et al., 2012) where more women (267 million) than men (150 million) had prevalent infection. Greater biological susceptibility to HSV-2 by women may be attributed to the most likely reason for this [Holmes et al., 2008; Corey et al., 2004; Kaushic et al., 2011; Glynn et al., 2001). The global burden of HSV infection is huge, exposing more than 400 million people at increased risk of genital ulcer disease, transmission of HSV-2 to partners or neonates and HIV acquisition. These estimates highlight the critical need for development of vaccines and other new prevention strategies to combat HSV associated infections (Looker et al., 2012).

Despite the deliberate research for more than 60 years, development of an effective HSV vaccine has remained elusive and challenging. To date, FDA-approved licensed HSV vaccine is not available yet (Whitley & Baines, 2018). Vaccine development is still complicated owing to some unique characteristics of herpes viruses which include complexity of the virus replication cycle (i.e., primary, latent and recurrent phases of infection), sophisticated immune evasion strategies of the virus, a high number of protein candidates by the large and complex herpes genome (Dasgupta et al., 2009). Development of vaccines often requires prior understanding of the immunological insights during the natural course of an infection. So far, researcher are trying to test efficacy of four types of experimental vaccines against HSV (Raghuwanshi et al., 2018) which are classical formalin-inactivated crude or partially purified infected cell extract, semi-purified or highly purified infected cell extract showing low concentration of non-viral proteins and a minimum of residual vDNA, recombinant virus glycoproteins along with nonstructural polypeptides prepared as fusion proteins and live attenuated HSV1 or HSV2 virus vaccine. Though many attempts have been taken, neither a therapeutic nor preventive vaccine exists for HSV-1 or HSV-2 (Whitley & Baines, 2018). Many vaccines were investigated which ultimately yielded disappointing results in human trials (Whitley & Baines, 2018). The development of several promising HSV vaccines has been terminated recently due to modest or controversial therapeutic effects in humans (Whitley and Baines, 2018). A vaccine trial named VICal was terminated in June 2018 as the magnitude of the effect in the test population was unsatisfactory (Whitley & Baines, 2018; Veselenak et al., 2012). Genocea, a company in USA has confirmed not to develop its GEN-003 vaccine anymore after results from a partially conducted phase III trial revealed its limited clinical efficacy (Whitley & Baines, 2018; Skoberne et al., 2013). A prophylactic vaccine also failed to prevent HSV-2 infection and diseases in clinical trials (Belshe et al., 2012) where the study population was representative of the general population of HSV-1 and HSV-2 seronegative women (Veselenak et al., 2012). The vaccine was effective in preventing HSV-1 genital disease but not in preventing HSV-2 infection. Overall, the vaccine was not efficacious (20%) (Veselenak et al., 2012). If a vaccine were found to give type-common protection against both HSV-1 and HSV-2, early childhood vaccination would have been optimal (Koelle et al., 2003).

However, Conventional vaccines development by either attenuated or inactivated whole pathogen witnessed a number of limitations (Rappuoli, 2000; Purcell et al., 2007). First, genetic variations of pathogens around the globemay results in reduced efficiency of these vaccines in distinctive parts of the world. Second, enormous vaccine trials failed to reach in phase III (Sabchareon et al., 2012; Gray et al., 2011; Tameris et al., 2013). Third, experimental assay based identification of conserved regions that maintain their structure and function of glycoprotein is a tedious process (Rappuoli, 2000). Such shorts of limitation apparently indicates the better understanding of host pathogen interaction, focusing prior *in-silico* study before lab based preparation of vaccines (Mora et al., 2003; Rappuoli et al., 2016). Immunoinformatics, an emerging field of the present era addressed the complex biological shortcomings of decrypting the immune response for vaccine designing and development (Nielsen et al., 2007). For completely eradicating the chance of re-infection, an ideal vaccine has to initiate humoral or cell mediated immunity. The Cytotoxic T lymphocytes (CTL) and Helper T lymphocytes (HTL) recognize the foreign antigen that are presented by Major Histocompatibility Complex (MHC) and expressed on the surface of all nucleated cells. T-cell epitope prediction tools help to identify allele specific peptides. Hence, it reduces the number of potential peptides to be considered as vaccine candidates. *In silico* epitope predictions tools have already proved advantages in determining the potential candidates reducing the number of validation experiments and time (Korber et al., 2006; Nielsen et al., 2007). In recent times, enormous computational tools are available for predicting peptides (T and B cell) with necessary properties for vaccine design (Yang & Yu 2009). On top of that, several works were done by the researcher working on *in silico* vaccine design of several deadly and worrisome viruses including Nipah (Saha et al., 2017), Ebola (Dash et al., 2017), HIV (Saxena et al., 2013; Yang et al., 2015) etc. But still now, no attempt has been taken for determining *in silico* vaccine target against both types of HSV. Therefore, the present research will be a promising study for the computer aided design of epitope based vaccine target against Type-1 and Type-2 HSV in together. Meanwhile, the study could be useful for laboratory based development of effective multiepitope polyvalent vaccine as well as further development of existing experimental subunit vaccines of HSV.

## Materials and methods

In the present study, an immunoinformatic approach was adopted for predicting vaccine candidates against Type-1 and Type-2 Herpes Simplex Virus (HSV). The flow chart summarizing the protocol over *in silico* vaccinomics strategy for designing a unique multiepitope subunit vaccine has been illustrated in figure 1.

**Figure 1.**
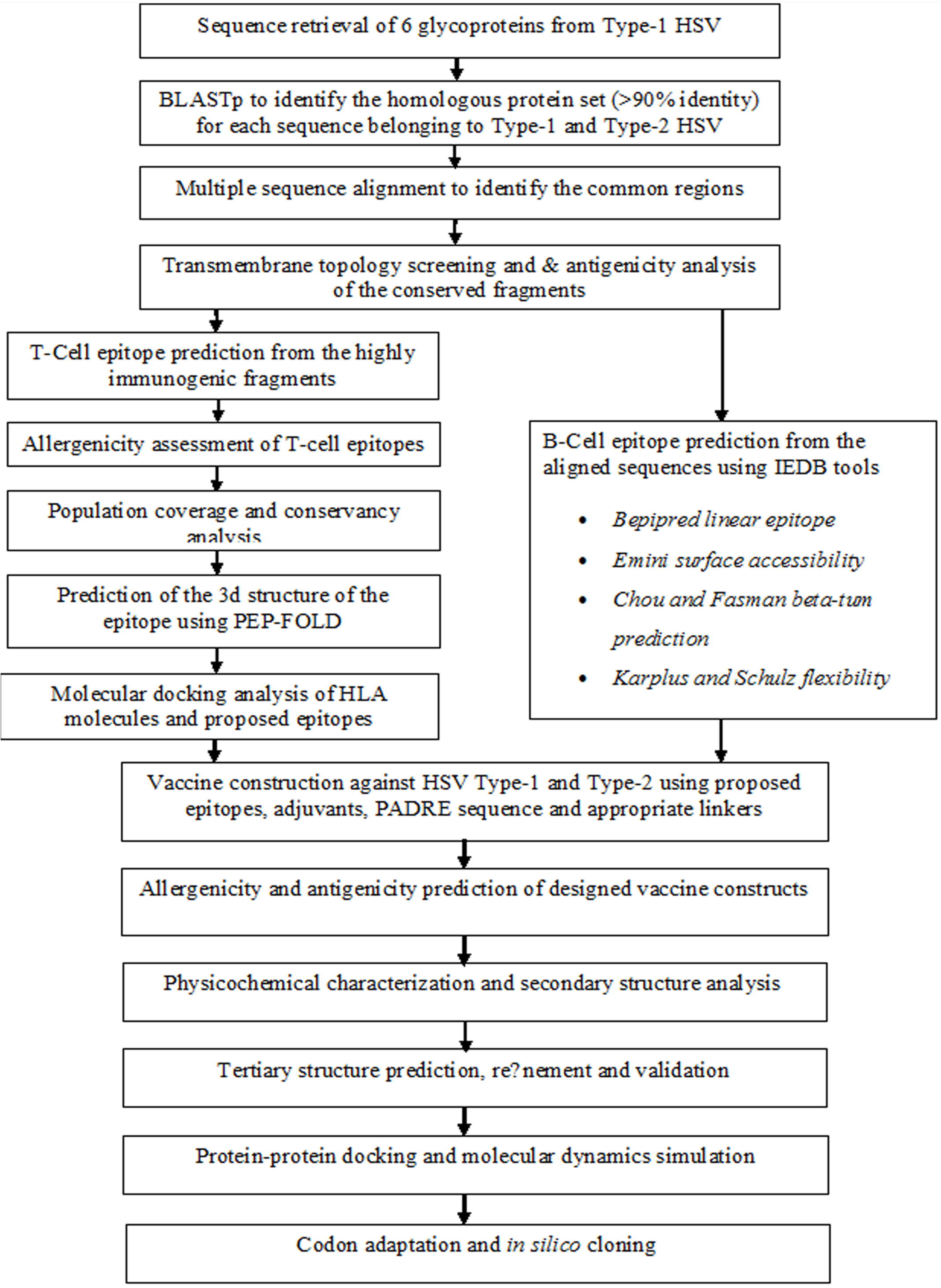
Flow chart summarizing the protocols for the prediction of epitope based vaccine candidate by *in silico* reverse vaccinology technique.

### Selection of Viral strain and retrieval of protein sequences

The National Center for Biotechnology Information (NCBI) provides access to biomedical and genomic information over many organisms. The server was used for the selection of Type-1 Herpes Simplex Virus (HSV) (https://www.ncbi.nlm.nih.gov/genome/). Study of other associated information including the genus, family, host, transmission, disease, genome and proteome analysis were performed by using ViralZone, a web-resource of Swiss Institute of Bioinformatics (https://viralzone.expasy.org/). A total 6 glycoproteins were considered as vaccine targets for *in silico* analysis and vaccine development against HSV.

### Retrieval of homologous protein sets and multiple sequence alignment

Homologous sequences of the selected glycoproteins were retrieved from the NCBI database by using BLASTp tool. All 6 glycoproteins were used as query and the searches were restricted to Type-1 HSV (Taxid: 10298) and Type-2 HSV (Taxid: 10310). Multiple sequence alignment (MSA) was further performed to find out the common regions for both sets of proteins using using MEGA 5.05 software package (http://www.megasoftware.net). The CLUSTALW algorithm along with 1000 bootstrap value and other default parameters were used to fabricate the alignment. The sequences were investigated in order to recognize the immunologically pertinent regions that were achieved by predicting epitopic peptides.

### Prediction of antigenicity and transmembrane properties of the conserved sequences

Antigenicity refers to the capacity of antigens to be recognized by the by the antibodies or immune system. The conserved amino acid sequences from each of the 6 proteins were screened for predicting their antigenicity using an online antigen prediction server, VaxiJen v2.0 (Doytchinova & Flower, 2007). Each selected conserved fragments were subjected to transmembrane topology prophecy using TMHMM v0.2 server (Krogh et al., 2001) with a view to discriminate intracellular and surface proteins with high degree of accuracy.

### Prediction of T Cell Epitopes from the Conserved Regions

Both MHC-I restricted CD8+ cytotoxic T lymphocytes (CTLs) and MHC-II restricted CD4+ cytotoxic T lymphocytes play a pivotal role in controlling viral infections. Hence, identification of T cell epitopes is crucial for understanding the mechanism of T cell activation and epitope driven vaccine design. The IEDB offers easy searching of experimental data characterizing antibody and T cell epitopes studied in human and other non-human primates. From this Immune Epitope Database, MHC-I prediction tool (http://tools.iedb.org/mhci/) and MHC-II prediction tool (http://tools.iedb.org/mhcii/) were used to predict the MHC-I binding and MHC-II binding respectively (Vita et al., 2014).

### Transmembrane topology prediction and antigenicity analysis of epitopes

The TMHMM server (http://www.cbs.dtu.dk/services/TMHMM/) predicted the transmembrane helices in proteins. The topology was determined according to the position of the transmembrane helices separated by ‘i’ if the loop is on the inside or ‘o’ if it is on the outside (Krogh et al., 2001). Again, VaxiJen v2.0 server (http://www.ddg-pharmfac.net/vaxijen/) was used to determine the epitope antigenicity (Doytchinova and Flower, 2007). The most potent antigenic epitopes were selected for further investigation.

### Population coverage analysis

HLA distribution varies among different ethnic groups and geographic regions around the world. In this study, population coverage for each individual epitope was analyzed by the IEDB population coverage calculation tool analysis resource (http://tools.iedb.org/population/).

### Allergenicity assessment and toxicity analysis of T-Cell epitopes

The prediction of allergens has been explored widely using bioinformatics, with many tools being developed in the last decade. Four different tools, i.e.AllerTOP (http://www.ddg-pharmfac.net/AllerTop/) (Dimitrov et al., 2013), AllergenFP (http://www.ddg-pharmfac.net/AllergenFP/) (Dimitrov et al., 2014), PA3P (http://www.Ipa.saogabriel.unipampa.edu.br:8080/pa3p/) (Chrysostomou & Seker, 2014) and Allermatch (http://www.allermatch.org/allermatch.py/form) (Fiers et al., 2004) were used to predict the allergenicity of our proposed epitopes for vaccine development. Only the non-allergenic epitopes were allowed to demonstrate the toxicity level by ToxinPred server (http://crdd.osdd.net/raghava/toxinpred/).

### Conservancy analysis

Epitope conservancy is a vital step in the immunoinformatic approach as it determines the extent of desired epitope distributions in the homologous protein set. IEDB’s epitope conservancy analysis tool (http://tools.iedb.org/conservancy/) was selected for the analysis of conservancy level by concentrating on the identities of the selected proteins.

### Designing three-dimensional (3D) epitope structure

The top ranked epitopes were subjected for the docking study after analyzing through different bioinformatics tools. PEP-FOLD is a *de novo* approach aimed at predicting peptide structures from amino acid sequences (Maupetit et al., 2010). By Folding peptides on a user specified patch of a protein, it comes with the possibility to generate candidate conformations of peptide-protein complexes (Kaur et al., 2007). The server was used to design and retrieve the 3D structure of most potent selected epitopes for further analysis.

### Molecular docking analysis

MGLTools is a software developed for the visualization and analysis of molecular structures (Sanner, 1999). It includes AutoDock, an automated docking software designed to predict the interactions between small molecules (i.e. substrates or drug candidates) and receptor of known 3D structure (Morris et al., 2009). All the operations were performed at 1.00°A space keeping the exhaustiveness parameter at 8.00. The numbers of outputs were set at 10. The docking was conducted using AutoDOCKVina program based on the above-mentioned parameters. OpenBabel (version 2.3.1) was used to convert the output PDBQT files in PDB format. The best output was selected on the basis of higher binding energy. The docking interaction was visualized with the PyMOL molecular graphics system, version 1.5.0.4 (https://www.pymol.org/).

### B-Cell epitope identification

B cell epitope prediction tools from IEDB were used to identify the B cell antigenicity based on six different algorithms which include Emini surface accessibility prediction (Emini et al., 1985), Karplus and Schulz flexibility prediction (Karplus & Schulz, 1985), Bepipred linear epitope prediction analysis (Jespersen et al., 2017), Chou and Fasman beta turn prediction (Chou & Fasman, 1978), Kolaskar & Tongaonkar antigenicity scale (Kolaskar & Tongaonkar, 1990) and Parker hydrophilicity prediction (Parker et al., 1986).

### Vaccine construction

Subunit vaccines consist of antigenic parts of a pathogen to stimulate an immunogenic reaction in the host. The predicted T-cell and B-cell epitopes were conjugated in a sequential manner to design the final vaccine construct. All three vaccine proteins started with an adjuvant followed by the top CTL epitopes for both capsid protein VP1 and protein VP2, then by top HTL epitopes and BCL epitopes respectively, in the similar fashion. Three vaccine sequence were constructed named V1, V2 and V3, each associated with different adjuvants including beta defensin (a 45 mer peptide), L7/L12 ribosomal protein and HABA protein (*M. tuberculosis*, accession number: AGV15514.1). Interactions of adjuvants with toll like receptors (TLRs) polarize CTL responses and induce robust immunoreactions (Rana and Akhter, 2016). Beta defensin adjuvant can act as an agonist to TLR1, TLR2 and TLR4 receptor. On the contrary, L7/L12 ribosomal protein and HBHA protein are agonists to TLR4 only. To overcome the problems caused by highly polymorphic HLA alleles, PADRE sequence was also incorporated along with the adjuvant peptides. EAAAK linkers were used to join the adjuvant and CTL epitopes. Similarly, GGGS, GPGPG and KK linkers were used to conjugate the CTL, HTL and BCL epitopes respectively. Utilized linkers ensured the effective separation of individual epitopes in vivo (Hajighahramani et al., 2017; Pandey et al., 2016).

### Allergenicity and antigenicity prediction of different vaccine constructs

AlgPred v.2.0 (Saha and Raghava, 2000) sever was used to predict the non-allergic nature of the constructed vaccines. The server developed an algorithm by considering the auto cross covariance transformation of proteins into uniform vectors of similar length. The accuracy of results ranges from 70% to 89% depending on species. We further used VaxiJen v2.0 server (Doytchinova and Flower, 2007) to evaluate the probable antigenicity of the vaccine constructs in order to suggest the superior vaccine candidate. The server analyzed the immunogenic potential of the given proteins through an alignment-independent algorithm.

### Physicochemical characterization and secondary structure analysis of vaccine protein

ProtParam, a tool provided by Expasy server (Gasteiger et al., 2003; Hasan et al., 2015a; Das et al., 2016) was used to functionally characterize the vaccine constructs according to molecular weight, aliphatic index, isoelectric pH, hydropathicity, instability index, GRAVY values, estimated half-life and various physicochemical properties. By comparing the pK values of different amino acids, the server computes these parameters of a given protein sequence. Aliphatic index is the volume occupied by the aliphatic side chains of protein. Grand average of hydropathicity was computed by summing the hydropathicity of all amino acid residues present in the protein sequenceand then by dividing it by total number of amino acid residues. The PSIPRED v3.3 (Kosciolek & Jones, 2014) and NetTurnP 1.0 program (Petersen et al., 2010; Thaysen-Andersen & Packer, 2012) was used to predict the alpha helix, beta sheet and coil structure of the vaccine constructs.

### Vaccine tertiary structure prediction, refinement and validation

The RaptorX server performed 3D modeling of the designed vaccines depending on the degree of similarity between target protein and available template structure from PDB (Kallberg et al., 2014; Peng and Xu, 2011). Refinement was conducted using ModRefiner (Xu and Zhang, 2011) followed by FG-MD refinement server (Zhang et al., 2011) to improve the accuracy of the predicted 3D modeled structure. ModRefiner drew the initial model closer to its native state based on hydrogen bonds, side-chain positioning and backbone topology, thus resulting in significant improvement in the physical quality of the local structure. FG-MD is another molecular dynamics based algorithm for structure refinement at atomic level. The refined protein structure was further validated by Ramachandran plot assessment at RAMPAGE (Lovell et al., 2002; Hasan et al., 2015b).

### Vaccine protein disulfide engineering

Disulfide bonds enhance the geometric conformation of proteins and provide significant stability. DbD2, an online tool was used to design such bonds for the constructed vaccine protein (Craig & Dombkowski, 2013). The server detects and provides a list of residue pairs with proper geometry which have the capacity to form disulfide bond when individual amino acids are mutated to cysteine.

### Protein-protein docking

Molecular docking aims to determine the binding affinity between a receptor molecule and ligand (Solanki & Tiwari, 2018). An approach for protein-protein docking was employed to determine the binding affinity of designed subunit vaccines with different HLA alleles and TLR-2 immune receptor by using ClusPro 2.0 (Comeau et al., 2004), hdoc (Macalino et al., 2018; Kangueane & Nilofer, 2018) and PatchDock server (Schneidman-Duhovny et al., 2005). During HSV infection, viral glycoproteins on the envelope serve as PAMPs for TLR2 and act as a ligand (Leoni et al., 2012; Sato et al., 2006). The 3D structure of different MHC molecules and human TLR-2 receptor was retrieved from RCSB protein data bank. The above mentioned servers were used to obtain the desirable complexes in terms of better electrostatic interaction and free binding energy. PatchDock generated a number of solutions which were again subjected to the FireDock server to refine the complexes.

### Molecular dynamics simulation

Molecular dynamics study is important to strengthen any *in silico* prediction and demonstrate the stability of protein-protein complex. Stability can be determined by comparing the essential dynamics of proteins to their normal modes (Aalten et al., 1997; Wuthrich et al., 1980). This powerful tool is an alternative to the costly atomistic simulation (Tama & Brooks, 2006; Cui & Bahar, 2007). iMODS server explains the collective motion of proteins by analyzing the normal modes (NMA) in internal coordinates (Lopez-Blanco et al., 2017). The structural dynamics of protein complex was investigated by using this server due to its much faster and effective assessments than other molecular dynamics (MD) simulations tools (Awan et al., 2017; Prabhakar et al., 2016). It predicted the direction and extent of the immanent motions of the complex in terms of deformability, eigenvalues, B-factors and covariance. The deformability of the main chain depends on the ability to deform at each of its residues for a given molecule. The eigenvalue related to each normal mode describes the motion stiffness. This value is directly linked to the energy required to deform the structure. Deformation is much easier if the eigenvalue is low and vice versa (Lopez-Blanco et al., 2011).

### Codon adaptation and in silico cloning

Codon adaptation tools are used for adapting the codon usage to the well characterized prokaryotic organisms to accelerate the expression rate in them. *E. coli* strain K12 was selected as host for cloning purpose of the designed vaccine construct. Due to the lack of similarities between the codon usage of human and *E. coli*, the approach was adopted to achieve higher expression of vaccine protein V1 in the selected host. Rho independent transcription termination, prokaryote ribosome-binding site and cleavage sites of several restriction enzymes (i.e. XhoI and EcoRI) were avoided during the operation performed by JCAT server (Grote et al., 2005). The optimized sequence of vaccine protein V1 was reversed and then conjugated with XhoI and EcoRI restriction site at the N-terminal and C-terminal sites respectively. SnapGene (Solanki & Tiwari, 2018) restriction cloning module was used to insert the adapted sequence between XhoI (158) and EcoRI (192) of pET28a(+) vector.

## Results

### Retrieval of Protein Sequences

The sequences of 6 glycoproteins i.e. GP B39, GP C, GP D, GP E, GP G and envelope protein UL45 of Type-1 Herpes Simplex virus were retrieved from NCBI protein database(http://www.ncbi.nlm.nih.gov/protein) in FASTA format. Again, 6 sets of homologous proteins for each glycoproteins were generated after BLASTp search using NCBI BLAST tools (Supplementary file 1-6). Each set comprisedprotein sequences belonging to different isolates of Type-1 and Type-2 HSV.

### Identification and Selection of Conserved Sequences

A total of 27, 17, 14, 21, 15 and 7 conserved sequences were found inGP B39, GP C, GP D, GP E, GP G and envelope protein UL45, respectively. Conserved sequences generated in this method have been presented in supplementary file 7-12.

### Antigenicity and Transmembrane Properties of the Conserved Sequences

Results showed that 20, 10, 9, 10, 11 and 2conserved sequencesfrom GP B39, GP C, GP D, GP E, GP G and GP UL45 respectively met the criteria of default threshold level in VaxiJen (Table 1). Moreover, transmembrane topology screening revealed, among the immunogenic conserved sequences8, 2, 5, 3, 8 and 2 sequences from the corresponding proteins fulfilled the criteria of exomembrane characteristics (Table 1).

**Table 1.**
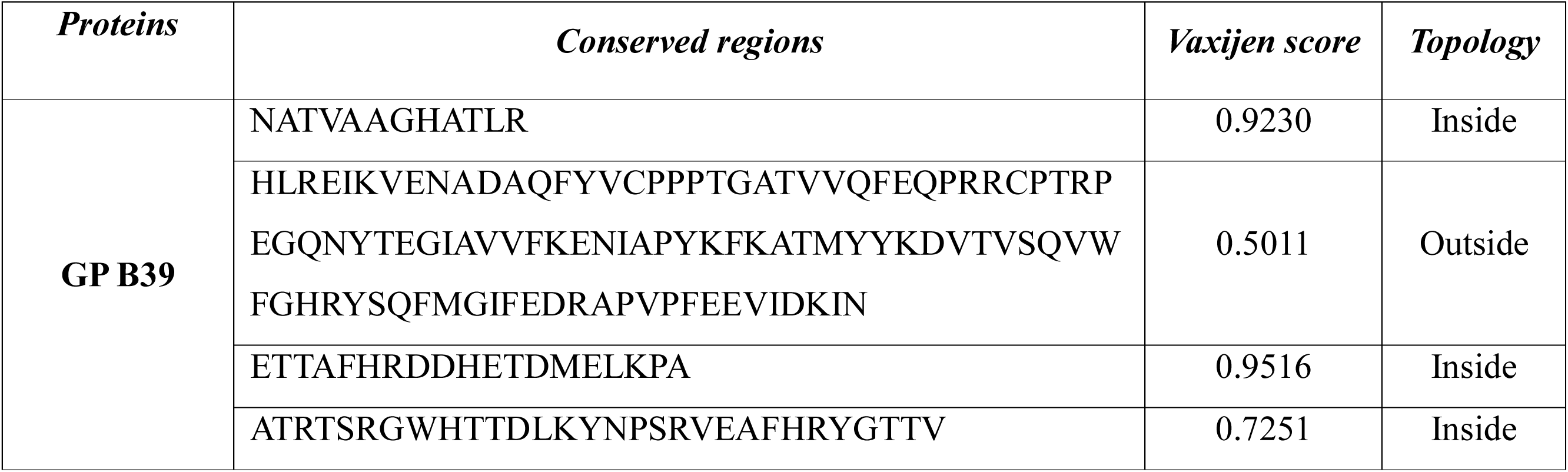
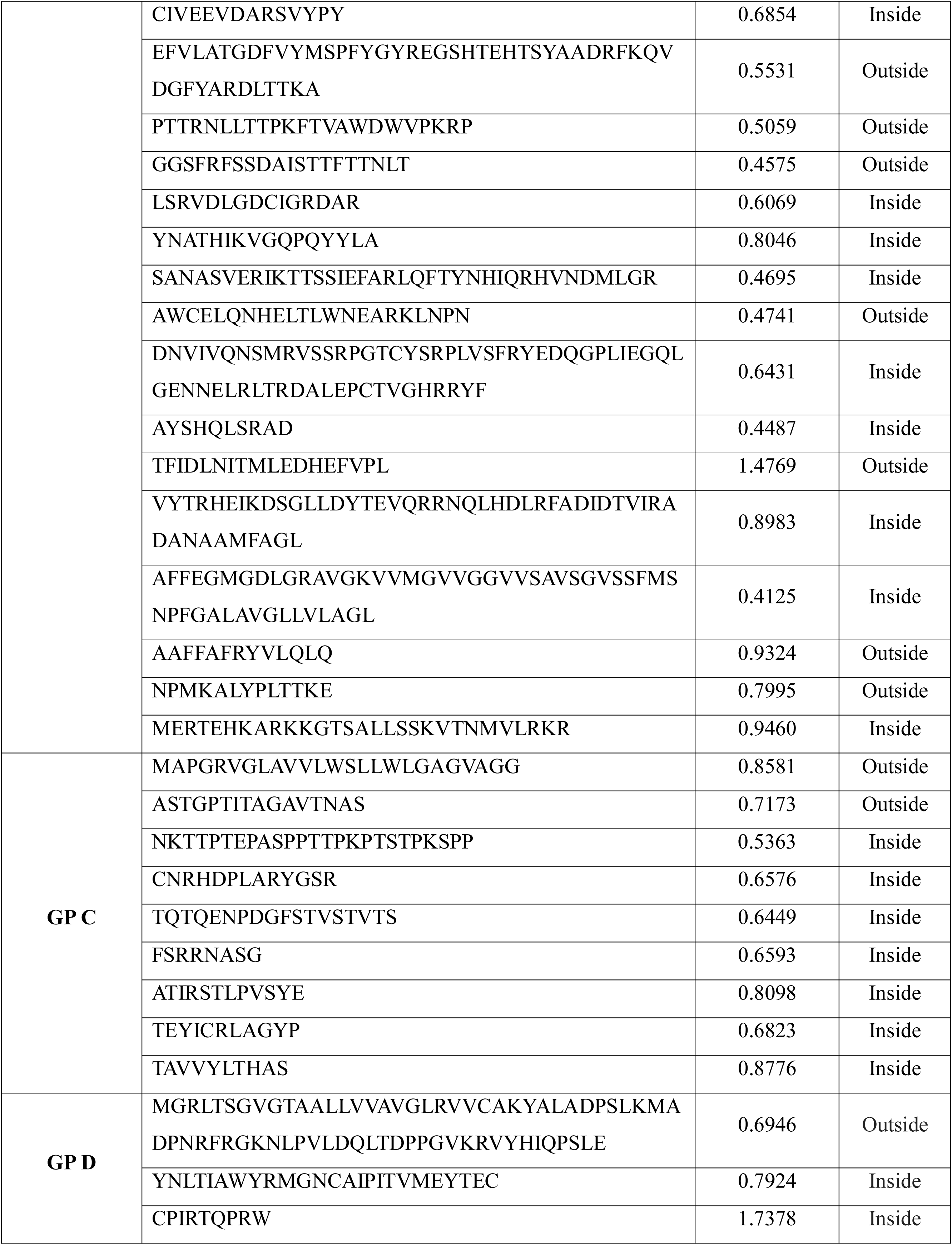
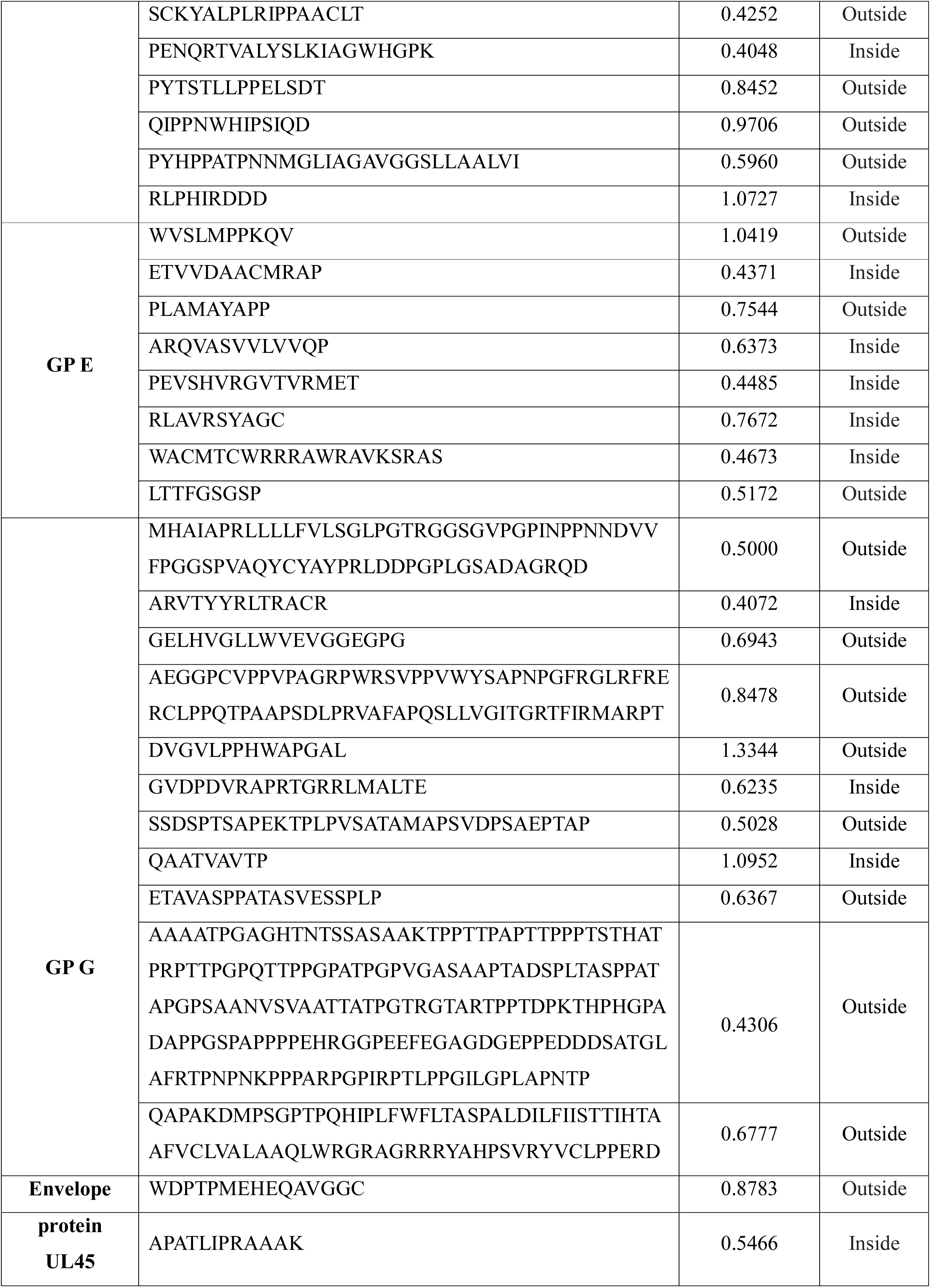
Highly antigenic conserved regions identified among different sets of HSV proteins.

### T-Cell epitope prediction

Numerous immunogenic epitopes from the selected proteins were identified to be potent T cell epitopes using both MHC-I and MHC-II binding predictions of IEDB. Epitopes that bind to the maximum number of HLA cells with high binding affinity were selected (Table 2 and Table 3).

**Table 2.** Predicted CTL epitopes among HSV proteins to construct multiepitope polyvalent vaccine.

| *Proteins* | *Top CTL Epitopes* | *Interacting MHC I Molecules* | *Allergenicity* | *VaxiJen Score* |
| --- | --- | --- | --- | --- |
| GP B39 | GLAAFFAFR | HLA-A*31:01, HLA-A*68:01, HLA-A*33:01,<br>HLA-A*11:01, HLA-A*03:01 | Non Allergen | 0.8634 |
| GP C | SLLWLGAGV | HLA-A*02:01, HLA-A*02:03, HLA-A*02:06 | Non Allergen | 1.1557 |
| GP D | LLAALVIRL | HLA-A*02:03, HLA-A*02:01, HLA-A*02:06 | Non Allergen | 0.8730 |
| GP E | KQVPLAMAY | HLA-B*15:01, HLA-A*02:06, HLA-A*30:02 | Non Allergen | 0.9020 |
| GP G | HTAAFVCLV | HLA-A*02:06, HLA-A*68:02, HLA-A*02:03 | Non Allergen | 1.1073 |
| Envelope Protein UL45 | HEQAVGGCW | HLA-A*11:01, HLA-B*44:02, HLA-B*44:03 | Non Allergen | 0.6080 |

**Table 3.** Predicted HTL epitopes among HSV proteins to be a part of multiepitope polyvalent vaccine.

| *Proteins* | *Top HTL Epitopes* | *Interacting MHC II Molecules* | *Allergenicity* | *VaxiJen Score* |
| --- | --- | --- | --- | --- |
| GP B39 | NPFGALAVGLLVLAG | HLA-DRB1*01:01, HLA-DQA1*05:01,<br>HLA-DRB1*09:01,<br>HLA-DPA1*03:01/DPB1*04:02 | Non Allergen | 0.8490 |
| GP C | RVGLAVVLWSLLWLG | HLA-DPA1*03:01/DPB1*04:02,<br>HLA-DPA1*02:01/DPB1*01:01,<br>HLA-DPA1*01:03/DPB1*02:01,<br>HLA-DRB1*01:01, HLA-DRB4*01:01,<br>HLA-DRB1*15:01, HLA-DRB1*07:01,<br>HLA-DRB1*04:05 | Non Allergen | 0.7390 |
| GP D | AALLVVAVGLRVVCA | HLA-DRB1*07:01, HLA-DRB1*01:01,<br>HLA-DRB5*01:01, HLA-DRB1*09:01,<br>HLA-DRB1*13:02, HLA-DQA1*05:01, | Non Allergen | 1.2353 |
|  |  | HLA- DQB1*06:0, HLA-DRB1*15:01 |  |  |
| GP E | No potential fragments found with 15 residues | - | - | - |
| GP G | ALDILFIISTTIHTA | HLA-DRB1*07:01, HLA-DRB1*01:01, HLA-DRB1*15:01, HLA-DRB1*04:01, HLA-DRB1*13:02, HLA-DRB1*11:01, HLA-DRB1*04:05, HLA-DRB5*01:01, HLA-DQA1*01:02/DQB1*06:02, HLA-DRB4*01:01, HLA-DPA1*02:01 | Non Allergen | 0.8169 |
| Envelope Protein UL45 | HLAALARVQAERSSG | HLA-DRB1*01:01, HLA-DQA1*03:01, HLA-DRB3*01:01, HLA-DRB1*08:02, HLA-DRB1*04:01, HLA-DRB1*11:01 | Non Allergen | 0.5567 |

### Transmembrane topology prediction and antigenicity analysis

Top epitopes from both proteins were selected as putative T cell epitope candidates based on their transmembrane topology screening and antigenic scoring (Table 2 and Table 3). Epitopes with a positive score of immunogenicity exhibited potential to elicit effective T-cell response.

### Allergenicity assessment and toxicity analysis of T-Cell epitopes

Based on the allergenicity assessment by four servers (i.e. AllerTOP, AllergenFP, PA^3^P, Allermatch), epitopes that were found to be non-allergen for human were identified (Table 2 and Table 3). Epitopes those were indicated as allergen for human and classified as toxic or undefined were removed from the predicted list of epitopes.

### Population coverage analysis

All indicated alleles in supplementary data were identified as optimum binders with the predicted epitopes and were used to determine the population coverage. Population coverage results for six glucoproteins are shown in figure 2.

**Figure 2.**
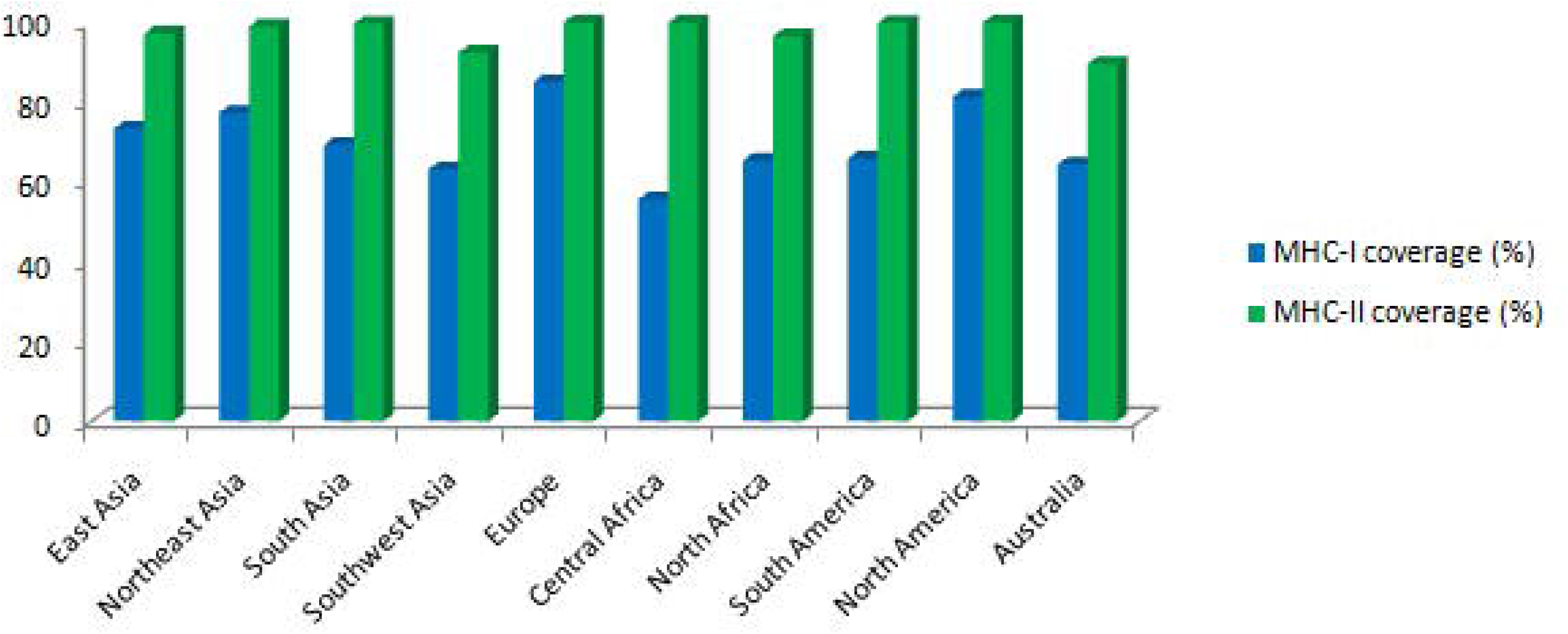
Population coverage analysis of HSV glycoproteins.

### Conservancy analysis

Putative epitopes generated from 6 glycoproteins were found to be highly conserved with conservancy level ranging from 90% to 100%. The top epitopes showing conservancy at a superior level were allowed for further docking study and used to construct the final vaccine candidates to ensure a broad spectrum vaccine efficacy.

### Molecular docking analysis and HLA allele interaction

11 T-cell epitopes were subjected to PEP-FOLD3 web-based server for 3D structure conversion in order to analyze their interactions with HLA molecules. On the basis of available Protein Data Bank (PDB) structures deposited in the database, HLA-A*11:01, HLA-A*02:06 and HLA-DRB1*01:01was selected for docking analysis and the binding energies were analyzed (Table 4). Results showed that all the predicted epitopes bound in the groove of MHCmolecules with a negative bindingenergy which were biologically significantly (Table 4, Figure 3 and Figure 4).

**Figure 3.**
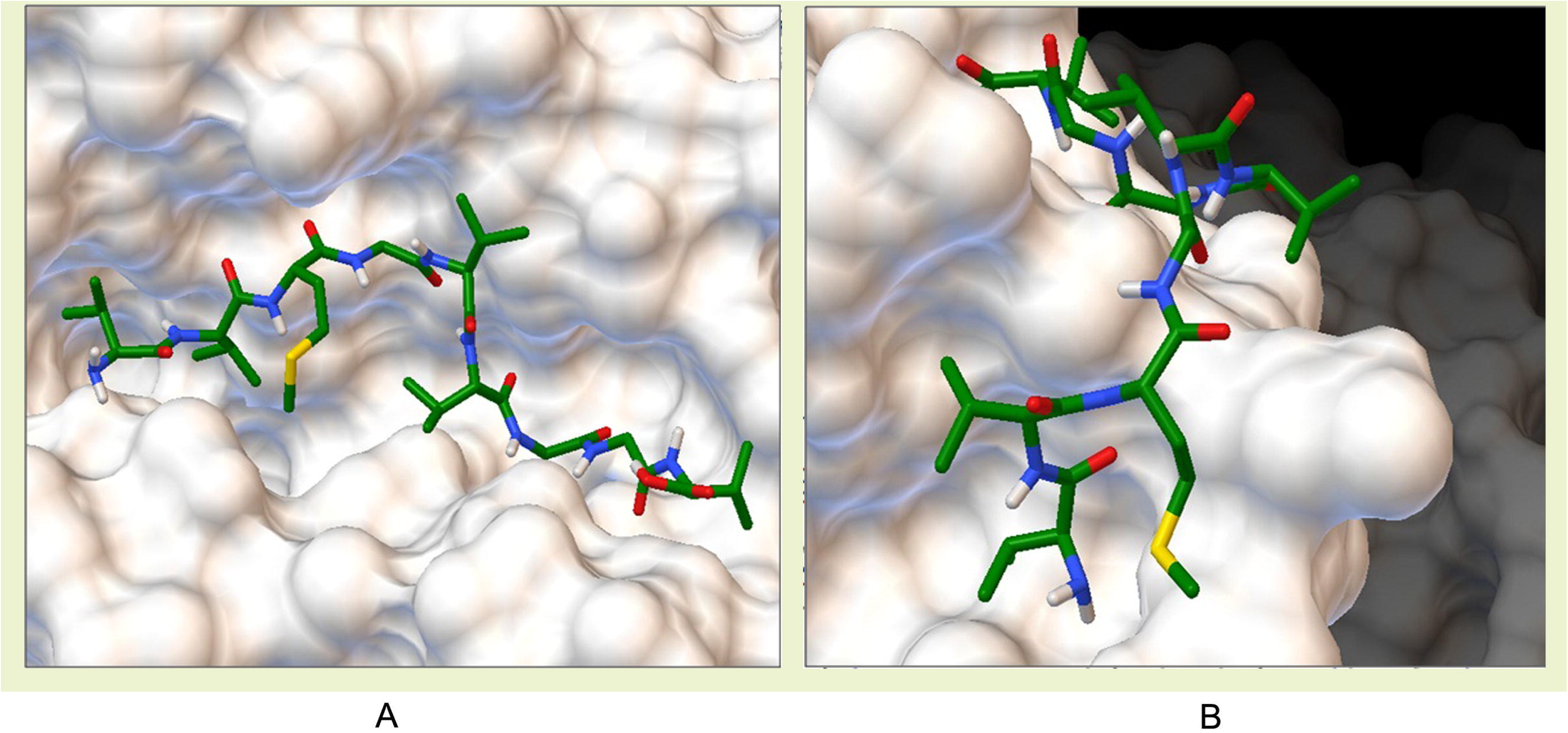
Docking of predicted GP B39 epitope ‘GLAAFFAFR’ to HLA-A*11:01 and GP G epitope ‘HTAAFVCLV’ to HLA-A*02:06 with a binding energy of −6.4 Kcal/mol and −7.0 Kcal/mol, respectively.

**Figure 4.**
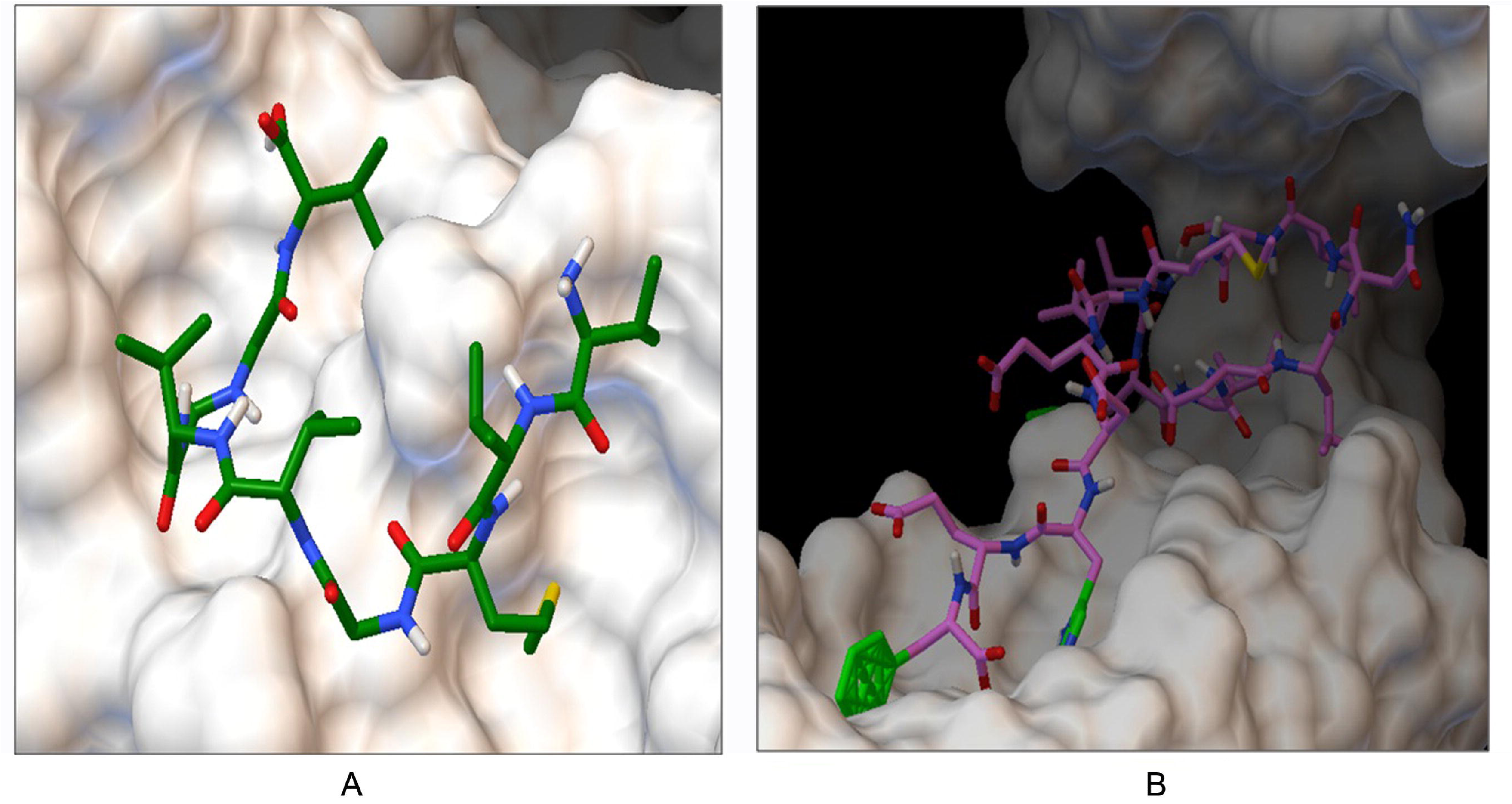
Docking of predicted GP C epitope ‘RVGLAVVLWSLLWLG’ and GP D epitope ‘AALLVVAVGLRVVCA’ to HLA-DRB1*01:01 with a binding energy of −8.3 Kcal/mol and −6.6 Kcal/mol, respectively.

**Table 4.** Binding energy of suggested T-cell epitopes with selected MHC class I and class-II molecules generated from molecular docking analysis.

| *Proteins* | *Epitopes* | *MHC alleles* | *Binding energy (kcal/mol)* | *Epitopes* | *MHC alleles* | *Binding energy (kcal/mol)* |
| --- | --- | --- | --- | --- | --- | --- |
| GP B39 | GLAAFFAFR | HLA-A*11:01 | -6.4 | NPFGALAVGLLVLAG | HLA-DRB1*01:01 | -5.3 |
| GP C | SLLWLGAGV | HLA-A*02:06 | -5.6 | RVGLAVVLWSLLWLG |  | -8.3 |
| GP D | LLAALVIRL |  | -5.9 | AALLVVAVGLRVVCA |  | -6.6 |
| GP E | KQVPLAMAY |  | -3.0 | - |  | - |
| GP G | HTAAFVCLV |  | -7.0 | ALDILFIISTTIHTA |  | -6.2 |
| Envelope Protein UL45 | HEQAVGGCW | HLA-A*11:01 | -6.2 | HLAALARVQAERSSG |  | -5.6 |

### B-Cell epitope identification

B cell epitopes were predicted according to the analysis via four different algorithms from IEDB. Top BCL epitopes for selected six glycoproteins were further screened based on their antigenicity scoring and allergenicity pattern.The top BCL epitope for each protein was used to design the final vaccine construct (Table 5).

**Table 5.** Predicted B cell epitopes among HSV proteins to design multiepitope polyvalent vaccine.

| Proteins | Top BCL Epitopes | Allergenicity | Antigenicity (VaxiJen Score) |
| --- | --- | --- | --- |
| GP B39 | PTGATVVQFE | Non Allergen | 1.19 |
| GP C | MAPGRVGLAV | Non Allergen | 1.42 |
| GP D | VKRVYHIQPS | Non Allergen | 1.17 |
| GP E | PKQVPLAMAY | Non Allergen | 1.34 |
| GP G | AFAPQSLLVG | Non Allergen | 1.13 |
| GP UL45 | WDPTPMEHEQ | Non Allergen | 1.64 |

### Vaccine construction

Each of the constructs consisted of a protein adjuvant followed by PADRE peptide sequence, while the rest was occupied by the T-cell and B-cell epitopes and their respective linkers. PADRE sequence was incorporated to maximize the efficacy and potency of the peptide vaccine. All three designed vaccines comprised 6 CTL epitopes, 6 HTL epitopes and 8 BCL epitopes combined together in a sequential manner. A total 3 vaccines of 403 (V1), 488 (V2) and 517 (V3) amino acid long were constructed (Table 6) and further analyzed to investigate their immunogenic potential.

**Table 6.**
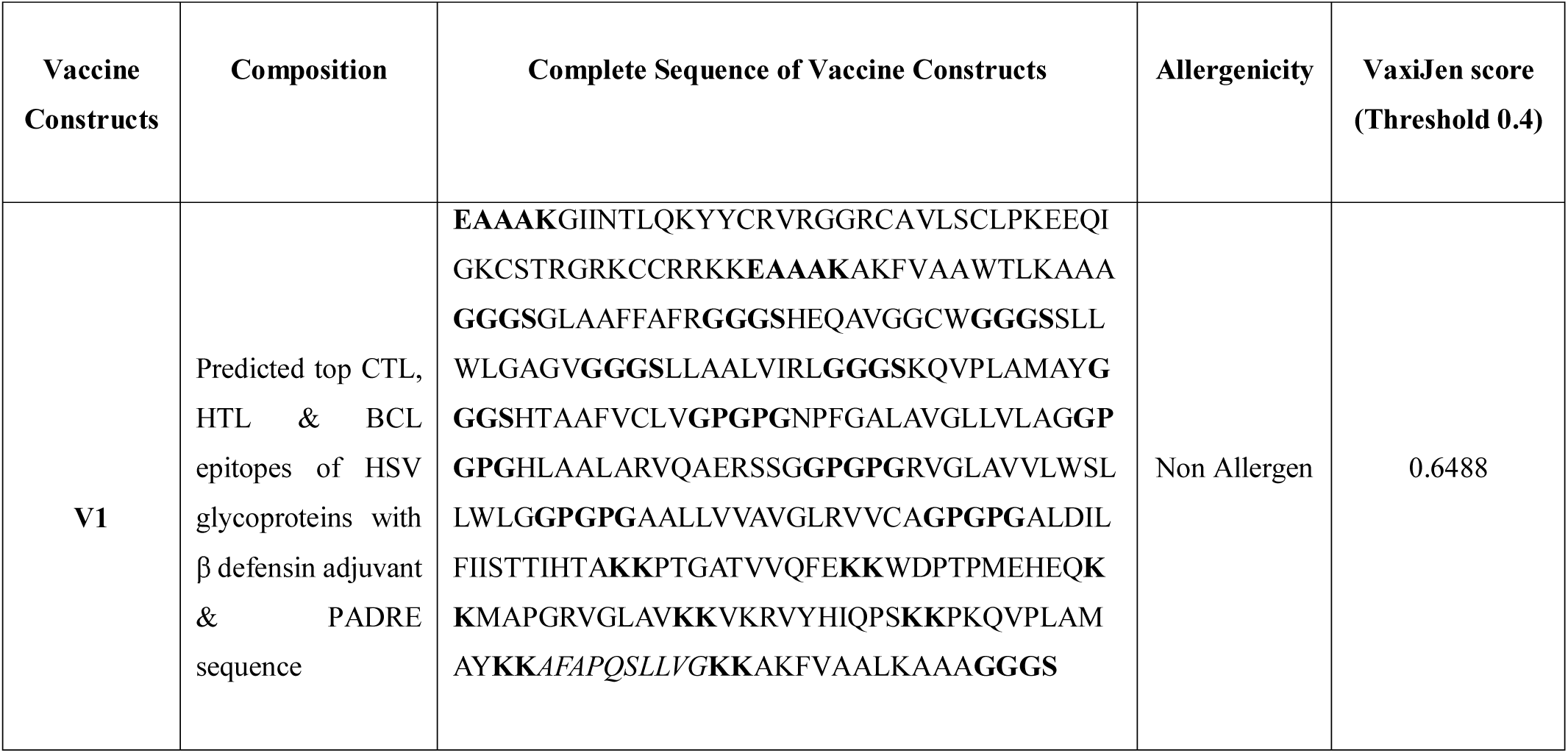
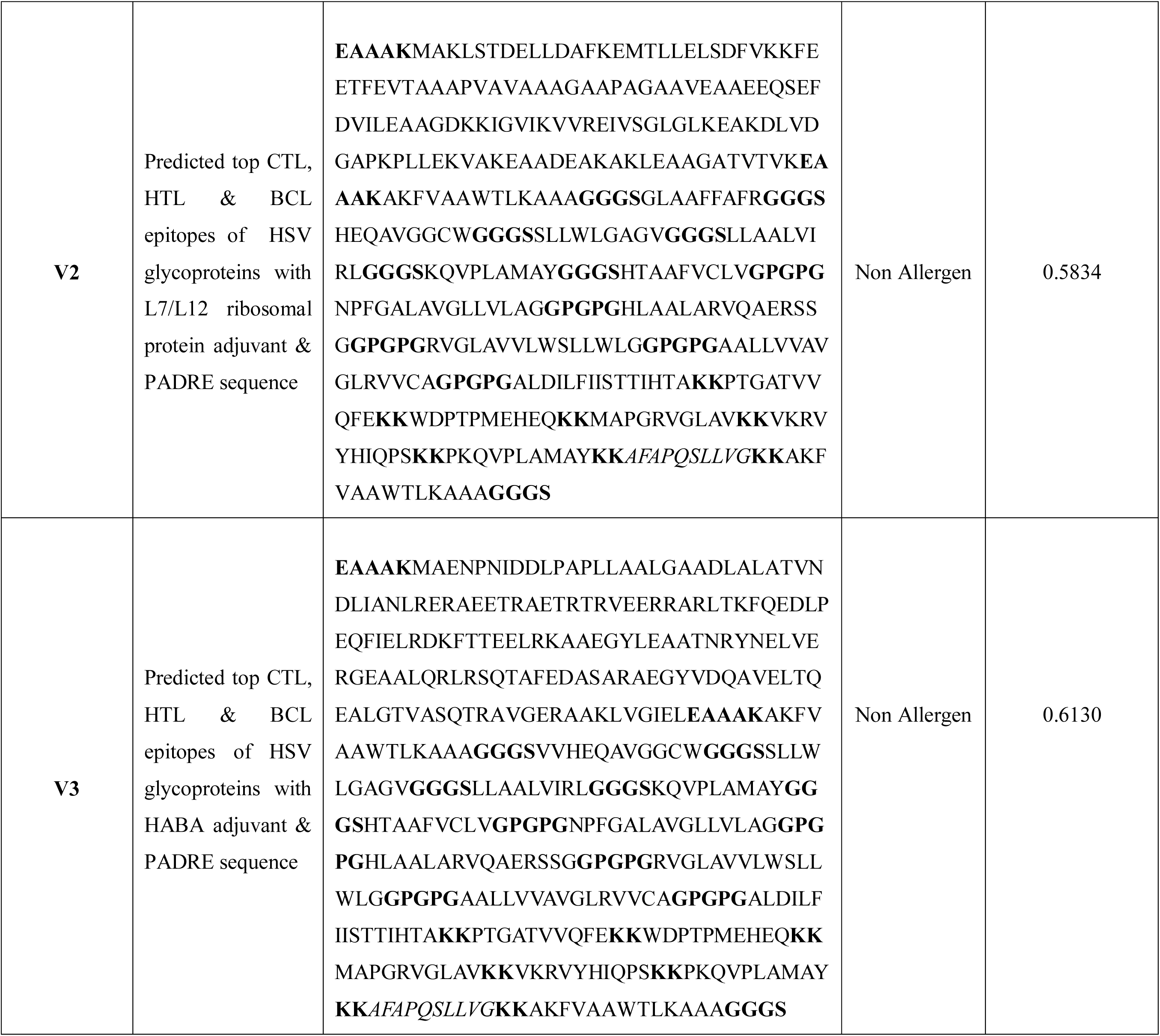
Allergenicity and antigenicity analysis of the constructed vaccines.

### Allergenicity and antigenicity prediction of different vaccine constructs

Results showed that all three constructs (V1, V2 and V3) were non-allergic in behavior. However, V1 was best in terms of safety and found superior as potential vaccine candidate with better antigenicity (0.772) and ability to stimulate preferred immune response (Table 6).

### Physicochemical characterization of vaccine protein

The final vaccine construct was characterized on the basis of physical and chemical properties. The molecular weight of the vaccine construct V1 was 33.83kDa which ensured its good antigenic potential. The theoretical pI 10.33 indicated that the protein will have net negative charge above the pI and vice versa. The extinction coefficient was 40450, assuming all cysteine residues are reduced at 0.1% absorption. The estimated half-life of the constructed vaccine was expected to be 1h in mammalian reticulocytes in vitro while more than 10 h in *E. coli* in vivo. Thermostability and hydrophilic nature of the vaccine protein was represented by aliphatic index and GRAVY value which were 91.55 and 0.229 respectively. The computed instability index of the protein was 34.91 which classified it as a stable one.

### Secondary and tertiary structure prediction

Secondary structure of the construct V1 confirmed to have 30.44% alpha helix, 21.79% sheet and 47.77% coil structure (Fig. 6). RaptorX generated the tertiary structure of the designed construct V1 consisting single domain (Figure 5). Homologymodeling was performed by detecting and using 1vh4A from protein data bank (PDB) as best suited template for Vaccine V1. All 335 amino acid residues (100%) were modeled with only 15% residues in the disordered region. The quality of the 3D model was defined by P value which was 2.12e^-03^ for the predicted vaccine protein. The low P value ensured better model quality of the predicted vaccine.

**Figure 5.**
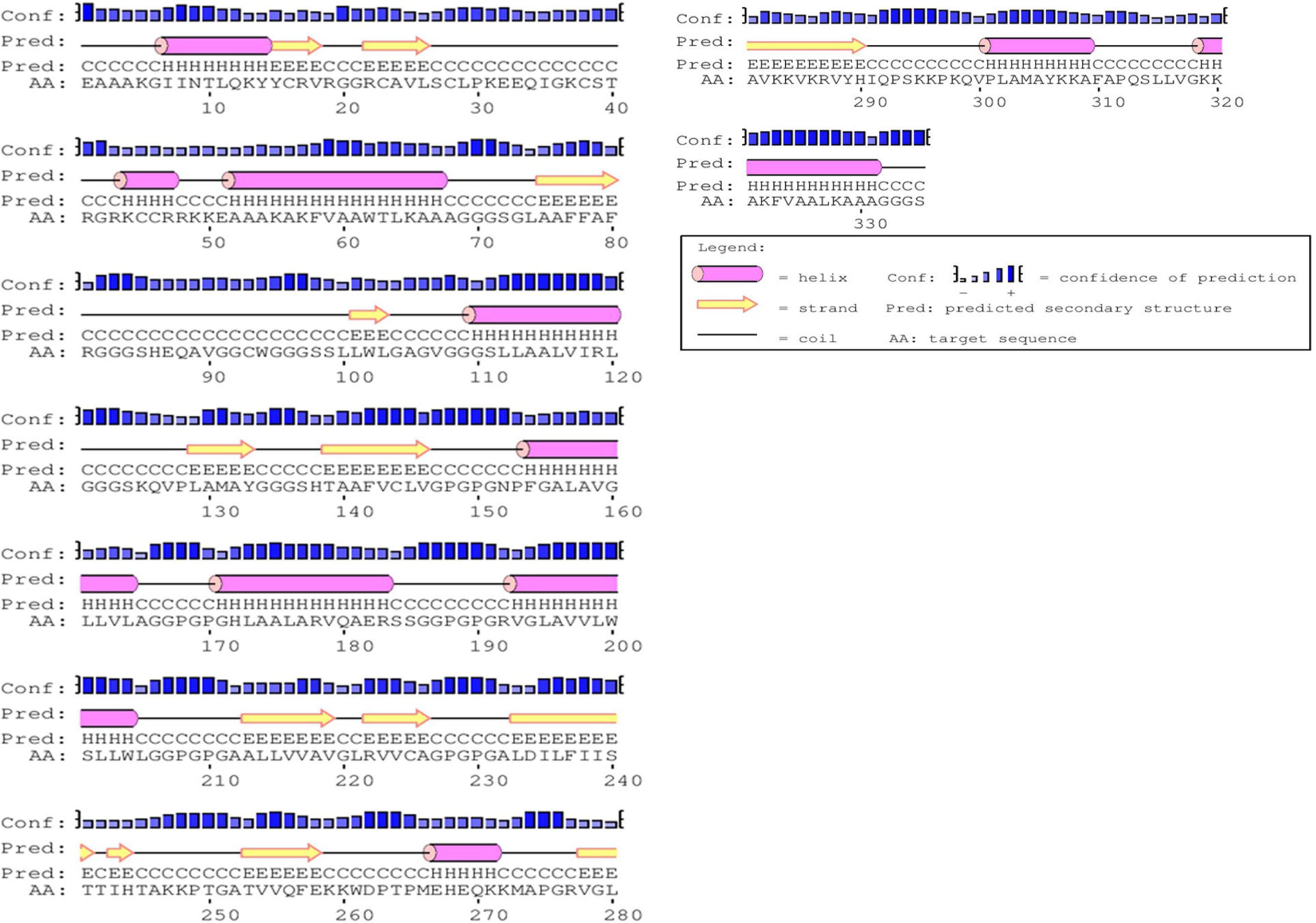
Secondary structure prediction of designed vaccine V1 using PESIPRED server

**Figure 6.**
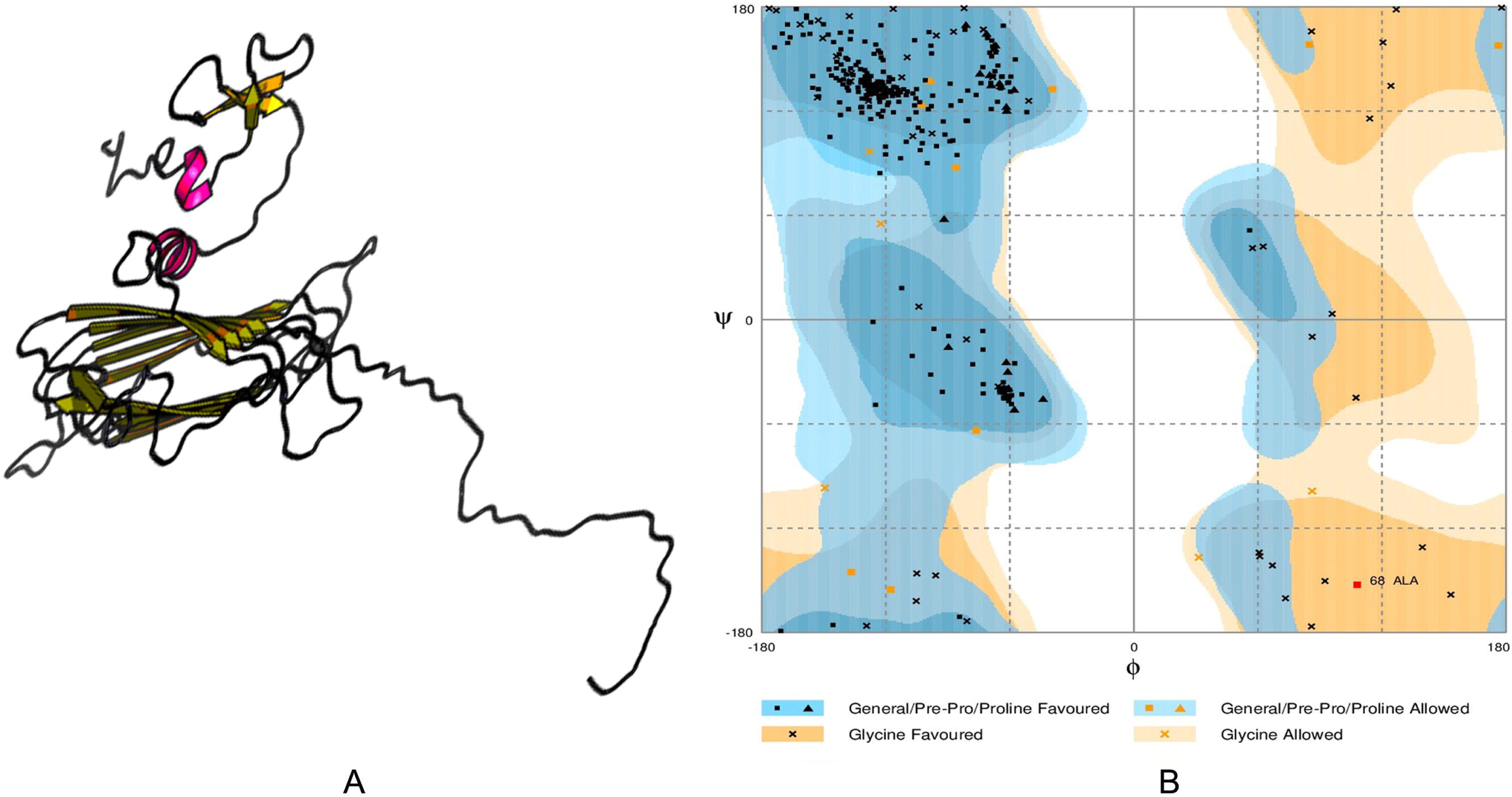
Tertiary structure prediction (A) and validation of the 3D structure of vaccine protein V1 through Ramachandran plot analysis (B).

### Tertiary structure refinement and validation

Refinement was performed to improve the quality of predicted 3D modeled structure beyond the accuracy. Before refinement Ramachandran plot analysis revealed that 92.2% residues were in the favored, 6.3% residues in the allowed and 1.5% residues in the outlier region. However, after refinement 95.5% and 4.2% residues were in the favored and allowed region respectively, while only 0.3% residues were found in the outlier region (Figure 6). Modeled tertiary structure of vaccine construct V2 and V3 have been shown in figure 7.

**Figure 7.**
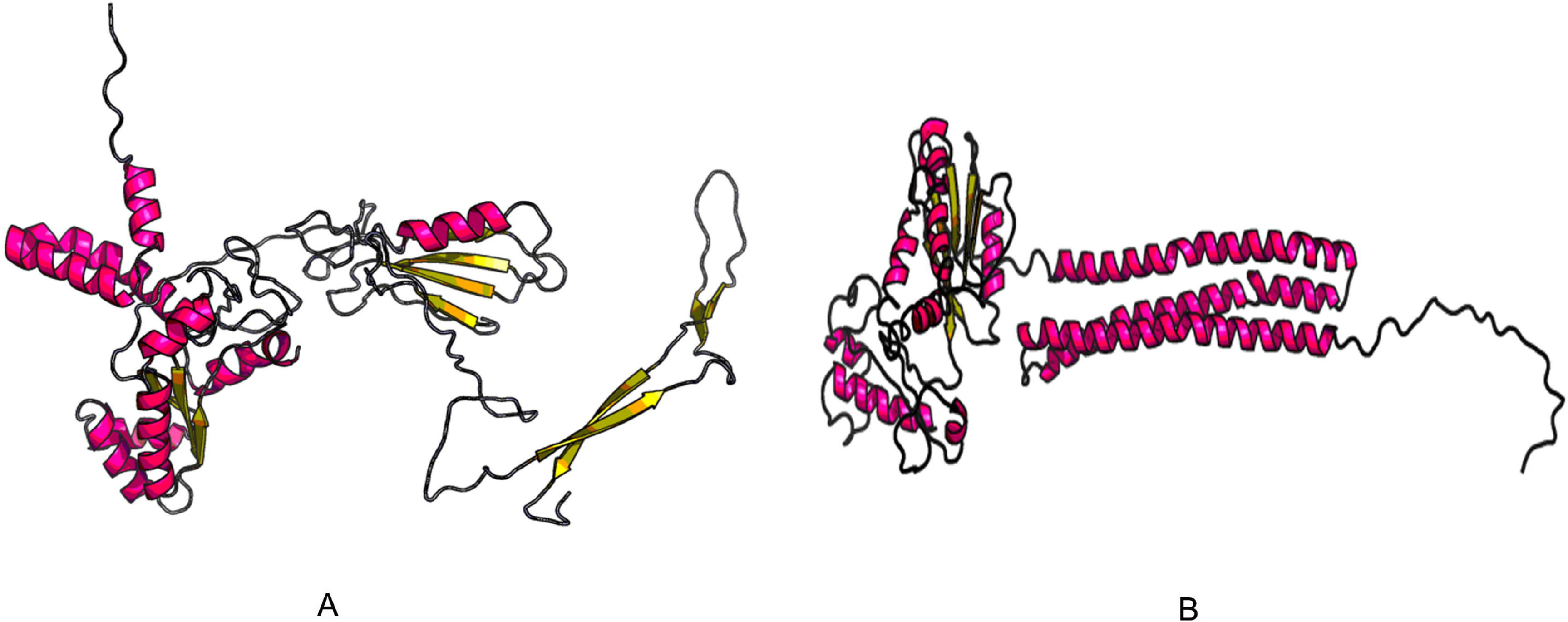
3D modelled structure of vaccine protein V2 (A) and V3 (B) generated via RaptorX server.

### Vaccine protein disulfide engineering

Residues in the highly mobile region of the protein sequence were mutated with cysteine to perform Disulfide engineering. A total 54 pairs of amino acid residue were identified having the capability to form disulfide bond by DbD2 server. After evaluation of the residue pairs in terms of energy, chi3 and B-factor parameter, only 2 pair satisfied the disulfide bond formation. Those residue pairs were GLY 82-GLY 121 and ALA 105-GLY 136 and LYS 284-ARG287 (Figure 8). All these 4 residues were replaced with cysteine residue. The value of chi3 considered for the residue screening was between −87 to +97 while the energy value was less than 2.5.

**Figure 8.**
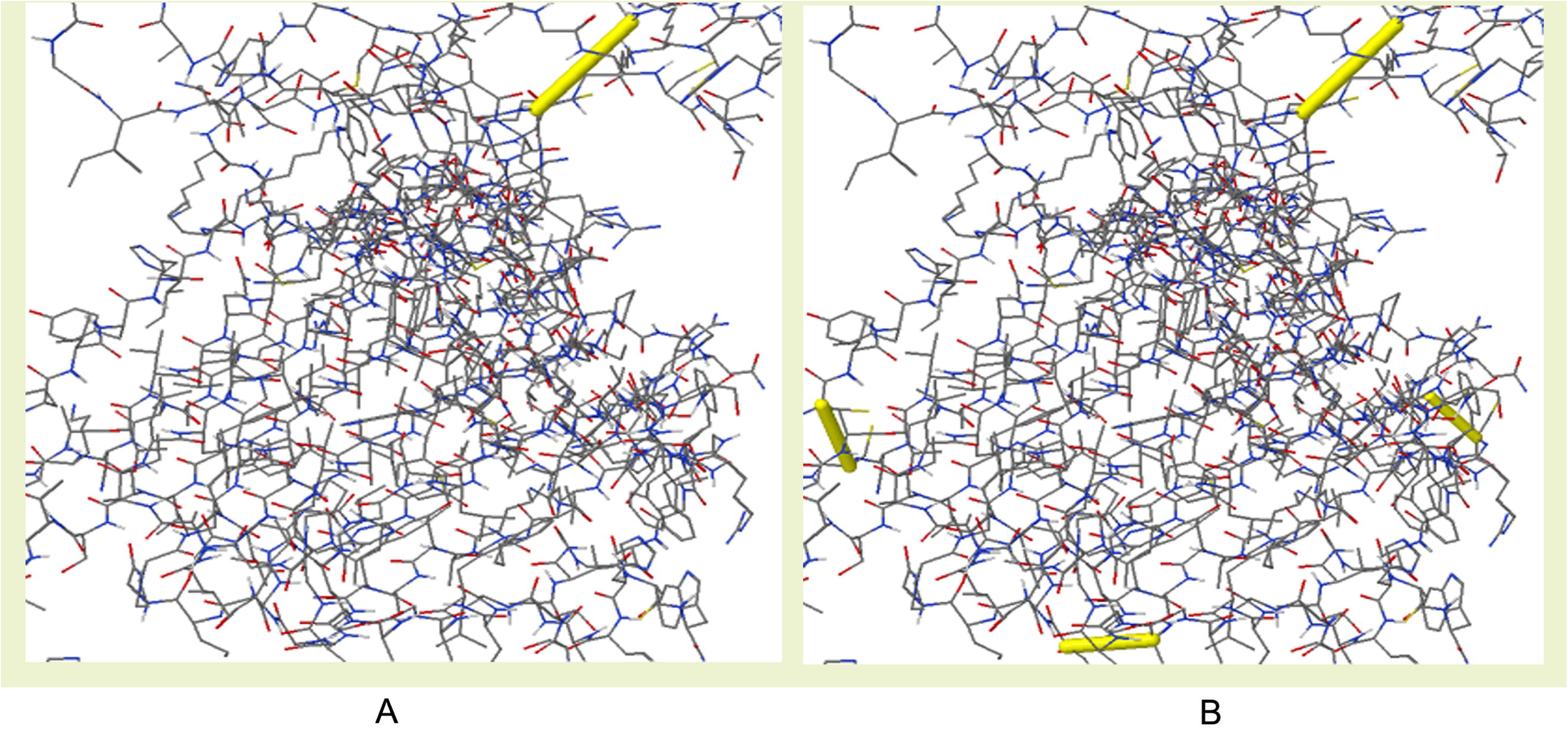
Disulfide engineering of vaccine protein V1; (A) Initial form, (B) Mutant form.

### Protein-protein docking

Docking analysis was performed between vaccine construct and different HLA alleles (Table 7). Construct V1 showed biologically significant results in terms of free binding energy. Besides, docking was also conducted to evaluate the binding affinity of designed vaccine with human TLR2 receptor using ClusPro, hdoc and PatchDock web servers (Figure 9). The ClusPro server generated 30 protein-ligand complexes as output along with respective free binding energy. The lowest energy of −1200.2 was obtained for the complex named cluster 1. The hdoc server predicted the binding energy for the protein-protein complex was −293.17, while FireDock output refinement of Patch Dock server showed the lowest global energy of - 28.07 for solution 7. The lowest binding energy of the complexes indicates the highest binding affinity between TLR-8 and vaccine construct.

**Figure 9.**
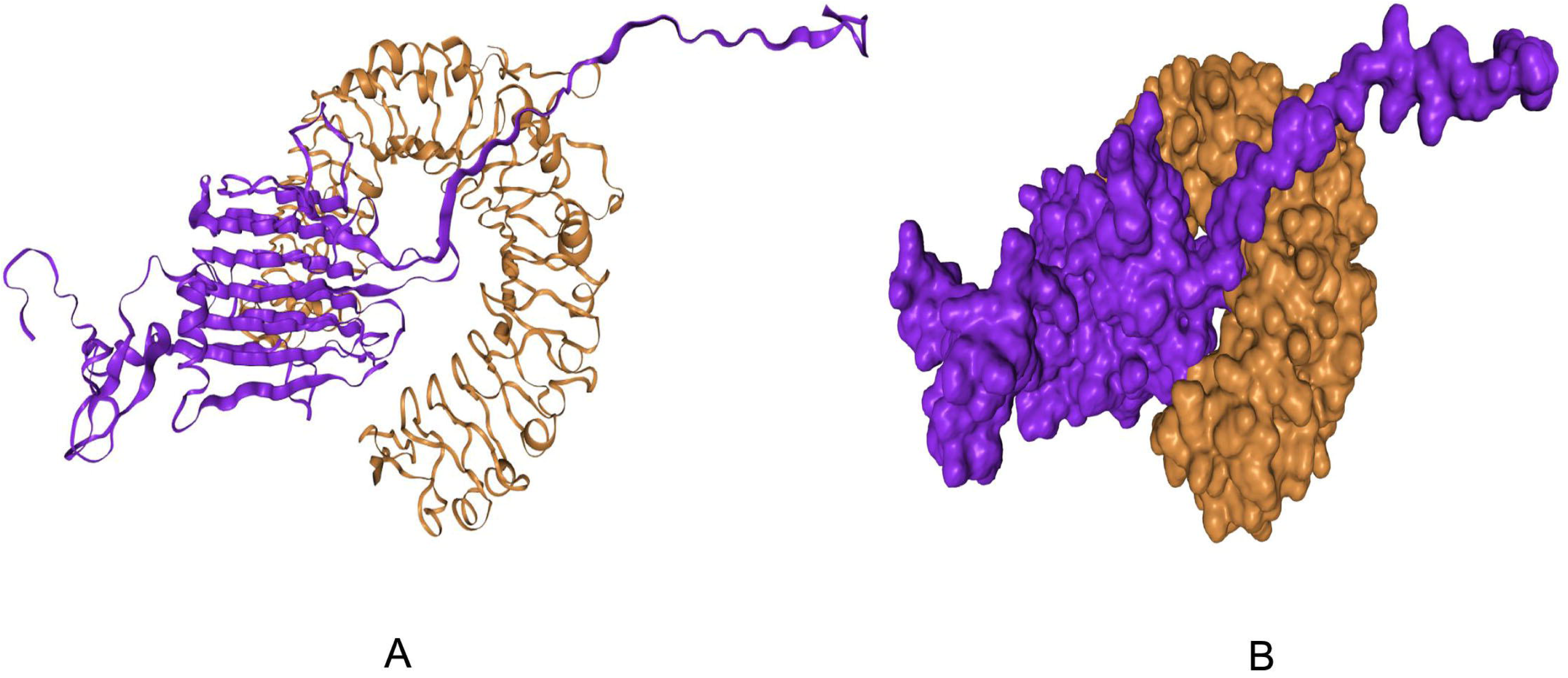
Docked complex of vaccine construct V1 with human TLR2; (A) Cartoon format and (B) Ball structure.

**Table 7.**
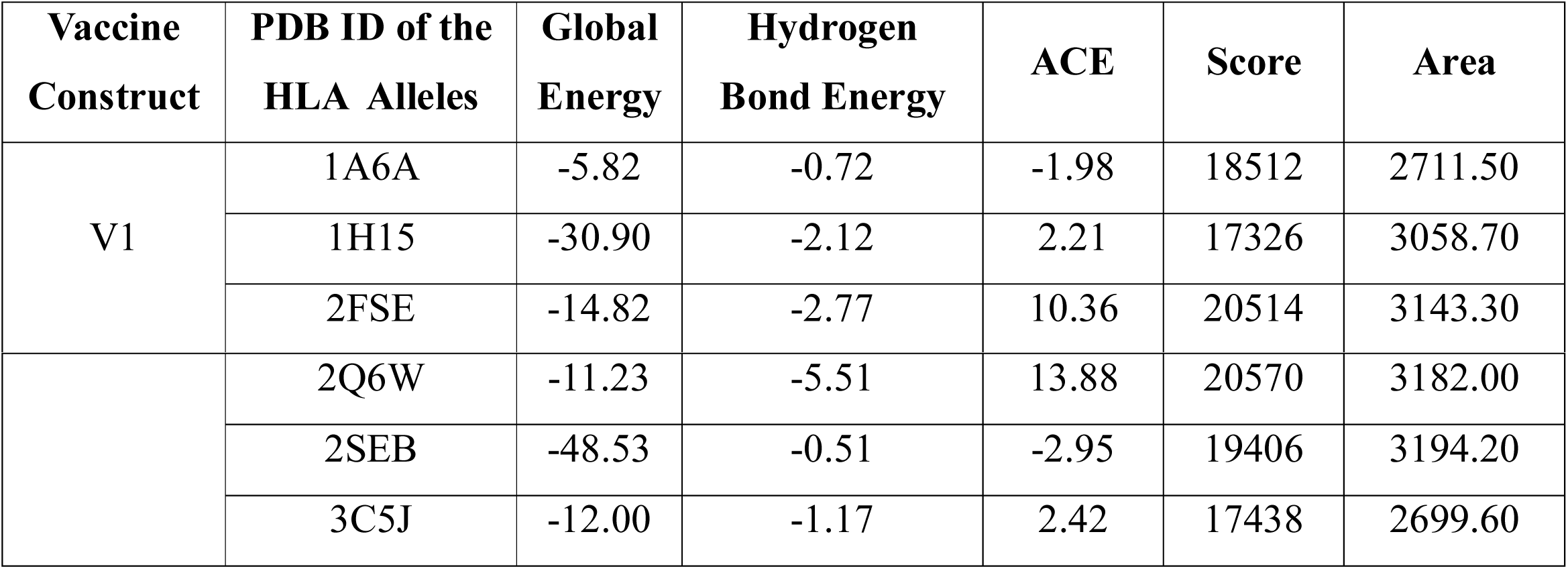
Docking score of vaccine construct V1 with different HLA alleles including HLA-DRB1*03:01 (1A6A), HLA-DRB5*01:01 (1H15), HLA-DRB1*01:01 (2FSE), HLA-DRB3*01:01 (2Q6W), HLA-DRB1*04:01 (2SEB) and HLA-DRB3*02:02 (3C5J).

| Vaccine Construct | PDB ID of the HLA Alleles | Global Energy | Hydrogen Bond Energy | ACE | Score | Area |
| --- | --- | --- | --- | --- | --- | --- |
| V1 | 1A6A | -5.82 | -0.72 | -1.98 | 18512 | 2711.50 |
|  | 1H15 | -30.90 | -2.12 | 2.21 | 17326 | 3058.70 |
|  | 2FSE | -14.82 | -2.77 | 10.36 | 20514 | 3143.30 |
|  | 2Q6W | -11.23 | -5.51 | 13.88 | 20570 | 3182.00 |
|  | 2SEB | -48.53 | -0.51 | -2.95 | 19406 | 3194.20 |
|  | 3C5J | -12.00 | -1.17 | 2.42 | 17438 | 2699.60 |

### Molecular dynamics simulation

Normal mode analysis (NMA) was performed throughiMODS server by considering the internal coordinates of the docked complex. The deformabilty of the complex depends on the individual distortion of each residues, represented by hinges in the chain (Figure 10:B). The eigenvalue found for the complex was 1.051e-05 (Figure 10:D). There was an inverse relationship between eigenvalue and the variance associated to each normal mode (Figure 10:C). The B-factor values inferred via NMA, was equivalent to RMS (Figure 10:A). Coupling between pairs of residues was indicated by the covariance matrix where different pairs showed correlated, uncorrelated or anti-correlated motions, represented by red, white and blue colors respectively (Figure 10:E). An elastic network model was also generated (Figure 10:F) according to the level of stiffness between the pairs of atoms connected via springs (the darker the grays, the stiffer the springs).

**Figure 10:**
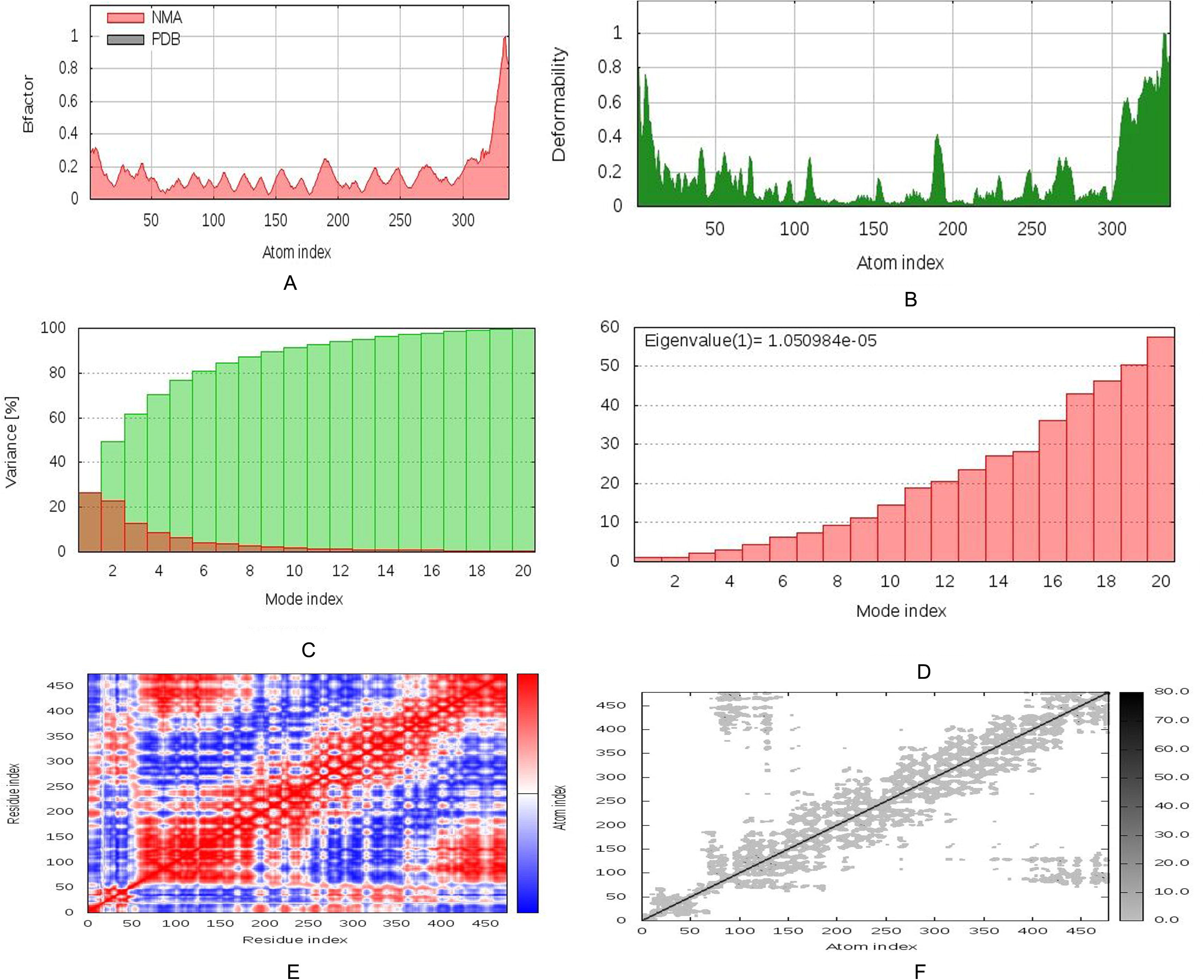
Molecular dynamics simulation of vaccine protein V1-TLR2 complex; stability of the protein-protein complex was investigated through B-factor values (A), deformability (B), variance (C), eigenvalue (D), covariance of residue index (E) and elastic network (F) analysis.

### Codon adaptation and in silico cloning

Due to dissimilarity in the regulatory systems of human and *E. coli*, codon adaptation was performed considering the expression system of the host. Construct V1 was reverse-transcribed where in the adapted codons, codon adaptation index (CAI) was 0.968 ensuring the higher proportion of most abundant codons. The GC content of the optimized codons (53.081%) was significant as well. The construct did not contain restriction sites for XhoI and EcoRI and thus indicating its safety for cloning purpose. Finally, the optimized codons were inserted into pET28a(+) vector along with XhoI and EcoRI restriction sites. A clone of 6346 base pair was produced comprising 1011bp desired sequence (shown in red color in between the sequence of pET28a(+) vector and the rest belonging to the vector only (Figure 11).

**Figure 11.**
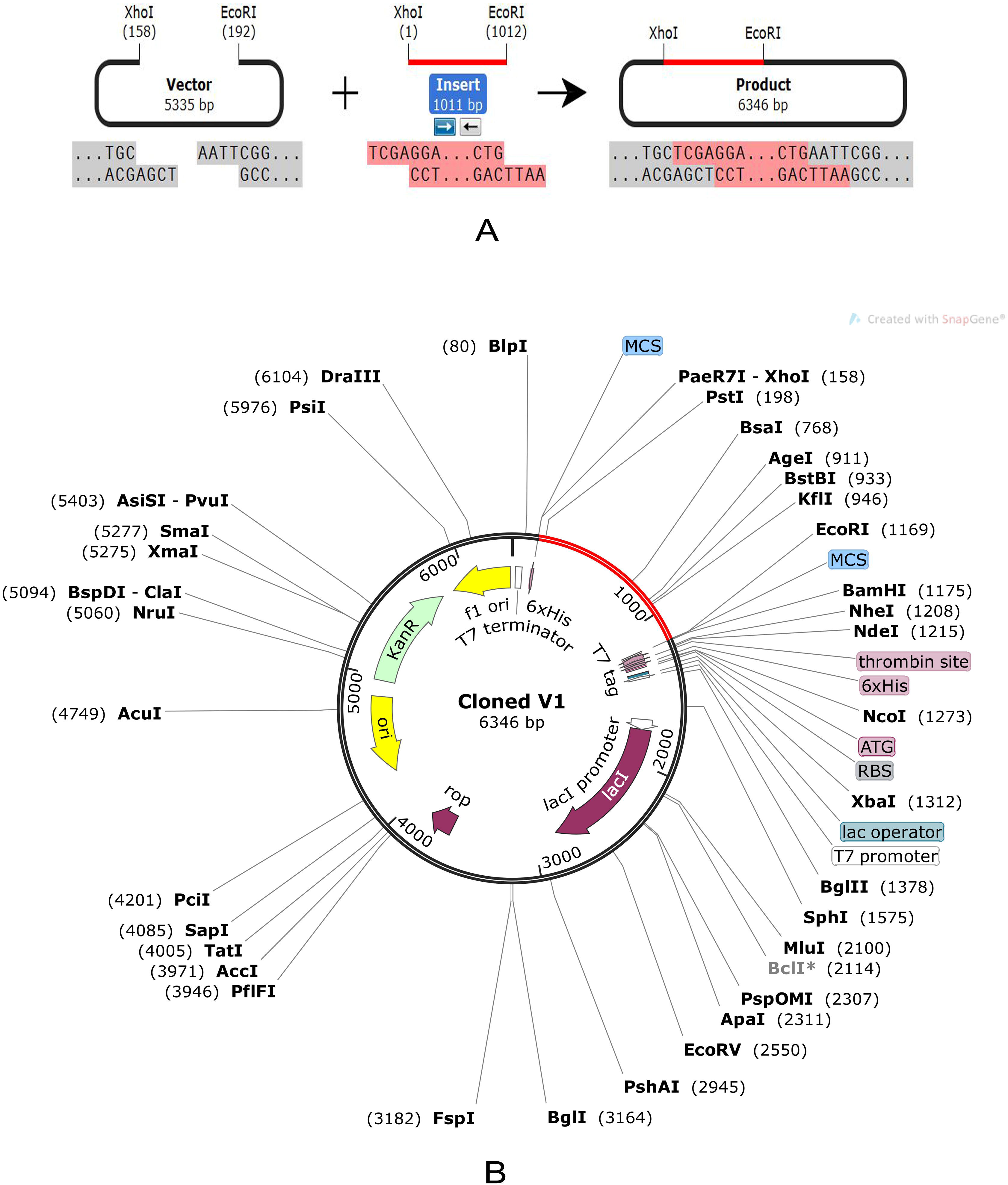
Restriction digestion (A) and *in silico* cloning (B) of the gene sequence of final vaccine construct V1 into pET28a(+) expression vector. Target sequence was inserted between XhoI and EcoRI.

## Discussion

Although studies regarding HSV have been performed extensively over the last six decades, development of an effective vaccine has remained still challenging (Whitley & Baines, 2018). No FDA verified effective vaccine has been reported that can prevent the spread or acquisition of either HSV serotype (Whitley & Baines, 2018; Boukhvalova et al., 2005). Hence, the present study aids in the path of broad spectrum vaccine development against HSV. The conventional approaches for vaccine development may take decades to experimentally validate and commercialize a product in the drug market (Babu et al., 2014; Hasan et al., 2019a). These include live or attenuated vaccines which require in vitro culture and antigen expression. Moreover most of the techniques used so far to purge antigens resulted in undesirable consequences (Rappuoli, 2000) or adverse reactions (Purcell et al., 2007; Wong et al., 2014). In this study we focused on a different strategy by taking the advantages of genome and proteome database through adopting reverse vaccinology approach (Azim et al., 2019; Hasan et al., 2015b).

The major glycoproteins (six) of Herpes Simplex Virus was retrieved in FASTA format and investigated through BLASTp analysis to find the homologous protein sets (> 90% identity) for each glycoprotein. The retrieved protein sequences belong to different strains of Type-1 and Type-2 Herpes Simplex Virus isolated from different geographic regions. Multiple sequence alignment (MSA) allowed identification of the conserved regions among these proteins. This step was followed by the prediction of potential B-cell and T-cell epitopes for each glycoproteins separately. Cytotoxic CD8+T lymphocytes (CTL) control the spread of infectious agents by recognizing and killing infected cells or secreting specific antiviral cytokines (Garcia et al., 1999). Thus, T cell epitope-based vaccination is a unique process to stimulate strong defensive response against infectious pathogen (Shrestha, 2004). However, most vaccines trigger both T cell and B cell response. Vaccine induces production of antibodies that are synthesized by B cells and mediates effector functions by interacting specifically to a pathogen or toxin (Cooper & Nemerow, 1984). Here, the most immunogenic epitopes for HTL, CTL and BCL receptors were screened through analyzing antigenicity, allergenicity, toxicity, conservancy and other physiochemical properties using a number of bioinfomatics tools. The top and finalized epitopes used to design the final vaccine constructs were validated through molecular docking approach with corresponding HLA alleles (Table 4). Predicted CTL, HTL and BCL epitopes from each protein were combined together using suitable linkers and adjuvant to develop a novel multiepitope polyvalent vaccine against Herpes Simplex Virus. It has been reported that PADRE sequence reduces the polymorphism of HLA molecules in the population (Ghaffari-Nazari et al., 2015). Therefore it was also incorporated to construct the final vaccines in this study. The selected adjuvants and linkers were used to enhance the immunogenicity and effective separation of epitopes within host (Yang et al., 2015).

Moreover, the designed vaccines were characterized based on allergenicity, antigenicity, physiochemical properties, three dimensional structure, di-sulfide engineering and stability. Construct V1 was superior and found best considering each parameter discussed above. Finally, the molecular docking and dynamics study was performed to investigate the binding affinity and complex stability of the vaccine with different HLA molecules (i.e. DRB1*0101, DRB3*0202, DRB5*0101, DRB3*0101, DRB1*0401, and DRB1*0301) and human TLR-2 receptor. The vaccine-TLR2 complex showed minimum deformability at molecular level which further strengthened our prediction. *In-silico* cloning was conducted to ensure the stability and effective expression of designed vaccine construct.

## Conclusion

*In-silico* studies can guide experimental works with higher probabilities of finding the appropriate solutions through fewer trials and error repeats thus saving time and costs for the researchers. In this study, we developed a unique chimeric vaccine with the potential to confer immunity against both type (Type-1 and Type-2) of Herpes Simplex Virus and the expression system was also analyzed. Our predicted results were based on diligent analysis of sequence and various immune databases. However, further *in vivo* trials using model organisms are highly recommended to validate our findings.

## Acknowledgments

Authors like to acknowledge the authority of the Bioinformatics Laboratory of Sylhet Agricultural University for the technical support of the project.

## Author contributions

*Mahmudul Hasan:* Conceptualization, Supervision, Project Administration and Reviewing;

*Shiful Islam:* Experiment Design, Data Handling, T-Cell Epitope Prediction and Docking Study

*Sourav Chakraborty:* Experiment Design, Data Handling and B-Cell Epitope Prediction

*Abu Hasnat Mustafa:* Data Handling, Conservancy Analysis and Docking Study

*Kazi Faizul Azim:* Data Handling, Data Analysis, Vaccine Modeling, Manuscript writing

*Ziaul Faruque Joy:* Simulation Study and Expression Analysis

*Md Nazmul Hossain:* Data Organization, Manuscript writing

*Shakhawat Hossain Foysal:* Data Organization, Disulfide Engineering and Homology modeling

*Md Nazmul Hasan:* Population Coverage Study and Manuscript Reviewing

