## Supplementary file 1 for "Contriving a chimeric polyvalent vaccine to prevent infections caused by Herpes Simplex Virus (Type-1 and Type-2): an exploratory immunoinformatic approach"

**GLYCOPROTEIN B**

**HSV1=20 HSV2=19**

>HS1GPB_1 MRQGAPARGRRWFVVWALLGLTLGVLVASAAPSSPGTPGVAAATQAANGGPATPAPPAPGAPPTGDPKPK

KNRKPKPPKPPRPAGDNATVAAGHATLREHLRDIKAENTDANFYVCPPPTGATVVQFEQPRRCPTRPEGQ

NYTEGIAVVFKENIAPYKFKATMYYKDVTVSQVWFGHRYSQFMGIFEDRAPVPFEEVIDKINAKGVCRST

AKYVRNNLETTAFHRDDHETDMELKPANAATRTSRGWHTTDLKYNPSRVEAFHRYGTTVNCIVEEVDARS

VYPYDEFVLATGDFVYMSPFYGYREGSHTEHTSYAADRFKQVDGFYARDLTTKARATAPTTRNLLTTPKF

TVAWDWVPKRPSVCTMTKWQEVDEMLRSEYGGSFRFSSDAISTTFTTNLTEYPLSRVDLGDCIGKDARDA

MDRIFARRYNATHIKVGQPQYYLANGGFLIAYQPLLSNTLAELYVREHLREQSRKPPNPTPPPPGASANA

SVERIKTTSSIEFARLQFTYNHIQRHVNDMLGRVAIAWCELQNHELTLWNEARKLNPNAIASATVGRRVS

ARMLGDVMAVSTCVPVAADNVIVQNSMRISSRPGACYSRPLVSFRYEDQGPLVEGQLGENNELRLTRDAI

EPCTVGHRRYFTFGGGYVYFEEYAYSHQLSRADITTVSTFIDLNITMLEDHEFVPLEVYTRHEIKDSGLL

DYTEVQRRNQLHDLRFADIDTVIHADANAAMFAGLGAFFEGMGDLGRAVGKVVMGIVGGVVSAVSGVSSF

MSNPFGALAVGLLVLAGLAAAFFAFRYVMRLQSNPMKALYPLTTKELKNPTNPDASGEGEEGGDFDEAKL

AEAREMIRYMALVSAMERTEHKAKKKGTSALLSAKVTDMVMRKRRNTNYTQVPNKDGDADEDDL

>HS1GPB_2 MRQGAPARGRRWFVVWALLGLTLGVLVASAAPSSPGTPGVAAATQAANGGPATPAPPAPGAPPTGDPKPK

KNKKPKPPKPPRPAGDNATVAAGHATLREHLRDIKAENTDANFYVCPPPTGATVVQFEQPRRCPTRPEGQ

NYTEGIAVVFKENIAPYKFKATMYYKDVTVSQVWFGHRYSQFMGIFEDRAPVPFEEVIDKINAKGVCRST

AKYVRNNLETTAFHRDDHETDMELKPANAATRTSRGWHTTDLKYNPSRVEAFHRYGTTVNCIVEEVDARS

VYPYDEFVLATGDFVYMSPFYGYREGSHTEHTSYAADRFKQVDGFYARDLTTKARATAPTTRNLLTTPKF

TVAWDWVPKRPSVCTMTKWQEVDEMLRSEYGGSFRFSSDAISTTFTTNLTEYPLSRVDLGDCIGKDARDA

MDRIFARRYNATHIKVGQPQYYLANGGFLIAYQPLLSNTLAELYVREHLREQSRKPPNPTPPPPGASANA

SVERIKTTSSIEFARLQFTYNHIQRHVNDMLGRVAIAWCELQNHELTLWNEARKLNPNAIASATVGRRVS

ARMLGDVMAVSTCVPVAADNVIVQNSMRISSRPGACYSRPLVSFRYEDQGPLVEGQLGENNELRLTRDAI

EPCTVGHRRYFTFGGGYVYFEEYAYSHQLSRADITTVSTFIDLNITMLEDHEFVPLEVYTRHEIKDSGLL

DYTEVQRRNQLHDLRFADIDTVIHADANAAMFAGLGAFFEGMGDLGRAVGKVVMGIVGGVVSAVSGVSSF

MSNPFGALAVGLLVLAGLAAAFFAFRYVMRLQSNPMKALYPLTTKELKNPTNPDASGEGEEGGDFDEAKL

AEAREMIRYMALVSAMERTEHKAKKKGTSALLSAKVTDMVMRKRRNTNYTQVPNKDGDADEDDL

>HS1GPB_3

MRQGAPARGRRWFVVWALLGLTLGVLVVSAAPSSPGTPGVAAATQAANGGPATPAPPAPGAPPTGDPKPK

KNKKPKPPKPPRPAGDNATVAAGHATLREHLRDIKAENTDANFYVCPPPTGATVVQFEQPRRCPTRPEGQ

NYTEGIAVVFKENIAPYKFKATMYYKDVTVSQVWFGHRYSQFMGIFEDRAPVPFEEVIDKINAKGVCRST

AKYVRNNLETTAFHRDDHETDMELKPANAATRTSRGWHTTDLKYNPSRVEAFHRYGTTVNCIVEEVDARS

VYPYDEFVLATGDFVYMSPFYGYREGSHTEHTSYAADRFKQVDGFYARDLTTKARATAPTTRNLLTTPKF

TVAWDWVPKRPSVCTMTKWQEVDEMLRSEYGGSFRFSSDAISTTFTTNLTEYPLSRVDLGDCIGKDARDA

MDRIFARRYNATHIKVGQPQYYLANGGFLIAYQPLLSNTLAELYVREHLREQSRKPPNPTPPPPGASANA

SVERIKTTSSIEFARLQFTYNHIQRHVNDMLGRVAIAWCELQNHELTLWNEARKLNPNAIASATVGRRVS

ARMLGDVMAVSTCVPVAADNVIVQNSMRISSRPGACYSRPLVSFRYEDQGPLVEGQLGENNELRLTRDAI

EPCTVGHRRYFTFGGGYVYFEEYAYSHQLSRADITTVSTFIDLNITMLEDHEFVPLEVYTRHEIKDSGLL

DYTEVQRRNQLHDLRFADIDTVIHADANAAMFAGLGAFFEGMGDLGRAVGKVVMGIVGGVVSAVSGVSSF

MSNPFGALAVGLLVLAGLAAAFFAFRYVMRLQSNPMKALYPLTTKELKNPTNPDASGEGEEGGDFDEAKL

AEAREMIRYMALVSAMERTEHKAKKKGTSALLSAKVTDMVMRKRRNTNYTQVPNKDGDADEDDL

>HS1GPB_4 MRQGAPARGRRWFVVWALLGLTLGVLVASAAPSSPGTPGVAAATQAANGGPATPAPPAPGAPPTGDPKPK

KNKKPKPPKPPRPAGDNATVAAGHATLREHLRDIKAENTDANFYVCPPPTGATVVQFEQPRRCPTRPEGQ

NYTEGIAVVFKENIAPYKFKATMYYKDVTVSQVWFGHRYSQFMGIFEDRAPVPFEEVIDKINAKGVCRST

AKYVRNNLETTAFHRDDHETDMELKPANAATRTSRGWHTTDLKYNPSRVEAFHRYGTTVNCIVEEVDARS

VYPYDEFVLATGDFVYMSPFYGYREGSHTEHTSYAADRFKQVDGFYARDLTTKARATAPTTRNLLTTPKF

TVAWDWVPKRPSVCTMTKWQEVDEMLRSEYGGSFRFSSDAISTTFTTNLTEYPLSRVDLGDCIGKDARDA

MDRIFARRYNATHIKVGQPQYYLANGGFLIAYQPLLSNTLAELYVREHLREQSRKPPNPTPPPPGASANA

SVERIKTTSSIEFARLQFTYNHIQRHVNDMLGRVAIAWCELQNHELTLWNEARKLNPNAIASATVGRRVS

ARMLGDVMAVSTCVPVAADNVIVQNSMRISSRPGACYSRPLVSFRYEDQGPLVEGQLGENNELRLTRDAI

EPCTVGHRRYFTFGGGYVYFEEYAYSHQLSRADITTVSTFIDLNITMLEDHEFVPLEVYTRHEIKDSGLL

DYTEVQRRNQLHDLRFADIDTVIHADANAAMFAGLGAFFEGMGDLGRAVGKVVMGIVGGVVSAVSGVSSF

ISNPFGALAVGLLVLAGLAAAFFAFRYVMRLQSNPMKALYPLTTKELKNPTNPDASGEGEEGGDFDEAKL

AEAREMIRYMALVSAMERTEHKAKKKGTSALLSAKVTDMVMRKRRNTNYTQVPNKDGDADEDDL

>HS1GPB_5

MRQGAPARGRRWFVVWALLGLTLGVLVASAAPSSPGTPGVAAATQAANGGPATPAPPAPGAPPTGDPKPK

KNKKPKPPKPPRPAGDNATVAAGHATLREHLRDIKAENTDANFYVCPPPTGATVVQFEQPRRCPTRPEGQ

NYTEGIAVVFKENIAPYKFKATMYYKDVTVSQVWFGHRYSQFMGIFEDRAPVPFEEVIDKINAKGVCRST

AKYVRNNLETTAFHRDDHETDMELKPANAATRTSRGWHTTDLKYNPSRVEAFHRYGTTVNCIVEEVDARS

VYPYNEFVLATGDFVYMSPFYGYREGSHTEHTSYAADRFKQVDGFYARDLTTKARATAPTTRNLLTTPKF

TVAWDWVPKRPSVCTMTKWQEVDEMLRSEYGGSFRFSSDAISTTFTTNLTEYPLSRVDLGDCIGKDARDA

MDRIFARRYNATHIKVGQPQYYLANGGFLIAYQPLLSNTLAELYVREHLREQSRKPPNPTPPPPGASANA

SVERIKTTSSIEFARLQFTYNHIQRHVNDMLGRVAIAWCELQNHELTLWNEARKLNPNAIASATVGRRVS

ARMLGDVMAVSTCVPVAADNVIVQNSMRISSRPGACYSRPLVSFRYEDQGPLVEGQLGENNELRLTRDAI

EPCTVGHRRYFTFGGGYVYFEEYAYSHQLSRADITTVSTFIDLNITMLEDHEFVPLEVYTRHEIKDSGLL

DYTEVQRRNQLHDLRFADIDTVIHADANAAMFAGLGAFFEGMGDLGRAVGKVVMGIVGGVVSAVSGVSSF

MSNPFGALAVGLLVLAGLAAAFFAFRYVMRLQSNPMKALYPLTTKELKNPTNPDASGEGEEGGDFDEAKL

AEAREMIRYMALVSVMERTEHKAKKKGTSALLSAKVTDMVMRKRRNTNYTQVPNKDGDADEDDL

>HS1GPB_6

MRQGAPARGRRWFVVWALLGLTLGVLVASAAPSSPGTPGVAAATQAANGGPATPAPPALGAAPTGDPKPK

KNKKPKNPTPPRPAGDNATVAAGHATLREHLRDIKAENTDANFYVCPPPTGATVVQFEQPRRCPTRPEGQ

NYTEGIAVVFKENIAPYKFKATMYYKDVTVSQVWFGHRYSQFMGIFEDRAPVPFEEVIDKINAKGVCRST

AKYVRNNLETTAFHRDDHETDMELKPANAATRTSRGWHTTDLKYNPSRVEAFHRYGTTVNCIVEEVDARS

VYPYDEFVLATGDFVYMSPFYGYREGSHTEHTSYAADRFKQVDGFYARDLTTKARATAPTTRNLLTTPKF

TVAWDWVPKRPSVCTMTKWQEVDEMLRSEYGGSFRFSSDAISTTFTTNLTEYPLSRVDLGDCIGKDARDA

MDRIFARRYNATHIKVGQPQYYLANGGFLIAYQPLLSNTLAELYVREHLREQSRKPPNPTPPPPGASANA

SVERIKTTSSIEFARLQFTYNHIQRHVNDMLGRVAIAWCELQNHELTLWNEARKLNPNAIASATVGRRVS

ARMLGDVMAVSTCVPVAADNVIVQNSMRISSRPGACYSRPLVSFRYEDQGPLVEGQLGENNELRLTRDAI

EPCTVGHRRYFTFGGGYVYFEEYAYSHQLSRADITTVSTFIDLNITMLEDHEFVPLEVYTRHEIKDSGLL

DYTEVQRRNQLHDLRFADIDTVIHADANAAMFAGLGAFFEGMGDLGRAVGKVVMGIVGGVVSAVSGVSSF

MSNPFGALAVGLLVLAGLAAAFFAFRYVMRLQSNPMKALYPLTTKELKNPTNPDASGEGEEGGDFDEAKL

AEAREMIRYMALVSAMERTEHKAKKKGTSALLSAKVTDMVMRKRRNTNYTQVPNKDGDADEDDL

>HS1GPB_7 MRQGAPARGRRWFVVWALLGLTLGVLVASAAPSSPGTPGVAAATQAANGGPATPAPPAPGAPPTGDPKPK

KNKKPKPPKLPRPAGDNATVAAGHATLREHLRDIKAENTDANFYVCPPPTGATVVQFEQPRRCPTRPEGQ

NYTEGIAVVFKENIAPYKFKATMYYKDVTVSQVWFGHRYSQFMGIFEDRAPVPFEEVIDKINAKGVCRST

AKYVRNNLETTAFHRDDHETDMELKPANAATRTSRGWHTTDLKYNPSRVEAFHRYGTTVNCIVEEVDARS

VYPYDEFVLATGDFVYMSPFYGYREGSHTEHTSYAADRFKQVDGFYARDLTTKARATAPTTRNLLTTPKF

TVAWDWVPKRPSVCTMTKWQEVDEMLRSEYGGSFRFSSDAISTTFTTNLTEYPLSRVDLGDCIGKDARDA

MDRIFARRYNATHIKVGQPQYYLANGGFLIAYQPLLSNALAELYVREHLREQSRKPPNPTPPPPGASANA

SVERIKTTSSIEFARLQFTYNHIQRHVNDMLGRVAIAWCELQNHELTLWNEARKLNPNAIASATMGRRVS

ARMLGDVMAVSTCVPVAADNVIVQNSMRISSRPGACYSRPLVSFRYEDQGPLVEGQLGENNELRLTRDAI

EPCTVGHRRYFTFGGGYVYFEEYAYSHQLSRADITTVNTFIDLNITMLEDHEFVPLEVYTRHEIKDSGLL

DYTEVQRRNQLHDLRFADIDTVIHADANAAMFAGLGAFFEGMGDLGRAVGKVVMGIVGGVVSAVSGVSSF

MSNPFGALAVGLLVLAGLAAAFFAFRYVMRLQSNPMKALYPLTTKEPKNPTNPDASGEGEEGGDFDEAKL

AEAREMIRYMALVSAMERTEHKAKKKGTSALLSAKVTDMVMRKRRNTNYTQVPNKDGDADEDDL

>HS1GPB_8 MRQGAPARGRRWFVVWALLGLTLGVLVASAAPSAPGTPGVAAATQAANGGPATPAPPAPGAAPTGDTKPK

KNKKPKNPTPPRPAGDNATVAAGHATLREHLRDIKAENTDANFYVCPPPTGATVVQFEQPRRCPTRPEGQ

NYTEGIAVVFKENIAPYKFKATMYYKDVTVSQVWFGHRYSQFMGIFEDRAPVPFEEVIDKINAKGVCRST

AKYVRNNLETTAFHRDDHETDMELKPANAATRTSRGWHTTDLKYNPSRVEAFHRYGTTVNCIVEEVDARS

VYPYDEFVLATGDFVYMSPFYGYREGSHTEHTSYAADRFKQVDGFYARDLTTKARATAPTTRNLLTTPKF

TVAWDWVPKRPSVCTMTKWQEVDEMLRSEYGGSFRFSSDAISTTFTTNLTEYPLSRVDLGDCIGKDARDA

MDRIFARRYNATHIKVGQPQYYLANGGFLIAYQPLLSNTLAELYVREHLREQSRKPPNPTPPPPGASANA

SVERIKTTSSIEFARLQFTYNHIQRHVNDMLGRVAIAWCELQNHELTLWNEARKLNPNAIASATVGRRVS

ARMLGDVMAVSTCVPVAADNVIVQNSMRISSRPGACYSRPLVSFRYEDQGPLVEGQLGENNELRLTRDAI

EPCTVGHRRYFTFGGGYVYFEEYAYSHQLSRADITTVSTFIDLNITMLEDHEFVPLEVYTRHEIKDSGLL

DYTEVQRRNQLHDLRFADIDTVIHADANAAMFAGLGAFFEGMGDLGRAVGKVVMGIVGGVVSAVSGVSSF

MSNPFGALAVGLLVLAGLAAAFFAFRYVMRLQSNPMKALYPLTTKELKNPTNPDASGEGEEGGDFDEAKL

AEAREMIRYMALVSAMERTEHKAKKKGTSALLSAKVTDMVMRKRRNTNYTQVPNKDGDADEDDL

>HS1GPB_9

MRQGAPARGRRWFVVWALLGLTLGVLVASAAPSSPGTPGVAAATQAANGGPATPAPPAPGAPPTGDPKPK

KNKKPKPPKLPRPAGDNATVAAGHATLREHLRDIKAENTDANFYVCPPPTGATVVQFEQPRRCPTRPEGQ

NYTEGIAVVFKENIAPYKFKATMYYKDVTVSQVWFGHRYSQFMGIFEDRAPVPFEEVIDKINAKGVCRST

AKYVRNNLETTAFHRDDHETDMELKPANAATRTSRGWHTTDLKYNPSRVEAFHRYGTTVNCIVEEVDARS

VYPYDEFVLATGDFVYMSPFYGYREGSHTEHTSYAADRFKQVDGFYARDLTTKARATAPTTRNLLTTPKF

TVAWDWVPKRPSVCTMTKWQEVDEMLRSEYGGSFRFSSDAISTTFTTNLTEYPLSRVDLGDCIGKDARDA

MDRIFARRYNATHIKVGQPQYYLANGGFLIAYQPLLSNTLAELYVREHLREQSRKPPNPTPPPPGASANA

SVERIKTTSSIEFARLQFTYNHIQRHVNDMLGRVAIAWCELQNHELTLWNEARKLNPNAIASATMGRRVS

ARMLGDVMAVSMCVPVAADNVIVQNSMRISSRPGACYSRPLVSFRYEDQGPLVEGQLGENNELRLTRDAI

EPCTVGHRRYFTFGGGYVYFEEYAYSHQLSRADITTVNTFIDLNITMLEDHEFVPLEVYTRHEIKDSGLL

DYTEVQRRNQLHDLRFADIDTVIHADANAAMFAGLGAFFEGMGDLGRAVGKVVMGIVGGVVSAVSGVSSF

MSNPFGALAVGLLVLAGLAAAFFAFRYVMRLQSNPMKALYPLTTKEPKNPTNPDASGEGEEGGDFDEAKL

AEAREMIRYMALVSAMERTEHKAKKKGTSALLSAKVTDMVMRKRRNTNYTQVPNKDGDADEDDL

>HS1GPB_10

MRQGAPARGRRWFVVWALLGLTLGVLVASAAPSSPGTPGVAAATQAANGGPATPAPPAPGPAPTGDPKPR

KNKKPKNPTPPRPAGDNATVAAGHATLREHLRDIKAENTDANFYVCPPPTGATVVQFEQPRRCPTRPEGQ

NYTEGIAVVFKENIAPYKFKATMYYKDVTVSQVWFGHRYSQFMGIFEDRAPVPFEEVIDKINAKGVCRST

AKYVRNNLETTAFHRDDHETDMELKPANAATRTSRGWHTTDLKYNPSRVEAFHRYGTTVNCIVEEVDARS

VYPYDEFVLATGDFVYMSPFYGYREGSHTEHTSYAADRFKQVDGFYARDLTTKARATAPTTRNLLTTPKF

TVAWDWVPKRPSVCTMTKWQEVDEMLRSEYGGSFRFSSDAISTTFTTNLTEYPLSRVDLGDCIGKDARDA

MDRIFARRYNATHIKVGQPQYYLANGGFLIAYQPLLSNTLAELYVREHLREQSRKPPNPTPPPPGASANA

SVERIKTTSSIEFARLQFTYNHIQRHVNDMLGRVAIAWCELQNHELTLWNEARKLNPNAIASATVGRRVS

ARMLGDVMAVSTCVPVAADNVIVQNSMRISSRPGACYSRPLVSFRYEDQGPLVEGQLGENNELRLTRDAI

EPCTVGHRRYFTFGGGYVYFEEYAYSHQLSRADITTVSTFIDLNITMLEDHEFVPLEVYTRHEIKDSGLL

DYTEVQRRNQLHDLRFADIDTVIHADANAAMFAGLGAFFEGMGDLGRAVGKVVMGIVGGVVSAVSGVSSF

MSNPFGALAVGLLVLAGLAAAFFAFRYVMRLQSNPMKALYPLTTKELKNPTNPDASGEGEEGGDFDEAKL

AEAREMIRYMALVSAMERTEHKAKKKGTSALLSAKVTDMVMRKRRNTNYTQVPNKDGDADEDDL

>HS1GPB_11

MRQGAPARGRRWFVVWALLGLTLGVLVASAAPSSPGTPGVAAATPAANGGPATPAPPALGAAPTGDPKPK

KNKKPKNPTPPRPAGDNATVAAGHATLREHLRDIKAENTDANFYVCPPPTGATVVQFEQPRRCPTRPEGQ

NYTEGIAVVFKENIAPYKFKATMYYKDVTVSQVWFGHRYSQFMGIFEDRAPVPFEEVIDKINAKGVCRST

AKYVRNNLETTAFHRDDHETDMELKPANAATRTSRGWHTTDLKYNPSRVEAFHRYGTTVNCIVEEVDARS

VYPYDEFVLATGDFVYMSPFYGYREGSHTEHTSYAADRFKQVDGFYARDLTTKARATAPTTRNLLTTPKF

TVAWDWVPKRPSVCTMTKWQEVDEMLRSEYGGSFRFSSDAISTTFTTNLTEYPLSRVDLGDCIGKDARDA

MDRIFARRYNATHIKVGQPQYYLANGGFLIAYQPLLSNTLAELYVREHLREQSRKPPNPTPPPPGASANA

SVERIKTTSSIEFARLQFTYNHIQRHVNDMLGRVAIAWCELQNHELTLWNEARKLNPNAIASATVGRRVS

ARMLGDVMAVSTCVPVAADNVIVQNSMRISSRPGACYSRPLVSFRYEDQGPLVEGQLGENNELRLTRDAI

EPCTVGHRRYFTFGGGYVYFEEYAYSHQLSRADITTVSTFIDLNITMLEDHEFVPLEVYTRHEIKDSGLL

DYTEVQRRNQLHDLRFADIDTVIHADANAAMFAGLGAFFEGMGDLGRAVGKVVMGIVGGVVSAVSGVSSF

MSNPFGALAVGLLVLAGLAAAFFAFRYVMRLQSNPMKALYPLTTKELKNPTNPDASGEGEEGGDFDEAKL

AEAREMIRYMALVSAMERTEHKAKKKGTSALLSAKVTDMVMRKRRNTNYTQVPNKDGDADEDDL

>HS1GPB_12

MRQGAPTRGCRWFVVWALLGLTLGVLVASAAPSSPGTPGVAAATQAANGGPATPAPPAPGPAPTGDPKPR

KNKKPKPPTPPRPAGDNATVAAGHATLREHLRDIKAENTDANFYVCPPPTGATVVQFEQPRRCPTRPEGQ

NYTEGIAVVFKENIAPYKFKATMYYKDVTVSQVWFGHRYSQFMGIFEDRAPVPFEEVIDKINAKGVCRST

AKYVRNNLETTAFHRDDHETDMELKPANAATRTSRGWHTTDLKYNPSRVEAFHRYGTTVNCIVEEVDARS

VYPYDEFVLATGDFVYMSPFYGYREGSHTEHTSYAADRFKQVDGFYARDLTTKARATAPTTRNLLTTPKF

TVAWDWVPKRPSVCTMTKWQEVDEMLRSEYGGSFRFSSDAISTTFTTNLTEYPLSRVDLGDCIGKDARDA

MDRIFARRYNATHIKVGQPQYYLANGGFLIAYQPLLSNTLAELYVREHLREQSRKPPNPTPPPPGASANA

SVERIKTTSSIEFARLQFTYNHIQRHVNDMLGRVAIAWCELQNHELTLWNEARKLNPNAIASATVGRRVS

ARMLGDVMAVSTCVPVAADNVIVQNSMRISSRPGACYSRPLVSFRYEDQGPLVEGQLGENNELRLTRDAI

EPCTVGHRRYFTFGGGYVYFEEYAYSHQLSRADITTVSTFIDLNITMLEDHEFVPLEVYTRHEIKDSGLL

DYTEVQRRNQLHDLRFADIDTVIHADANAAMFAGLGAFFEGMGDLGRAVGKVVMGIVGGVVSAVSGVSSF

MSNPFGALAVGLLVLAGLAAAFFAFRYVMRLQSNPMKALYPLTTKELKNPTNPDASGEGEEGGDFDEAKL

AEAREMIRYMALVSAMERTEHKAKKKGTSALLSAKVTDMVMRKRRNTNYTQVPNKDGDADEDDL

>HS1GPB_13

MRQGAPARGCRWFVVWALLGLTLGVLVASAAPSSPGTPGVAAATQAANGGPATPAPPALGAAPTGDPKPK

KNKKPKNPTPPRPAGDNATVAAGHATLREHLRDIKAENTDANFYVCPPPTGATVVQFEQPRRCPTRPEGQ

NYTEGIAVVFKENIAPYKFKATMYYKDVTVSQVWFGHRYSQFMGIFEDRAPVPFEEVIDKINAKGVCRST

AKYVRNNLETTAFHRDDHETDMELKPANAATRTSRGWHTTDLKYNPSRVEAFHRYGTTVNCIVEEVDARS

VYPYDEFVLATGDFVYMSPFYGYREGSHTEHTSYAADRFKQVDGFYARDLTTKARATAPTTRNLLTTPKF

TVAWDWVPKRPSVCTMTKWQEVDEMLRSEYGGSFRFSSDAISTTFTTNLTEYPLSRVDLGDCIGKDARDA

MDRIFARRYNATHIKVGQPQYYLANGGFLIAYQPLLSNTLAELYVREHLREQSRKPPNPTPPPPGASANA

SVERIKTTSSIEFARLQFTYNHIQRHVNDMLGRVAIAWCELQNHELTLWNEARKLNPNAIASATVGRRVS

ARMLGDVMAVSTCVPVAADNVIVQNSMRISSRPGACYSRPLVSFRYEDQGPLVEGQLGENNELRLTRDAI

EPCTVGHRRYFTFGGGYVYFEEYAYSHQLSRADITTVSTFIDLNITMLEDHEFVPLEVYTRHEIKDSGLL

DYTEVQRRNQLHDLRFADIDTVIHADANAAMFAGLGAFFEGMGDLGRAVGKVVMGIVGGVVSAVSGVSSF

MSNPFGALAVGLLVLAGLAAAFFAFRYVMRLQSNPMKALYPLTTKELKNPTNPDASGEGEEGGDFDEAKL

AEAREMIRYMALVSAMERTEHKAKKKGTSALLSAKVTDMVMRKRRNTNYTQVPNKDGDADEDDL

>HS1GPB_14 MRQGAPARGRRWFVVWALLGLTLGVLVASAAPSSPGTPGVAAATQAANGGPATPAPPAPGPAPTGDTKPK

KNKKPKNPPPPRPAGDNATVAAGHATLREHLRDIKAENTDANFYVCPPPTGATVVQFEQPRRCPTRPEGQ

NYTEGIAVVFKENIAPYKFKATMYYKDVTVSQVWFGHRYSQFMGIFEDRAPVPFEEVIDKINAKGVCRST

AKYVRNNLETTAFHRDDHETDMELKPANAATRTSRGWHTTDLKYNPSRVEAFHRYGTTVNCIVEEVDARS

VYPYDEFVLATGDFVYMSPFYGYREGSHTEHTSYAADRFKQVDGFYARDLTTKARATAPTTRNLLTTPKF

TVAWDWVPKRPSVCTMTKWQEVDEMLRSEYGGSFRFSSDAISTTFTTNLTEYPLSRVDLGDCIGKDARDA

MDRIFARRYNATHIKVGQPQYYLANGGFLIAYQPLLSNTLAELYVREHLREQSRKPPNPTPPPPGASANA

SVERIKTTSSIEFARLQFTYNHIQRHVNDMLGRVAIAWCELQNHELTLWNEARKLNPNAIASATVGRRVS

ARMLGDVMAISTCVPVAADNVIVQNSMRISSRPGACYSRPLVSFRYEDQGPLVEGQLGENNELRLTRDAI

EPCTVGHRRYFTFGGGYVYFEEYAYSHQLSRADITTVSTFIDLNITMLEDHEFVPLEVYTRHEIKDSGLL

DYTEVQRRNQLHDLRFADIDTVIHADANAAMFAGLGAFFEGMGDLGRAVGKVVMGIVGGVVSAVSGVSSF

MSNPFGALAVGLLVLAGLAAAFFAFRYVMRLQSNPMKALYPLTTKELKNPTNPDASGEGEEGGDFDEAKL

AEAREMIRYMALVSAMERTEHKAKKKGTSALLSAKVTDMVMRKRRNTNYTQVPNKDGDADEDDL

>HS1GPB_15

MRQGAPARGRRWFVVWALLGLTLGVLVASAAPSSPGTPGGAAATQAADGGPATPAPPALGAAPTGDPKPK

KNKKPKNPTPPRPAGDNATVAAGHATLREHLRDIKAENTDANFYVCPPPTGATVVQFEQPRRCPTRPEGQ

NYTEGIAVVFKENIAPYKFKATMYYKDVTVSQVWFGHRYSQFMGIFEDRAPVPFEEVIDKINAKGVCRST

AKYVRNNLETTAFHRDDHETDMELKPANAATRTSRGWHTTDLKYNPSRVEAFHRYGTTVNCIVEEVDARS

VYPYDEFVLATGDFVYMSPFYGYREGSHTEHTSYAADRFKQVDGFYARDLTTKARATAPTTRNLLTTPKF

TVAWDWVPKRPSVCTMTKWQEVDEMLRSEYGGSFRFSSDAISTTFTTNLTEYPLSRVDLGDCIGKDARDA

MDRIFARRYNATHIKVGQPQYYLANGGFLIAYQPLLSNTLAELYVREHLREQSRKPPNPTPPPPGASANA

SVERIKTTSSIEFARLQFTYNHIQRHVNDMLGRVAIAWCELQNHELTLWNEARKLNPNAIASATVGRRVS

ARMLGDVMAVSTCVPVAADNVIVQNSMRISSRPGACYSRPLVSFRYEDQGPLVEGQLGENNELRLTRDAI

EPCTVGHRRYFTFGGGYVYFEEYAYSHQLSRADITTVSTFIDLNITMLEDHEFVPLEVYTRHEIKDSGLL

DYTEVQRRNQLHDLRFADIDTVIHADANAAMFAGLGAFFEGMGDLGRAVGKVVMGIVGGVVSAVSGVSSF

MSNPFGALAVGLLVLAGLAAAFFAFRYVMRLQSNPMKALYPLTTKELKNPTNPDASGEGEEGGDFDEAKL

AEAREMIRYMALVSAMERTEHKAKKKGTSALLSAKVTDMVMRKRRNTNYTQVPNKDGDADEDDL

>HS1GPB_16

MRQGAPARGCRWFVVWVLLGLTLGVLVASAAPSSPGTPGVAAATQAANGGPATPAPPALGAAPTGDPKPK

KNKKPKNPTPPRPAGDNATVAAGHATLREHLRDIKAENTDANFYVCPPPTGATVVQFEQPRRCPTRPEGQ

NYTEGIAVVFKENIAPYKFKATMYYKDVTVSQVWFGHRYSQFMGIFEDRAPVPFEEVIDKINAKGVCRST

AKYVRNNLETTAFHRDDHETDMELKPANAATRTSRGWHTTDLKYNPSRVEAFHRYGTTVNCIVEEVDARS

VYPYDEFVLATGDFVYMSPFYGYREGSHTEHTSYAADRFKQVDGFYARDLTTKARATAPTTRNLLTTPKF

TVAWDWVPKRPSVCTMTKWQEVDEMLRSEYGGSFRFSSDAISTTFTTNLTEYPLSRVDLGDCIGKDARDA

MDRIFARRYNATHIKVGQPQYYLANGGFLIAYQPLLSNTLAELYVREHLREQSRKPPNPTPPPPGASANA

SVERIKTTSSIEFARLQFTYNHIQRHVNDMLGRVAIAWCELQNHELTLWNEARKLNPNAIASATVGRRVS

ARMLGDVMAVSTCVPVAADNVIVQNSMRISSRPGACYSRPLVSFRYEDQGPLVEGQLGENNELRLTRDAI

EPCTVGHRRYFTFGGGYVYFEEYAYSHQLSRADITTVSTFIDLNITMLEDHEFVPLEVYTRHEIKDSGLL

DYTEVQRRNQLHDLRFADIDTVIHADANAAMFAGLGAFFEGMGDLGRAVGKVVMGIVGGVVSAVSGVSSF

MSNPFGALAVGLLVLAGLAAAFFAFRYVMRLQSNPMKALYPLTTKELKNPTNPDASGEGEEGGDFDEAKL

AEAREMIRYMALVSAMERTEHKAKKKGTSALLSAKVTDMVMRKRRNTNYTQVPNKDGDADEDDL

>HS1GPB_17

MRQGAPARGCRWFVVWALLGLTLGVLVASAAPSSPGTPGVAAATQAANGGPATPAPPAPGPAPTGDTKPK

KNKKPKNPPPPRPAGDNATVAAGHATLREHLRDIKAENTDANFYVCPPPTGATVVQFEQPRRCPTRPEGQ

NYTEGIAVVFKENIAPYKFKATMYYKDVTVSQVWFGHRYSQFMGIFEDRAPVPFEEVIDKINAKGVCRST

AKYVRNNLETTAFHRDDHETDMELKPANAATRTSRGWHTTDLKYNPSRVEAFHRYGTTVNCIVEEVDARS

VYPYDEFVLATGDFVYMSPFYGYREGSHTEHTSYAADRFKQVDGFYARDLTTKARATAPTTRNLLTTPKF

TVAWDWVPKRPSVCTMTKWQEVDEMLRSEYGGSFRFSSDAISTTFTTNLTEYPLSRVDLGDCIGKDARDA

MDRIFARRYNATHIKVGQPQYYLANGGFLIAYQPLLSNTLAELYVREHLREQSRKPPNPTPPPPGASANA

SVERIKTTSSIEFARLQFTYNHIQRHVNDMLGRVAIAWCELQNHELTLWNEARKLNPNAIASATVGRRVS

ARMLGDVMAVSTCVPVAADNVIVQNSMRISSRPGACYSRPLVSFRYEDQGPLVEGQLGENNELRLTRDAI

EPCTVGHRRYFTFGGGYVYFEEYAYSHQLSRADITTVSTFIDLNITMLEDHEFVPLEVYTRHEIKDSGLL

DYTEVQRRNQLHDLRFADIDTVIHADANAAMFAGLGAFFEGMGDLGRAVGKVVMGIVGGVVSAVSGVSSF

MSNPFGALAVGLLVLAGLAAAFFAFRYVMRLQSNPMKALYPLTTKELKNPTNPDASGEGEEGGDFDEAKL

AEAREMIRYMALVSAMERTEHKAKKKGTSALLSAKVTDMVMRKRRNTNYTQVPNKDGDADEDDL

>HS1GPB_18

MRQGAPARGCRWFVVWALLGLTLGVLVASAAPSSPGTPGVAAATQAANGGPATPAPPAPGPAPTGDTKPQ

KNKKPKPPXXPRPAGDNATVAAGHATLREHLRDIKAENTDANFYVCPPPTGATVVQFEQPRRCPTRPEGQ

NYTEGIAVVFKENIAPYKFKATMYYKDVTVSQVWFGHRYSQFMGIFEDRAPVPFEEVIDKINAKGVCRST

AKYVRNNLETTAFHRDDHETDMELKPANAATRTSRGWHTTDLKYNPSRVEAFHRYGTTVNCIVEEVDARS

VYPYDEFVLATGDFVYMSPFYGYREGSHTEHTSYAADRFKQVDGFYARDLTTKARATAPTTRNLLTTPKF

TVAWDWVPKRPSVCTMTKWQEVDEMLRSEYGGSFRFSSDAISTTFTTNLTEYPLSRVDLGDCIGKDARDA

MDRIFARRYNATHIKVGQPQYYLANGGFLIAYQPLLSNTLAELYVREHLREQSRKPPNPTPPPPGASANA

SVERIKTTSSIEFARLQFTYNHIQRHVNDMLGRVAIAWCELQNHELTLWNEARKLNPNAIASATVGRRVS

ARMLGDVMAVSTCVPVAADNVIVQNSMRISSRPGACYSRPLVSFRYEDQGPLVEGQLGENNELRLTRDAI

EPCTVGHRRYFTFGGGYVYFEEYAYSHQLSRADITTVSTFIDLNITMLEDHEFVPLEVYTRHEIKDSGLL

DYTEVQRRNQLHDLRFADIDTVIHADANAAMFAGLGAFFEGMGDLGRAVGKVVMGIVGGVVSAVSGVSSF

MSNPFGALAVGLLVLAGLAAAFFAFRYVMRLQSNPMKALYPLTTKELKNPTNPDASGEGEEGGDFDEAKL

AEAREMIRYMALVSAMERTEHKAKKKGTSALLSAKVTDMVMRKRRNTNYTQVPNKDGDADEDDL

>HS1GPB_19 MRQGAPARGCRWFVVWALLGLTLGVLVASAAPSSPGTPGVAAATQAANGGPATPAPPALGAAPTGDPKPK

KNKKPKNPTPPRPAGDNATVAAGHATLREHLRDIKAENTDANFYVCPPPTGATVVQFEQPRRCPTRPEGQ

NYTEGIAVVFKENIAPYKFKATMYYKDVTVSQVWFGHRYSQFMGIFEDRAPVPFEEVIDKINAKGVCRST

AKYVRNNLETTAFHRDDHETDMELKPANAATRTSRGWHTTDLKYNPSRVEAFHRYGTTVNCIVEEVDARS

VYPYDEFVLATGDFVYMSPFYGYREGSHTEHTSYAADRFKQVDGFYARDLTTKARATAPTTRNLLTTPKF

TVAWDWVPKRPSVCTMTKWQEVDEMLRSEYGGSFRFSSDAISTTFTTNLTEYPLSRVDLGDCIGKDARDA

MDRIFARRYNATHIKVGQPQYYLANGGFLIAYQPLLSNTLAELYVREHLREQSRKPPNPTPPPPGASANA

SVERIKTTSSIEFARLQFTYNHIQRHVNDMLGRVAIAWCELQNHELTLWNEARKLNPNAIASATVGRRVS

ARMLGDVMAVSTCVPVAADNVIVQNSMRISSRPGACYSRPLVSFRYEDQGPLVEGQLGENNELRLTRDAI

EPCTVGHRRYFTFGGGYVYFEESAYSHQLSRADITTVSTFIDLNITMLEDHEFVPLEVYTRHEIKDSGLL

DYTEVQRRNQLHDLRFADIDTVIHADANAAMFAGLGAFFEGMGDLGRAVGKVVMGIVGGVVSAVSGVSSF

MSNPFGALAVGLLVLAGLAAAFFAFRYVMRLQSNPMKALYPLTTKELKNPTNPDASGEGEEGGDFDEAKL

AEAREMIRYMALVSAMERTEHKAKKKGTSALLSAKVTDMVMRKRRNTNYTQVPNKDGDADEDDL

>HS1GPB_20 MRQGAPARGCRWFVVWALLGLTLGVLVASAAPSSPGTPGVAAATQAANGGPATPAPPAPGPAPTGDTKPK

KNKKPKNPPPPRPAGDNATVAAGHATLREHLRDIKAENTDANFYVCPPPTGATVVQFEQPRRCPTRPEGQ

NYTEGIAVVFKENIAPYKFKATMYYKDVTVSQVWFGHRYSQFMGIFEDRAPVPFEEVIDKINAKGVCRST

AKYVRNNLETTAFHRDDHETDMELKPANAATRTSRGWHTTDLKYNPSRVEAFHRYGTTVNCIVEEVDARS

VYPYDEFVLATGDFVYMSPFYGYREGSHTEHTSYAADRFKQVDGFYARDLTTKARATAPTTRNLLTTPKF

TVAWDWVPKRPSVCTMTKWQEVDEMLRSEYGGSFRFSSDAISTTFTTNLTEYPLSRVDLGDCIGKDARDA

MDRIFARRYNATHIKVGQPQYYLANGGFLIAYQPLLSNTLAELYVREHLREQSRKPPNPTPPPPGASANA

SVERIKTTSSIEFARLQFTYNHIQRHVNDMLGRVAIAWCELQNHELTLWNEARKLNPNAIASATVGRRVS

ARMLGDVMAVSTCVPVAADNVIVQNSMRISSRPGACYSRPLVSFRYEDQGPLVEGQLGENNELRLTRDAI

EPCTVGHRRYFTFGGGYVYFEEYAYSHQLSRADITTVSTFIDLNITMLEDHEFVPLEVYTRHEIKDSGLL

DYTEVQRRNQLHDLRFADIDTVIHADANAAMFAGLGAFFEGMGDLGRAVGKVVMGIVGGVVSAVSGVSSF

MSNPFGALAVGLLVLAGLAAAFFAFRYVMRLQSNPMKALYPLTTKELKNPTNPDASGEGEEGGDFDEAKL

AEAREMIRYMALVSAMERTEHKAKKKGTSALLSAKVTDMVMRKRRNTNYTQVPNKDGDADEDNL

>HS2GPB_21

MRGGGLICALVVGALVAAVASAAPAAPAAPRASGGVAATVAANGGPASRPPPVPSPATTKARKRKTKKPP

KRPEATPPPDANATVAAGHATLRAHLREIKVENADAQFYVCPPPTGATVVQFEQPRRCPTRPEGQNYTEG

IAVVFKENIAPYKFKATMYYKDVTVSQVWFGHRYSQFMGIFEDRAPVPFEEVIDKINTKGVCRSTAKYVR

NNMETTAFHRDDHETDMELKPAKVATRTSRGWHTTDLKYNPSRVEAFHRYGTTVNCIVEEVDARSVYPYD

EFVLATGDFVYMSPFYGYREGSHTEHTSYAADRFKQVDGFYARDLTTKARATSPTTRNLLTTPKFTVAWD

WVPKRPAVCTMTKWQEVDEMLRAEYGGSFRFSSDAISTTFTTNLTEYSLSRVDLGDCIGRDAREAIDRMF

ARKYNATHIKVGQPQYYLATGGFLIAYQPLLSNTLAELYVREYMREQDRKPRNATPAPLREAPSANASVE

RIKTTSSIEFARLQFTYNHIQRHVNDMLGRIAVAWCELQNHELTLWNEARKLNPNAIASATVGRRVSARM

LGDVMAVSTCVPVAPDNVIVQNSMRVSSRPGTCYSRPLVSFRYEDQGPLIEGQLGENNELRLTRDALEPC

TVGHRRYFIFGGGYVYFEEYAYSHQLSRADVTTVSTFIDLNITMLEDHEFVPLEVYTRHEIKDSGLLDYT

EVQRRNQLHDLRFADIDTVIRADANAAMFAGLCAFFEGMGDLGRAVGKVVMGVVGGVVSAVSGVSSFMSN

PFGALAVGLLVLAGLVAAFFAFRYVLQLQRNPMKALYPLTTKELKTSDPGGVGGEGEEGAEGGGFDEAKL

AEAREMIRYMALVSAMERTEHKARKKGTSALLSSKVTNMVLRKRNKARYSPLHNEDEAGDEDEL

> HS2GPB_22

MRGGGLICALVVGALVAAVASAAPAAPAAPRASGGVAATVAANGGPASRPPPVPSPATTKARKRKTKKPP

KRPEATPPPDANATVAAGHATLRAHLREIKVENADAQFYVCPPPTGATVVQFEQPRRCPTRPEGQNYTEG

IAVVFKENIAPYKFKATMYYKDVTVSQVWFGHRYSQFMGIFEDRAPVPFEEVIDKINAKGVCRSTAKYVR

NNMETTAFHRDDHETDMELKPAKVATRTSRGWHTTDLKYNPSRVEAFHRYGTTVNCIVEEVDARSVYPYD

EFVLATGDFVYMSPFYGYREGSHTEHTSYAADRFKQVDGFYARDLTTKARATSPTTRNLLTTPKFTVAWD

WVPKRPAVCTMTKWQEVDEMLRAEYGGSFRFSSDAISTTFTTNLTEYSLSRVDLGDCIGRDAREAIDRMF

ARKYNATHIKVGQPQYYLATGGFLIAYQPLLSNTLAELYVREYMREQDRKPRNATPAPLREAPSANASVE

RIKTTSSIEFARLQFTYNHIQRHVNDMLGRIAVAWCELQNHELTLWNEARKLNPNAIASATVGRRVSARM

LGDVMAVSTCVPVAPDNVIVQNSMRVSSRPGTCYSRPLVSFRYEDQGPLIEGQLGENNELRLTRDALEPC

TVGHRRYFIFGGGYVYFEEYAYSHQLSRADVTTVSTFIDLNITMLEDHEFVPLEVYTRHEIKDSGLLDYT

EVQRRNQLHDLRFADIDTVIRADANAAMFAGLCAFFEGMGDLGRAVGKVVMGVVGGVVSAVSGVSSFMSN

PFGALAVGLLVLAGLVAAFFAFRYVLQLQRNPMKALYPLTTKELKTSDPGGVGGEGEEGAEGGGFDEAKL

AEAREMIRYMALVSAMERTEHKARKKGTSALLSSKVTNMVLRKRNKARYSPLHNEDEAGDEDEL

>HS2GPB_23 MRGGGLICALVVGALVAAVASAAPAAPAAPRASGGVAATVAANGGPASRPPPVPSPATTKARKRKTKKPP

KRPEATPPPDANATVAAGHATLRAHLREIKVENADAQFYVCPPPTGATVVQFEQPRRCPTRPEGQNYTEG

IAVVFKENIAPYKFKATMYYKDVTVSQVWFGHRYSQFMGIFEDRAPVPFEEVIDKINAKGVCRSTAKYVR

NNMETTAFHRDDHETDMELKPAKVATRTSRGWHTTDLKYNPSRVEAFHRYGTTVNCIVEEVDARSVYPYD

EFVLATGDFVYMSPFYGYREGSHTEHTSYAADRFKQVDGFYARDLTTKARATSPTTRNLLTTPKFTVAWD

WVPKRPAVCTMTKWQEVDEMLRAEYGGSFRFSSDAISTTFTTNLTQYSLSRVDLGDCIGRDAREAIDRMF

ARKYNATHIKVGQPQYYLATGGFLIAYQPLLSNTLAELYVREYMREQDRKPRNATPAPLREAPSANASVE

RIKTTSSIEFARLQFTYNHIQRHVNDMLGRIAVAWCELQNHELTLWNEARKLNPNAIASATVGRRVSARM

LGDVMAVSTCVPVAPDNVIVQNSMRVSSRPGTCYSRPLVSFRYEDQGPLIEGQLGENNELRLTRDALEPC

TVGHRRYFIFGGGYVYFEEYAYSHQLSRADVTTVSTFIDLNITMLEDHEFVPLEVYTRHEIKDSGLLDYT

EVQRRNQLHDLRFADIDTVIRADANAAMFAGLCAFFEGMGDLGRAVGKVVMGVVGGVVSAVSGVSSFMSN

PFGALAVGLLVLAGLVAAFFAFRYVLQLQRNPMKALYPLTTKELKTSDPGGVGGEGEEGAEGGGFDEAKL

AEAREMIRYMALVSAMERTEHKARKKGTSALLSSKVTNMVLRKRNKARYSPLHNEDEAGDEDEL

>HS2GPB_24

MRGGGLICALVVGALVAAVASAAPAAPAAPRASGGVAATVAANGGPASRPPPVPSPATTKARKRKTKKPP

KRPEATPPPDANATVAAGHATLRAHLREIKVENADAQFYVCPPPTGATVVQFEQPRRCPTRPEGQNYTEG

IAVVFKENIAPYKFKATMYYKDVTVSQVWFGHRYSQFMGIFEDRAPVPFEEVIDKINAKGVCRSTAKYVR

NNMETTAFHRDDHETDMELKPAKVATRTSRGWHTTDLKYNPSRVEAFHRYGTTVNCIVEEVDARSVYPYD

EFVLATGDFVYMSPFYGYREGSHTEHTSYAADRFKQVDGFYARDLTTKARATSPTTRNLLTTPKFTVAWD

WVPKRPVVCTMTKWQEVDEMLRAEYGGSFRFSSDAISTTFTTNLTQYSLSRVDLGDCIGRDAREAIDRMF

ARKYNATHIKVGQPQYYLATGGFLIAYQPLLSNTLAELYVREYMREQDRKPRNATPAPLREAPSANASVE

RIKTTSSIEFARLQFTYNHIQRHVNDMLGRIAVAWCELQNHELTLWNEARKLNPNAIASATVGRRVSARM

LGDVMAVSTCVPVAPDNVIVQNSMRVSSRPGTCYSRPLVSFRYEDQGPLIEGQLGENNELRLTRDALEPC

TVGHRRYFIFGGGYVYFEEYAYSHQLSRADVTTVSTFIDLNITMLEDHEFVPLEVYTRHEIKDSGLLDYT

EVQRRNQLHDLRFADIDTVIRADANAAMFAGLCAFFEGMGDLGRAVGKVVMGVVGGVVSAVSGVSSFMSN

PFGALAVGLLVLAGLVAAFFAFRYVLQLQRNPMKALYPLTTKELKTSDPGGVGGEGEEGAEGGGFDEAKL

AEAREMIRYMALVSAMERTEHKARKKGTSALLSSKVTNMVLRKRNKARYSPLHNEDEAGDEDEL

>HS2GPB_25 MRGGGLICALVVGALVAAVASAAPAAPAAPRASGGVAATVAANGGPASRPPPVPSPATTRARKRKTKKPP

ERPEATPPPDANATVAAGHATLRAHLREIKVENADAQFYVCPPPTGATVVQFEQPRRCPTRPEGQNYTEG

IAVVFKENIAPYKFKATMYYKDVTVSQVWFGHRYSQFMGIFEDRAPVPFEEVIDKINAKGVCRSTAKYVR

NNMETTAFHRDDHETDMELKPAKVATRTSRGWHTTDLKYNPSRVEAFHRYGTTVNCIVEEVDARSVYPYD

EFVLATGDFVYMSPFYGYREGSHTEHTSYAADRFKQVDGFYARDLTTKARATSPTTRNLLTTPKFTVAWD

WVPKRPAVCTMTKWQEVDEMLRAEYGGSFRFSSDAISTTFTTNLTQYSLSRVDLGDCIGRDAREAIDRMF

ARKYNATHIKVGQPQYYLATGGFLIAYQPLLSNTLAELYVREYMREQDRKPRNATPAPLREAPSANASVE

RIKTTSSIEFARLQFTYNHIQRHVNDMLGRIAVAWCELQNHELTLWNEARKLNPNAIASATVGRRVSARM

LGDVMAVSTCVPVAPDNVIVQNSMRVSSRPGTCYSRPLVSFRYEDQGPLIEGQLGENNELRLTRDALEPC

TVGHRRYFIFGGGYVYFEEYAYSHQLSRADVTTVSTFIDLNITMLEDHEFVPLEVYTRHEIKDSGLLDYT

EVQRRNQLHDLRFADIDTVIRADANAAMFAGLCAFFEGMGDLGRAVGKVVMGVVGGVVSAVSGVSSFMSN

PFGALAVGLLVLAGLVAAFFAFRYVLQLQRNPMKALYPLTTKELKTSDPGGVGGEGEEGAEGGGFDEAKL

AEAREMIRYMALVSAMERTEHKARKKGTSALLSSKVTNMVLRKRNKARYSPLHNEDEAGDEDEL

>HS2GPB_26

MRGGGLICALVVGALVAAVASAAPAAPAAPRASGGVAATVAANGGPASRPPPVPSPATTRARKRKTKKPP

KRPEATPPPDANATVAAGHATLRAHLREIKVENADAQFYVCPPPTGATVVQFEQPRRCPTRPEGQNYTEG

IAVVFKENIAPYKFKATMYYKDVTVSQVWFGHRYSQFMGIFEDRAPVPFEEVIDKINAKGVCRSTAKYVR

NNMETTAFHRDDHETDMELKPAKVATRTSRGWHTTDLKYNPSRVEAFHRYGTTVNCIVEEVDARSVYPYD

EFVLATGDFVYMSPFYGYREGSHTEHTSYAADRFKQVDGFYARDLTTKAQATSPTTRNLLTTPKFTVAWD

WVPKRPAVCTMTKWQEVDEMLRAEYGGSFRFSSDAISTTFTTNLTQYSLSRVDLGDCIGRDAREAIDRMF

ARKYNATHIKVGQPQYYLATGGFLIAYQPLLSNTLAELYVREYMREQDRKPRNATPAPLREAPSANASVE

RIKTTSSIEFARLQFTYNHIQRHVNDMLGRIAVAWCELQNHELTLWNEARKLNPNAIASATVGRRVSARM

LGDVMAVSTCVPVAPDNVIVQNSMRVSSRPGTCYSRPLVSFRYEDQGPLIEGQLGENNELRLTRDALEPC

TVGHRRYFIFGGGYVYFEEYAYSHQLSRADVTTVSTFIDLNITMLEDHEFVPLEVYTRHEIKDSGLLDYT

EVQRRNQLHDLRFADIDTVIRADANAAMFAGLCAFFEGMGDLGRAVGKVVMGVVGGVVSAVSGVSSFMSN

PFGALAVGLLVLAGLVAAFFAFRYVLQLQRNPMKALYPLTTKELKTSDPGGVGGEGEEGAEGGGFDEAKL

AEAREMIRYMALVSAMERTEHKARKKGTSALLSSKVTNMVLRKRNKARYSPLHNEDEAGDEDEL

>HS2GPB_27

MRGGGLICALVVGALVAAVASAAPAAPAAPRASGGVAATVAANGGPASRPPPVPSPATTKARKRKTKKPP

KRPEATPPPDANATVAAGHATLRAHLREIKVENADAQFYVCPPPTGATVVQFEQPRRCPTRPEGQNYTEG

IAVVFKENIAPYKFKATMYYKDVTVSQVWFGHRYSQFMGIFEDRAPVPFEEVIDKINAKGVCRSTAKYVR

NNMETTAFHRDDHETDMELKPAKVATRTSRGWHTTDLKYNPSRVEAFHRYGTTVNCIVEEVDARSVYPYD

EFVLATGDFVYMSPFYGYREGSHTEHTSYAADRFKQVDGFYARDLTTKARATSPTTRNLLTTPKFTVAWD

WVPKRPAVCTMTKWQEVDEMLRAEYGGSFRFSSDAISTTFTTNLTQYSLSRVDLGDCIGRDAREAIDRMF

ARKYNATHIKVGQPQYYLATGGFLIAYQPLLSNTLAELYVREYMREQDRKPRNATPAPLREAPSANASVE

RIKTTSSIEFARLQFTYNHIQRHVNDMLGRIAVAWCELQNHELTLWNEARKLNPNAIASATVGRRVSARM

LGDVMAVSTCVPVAPDNVIVQNSMRVSSRPGTCYSRPLVSFRYEDQGPLIEGQLGENNELRLTRDALEPC

TVGHRRYFIFGGGYVYFEEYAYSHQLSRADVTTVSTFIDLNITMLEDHEFVPLGVYTRHEIKDSGLLDYT

EVQRRNQLHDLRFADIDTVIRADANAAMFAGLCAFFEGMGDLGRAVGKVVMGVVGGVVSAVSGVSSFMSN

PFGALAVGLLVLAGLVAAFFAFRYVLQLQRNPMKALYPLTTKELKTSDPGGVGGEGEEGAEGGGFDEAKL

AEAREMIRYMALVSAMERTEHKARKKGTSALLSSKVTNMVLRKRNKARYSPLHNEDEAGDEDEL

>HS2GPB_28

MRGGGLICALVVGALVAAVASAAPAAPAAPRASGGVAATVAANGGPASRPPPVPSPATTKARKRKTKKPP

ERPEATPPPDANATVAAGHATLRAHLREIKVENADAQFYVCPPPTGATVVQFEQPRRCPTRPEGQNYTEG

IAVVFKENIAPYKFKATMYYKDVTVSQVWFGHRYSQFMGIFEDRAPVPFEEVIDKINAKGVCRSTAKYVR

NNMETTAFHRDDHETDMELKPAKVATRTSRGWHTTDLKYNPSRVEAFHRYGTTVNCIVEEVDARSVYPYD

EFVLATGDFVYMSPFYGYREGSHTEHTSYAADRFKQVDGFYARDLTTKARATSPTTRNLLTTPKFTVAWD

WVPKRPAVCTMTKWQEVDEMLRAEYGGSFRFSSDAISTTFTTNLTQYSLSRVDLGDCIGRDAREAIDRMF

ARKYNATHIKVGQPQYYLATGGFLIAYQPLLSNTLAELYVREYMREQDRKPRNATPAPLREAPSANASVE

RIKTTSSIEFARLQFTYNHIQRHVNDMLGRIAVAWCELQNHELTLWNEARKLNPNAIASATVGRRVSARM

LGDVMAVSTCVPVAPDNVIVQNSMRVSSRPGTCYSRPLVSFRYEDQGPLIEGQLGENNELRLTRDALEPC

TVGHRRYFIFGGGYVYFEEYAYSHQLSRADVTTVSTFIDLNITMLEDHEIVPLEVYTRHEIKDSGLLDYT

EVQRRNQLHDLRFADIDTVIRADANAAMFAGLCAFFEGMGDLGRAVGKVVMGVVGGVVSAVSGVSSFMSN

PFGALAVGLLVLAGLVAAFFAFRYVLQLQRNPMKALYPLTTKELKTSDPGGVGGEGEEGAEGGGFDEAKL

AEAREMIRYMALVSAMERTEHKARKKGTSALLSSKVTNMVLRKRNKARYSPLHNEDEAGDEDEL

>HS2GPB_29 MRGGGLICALVVGALVAAVASAAPAAPAAPRASGGVAATVAANGGPASRPPPVPSPATTKARKRKTKKPP

KRPEATPPPDANATVAAGHATLRAHLREIKVENADAQFYVCPPPTGATVVQFEQPRRCPTRPEGQNYTEG

IAVVFKENIAPYKFKATMYYKDVTVSQVWFGHRYSQFMGIFEDRAPVPFEEVIDKINAKGVCRSTAKYVR

NNMETTAFHRDDHETDMELKPAKVATRTSRGWHTTDLKYNPSRVEAFHRYGTTVNCIVEEVDARSVYPYD

EFVLATGDFVYMSPFYGYREGSHTEHTSYAADRFKQVDGFYARDLTTKARATSPTTRNLLTTPKFTVAWD

WVPKRPAVCTMTKWQEVDEMLRAEYGGSFRFSSDAISTTFTTNLTQYSLSRVDLGDCIGRDAREAIDRMF

ARKYNATHIKVGQPQYYLATGGFLIAYQPLLSNTLAELYVREYMREQDRKPRNATPAPLREAPSANASVE

RIKTTSSIEFARLQFTYNHIQRHVNDMLGRIAVAWCELQNHELTLWNEARKLNPNAIASATVGRRVSARM

LGDVMAVSTCVPVAPDNVIVQNSMRVSSRPGTCYSRPLVSFRYEDQGPLIEGQLGENNELRLTRDALEPC

TVGHRRYFIFGGGYVYFEEYAYSHQLSRADVTTVSTFIDLNITMLEDHEFVPLGVYTRHEIKDSGLLDYT

EVQRRNQLHDLRFADIDTVIRADANAAMFAGLCAFFEGMGDLGRAVGKVVMGVVGGVVSAVSGVSSFMSN

PFGALAVGLLVLAGLVAAFFAFRYVLQLQRNPMKALYPLTTKELKTSDPGGVGGEGEEGAEGGGFDEAKL

AEAQEMIRYMALVSAMERTEHKARKKGTSALLSSKVTNMVLRKRNKARYSPLHNEDEAGDEDEL

>HS2GPB_30

MRGGGLICALVVGALVAAVASAAPAAPAAPRASGGVAATVAANGGPASRPPPVPSPATTKARKRKTKKPP

KRPEATPPPDANATVAAGHATLRAHLREIKVENADAQFYVCPPPTGATVVQFEQPRRCPTRPEGQNYTEG

IAVVFKENIAPYKFKATMYYKDVTVSQVWFGHRYSQFMGIFEDRAPVPFEEVIDKINAKGVCRSTAKYVR

NNMETTAFHRDDHETDMELKPAKVATRTSRGWHTTDLKYNPSRVEAFHRYGTTVTCIVEEVDARSVYPYD

EFVLATGDFVYMSPFYGYREGSHTEHTSYAADRFKQVDGFYARDLTTKARATSPTTRNLLTTPKFTVAWD

WVPKRPAVCTMTKWQEVDEMLRAEYGGSFRFSSDAISTTFTTNLTQYSLSRVDLGDCIGRDAREAIDRMF

ARKYNATHIKVGQPQYYLATGGFLIAYQPLLSNTLAELYVREYMREQDRKPRNATPAPLREAPSANASVE

RIKTTSSIEFARLQFTYNHIQRHVNDMLGRIAVAWCELQNHELTLWNEARKLNPNAIASATVGRRVSARM

LGDVMAVSTCVPVAPDNVIVQNSMRVSSRPGTCYSRPLVSFRYEDQGPLIEGQLGENNELRLTRDALEPC

TVGHRRYFIFGGGYVYFEEYAYSHQLSRADVTTVSTFIDLNITMLEDHEFVPLGVYTRHEIKDSGLLDYT

EVQRRNQLHDLRFADIDTVIRADANAAMFAGLCAFFEGMGDLGRAVGKVVMGVVGGVVSAVSGVSSFMSN

PFGALAVGLLVLAGLVAAFFAFRYVLQLQRNPMKALYPLTTKELKTSDPGGVGGEGEEGAEGGGFDEAKL

AEAREMIRYMALVSAMERTEHKARKKGTSALLSSKVTNMVLRKRNKARYSPLHNEDEAGDEDEL

>HS2GPB_31

MRGGGLICALVVGALVAAVASAAPAAPAAPRASGGVAATVAANGGPASRPPPVPSPATTKARKRKTKKPP

KRPEATPPPDANATVAAGHATLRAHLREIKVENADAQFYVCPPPTGATVVQFEQPRRCPTRPEGQNYTEG

IAVVFKENIAPYKFKATMYYKDVTVSQVWFGHRYSQFMGIFEDRAPVPFEEVIDKINAKGVCRSTAKYVR

NNMETTAFHRDDHETDMELKPAKVATRTSRGWHTTDLKYNPSRVEAFHRYGTTVNCIVEEVDARSVYPYD

EFVLATGDFVYMSPFYGYREGSHTEHTSYAADRFKQVDGFYARDLTTKARATSPTTRNLLTTPKFTVAWD

WVPKRPAVCTMTKWQEVDEMLRAEYGGSFRFSSDAISTTFTTNLTQYSLSRVDLGDCIGRDAREAIDRMF

ARKYNATHIKVGQPQYYLATGGFLIAYQPLLSNTLAELYVREYMREQDRKPRNATPAPLREAPSANASVE

RIKTTSSIEFARLQFTYNHIQRHVNDMLGRIAVAWCELQNHELTLWNEARKLNPNAIASATVGRRVSARM

LGDVMAVSTCVPVAPDNVIVQNSMRVSSRPGTCYSRPLVSFRYEDQGPLIEGQLGENNELRLTRDALEPC

TVGHRRYFIFGGGYVYFEEYAYSHQLSRADVTTVSTFIDLNITMLEDHEFVPLGVYTRHEIKDSGLLDYT

EVQRRNQLHDLRFADIDTVIRADANAAMFAGLCAFFEGMGDLGRAVGKVVMGVVGGVVSAVSGVSSFMSN

PFGALAVGLLVLAGLVAAFFAFRYVLQLQRNPMKALYPLTTKEPKTSDPGGVGGEGEEGAEGGGFDEAKL

AEAREMIRYMALVSAMERTEHKARKKGTSALLSSKVTNMVLRKRNKARYSPLHNEDEAGDEDEL

>HS2GPB_32 MRGGGLICALVVGALVAAVASAAPAAPAAPRASGGVAATVAANGGPASRPPPVPSPATTKARKRKTKKPP

KRPEATPPPDANATVAAGHATLRAHLREIKVENADAQFYVCPPPTGATVVQFEQPRRCPTRPEGQNYTEG

IAVVFKENIAPYKFKATMYYKDVTVSQVWFGHRYSQFMGIFEDRAPVPFEEVIDKINAKGVCRSTAKYVR

NNMETTAFHRDDHETDMELKPAKVATRTSRGWHTTDLKYNPSRVEAFHRYGTTVNCIVEEVDARSVYPYD

EFVLATGDFVYMSPFYGYREGSHTEHTSYAADRFKQVDGFYARDLTTKARATSPTTRNLLTTPKFTVAWD

WVPKRPAVCTMTKWQEVDEMLRAEHGGSFRFSSDAISTTFTTNLTQYSLSRVDLGDCIGRDAREAIDRMF

ARKYNATHIKVGQPQYYLATGGFLIAYQPLLSNTLAELYVREYMREQDRKPRNATPAPLREAPSANASVE

RIKTTSSIEFARLQFTYNHIQRHVNDMLGRIAVAWCELQNHELTLWNEARKLNPNAIASATVGRRVSARM

LGDVMAVSTCVPVAPDNVIVQNSMRVSSRPGTCYSRPLVSFRYEDQGPLIEGQLGENNELRLTRDALEPC

TVGHRRYFIFGGGYVYFEEYAYSHQLSRADVTTVSTFIDLNITMLEDHEFVPLGVYTRHEIKDSGLLDYT

EVQRRNQLHDLRFADIDTVIRADANAAMFAGLCAFFEGMGDLGRAVGKVVMGVVGGVVSAVSGVSSFMSN

PFGALAVGLLVLAGLVAAFFAFRYVLQLQRNPMKALYPLTTKEPKTSDPGGVGGEGEEGAEGGGFDEAKL

AEAREMIRYMALVSAMERTEHKARKKGTSALLSSKVTNMVLRKRNKARYSPLHNEDEAGDEDEL

>HS2GPB_33 MRGGGLICALVVGALVAAVASAAPAAPRASGGVAATVAANGGPASRPPPVPSPATTKARKRKTKKPPKRP

EATPPPDANATVAAGHATLRAHLREIKVENADAQFYVCPPPTGATVVQFEQPRRCPTRPEGQNYTEGIAV

VFKENIAPYKFKATMYYKDVTVSQVWFGHRYSQFMGIFEDRAPVPFEEVIDKINAKGVCRSTAKYVRNNM

ETTAFHRDDHETDMELKPAKVATRTSRGWHTTDLKYNPSRVEAFHRYGTTVNCIVEEVDARSVYPYDEFV

LATGDFVYMSPFYGYREGSHTEHTSYAADRFKQVDGFYARDLTTKARATSPTTRNLLTTPKFTVAWDWVP

KRPAVCTMTKWQEVDEMLRAEYGGSFRFSSDAISTTFTTNLTEYSLSRVDLGDCIGRDAREAIDRMFARK

YNATHIKVGQPQYYLATGGFLIAYQPLLSNTLAELYVREYMREQDRKPRNATPAPLREAPSANASVERIK

TTSSIEFARLQFTYNHIQRHVNDMLGRIAVAWCELQNHELTLWNEARKLNPNAIASATVGRRVSARMLGD

VMAVSTCVPVAPDNVIVQNSMRVSSRPGTCYSRPLVSFRYEDQGPLIEGQLGENNELRLTRDALEPCTVG

HRRYFIFGGGYVYFEEYAYSHQLSRADVTTVSTFIDLNITMLEDHEFVPLEVYTRHEIKDSGLLDYTEVQ

RRNQLHDLRFADIDTVIRADANAAMFAGLCAFFEGMGDLGRAVGKVVMGVVGGVVSAVSGVSSFMSNPFG

ALAVGLLVLAGLVAAFFAFRYVLQLQRNPMKALYPLTTKELKTSDPGGVGGEGEEGAEGGGFDEAKLAEA

REMIRYMALVSAMERTEHKARKKGTSALLSSKVTNMVLRKRNKARYSPLHNEDEAGDEDEL

>HS2GPB_34

MRGGGLICALVVGALVAAVASAAPAAPAAPRASGGVAATVAAKGGPASRPPPVPSPATTKARKRKTKKPP

KRPEATPPPDANATVAAGHATLRAHLREIKVENADAQFYVCPPPTGATVVQFEQPRRCPTRPEGQNYTEG

IAVVFKENIAPYKFKATMYYKDVTVSQVWFGHRYSQFMGIFEDRAPVPFEEVIDKINAKGVCRSTAKYVR

NNMETTAFHRDDHETDMELKPAKVATRTSRGWHTTDLKYNPSRVEAFHRYGTTVNCIVEEVDARSVYPYD

EFVLATGDFVYMSPFYGYREGSHTEHTSYAADRFKQVDGFYARDLTTKARATSPTTRNLLTTPKFTVAWD

WVPKRPAVCTMTKWQEVDEMLRAEYGGSFRFSSDAISTTFTTNLTQYSLSRVDLGDCIGRDAREAIDRMF

ARKYNATHIKVGQPQYYLATGGFLIAYQPLLSNTLAELYVREYMREQDRKPRNATPAPLREAPSANASVE

RIKTTSSIEFARLQFTYNHIQRHVNDMLGRIAVAWCELQNHELTLWNEARKLNPNTIASATVGRRVSARM

LGDVMAVSTCVPVAPDNVIVQNSMRVSSRPGTCYSRPLVSFRYEDQGPLIEGQLGENNELRLTRDALEPC

TVGHRRYFIFGGGYVYFEEYAYSHQLSRADVTTVSTFIDLNITMLEDHEFVPLGVYTRHEIKDSGLLDYT

EVQRRNQLHDLRFADIDTVIRADANAAMFAGLCAFFEGMGDLGRAVGKVVMGVVGGVVSAVSGVSSFMSN

PFGALAVGLLVLAGLVAAFFAFRYVLQLQRNPMKALYPLTTKEPKTSDPGGVGGEGEEGAEGGGFDEAKL

AEAREMIRYMALVSAMERTEHKARKKGTSALLSSKVTNMVLRKRNKARYSPLHNEDEAGDEDEL

>HS2GPB_35

MRGGGLICALVVGALVAAVASAAPAAPRASGGVAATVAANGGPASRPPPVPSPATTRARKRKTKKPPKRP

EATPPPDANATVAAGHATLRAHLREIKVENADAQFYVCPPPTGATVVQFEQPRRCPTRPEGQNYTEGIAV

VFKENIAPYKFKATMYYKDVTVSQVWFGHRYSQFMGIFEDRAPVPFEEVIDKINTKGVCRSTAKYVRNNM

ETTAFHRDDHETDMELKPAKVATRTSRGWHTTDLKYNPSRVEAFHRYGTTVNCIVEEVDARSVYPYDEFV

LATGDFVYMSPFYGYREGSHTEHTSYAADRFKQVDGFYARDLTTKARATSPTTRNLLTTPKFTVAWDWVP

KRPAVCTMTKWQEVDEMLRAEYGGSFRFSSDAISTTFTTNLTEYSLSRVDLGDCIGRDAREAIDRMFARK

YNATHIKVGQPQYYLATGGFLIAYQPLLSNTLAELYVREYMREQDRKPRNATPAPLREAPSANASVERIK

TTSSIEFARLQFTYNHIQRHVNDMLGRIAVAWCELQNHELTLWNEARKLNPNAIASATVGRRVSARMLGD

VMAVSTCVPVAPDNVIVQNSMRVSSRPGTCYSRPLVSFRYEDQGPLIEGQLGENNELRLTRDALEPCTVG

HRRYFIFGGGYVYFEEYAYSHQLSRADVTTVSTFIDLNITMLEDHEFVPLEVYTRHEIKDSGLLDYTEVQ

RRNQLHDLRFADIDTVIRADANAAMFAGLCAFFEGMGDLGRAVGKVVMGVVGGVVSAVSGVSSFMSNPFG

ALAVGLLVLAGLVAAFFAFRYVLQLQRNPMKALYPLTTKELKTSDPGGVGGEGEEGAEGGGFDEAKLAEA

RQMIRYMALVSAMERTEHKARKKGTSALLSSKVTNMVLRKRNKARYSPLHNEDEAGDEDEL

>HS2GPB_36

MRGGGLICALVVGALVAAVASAAPAAPRASGGVAATVAANGGPASRPPPVPSPATTKARKRKTKKPPERP

EATPPPDANATVAAGHATLRAHLREIKVENADAQFYVCPPPTGATVVQFEQPRRCPTRPEGQNYTEGIAV

VFKENIAPYKFKATMYYKDVTVSQVWFGHRYSQFMGIFEDRAPVPFEEVIDKINAKGVCRSTAKYVRNNM

ETTAFHRDDHETDMELKPAKVATRTSRGWHTTDLKYNPSRVEAFHRYGTTVNCIVEEVDARSVYPYDEFV

LATGDFVYMSPFYGYREGSHTEHTSYAADRFKQVDGFYARDLTTKARATSPTTRNLLTTPKFTVAWDWVP

KRPAVCTMTKWQEVDEMLRAEYGGSFRFSSDAISTTFTTNLTEYSLSRVDLGDCIGRDAREAIDRMFARK

YNATHIKVGQPQYYLATGGFLIAYQPLLSNTLAELYVREYMREQDRKPRNATPAPLREAPSANASVERIK

TTSSIEFARLQFTYNHIQRHVNDMLGRIAVAWCELQNHELTLWNEARKLNPNAIASATVGRRVSARMLGD

VMAVSTCVPVAPDNVIVQNSMRVSSRPGTCYSRPLVSFRYEDQGPLIEGQLGENNELRLTRDALEPCTVG

HRRYFIFGGGYVYFEEYAYSHQLSRADVTTVSTFIDLNITMLEDHEFVPLEVYTRHEIKDSGLLDYTEVQ

RRNQLHDLRFADIDTVIRADANAAMFAGLCAFFEGMGDLGRAVGKVVMGVVGGVVSAVSGVSSFMSNPFG

ALAVGLLVLAGLVAAFFAFRYVLQLQRNPMKALYPLTTKELKTSDPGGVGGEGEEGAEGGGFDEAKLAEA

REMIRYMALVSAMERTEHKARKKGTSALLSSKVTNMVLRKRNKARYSPLHNEDEAGDEDEL

>HS2GPB_37

MRGGGLICALVVGALVAAVASAAPAAPRASGGVAATVAANGGPASRPPPVPSPATTKARKRKTKKPPKRP

EATPPPDANATVAAGHATLRAHLREIKVENADAQFYVCPPPTGATVVQFEQPRRCPTRPEGQNYTEGIAV

VFKENIAPYKFKATMYYKDVTVSQVWFGHRYSQFMGIFEDRAPVPFEEVIDKINAKGVCRSTAKYVRNNM

ETTAFHRDDHETDMELKPAKVATRTSRGWHTTDLKYNPSRVEAFHRYGTTVNCIVEEVDARSVYPYDEFV

LATGDFVYMSPFYGYREGSHTEHTSYAADRFKQVDGFYARDLTTKARATSPTTRNLLTTPKFTVAWDWVP

KRPAVCTMTKWQEVDEMLRAEYGGSFRFSSDAISTTFTTNLTEYSLSRVDLGDCIGRDAREAIDRMFARK

YNATHIKVGQPQYYLATGGFLIAYQPLLSNTLAELYVREYMREQDRKPRNATPAPLREAPSANASVERIK

TTSSIEFARLQFTYNHIQRHVNDMLGRIAVAWCELQNHELTLWNEARKLNPNAIASATVGRRVSARMLGD

VMAVSTCVPVAPDNVIVQNSMRVSSRPGTCYSRPLVSFRYEDQGPLIEGQLGENNELRLTRDALEPCTVG

HRRYFIFGGGYVYFEEYAYSHQLSRADVTTVSTFIDLNITMLEDHEFVPLEVYTRHEIKDSGLLDYTEVQ

RRNQLHDLRFADIDTVIRADANAAMFAGLCAFFEGMGDLGRAVGKVVMGVVGGVVSAVSGVSSFMSNPFG

ALAVGLLVLAGLVAAFFAFRYVLQLQRNPMKALYPLTTKELKTSDPGGVGGEGEEGAEGGGFDEAKLAEA

RQMIRYMALVSAMERTEHKARKKGTSALLSSKVTNMVLRKRNKARYSPLHNEDEAGDEDEL

>HS2GPB_38 MRGGGLICALVVGALVAAVASAAPAAPRASGGVAATVAANGGPASRPPPVPSPATTKARKRKTKRPPKRP

EATPPPDANATVAAGHATLRAHLREIKVENADAQFYVCPPPTGATVVQFEQPRRCPTRPEGQNYTEGIAV

VFKENIAPYKFKATMYYKDVTVSQVWFGHRYSQFMGIFEDRAPVPFEEVIDKINAKGVCRSTAKYVRNNM

ETTAFHRDDHETDMELKPAKVATRTSRGWHTTDLKYNPSRVEAFHRYGTTVNCIVEEVDARSVYPYDEFV

LATGDFVYMSPFYGYREGSHTEHTSYAADRFKQVDGFYARDLTTKARATSPTTRNLLTTPKFTVAWDWVP

KRPAVCTMTKWQEVDEMLRAEYGGSFRFSSDAISTTFTTNLTEYSLSRVDLGDCIGRDAREAIDRMFARK

YNATHIKVGQPQYYLATGGFLIAYQPLLSNTLAELYVREYMREQDRKPRNATPAPLREAPSANASVERIK

TTSSIEFARLQFTYNHIQRHVNDMLGRIAVAWCELQNHELTLWNEARKLNPNAIASATVGRRVSARMLGD

VMAVSTCVPVAPDNVIVQNSMRVSSRPGTCYSRPLVSFRYEDQGPLIEGQLGENNELRLTRDALEPCTVG

HRRYFIFGGGYVYFEEYAYSHQLSRADVTTVSTFIDLNITMLEDHEFVPLEVYTRHEIKDSGLLDYTEVQ

RRNQLHDLRFADIDTVIRADANAAMFAGLCAFFEGMGDLGRAVGKVVMGVVGGVVSAVSGVSSFMSNPFG

ALAVGLLVLAGLVAAFFAFRYVLQLQRNPMKALYPLTTKELKTSDPGGVGGEGEEGAEGGGFDEAKLAEA

REMIRYMALVSAMERTEHKARKKGTSALLSSKVTNMVLRKRNKARYSPLHNEDEAGDEDEL

>HS2GPB_39 MRGGGLICALVVGALVAAVASAAPAAPRASGGVAATVAANGGPASQPPPVPSPATTKARKRKTKKPPKRP

EATPPPDANATVAAGHATLRAHLREIKVENADAQFYVCPPPTGATVVQFEQPRRCPTRPEGQNYTEGIAV

VFKENIAPYKFKATMYYKDVTVSQVWFGHRYSQFMGIFEDRAPVPFEEVIDKINAKGVCRSTAKYVRNNM

ETTAFHRDDHETDMELKPAKVATRTSRGWHTTDLKYNPSRVEAFHRYGTTVNCIVEEVDARSVYPYDEFV

LATGDFVYMSPFYGYREGSHTEHTSYAADRFKQVDGFYARDLTTKARATSPTTRNLLTTPKFTVAWDWVP

KRPAVCTMTKWQEVDEMLRAEYGGSFRFSSDAISTTFTTNLTEYSLSRVDLGDCIGRDAREAIDRMFARK

YNATHIKVGQPQYYLATGGFLIAYQPLLSNTLAELYVREYMREQDRKPRNATPAPLREAPSANASVERIK

TTSSIEFARLQFTYNHIQRHVNDMLGRIAVAWCELQNHELTLWNEARKLNPNAIASATVGRRVSARMLGD

VMAVSTCVPVAPDNVIVQNSMRVSSRPGTCYSRPLVSFRYEDQGPLIEGQLGENNELRLTRDALEPCTVG

HRRYFIFGGGYVYFEEYAYSHQLSRADVTTVSTFIDLNITMLEDHEFVPLEVYTRHEIKDSGLLDYTEVQ

RRNQLHDLRFADIDTVIRADANAAMFAGLCAFFEGMGDLGRAVGKVVMGVVGGVVSAVSGVSSFMSNPFG

ALAVGLLVLAGLVAAFFAFRYVLQLQRNPMKALYPLTTKELKTSDPGGVGGEGEEGAEGGGFDEAKLAEA

REMIRYMALVSAMERTEHKARKKGTSALLSSKVTNMVLRKRNKARYSPLHNEDEAGDEDEL
