## Supplementary file 2 for "Contriving a chimeric polyvalent vaccine to prevent infections caused by Herpes Simplex Virus (Type-1 and Type-2): an exploratory immunoinformatic approach"

**Glycoprotein C**

**HSV1=20 AND HSV2=7**

>HSV1GPC_1 MAPGRVGLAVVLWSLLWLGAGVSGGSETASTGPTITAGAVTNASEAPTSGSPGSAASPEVTPTSTPNPNN

VTQNKTTPTEPASPPTTPKPTSTPKSPPTSTPDPKPKNNTTPAKSGRPTKPPGPVWCDRRDPLARYGSRV

QIRCRFRNSTRMEFRLQIWRYSMGPSPPIAPAPDLEEVLTNITAPPGGLLVYDSAPNLTDPHVLWAEGAG

PGADPPLYSVTGPLPTQRLIIGEVTPATQGMYYLAWGRMDSPHEYGTWVRVRMFRPPSLTLQPHAVMEGQ

PFKATCTAAAYYPRNPVEFVWFEDDHQVFNPGQIDTQTHEHPDGFTTVSTVTSEAVGGQVPPRTFTCQMT

WHRDSVTFSRRNATGLALVLPRPTITMEFGVRIVVCTAGCVPEGVTFAWFLGDDPSPAAKSAVTAQESCD

HPGLATVRSTLPISYDYSEYICRLTGYPAGIPVLEHHGSHQPPPRDPTERQVIEAIEWVGIGIGVLAAGV

LVVTAIVYVVRTSQSRQRHRR

>HSV1GPC_2

MAPGRVGLAVVLWSLLWLGAGVSGGSETASTGPTITAGAVTNASEAPTSGSPGSAASPEVTPTSTPNPNN

VTQNKTTPTEPASPPTTPKPTSTPKSPPTSTPDPKPKNNTTPAKSGRPTKPPGPVWCDRRDPLARYGSRV

QIRCRFRNSTRMEFRLQIWRYSMGPSPPIAPAPDLEEVLTNITAPPGGLLVYDSAPNLTDPHVLWAEGAG

PGADPPLYSVTGPLPTQRLIIGEVTPATQGMYYLAWGRMDSPHEYGTWVRVRMFRPPSLTLQPHAVMEGQ

PFKATCTAAAYYPRNPVEFVWFEDDHQVFNPGQIDTQTHEHPDGFTTVSTVTSEAVGGQVPPRTFTCQMT

WHRDSVTFSRRNATGLALVLPRPTITMEFGVRHVVCTAGCVPEGVTFAWFLGDDPSPAAKSAVTAQESCD

HPGLATVRSTLPISYDYSEYICRLTGYPAGIPVLEHHGSHQPPPRDPTERQVIEAIEWVGIGIGVLAAGV

LVVTAIVYVVRTSQSRQRHRR

>HSV1GPC_3

MAPGRVGLAVVLWSLLWLGAGVSGGSETASTGPTITAGAVTNASEAPTSGSPGSAANPEVTPTSTPNPNN

VTQNKTTPTEPASPPTTPKPTSTPKSPPTSTPDPKPKNNTTPAKSGRPTKPPGPVWCDRRDPLARYGSRV

QIRCRFRNSTRMEFRLQIWRYSMGPSPPIAPAPDLEEVLTNITAPPGGLLVYDSAPNLTDPHVLWAEGAG

PGADPPLYSVTGPLPTQRLIIGEVTPATQGMYYLAWGRMDSPHEYGTWVRVRMFRPPSLTLQPHAVMEGQ

PFKATCTAAAYYPRNPVEFVWFEDDHQVFNPGQIDTQTHEHPDGFTTVSTVTSEAVGGQVPPRTFTCQMT

WHRDSVTFSRRNATGLALVLPRPTITMEFGVRHVVCTAGCVPEGVTFAWFLGDDPSPAAKSAVTAQESCD

HPGLATVRSTLPISYDYSEYICRLTGYPAGIPVLEHHGSHQPPPRDPTERQVIEAIEWVGIGIGVLAAGV

LVVTAIVYVVRTSQSRQRHRR

>HSV1GPC_4

MAPGRVGLAVVLWSLLWLGAGVSGGSETASTGPTITAGAVTNASEAPTSGSPGSAASPEVTPTSTPNPNN

VTQNKTTPTEPASPPTTPKPTSTPKSPPTSTPDPKPKNNTTPAKSGRPTKPPGPVWCDRRDPLARYGSRV

QIRCRFRNSTRMEFRLQIWRYSMGPSPPIAPAPDLEEVLTNITAPPGGLLVYDSAPNLTDPHVLWAEGAG

PGADPPLYSVTGPLPTQRLIIGEVTPATQGMYYLAWGRMDSPHEYGTWVRVRMFRPPSLTLQPHAVMEGQ

PFKATCTADAYYPRNPVEFVWFEDDHQVFNPGQIDTQTHEHPDGFTTVSTVTSEAVGGQVPPRTFTCQMT

WHRDSVTFSRRNATGLALVLPRPTITMEFGVRHVVCTAGCVPEGVTFAWFLGDDPSPAAKSAVTAQESCD

HPGLATVRSTLPISYDYSEYICRLTGYPAGIPVLEHHGSHQPPPRDPTERQVIEAIEWVGIGIGVLAAGV

LVVTAIVYVVRTSQSRQRHRR

>HSV1GPC_5

MAPGRVGLAVVLWSLLWLGAGVSGGSETASTGPTITAGAVTNASEAPTSGSPGSAASPEVTPTSTPNPNN

VTQNKTTPTEPASPPTTPKPTSTPKSPPTSTPDPKPKNNTTPAKSGRPTKPPGPVWCDRRDPLARYGSRV

QIRCRFRNSTRMEFRLQIWRYSMGPSPPIAPAPDLEEVLTNITAPPGGLLVYDSAPNLTDPHVLWAEGAG

PGADPPLYSVTGPLPTQRLIIGEVTPATQGMYYLAWGRMDSPHEYGTWVRVRMFRPPSLTLQPHAVMEGQ

PFKATCTAAAYYPRNPVEFVWFEDDHQVFNPGQIDTQTHEHPDGFTTVSTVTSEAVGGQVPPRTFTCQMT

WHRDSVTFSRRNATGLALVLPRPTITMEFGVRHVVCTAGCVPEGVTFAWFLGDDPSPAAKSAVTAQESCD

RPGLATVRSTLPISYDYSEYICRLTGYPAGIPVLEHHGSHQPPPRDPTERQVIEAIEWVGIGIGVLAAGV

LVVTAIVYVVRTSQSRQRHRR

>HSV1GPC_6 MAPGRVGLAVVLWSLLWLGAGVAGGSETASTGPTITAGAVTNASEAPTSGSPGSAASPEVTPTSTPNPNN

VTQNKTTPTEPASPPTTPKPTSTPKSPPTSTPDPKPKNNTTPAKSGRPTKPPGPVWCDRRDPLARYGSRV

QIRCRFRNSTRMEFRLQIWRYSMGPSPPIAPAPDLEEVLTNITAPPGGLLVYDSAPNLTDPHVLWAEGAG

PGADPPLYSVTGPLPTQRLIIGEVTPATQGMYYLAWGRMDSPHEYGTWVRVRMFRPPSLTLQPHAVMEGQ

PFKATCTAAAYYPRNPVEFVWFEDDRQVFNPGQIDTQTHEHPDGFTTVSTVTSEAVGGQVPPRTFTCQMT

WHRDSVTFSRRNATGLALVLPRPTITMEFGVRHVVCTAGCVPEGVTFAWFLGDDPSPAAKSAVTAQESCD

HPGLATVRSTLPISYDYSEYICRLTGYPAGIPVLEHHGSHQPPPRDPTERQVIEAIEWVGIGIGVLAAGV

LVVTAIVYVVRTSQSRQRHRR

>HSV1GPC_7

MAPGRVGLAVVLWSLLWLGAGVSGGSETASTGPTITAGAVTNASEAPTSGSPGSAASPEVTPTSTPNPNN

VTQNKTTPTEPASPPTTPKPTSTPKSPPTSTPDPKPKNNTTPAKSGRPTKPPGPVWCDRRDPLARYGSRG

QIRCRFRNSTRMEFRLQIWRYSMGPSPPIAPAPDLEEVLTNITAPPGGLLVYDSAPNLTDPHVLWAEGAG

PGADPPLYSVTGPLPTQRLIIGEVTPATQGMYYLAWGRMDSPHEYGTWVRVRMFRPPSLTLQPHAVMEGQ

PFKATCTAAAYYPRNPVEFVWFEDDHQVFNPGQIDTQTHEHPDGFTTVSTVTSEAVGGQVPPRTFTCQMT

WHRDSVTFSRRNATGLALVLPRPTITMEFGVRHVVCTAGCVPEGVTFAWFLGDDPSPAAKSAVTAQESCD

RPGLATVRSTLPISYDYSEYICRLTGYPAGIPVLEHHGSHQPPPRDPTERQVIEAIEWVGIGIGVLAAGV

LVVTAIVYVVRTSQSRQRHRR

>HSV1GPC_8

MAPGRVGLAVVLWSLLWLGAGVAGGSETASTGPTITAGAVTNASEAPTSGSPGSAASPEVTPTSTPTPNN

VTQNKTTPTEPASPPTTPKPTSTPKSPPTSTPDPKPKNNTTPAKSGRPTKPPGPVWCDRRDPLARYGSRV

QIRCRFRNSTRMEFRLQIWRYSMGPSPPIAPAPDLEEVLTNITAPPGGLLVYDSAPNLTDPHVLWAEGAG

PGADPPLYSVTGPLPTQRLIIGEVTPATQGMYYLAWGRMDSPHEYGTWVRVRMFRPPSLTLQPHAVMEGQ

PFKATCTAAAYYPRNPVEFVWFEDDRQVFNPGQIDTQTHEHPDGFTTVSTVTSEAVGGQVPPRTFTCQMT

WHRDSVTFSRRNATGLALVLPRPTITMEFGVRHVVCTAGCVPEGVTFAWFLGDDPSPAAKSAVTAQESCD

HPGLATVRSTLPISYDYSEYICRLTGYPAGIPVLEHHGSHQPPPRDPTERQVIEAIEWVGIGIGVLAAGV

LVVTAIVYVVRTSQSRQRHRR

>HSV1GPC_9

MAPGRVGLAVVLWSLLWLGAGVSGDSETASTGPTITAGAVTNASEAPTSGSPGSAASPEVTPTSTPNPNN

VTQNKTIPTEPASPPTTPKPTSTPKSPPTSTPDPKPKNNTTPAKSGRPTKPPGPVWCDRRDPLARYGSRV

QIRCRFRNSTRMEFRLQIWRYSMGPSPPIAPAPDLEEVLTNITAPPGGLLVYDSAPNLTDPHVLWAEGAG

PGADPPLYSVTGPLPTQRLIIGEVTPATQGMYYLAWGRMDSPHEYGTWVRVRMFRPPSLTLQPHAVMEGQ

PFKATCTAAAYYPRNPVEFVWFEDDHQVFNPGQIDTQTHEHPDGFTTVSTVTSEAVGGQVPPRTFTCQIT

WHRDSVTFSRRNATGLALVLPRPTITMEFGVRHVVCTAGCVPEGVTFAWFLGDDPSPAAKSAVTAQESCD

HPGLATVRSTLPISYDYSEYICRLTGYPAGIPVLEHHGSHQPPPRDPTERQVIEAIEWVGIGIGVLAAGV

LVVTAIVYVVRTSQSRQRHRR

>HSV1GPC_10

MAPGRVGLAVVLWSLLWLGAGVAGGSETASTGPTITAGAVTNASEAPTSGSPGSAASPEVTPTSTPNPNN

VTQNKTTPTEPASPPTTPKPTSTPKSPPTSTPDPKPKNNTTPAKSGRPTKPPGPVWCDRRDPLARYGSRV

QIRCRFRNSTRMEFRLQIWRYSMGPSPPIAPAPDLEEVLTNITAPPGGLLVYDSAPNLTDPHVLWAEGAG

PGADPPLYSVTGPLPTQRLIIGEVTPATQGMYYLAWGRMDSPHEYGTWVRVRMFRPPSLTLQPHAVMEGQ

PFKATCTAAAYYPRNPVEFVWFEDDRQVFNPGQIDTQTHEHPDGFTTVSTVTSEAVGGQVPPRTFTCQMT

WHRDSVMFSRRNATGLALVLPRPTITMEFGVRHVVCTAGCVPEGVTFAWFLGDDPSPAAKSAVTAQESCD

HPGLATVRSTLPISYDYSEYICRLTGYPAGIPVLEHHGSHQPPPRDPTERQVIEAIEWVGIGIGVLAAGV

LVVTAIVYVVRTSQSRQRHRR

>HSV1GPC_11 MAPGRVGLAVVLWSLLWLGAGVSGGSETASTGPTITAGAVTNASEAPTSGSPGSAASPEVTPTSTPNPNN

VTQNKTTPTEPASPPTTPKPTSTPKSPPTSTPDPKPKNNTTPAKSGRPTKPPGPVWCDRRDPLARYGSRV

QIRCRFRNSTRMEFRLQIWRYSMGPSPPIAPAPDLEEVLTNITAPPGGLLVYDSAPNLTDPHVLWAEGAG

PGADPPLYSVTGPLPTQRLIIGEVTPATQGMYYLAWGRMDSPHEYGTWVRVRMFRPPSLTLQPHAVMEGQ

PFKATCTAAAYYPRNPVEFVWFEDDHQVFNPGQIDTQTHEHPDGFTTVSTVTSEAVGGQVPPRTFTCQMT

WHRDSVTFSRRNATGLALVLPRPTITMEFGVRHVVCTAGCVPEGVTFAWFLGDDPSPAAKSAVTAQESCD

RPGLATVRSTLPISYDYSEYICRLTGYPAGIPVLEHHGSHQPPPRDPTERQVIEAIECVGIGIGVLAAGV

LVVTAIVYVVRTSQSRQRHRR

>HSV1GPC_12

MAPGRVGLAVVLWSLVWLGAGVSGGSETASTGPTITAGAVMNASEAPTSGSPGSAASPEVTPTSTPNPNN

VTQNQTTPTEPASPPTTPKPTSTPKSPPTSTPDPKPKNNTTPAKSGRPTKPPGPVWCDRRDPLARYGSRV

QIRCRFRNSTRMEFRLQIWRYSMGPSPPIAPAPDLEEVLTNITAPPGGLLVYDSAPNLTDPHVLWAEGAG

PGADPPLYSVTGPLPTQRLIIGEVTPATQGMYYLAWGRMDSPHEYGTWVRVRMFRPPSLTLQPHAVMEGQ

PFKATCTAAAYYPRNPVEFVWFEDDHQVFNPVQIDTQTHEHPDGFTTVSTVTSEAVGGQVPPRTFTCQMT

WHRDSVTFSRRNATGLALVLPRPTITMEFGVRHVVCTAGCVPEGVTFAWFLGDDPSPAAKSAVTAQESCD

HPGLATVRSTLPISYDYSEYICRLTGYPAGIPVLEHHGSHQPPPRDPTERQVIEAIEWVGIGIGVLAAGV

LVVTAIVYVVRTSQSRQRHRR

>HSV1GPC_13

MAPGRVGLAVVLWSLLWLGAGVSGGSETASTGPTITAGAVTNASEAPTSGSPGSAASPEVTPTSTPNPNN

VTQNKTIPTEPASPPTTPKPTSTPKSPPTSTPDPKPKNNTTPAKSGRPTKPPGPVWCDRRDPLARYGSRV

QIRCRFRNSTRMEFRLQIWRYSMGPSPPIAPAPDLEEVLTNITAPPGGLLVYDSAPNLTDPHVLWAEGAG

PGADPPLYSVTGPLPTQRLIIGEVTPATQGMYYLAWGRMDSPHEYGTWVRVRMFRPPSLTLQPHAVMEGQ

PFKATCTADAYYPRNPVEFVWFEDDHQVFNPGQIDTQTHEHPDGFTTVSTVTSEAVGGQVPPRTFTCQMT

WHRDSVTFSRRNATGLALVLPRPTITMEFGVRHVVCTAGCVPEGVTFAWFLGDDPSPAAKSAVTAQESCD

HPGLATVRSTLPISYDYSEYICRLTGYPAGIPVLEHHGSHQPPPRDPTERKVIEAIEWVGIGIGVLAAGV

LVVTAIVYVVRTSQSRQRHRR

>HSV1GPC_14 MAPGRVGLAVVLWSLLWLGAGVAGGSETASTGPTITAGAVTNASEAPTSGSPGSAASPEVTPTSTPNPNN

VTQNKTTPTEPASPPTTPKPTSTPKSPPTSTPDPKPKNNTTPAKSGRPTKPPGPVWCDRRDPLARYGSRV

QIRCRFRNSTRMEFRLQIWRYSMGPSPPIAPAPDLEEVLTNITAPPGGLLVYDSAPNLTDPHVLWAEGAG

PGADPPLYSVTGPLPTQRLIIGEVTPATQGMYYLAWGRMDSPHEYGTWVRVRMFRPPSLTLQPHAVMEGQ

PFKATCTAAAYYPRNPVEFVWFEDDRQVFNXGQIDTQTHEHPDGFTTVSTVTSEAVGGQVPPRTFTCQMT

WHRDSVTFSRRNATGLALVLPRPTITMEFGVRHVVCTAGCVPEGVTFAWFLGDDPSPAAKSAVTAQESCD

HPGLATVRSTLPISYDYSEYICRLTGYPAGIPVLEHHGSHQPPPRDPTERQVIEAIEWVGIGIGVLAAGV

LVVTAIVYVVRTSQSRQRHRR

>HSV1GPC_15 MAPGRVGLAVVLWSLLWLGAGVAGGSETASTGPTITAGAVTNASEAPTSGSPGSAANPEVTPTSTPNPNN

VTQNKTTPTEPASPPTTPKPTSTPKSPPTSTPDPKPKNNTTPAKSGRPTKPPGPVWCDRRDPLARYGSRV

QIRCRFRNSTRMEFRLQIWRYSMGPSPPIAPAPDLEEVLTNITAPPGGLLVYDSAPNLTDPHVLWAEGAG

PGADPPLYSVTGPLPTQRLIIGEVTPATQGMYYLAWGRIDSPHEYGTWVRVRMFRPPSLTLQPHAVMEGQ

PFKATCTAAAYYPRNPVEFVWFEDDRQVFNPGQIDTQTHEHPDGFTTVSTVTSEAVGGQVPPRTFTCQMT

WHRDSVTFSRRNATGLALVLPRPTITMEFGVRHVVCTAGCVPEGVTFAWFLGDDPSPAAKSAVTAQESCD

HPGLATVRSTLPISYDYSEYICRLTGYPAGIPVLEHHGSHQPPPRDPTERQVIEAIEWVGIGIGVLAAGV

LVVTAIVYVVRTSQSRQRHRR

>HSV1GPC_16

MAPGRVGLAVVLWSLLWLGAGVAGGSETASTGPTITAGAVTNASEAPTSGSPGSAASPEVTPTSTPNPNN

VTQNKTTPTEPASPPTTPKPTSTPKSPPTSTPDPKPKNNTTPAKSGRPTKPPGPVWCDRRDPLARYGSRV

QIRCRFRNSTRMEFRLQIWRYSMGPSPPIAPAPDLEEVLTNITAPPGGLLVYDSAPNLTDPHVLWAEGAG

PGADPPLYSVTGPLPTQRLIIGEVTPATQGMYYLAWGRMDSPHEYGTWVRVRMFRPPSLTLQPHAVMEGQ

PFKATCTAAAYYPRNPVEFDWFEDDRQVFNPGQIDTQTHEHPDGFTTVSTVTSEAVGGQVPPRTFTCQMT

WHRDSVTFSRRNATGLALVLPRPTITMEFGVRHVVCTAGCVPEGVTFAWFLGDDPSPAAKSAVTAQESCD

HPGLATVRSTLPISYDYSEYICRLTGYPAGIPVLEHHGSHQPPPRDPTERQVIEAIEWVGIGIGVLAAGV

LVVTAIVYVVRTSQSRQRHRR

>HSV1GPC_17 MAPGRVGLAVVLWSLLWLGAGVSGGSETASTGPTITAGAVTNASEAPTSGSPGSAASPEVTPTSTPNPNN

VTQNKTTPTEPASPPTTPKPTSTPKSPPTSTPDPKPKNNTTPAKSGRPTKPPGPVWCDRRDPLARYGSRV

QIRCRFRNSTRMEFRLQIWRYSMGPSPPIAPAPDLEEVLTNIPAPPGGLLVYDSAPNLTDPHVLWAEGAG

PGADPPLYSVTGPLPTQRLIIGEVTPATQGMYYLAWGRMDSPHEYGTWVRVRMFRPPSLTLQPHAVMEGQ

PFKATCTAAAYYPRNPVEFVWFEDDHQVFNPGQIDTQTHEHPDGFTTVSTVTSEAVGGQVPPRTFTCQMT

WHRDSVTFSRRNATGLALVLPRPTITMEFGVRHVVCTAGCVPEGVTFAWFLGDDPSPAAKSAVTAQESCD

RPGLATVRSTLPISYDYSEYICRLTGYPAGIPVLEHHGSHQPPPRDPTERQVIEAIEWVGIGIGVLAAGV

LVATAIVYVVRTSQSRQRHRR

>HSV1GPC_18 MATGRVGLAVVLWSLLWLGAGVSGGSETASTGPTITAGAVTNASEAPTSGSPGSAASPEVTPTSTPNPNN

VTQNKTTPTEPASPPTTPKPTSTPKSPPTSTPDPKPKNNTTPAKSGRPTKPPGPVWCDRRDPLARYGSRV

QIRCRFRNSTRMEFRLQIWRYSTGPSPPIAPAPDLEEVLTNITAPPGGLLVYDSAPNLTDPHVLWAEGAG

PGADPPLYSVTGPLPTQRLIIGEVTPATQGMYYLAWGRMDSPHEYGTWVRVRMFRPPSLTLQPHAVMEGQ

PFKATCTAAAYYPRNPVEFVWFEDDHQVFNPGQIDTQTHEHPDGFTTVSTVTSEAVGGQVPPRTFTCQMT

WHRDSVTFSRRNATGLALVLPRPTITMEFGVRHVVCTAGCVPEGVTFAWFLGDDPSPAAKSAVTAQESCD

RPGLATVRSTLPISYDYSEYICRLTGYPAGIPVLEHHGSHQPPPRDPTERQVIEAIEWVGIGIGVLAAGV

LVVTAIVYVVRTSQSRQRHRR

>HSV1GPC_19

MAPGRVGLAVVLWSLLWLGAGVSGGSETASTGPTITAGAVTNASEAPTSGSPGSAASPEVTPTSTPNPNN

VTQNKTTPTEPASPPTTPKPTSTPKSPPTSTPDPKPKNNTTPAKSGRPTNPPGPVWCDRRDPLARYGSRV

QIRCRFRNSTRMEFRLQIWRYSMGPSPPIAPAPDLEEVLTNITAPPGGLLVYDSAPNLTDPHVLWAEGAG

PGADPPLYSVTGPLPTQRLIIGEVTPATQGMYYLAWGRMDSPHEYGTWVRVRMFRPPSLTLQPHAVMEGQ

PFKATCTATAYYPRNPVEFVWFEDDHQVFNPGQIDTQTHEHPDGFTTVSTVTSEAVGGQVPPRTFTCQMT

WHRDSVTFSRRNATGLALVLPRPTITMEFGVRHVVCTAGCVPEGVTFAWFLGDDPSPAAKSAVTAQESCD

RPGLATVRSTLPISYDYSEYICRLTGYPAGIPVLEHHGSHQPPPRDPTERQVIEAIEWVGIGIGVLAAGV

LVVTAIVYVVRTSQSRQRHRR

>HSV1GPC_20

MAPGRVGLAVVLWSLLWLGAGVAGGSETASTGPTITAGAVTNASEAPTSGSPGSAASPEVTPTSTPNPNN

VTQNKTTPTEPASPPTTPKPTSTPKSPPTSTPDPKPKNNTTPAKSGRPTKPPGPVWCDRRDPLARYGSRV

QIRCRFRNSTRMEFRLQIWRYSMGPSPPIAPAPDLEEVLTNITAPPGGLLVYDSAPNLTDPHVLWAEGAG

PGADPPLYSVTGPLPTQRLIIGEVTPATQGMYYLAWGRMDNPHEYGTWVRVRMFRPPSLTLQPHAVMEGQ

PFKATCTAAAYYPRNPVEFVWFEDDRQVFNPGQIDTQTHEHPDGFTTVSTVTSEAVGGQVPPRTFTCQMT

WHRDSVMFSRRNATGLALVLPRPTITMEFGVRHVVCTAGCVPEGVTFAWFLGDDPSPAAKSAVTAQESCD

HPGLATVRSTLPISYDYSEYICRLTGYPAGIPVLEHHGSHQPPPRDPTERQVIEAIEWVGIGIGVLAAGV

LVVTAIVYVVRTSQSRQRHRR

>HSV2GPC_21 gi|9629314|ref|NP_044514.1| envelope glycoprotein C [Human herpesvirus 2]

MALGRVGLAVGLWGLLWVGVVVVLANASPGRTITVGPRGNASNAAPSASPRNASAPRTTPTPPQPRKATK

SKASTAKPAPPPKTGPPKTSSEPVRCNRHDPLARYGSRVQIRCRFPNSTRTEFRLQIWRYATATDAEIGT

APSLEEVMVNVSAPPGGQLVYDSAPNRTDPHVIWAEGAGPGASPRLYSVVGPLGRQRLIIEELTLETQGM

YYWVWGRTDRPSAYGTWVRVRVFRPPSLTIHPHAVLEGQPFKATCTAATYYPGNRAEFVWFEDGRRVFDP

AQIHTQTQENPDGFSTVSTVTSAAVGGQGPPRTFTCQLTWHRDSVSFSRRNASGTASVLPRPTITMEFTG

DHAVCTAGCVPEGVTFAWFLGDDSSPAEKVAVASQTSCGRPGTATIRSTLPVSYEQTEYICRLAGYPDGI

PVLEHHGSHQPPPRDPTERQVIRAVEGAGIGVAVLVAVVLAGTAVVYLTHASSVRYRRLR

>HSV2GPC_22 MALGRVGLAVGLWGLLWVGVVVVLANASPGRTITVGPRGNASNAAPSASPRNASAPRTTPTPPQPRKATK

SKASTAKPAPPPKTGPPKTSSEPVRCNRHDPLARYGSRVQIRCRFPNSTRTEFRLQIWRYATATDAEIGT

APSLEEVMVNVSAPPGGQLVYDSAPNRTDPHVIWAEGAGPGASPRLYSVVGPLGRQRLIIEELTLETQGM

YYWVWGRTDRPSAYGTWVRVRVFRPPSLTIHPHAVLEGQPFKATCTAATYYPGNRAEFVWFEDGRRVFDP

AQIHTQTQENPDGFSTVSTVTSAAVGGQGPPRTFTCQLTWHRDSVSFSRRNASGTASVLPRPTITMEFTG

DHAVCTAGCVPEGVTFAWFLGDDSSPAEKVAVASQTSCGRPGTATIRSTLPVSYEQTEYICRLAGYPDGI

PVLEHHGSHQPPPRDPTKRQVIRAVEGAGIGVAVLVAVVLAGTAVVYLTHASSVRYRRLR

>HSV2GPC_23

MALGRVGLAVGLWGLLWVGVVVVLANASPGRTITVGPRGNASNAAPSASPRNASAPRTTPTPPQPRKATK

SKASTAKPAPPPKTGPPKTSSEPVRCNRHDPLARYGSRVQIRCRFPNSTRTEFRLQIWRYATATDAEIGT

APSLEEVMVNVSAPPGGQLVYDSAPNRTDPHVIWAEGAGPGASPRLYSVVGPLGRQRLIIEELTLETQGM

YYWVWGRTDRPSAYGTWVRVRVFRPPSLTIHPHAVLEGQPFKATCTAATYYPGNRAEFVWFEDGRRVFDP

AQIHTQTQENPDGFSTVSTVTSAAVGGQGPPRTFTCQLTWHRDSVSFSRRNASGTASVLPRPTITMEFTG

DHAVCTAGCVPEGVTFAWFLGDDSSPAEKVAVASQTSCGRPGTATIRSTLPVSYEQTEYICRLAGYPHGI

PVLEHHGSHQPPPRDPTERQVIRAVEGAGIGVAVLVAVVLAGTAVVYLTHASSVRYRRLR

>HSV2GPC_24

MALGRVGLAVGLWGLLWVGVVVVLANASPGRTITVGPRGNASNAAPSASPRNASAPRTTPTPPQPRKATK

SKASTAKPAPPPKTGPPKTSSEPVRCNRHDPLARYGSRVQIRCRFPNSTRTESRLQIWRYATATDAEIGT

APSLEEVMVNVSAPPGGQLVYDSAPNRTDPHVIWAEGAGPGASPRLYSVVGPLGRQRLIIEELTLETQGM

YYWVWGRTDRPSAYGTWVRVRVFRPPSLTIHPHAVLEGQPFKATCTAATYYPGNRAEFVWFEDGRRVFDP

AQIHTQTQENPDGFSTVSTVTSAAVGGQGPPRTFTCQLTWHRDSVSFSRRNASGTASVLPRPTITMEFTG

DHAVCTAGCVPEGVTFAWFLGDDSSPAEKVAVASQTSCGRPGTATIRSTLPVSYEQTEYICRLAGYPDGI

PVLEHHGSHQPPPRDPTERQVIRAVEGAGIGVAVLVAVVLAGTAVVYLTHASSVRYRRLR

>HSV2GPC_25

MALGRVGLAVGLWGLLWVGVVVVLANASPGRTITVGPRGNASNAAPSASPRNASAPRTTPTPPQPRKATK

SKASTAKPAPPPKTGPPKTSSEPVRCNRHDPLARYGSRVQIRCRFPNSTRTESRLQIWRYATATDAEIGT

APSLEEVMVNVSAPPGGQLVYDSAPNRTDPHVIWAEGAGPGASPRLYSVVGPLGRQRPIIEELTLETQGM

YYWVWGRTDRPSAYGTWVRVRVFRPPSLTIHPHAVLEGQPFKATCTAATYYPGNRAEFVWFEDGRRVFDP

AQIHTQTQENPDGFSTVSTVTSAAVGGQGPPRTFTCQLTWHRDSVSFSRRNASGTASVLPRPTITMEFTG

DHAVCTAGCVPEGVTFAWFLGDDSSPAEKVAVASQTSCGRPGTATIRSTLPVSYEQTEYICRLAGYPDGI

PVLEHHGSHQPPPRDPTERQVIRAVEGAGIGVAVLVAVVLAGTAVVYLTHASSVRYRRLR

>HSV2GPC_26

MALGRVGLAVGLWGLLWVGVVVVLANASPGRTITVGPRGNASNAAPSASPRNASAPRTTPTPPQPRKATK

SKASTAKPAPPPKTGPPKTSSEPVRCNRHDPLARYGSRVQIRCRFPNSTRTESRLQIWRYATATDAEIGT

APSLEEVMVNVSAPPGGQLVYDSPPNRTDPHVIWAEGAGPGASPRLYSVVGPLGRQRLIIEELTLETQGM

YYWVWGRTDRPSAYGTWVRVRVFRPPSLTIHPHAVLEGQPFKATCTAATYYPGNRAEFVWFEDGRRVFDP

AQIHTQTQENPDGFSTVSTVTSAAVGGQGPPRTFTCQLTWHRDSVSFSRRNASGTASVLPRPTITMEFTG

DHAVCTAGCVPEGVTFAWFLGDDSSPAEKVAVASQTSCGRPGTATIRSTLPVSYEQTEYICRLAGYPDGI

PVLEHHGSHQPPPRDPTERQVIRAVEGAGIGVAVLVAVVLAGTAVVYLTHASSVRYRRLR

>HSV2GPC_27

MALGRVGLTVGLWGLLWVGVVVVLANASPGRTITVGPRGNASNAAPSVPRNRSAPRTTPTPPQPRKATKS

KASTAKPAPPPKTGPPKTSSEPVRCNRHDPLARYGSRVQIRCRFPNSTRTESRLQIWRYATATDAEIGTA

PSLEEVMVNVSAPPGGQLVYDSAPNRTDPHVIWAEGAGPGASPRLYSVVGPLGRQRLIIEELTLETQGMY

YWVWGRTDRPSAYGTWVRVRVFRPPSLTIHPHAVLEGQPFKATCTAATYYPGNRAEFVWFEDGRRVFDPA

QIHTQTQENPDGFSTVSTVTSAAVGGQGPPRTFTCQLTWHRDSVSFSRRNASGTASVLPRPTITMEFTGD

HAVCTAGCVPEGVTFAWFLGDDSSPAEKVAVASQTSCGRPGTATIRSTLPVSYEQTEYICRLAGYPDGIP

VLEHHGSHQPPPRDPTERQVIRAVEGAGIGVAVLVAVVLAGTAVVYLTHASSVRYRRLR
