## Supplementary file 3 for "Contriving a chimeric polyvalent vaccine to prevent infections caused by Herpes Simplex Virus (Type-1 and Type-2): an exploratory immunoinformatic approach"

**GLYCOPROTEIN D**

**HSV1=20 AND HSV2=8**

>HSV1GPD_1

MGGAAARLGAVILFVVIVGLHGVRSKYALVDASLKMADPNRFRGKDLPVLDQLTDPPGVRRVYHIQAGLP

DPFQPPSLPITVYYAVLERACRSVLLNAPSEAPQIVRGASEDVRKQPYNLTIAWFRMGGNCAIPITVMEY

TECSYNKSLGACPIRTQPRWNYYDSFSAVSEDNLGFLMHAPAFETAGTYLRLVKINDWTEITQFILEHRA

KGSCKYALPLRIPPSACLSPQAYQQGVTVDSIGMLPRFIPENQRTVAVYSLKIAGWHGPKAPYTSTLLPP

ELSETPNATQPELAPEDPEDSALLEDPVGTVAPQIPPNWHIPSIQDAATPYHPPATPNNMGLIAGAVGGS

LLAALVICGIVYWMRRHTQKAPKRIRLPHIREDDQPSSHQPLFY

>HSV1GPD_2

MGGAAARLGAVILFVVIVGLHGVRGKYALVDASLKMADPNRFRGKDLPVLDQLTDPPGVRRVYHIQAGLP

DPFQPPSLPITVYYAVLERACRSVLLNAPSEAPQIVRGASEDVRKQPYNLTIAWFRMGGNCAIPITVMEY

TECSYNKSLGACPIRTQPRWNYYDSFSAVSEDNLGFLMHAPAFETAGTYLRLVKINDWTEITQFILEHRA

KGSCKYALPLRIPPSACLSPQAYQQGVTVDSIGMLPRFIPENQRTVAVYSLKIAGWHGPKAPYTSTLLPP

ELSETPNATQPELAPEDPEDSALLEDPVGTVAPQIPPNWHIPSIQDAATPYHPPATPNNMGLIAGAVGGS

LLAALVICGIVYWMRRHTQKAPKRIRLPHIREDDQPSSHQPLFY

>HSV1GPD_3 MGGAAARLGAVILFVVIVGLHGVRGKYALADASLKMADPNRFRGKDLPVLDQLTDPPGVRRVYHIQAGLP

DPFQPPSLPITVYYAVLERACRSVLLNAPSEAPQIVRGASEDVRKQPYNLTIAWFRMGGNCAIPITVMEY

TECSYNKSLGACPIRTQPRWNYYDSFSAVSEDNLGFLMHAPAFETAGTYLRLVKINDWTEITQFILEHRA

KGSCKYALPLRIPPSACLSPQAYQQGVTVDSIGMLPRFIPENQRTVAVYSLKIAGWHGPKAPYTSTLLPP

ELSETPNATQPELAPEDPEDSALLEDPVGTVAPQIPPNWHIPSIQDAATPYHPPATPNNMGLIAGAVGGS

LLAALVICGIVYWMRRRTQKAPKRIRLPHIREDDQPSSHQPLFY

>HSV1GPD_4

MGGAAARLGAVILFVVIVGLHGVRGKYALADASLKMADPNRFRGKDLPVLDQLTDPPGVRRVYHIQAGLP

DPFQPPSLPITVYYAVLERACRSVLLNAPSEAPQIVRGASEDVRKQPYNLTIAWFRMGGNCAIPITVMEY

TECSYNKSLGACPIRTQPRWNYYDSFSAVSEDNLGFLMHAPAFETAGTYLRLVKINDWTEITQFILEHRA

KGSCKYALPLRIPPSACLSPQAYQQGVTVDSIGMLPRFIPENQRTVAVYSLKIAGWHGPKAPYTSTLLPP

ELSETPNATQPELAPEDPEDSALLEDPVGTVAPQIPPNWHIPSIQDAATPYHPPATPNNMGLIAGAVGGS

LLAALVICGIVYWMRRRTRKAPKRIRLPHIREDDQPSSHQPLFY

>HSV1GPD_5

MGGAAARLGAVILFVVIVGLHGVRGKYALVDASLKMADPNRFRGKDLPVLDQLTDPPGVRRVYHIQAGLP

DPFQPPSLPITVYYAVLERACRSVLLNAPSEAPQIVRGASEDVRKQPYNLTIAWFRMGGNCAIPITVMEY

TECSYNKSLGACPIRTQPRWNYYDSFSAVSEDNLGFLMHAPAFETAGTYLRLVKINDWTEITQFILEHRA

KGSCKYALPLRIPPSACLSPQAYQQGVTVDSIGMLPRFIPENQRTVAVYSLKIAGWHGPKAPYTSTLLPP

ELSETPNATQPELAPEDPEDSALLEDPVGTVAPQIPPNWHIPSIQDAATPYHPPATPNNMGLIAGAVGGS

LLAALVICGIVYWMHRRTRKAPKRIRLPHIREDDQPSSHQPLFY

>HSV1GPD_6

MGGAAARLGAVILFVVIVGLHGVRGKYALADASLKMADPNRFRGKDLPVLDQLTDPPGVRRVYHIQAGLP

DPFQPPSLPITVYYAVLERACRSVLLNAPSEAPQIVRGASEDVRKQPYNLTIAWFRMGGNCAIPITVMEY

TECSYNKSLGACPIRTQPRWNYYDSFSAVSEDNLGFLMHAPAFETAGTYLRLVKINDWTEITQFILEHRA

KGSCKYALPLRIPPSACLSPQAYQQGVTVDSIGMLPRFIPENQRTVAVYSLKIAGWHGPKAPYTSTLLPP

ELSETPNATQPELTPEDPEDSALLEDPVGTVAPQIPPNWHIPSIQDAATPYHPPATPNNMGLIAGAVGGS

LLAALVICGIVYWMRRRTQKAPKRIRLPHIREDDQPSSHQPLFY

>HSV1GPD_7

MGGAAARLGAVILFVVIVGLHGVRGKYALADASLKMADPNRFRGKDHPVLDQLTDPPGVRRVYHIQAGLP

DPFQPPSLPITVYYAVLERACRSVLLNAPSEAPQIVRGASEDVRKQPYNLTIAWFRMGGNCAIPITVMEY

TECSYNKSLGACPIRTQPRWNYYDSFSAVSEDNLGFLMHAPAFETAGTYLRLVKINDWTEITQFILEHRA

KGSCKYALPLRIPPSACLSPQAYQQGVTVDSIGMLPRFIPENQRTVAVYSLKIAGWHGPKAPYTSTLLPP

ELSETPNATQPELAPEDPEDSALLEDPVGTVAPQIPPNWHIPSIQDAATPYHPPATPNNMGLIAGAVGGS

LLAALVICGIVYWMRRRTQKAPKRIRLPHIREDDQPSSHQPLFY

>HSV1GPD_8

MGGAAARLGAVILFVVLVGLHGVRGKYALADASLKMADPNRFRGKDLPVLDQLTDPPGVRRVYHIQAGLP

DPFQPPSLPITVYYAVLERACRSVLLNAPSEAPQIVRGASEDVRKQPYNLTIAWFRMGGNCAIPITVMEY

TECSYNKSLGACPIRTQPRWNYYDSFSAVSEDNLGFLMHAPAFETAGTYLRLVKINDWTEITQFILEHRA

KGSCKYALPLRIPPSACLSPQAYQQGVTVDSIGMLPRFIPENQRTVAVYSLKIAGWHGPKAPYTSTLLPP

ELSETPNATQPELAPEDPEDSALLEDPVGTVAPQIPPNWHIPSIQDAATPYHPPATPNNMGLIAGAVGGS

LLAALVICGIVYWMRRRTRKAPKRIRLPHIREDDQPSSHQPLFY

>HSV1GPD_9

MGGAAARLGAVILFVVIVGLHGVRGKYALADASLKMADPNRFRGKDLPVLDQLTDPPGVRRVYHIQAGLP

DPFQPPSLPITVYYAVLERACRSVLLNAPSEAPQIVRGASEDVRKQPYNLTIAWFRMGGNCAIPITVMEY

TECSYNKSLGACPIRTQPRWNYYDSFSAVSEDNLGFLMHAPAFETAGTYLRLVKINDWTEITQFILEHRA

KGSCKYAIPLRIPPSACLSPQAYQQGVTVDSIGMLPRFIPENQRTVAVYSLKIAGWHGPKAPYTSTLLPP

ELSETPNATQPELAPEDPEDSALLEDPVGTVAPQIPPNWHIPSIQDAATPYHPPATPNNMGLIAGAVGGS

LLVALVICGIVYWMRRRTQKAPKRIRLPHIREDDQPSSHQPLFY

>HSV1GPD_10

MGGAAARLGAVILFVVIVGLHGVRGKYALADASLKMADPNRFRGKDLPVLDQLTDPPGVRRVYHIQAGLP

DPFQPPSLPITVYYAVLERACRSVLLNAPSEAPQIVRGASEDVRKQPYNLTIAWFRMGGNCAIPITVMEY

TECSYNKSLGACPIRTQPRWNYYDSFSAVSEDNLGFLMHAPAFETAGTYLRLVKINDWTEITQFILEHRA

KGSCKYALPLRIPPSACLSPQAYQQGVTVDSIGMLPRFIPENQRTVAVYSLKIAGWHGPKAPYTSTLLPP

ELSETPNATQPELAPEDPEDSALLEDPVGTVAPQIPPNWHIPSIQDAATPYHPPATPNNMGLIAGAVGGS

LLAALVICGIVYWMHRRTRKAPKRIRLPHIREDDQPSSHQPLFY

>HSV1GPD_11

MGGAAARLGAVILFVVIVGLHGVRGKYALADASLKMADPNRFRGKDLPVLDQLTDPPGVRRVYHIQAGLP

DPFQPPSLPITVYYAVLERACRSVLLNAPSEAPQIVRGASEDVRKQLYNLTIAWFRMGGNCAIPITVMEY

TDCSYNKSLGACPIRTQPRWNYYDSFSAVSEDNLGFLMHAPAFETAGTYLRLVKINDWTEITQFILEHRA

KGSCKYALPLRIPPSACLSPQAYQQGVTVDSIGMLPRFIPENQRTVAVYSLKIAGWHGPKAPYTSTLLPP

ELSETPNATQPELAPEDPEDSALLEDPVGTVAPQIPPNWHIPSIQDAATPYHPPATPNNMGLIAGAVGGS

LLAALVICGIVYWMRRRTQKAPKRIRLPHIREDDQPSSHQPLFY

>HSV1GPD_12

MGGAAARLGAVILFVVIVGLHGVRGKYALADASLKMADPNRFRGKDLPVLDQLTDPPGVRRVYHIQAGLP

DPFQPPSLPITVYYAVLERACRSVLLNAPSEAPQIVRGASEDVRKQPYNLTIAWFRMGGNCAIPITVMEY

TECSYNKSLGACPIRTQPRWNYYDSFSAVSEDNLGFLMHAPAFETAGTYLRLVKINDWTEITQFILEHRA

KGSCKYALPLRIPPSACLSPQAYQQGVTVDSIGMLPRFIPENQRTVAVYSLKIAGWHGPKAPYTSTLLPP

ELSETPNATQPELAPEDPEDSALLEDPVGTVAPQIPPNWHIPSIQDAATPYHPPATPNNMGLIAGAVGGS

LLAALVICGIVYRMRRRTQKAPKRIRLPHIREDDQPSSHQPLFY

>HSV1GPD_13

MGGAAARLGAVILFVVIVGLHGVRGKYALADASLKMADPNRFRGKDLPVLDRLTDPPGVRRVYHIQAGLP

DPFQPPSLPITVYYAVLERACRSVLLNAPSEAPQIVRGASEDVRKQPYNLTIAWFRMGGNCAIPITVMEY

TECSYNKSLGACPIRTQPRWNYYDSFSAVSEDNLGFLMHAPAFETAGTYLRLVKINDWTEITQFILEHRA

KGSCKYALPLRIPPSACLSPQAYQQGVTVDSIGMLPRFIPENQRTVAVYSLKIAGWHGPKAPYTSTLLPP

ELSETPNATQPELAPEDPEDSALLEDPVGTVAPQIPPNWHIPSIQDAATPYHPPATPNNMGLIAGAVGGS

LLAALVICGIVYWMHRRTRKAPKRIRLPHIREDDQPSSHQPLFY

>HSV1GPD_14

MGGTAARLGAVILFVVIVGLHGVRGKYALADASLKMADPNRFRGKDLPVLDQLTDPPGVRRVYHIQAGLP

DPFQPPSLPITVYYAVLERACRSVLLNAPSEAPQIVRGASEDVRKQPYNLTIAWFRMGGNCAIPITVMEY

TECSYNKSLGACPIRTQPRWNYYDSFSAVSEDNLGFLMHAPAFETAGTYLRLVKINDWTEITQFILEHRA

KGSCKYALPLRIPPSACLSPQAYQQGVTVDSIGMLPRFIPENQRTVAVYSLKIAGWHGPKAPYTSTLLPP

ELSETPNATQPELAPEDPEDSALLEDPVGTVAPQIPPNWHIPSIQDAATPYHPPATPNNMGLIAGAVGGS

LLAALVICGIVYWMHRRTRKAPKRIRLPHIREDDQPSSHQPLFY

>HSV1GPD_15

MGGAAARLGAVILFVVIVGLHGVRGKYALADASLKLADPNRFRRKDLPVLDQLTDPPGVRRVYHIQAGLP

DPFQPPSLPITVYYAVLERACRSVLLNAPSEAPQIVRGASEDVRKQPYNLTIAWFRMGGNCAIPITVMEY

TECSYNKSLGACPIRTQPRWNYYDSFSAVSEDNLGFLMHAPAFETAGTYLRLVKINDWTEITQFILEHRA

KGSCKYALPLRIPPSACLSPQAYQQGVTVDSIGMLPRFIPENQRTVAVYSLKIAGWHGPKAPYTSTLLPP

ELSETPNATQPELAPEAPEDSALLEDPVGTVAPQIPPNWHIPSIQDAATPYHPPATPNNMGLIAGAVGGS

LLAALVICGIVYWMRRRTQKAPKRIRLPHIREDDQPSSHQPLFY

>HSV1GPD_16

MGGAAARLGAVILFVVIVGLHGVRGKYALADASLKMADPNRFRGKDLPVLDQLTDPPGVRRVYHIQAGLP

DPFQPPSLPITVYYAVLERACRSVLLNAPSEAPQIVRGASEDVRKQPYNLTIAWFRMGGNCAIPITVMEY

TECSYNKSLGACPIRTQPRWNYYDSFSAVSEDNLGFLMHAPAFETAGTYLRLVKINDWTEITQFILEHRA

KGSCKYALPLRIPPSACLSPQAYQQGVTVDSIGMLPRFIPENQRTVAVYSLKIAGWHGPKAPYTSTLLPP

ELSETPNATQPELAPEDPEDSALLEDPVGTVALQIPPNWHIPSIQDAATPYHPPATPNNMGLIAGAVGGS

LLAALVICGIVYWMHRRTRKAPKRIRLPHIREDDQPSSHQPLFY

>HSV1GPD_17

MGGAAARLGAVILFVVIVGLHGVRGKYALADASLKMADPNRFRGKDLPVPDQLTDPPGVRRVYHIQAGLP

DPFQPPSLPITVYYAVLERACRSVLLNAPSEAPQIVRGASEDVRKQPYNLTIAWFRMGGNCAIPITVMEY

TECSYNKSLGACPIRTQPRWNYYDSFSAVSEDNLGFLMHAPAFETAGTYLRLVKINDWTEITQFILEHRA

KGSCKYALPLRIPPSACLSPQAYQQGVTVDSIGMLPRFIPENQRTVAVYSLKIAGWHGPKAPYTSTLLPP

ELSETPNATQPELAPEDPEDSALLEDPVGTVAPQIPPNWHIPSIQDAATPYHPPATPNNMGLIAGAVGGS

LLAALVICGIVYWMHRRTRKAPKRIRLPHIREDDQPSSHQPLFY

>HSV1GPD_18

MGGAAARLGAVILFVVIVGLHGVRGKYALADASLKMADPNRFRGKDLPVLDPLTDPPGVRRVYHIQAGLP

DPFQPPSLPITVYYAVLERACRSVLLNAPSEAPQIVRGASEDVRKQPYNLTIAWFRMGGNCAIPITVMEY

TECSYNKSLGACPIRTQPRWNYYDSFSAVSEDNLGFLMHAPAFETAGTYLRLVKINDWTEITQFILEHRA

KGSCKYALPLRIPPSACLSPQAYQQGVTVDSIGMLPRFIPENQRTVAVYSLKIAGWHGPKAPYTSTLLPP

ELSETPNATQPELAPEDPEDSALLEDPVGTVAPQIPPNWHIPSIQDAATPYHPPATPNNMGLIAGAVGGS

LLAALVICGIVYWMHRRTRKAPKRIRLPHIREDDQPSSHQPLFY

>HSV1GPD_19

MGGAAARLGAVILFVVIVGLHGVRGKYALADASLKMADPNRFRGKDLPVPDRLTDPPGVRRVYHIQAGLP

DPFQPPSLPITVYYAVLERACRSVLLNAPSEAPQIVRGASEDVRKQPYNLTIAWFRMGGNCAIPITVMEY

TECSYNKSLGACPIRTQPRWNYYDSFSAVSEDNLGFLMHAPAFETAGTYLRLVKINDWTEITQFILEHRA

KGSCKYALPLRIPPSACLSPQAYQQGVTVDSIGMLPRFIPENQRTVAVYSLKIAGWHGPKAPYTSTLLPP

ELSETPNATQPELAPEDPEDSALLEDPVGTVVPQIPPNWHIPSIQDAATPYHPPATPNNMGLIAGAVGGS

LLVALVICGIVYWMRRRTQKAPKRIRLPHIREDDQPSSHQPLFY

>HSV1GPD_20

MGGAAARLGAVILFVVIVGLHGVRGKYALADASLKMADPNRFRGKDLPVPDRLTDPPGVRRVYHIQAGLP

DPFQPPSLPITVYYAVLERACRSVLLNAPSEAPQIVRGASEDVRKQPYNLTIAWFRMGGNCAIPITVMEY

TECSYNKSLGACPIRTQPRWNYYDSFSAVSEDNLGFLMHAPAFETAGTYLRLVKINDWTEITQFILEHRA

KGSCKYALPLRIPPSACLSPQAYQQGVTVDSIGMLPRFIPENQRTVAVYSLKIAGWHGPKAPYTSTLLPP

ELSETPNATQPELAPEDPEDSALLEDPVGTVAPQIPPNWHIPSIQDAATPYHPPATPNNMGLIAGAVGGS

LLAALVICGIVYWMHRRTRKAPKRIRLPHIREDDQPSSHQPLFY

>HSV2GPD_21

MGRLTSGVGTAALLVVAVGLRVVCAKYALADPSLKMADPNRFRGKNLPVLDQLTDPPGVKRVYHIQPSLE

DPFQPPSIPITVYYAVLERACRSVLLHAPSEAPQIVRGASDEARKHTYNLTIAWYRMGDNCAIPITVMEY

TECPYNKSLGVCPIRTQPRWSYYDSFSAVSEDNLGFLMHAPAFETAGTYLRLVKINDWTEITQFILEHRA

RASCKYALPLRIPPAACLTSKAYQQGVTVDSIGMLPRFIPENQRTVALYSLKIAGWHGPKPPYTSTLLPP

ELSDTTNATQPELVPEDPEDSALLEDPAGTVSSQIPPNWHIPSIQDVAPHHAPAAPSNPGLIIGALAGST

LAVLVIGGIAFWVRRRAQMAPKRLRLPHIRDDDAPPSHQPLFY

>HSV2GPD_22

MGRLTSGVGTAALLVVAVGLRVVCAKYALADPSLKMADPNRFRGKNLPVLDQLTDPPGVKRVYHIQPSLE

DPFQPPSIPITVYYAVLERACRSVLLHAPSEAPQIVRGASDEARKHTYNLTIAWYRMGDNCAIPITVMEY

TECPYNKSLGVCPIRTQPRWSYYDSFSAVSEDNLGFLIHAPAFETAGTYLRLVKINDWTEITQFILEHRA

RASCKYALPLRIPPAACLTSKAYQQGVTVDSIGMLPRFIPENQRTVALYSLKIAGWHGPKPPYTSTLLPP

ELSDTTNATQPELVPEDPEDSALLEDPAGTVSSQIPPNWHIPSIQDVAPHHAPAAPSNPGLIIGALAGST

LAVLVIGGIAFWVRRRAQMAPKRLRLPHIRDDDAPPSHQPLFY

>HSV2GPD_23

MGRLTSGVGTAALLVVAVGLRVVCAKYALADPSLKMADPNRFRGKNLPVLDQLTDPPGVKRVYHIQPSLE

DPFQPPSIPITVYYAVLERACRSVLLHAPSEAPQIVRGASDEARKHTYNLTIAWYRMGDNCAIPITVMEY

TECPYNKSLGVCPIRTQPRWSYYDSFSAVSEDNLGFLMHAPAFETAGTYLRLVKINDWTEITQFILEHRA

RASCKYALPLRIPPAACLTSKAYQQGVTVDSIGMLPRFIPENQRTVALYSLKIAGWHGPKPPYTSTLLPP

ELSDTTNATQPELVPEDPEDSALLEDPAGTVSSQIPPNWHIPSIQDVAPHHAPAAPSNPGLIIGALAGST

LAALVIGGIAFWVRRRAQMAPKRLRLPHIRDDDAPPSHQPLFY

>HSV2GPD_24

MGRLTSGVGTAALLVVAVGLRVVCAKYALADPSLKMADPNRFRGKNLPVLDQLTDPPGVKRVYHIQPSLE

DPFQPPSIPITVYYAVLERACRSVLLHAPSEAPQIVRGASDEARKHTYNLTIAWYRMGDNCAIPITVMEY

TECPYNKSLGVCPIRTQPRWSYYDSFSAASEDNLGFLMHAPAFETAGTYLRLVKINDWTEITQFILEHRA

RASCKYALPLRIPPAACLTSKAYQQGVTVDSIGMLPRFIPENQRTVALYSLKIAGWHGPKPPYTSTLLPP

ELSDTTNATQPELVPEDPEDSALLEDPAGTVSSQIPPNWHIPSIQDVAPHHAPAAPSNPGLIIGALAGST

LAVLVIGGIAFWVRRRAQMAPKRLRLPHIRDDDAPPSHQPLFY

>HSV2GPD_25

MGRLTSGVGTAALLVVAVGLRVVCAKYALADPSLKMADPNRFRGKNLPVLDRLTDPPGVKRVYHIQPSLE

DPFQPPSIPITVYYAVLERACRSVLLHAPSEAPQIVRGASDEARKHTYNLTIAWYRMGDNCAIPITVMEY

TECPYNKSLGVCPIRTQPRWSYYDSFSAVSEDNLGFLMHAPAFETAGTYLRLVKINDWTEITQFILEHRA

RASCKYALPLRIPPAACLTSKAYQQGVTVDSIGMLPRFIPENQRTVALYSLKIAGWHGPKPPYTSTLLPP

ELSDTTNATQPELVPEDPEDSALLEDPAGTVSSQIPPNWHIPSIQDVAPHHAPAAPSNPGLIIGALAGST

LAVLVIGGIAFWVRRRAQMAPKRLRLPHIRDDDAPPSHQPLFY

>HSV2GPD_26

MGRLTSGVGTAALLVVAVGLRVVCAKYALADPSLKMADPNRFRGKNLPVLDQLTDPPGVKRVYHIQPSLE

DPFQPPSIPITVYYAVLERACRSVLLHAPSEAPQIVRGASDEARKHTYNLTIAWYRMGDNCAIPITVMEY

TECPYNKSLGVCPIRTQPRWSYYDSFSAVSEDTLGFLMHAPAFETAGTYLRLVKINDWTEITQFILEHRA

RASCKYALPLRIPPAACLTSKAYQQGVTVDSIGMLPRFIPENQRTVALYSLKIAGWHGPKPPYTSTLLPP

ELSDTTNATQPELVPEDPEDSALLEDPAGTVSSQIPPNWHIPSIQDVAPHHAPAAPSNPGLIIGALAGST

LAVLVIGGIAFWVRRRAQMAPKRLRLPHIRDDDAPPSHQPLFY

>HSV2GPD_27

MGRLTSGVGTAALLVVAVGLRVVCAKYALADPSLKMADPNRFRGKNLPVLDQLTDPPGVKRVYHIQPSLE

DPFQPPSIPITVYYAVLERACRSVLLHAPSEAPQIVRGASDEARKHTYNLTIAWYRMGDNCAIPITVMEY

TECPYNKSLGVCPIRTQPRWSYYDSFSAVSEDNLGFLMHAPAFETAGTYLRLVKINDWTEITQFILEHRA

RASCKYALPLRIPPAACLTSKAYQQGVTVDSIGMLPRFIPENQRTVALYSLKIAGWHGPKPPYTSTLLPP

ELSDTTNATQPELVPEDPEDSALLEDPAGTVSSQIPPNWHIPSIQDVAPHHAPAAPSNPGLIIGALAGST

LAALVIGGIAFWVRRRAQMAPKRPRLPHIRDDDAPPSHQPLFY

>HSV2GPD_28

MGRLTSGVGTAALLVVAVGLRVVCAKYALADPSLKMADPNRFRGKNLPVLDQLTDPPGVKRVYHIQPSLE

DPFQPPSIPITVYYAVLERACRSVLLHAPSEAPQIVRGASDEARKHTYNLTIAWYRMGDNCAIPITVMEY

TECPYNKSLGVCPIRTQPRWSYYDSFSAVSEDNLGFLMHAPAFETAGTYLRLVKINDWTEITQFILEHRA

RASCKYALPLRIPPAACLTSKAYQQGVTVDSIGMLPRFTPENQRTVALYSLKIAGWHGPKPPYTSTLLPP

ELSDTTNATQPELVPEDPEDSALLEDPAGTVSSQIPPNWHIPSIQDVAPHHAPAAPANPGLIIGALAGST

LAALVIGGIAFWVRRRRSVAPKRLRLPHIRDDDAPPSHQPLFY
