## Supplementary file 4 for "Contriving a chimeric polyvalent vaccine to prevent infections caused by Herpes Simplex Virus (Type-1 and Type-2): an exploratory immunoinformatic approach"

**GLYCOPROTEIN E**

>HSV1GPE_1 MDRGAVVGFLLGVCVVSCLAGTPKTSWRRVSVGEDVSLLPAPGPTGRGPTQKLLWAVEPLDGCGPLHPSW

VSLMPPKQVPETVVDAACMRAPVPLAMAYAPPAPSATGGLRTDFVWQERAAVVNRSLVIHGVRETDSGLY

TLSVGDIKDPARQVASVVLVVQPAPVPTPPPTPADYDEDDNDEGEDESLAGTPASGTPRLPPPPAPPRSW

PSAPEVSHVRGVTVRMETPEAILFSPGETFSTNVSIHAIAHDDQTYSMDVVWLRFDVPTSCAEMRIYESC

LYHPQLPECLSPADAPCAASTWTSRLAVRSYAGCSRTNPPPRCSAEAHMEPVPGLAWQAASVNLEFRDAS

PQHSGLYLCVVYVNDHIHAWGHITISTAAQYRNAVVEQPLPQRGADLAEPTHPHVGAPPHAPPTHGALRL

GAVMGAALLLSALGLSVWACMTCWRRRAWRAVKSRASGKGPTYIRVADSELYADWSSDSEGERDQVPWLA

PPERPDSPSTNGSGFEILSPTAPSVYPRSDGHQSRRQLTTFGSGRPDRRYSQASDSSVFW

>HSV1GPE_2

MDRGAVVGFLLGVCVVSCLAGTPKTSWRRVSVGEDVSLLPAPGPTGRGPTQKLLWAVEPLDGCGPLHPSW

VSLMPPKQVPETVVDAACMRAPVPLAMAYAPPAPSATGGLRTDFVWQERAAVVNRSLVIHGVRETDSGLY

TLSVGDIKDPARQVASVVLVVQPAPVPTPPPTPADYDEDDNDEGEDESLAGTPASGTPRLPPPPAPPRSW

PSAPEVSHVRGVTVRMETPEAILFSPGETFSTNVSIHAIAHDDQTYAMDVVWLRFDVPTSCAEMRIYESC

LYHPQLPECLSPADAPCAASTWTSRLAVRSYAGCSRTNPPPRCSAEAHMEPVPGLAWQAASVNLEFRDAS

PQHSGLYLCVVYVNDHIHAWGHITISTAAQYRNAVVEQPLPQRGADLAEPTHPHVGAPPHAPPTHGALRL

GAVMGAALLLSALGLSVWACMTCWRRRAWRAVKSRASGKGPTYIRVADSELYADWSSDSEGERDQVPWLA

PPERPDSPSTNGSGFEILSPTAPSVYPRSDGHQSRRQLTTFGSGRPDRRYSQASDSSVFW

>HSV1GPE_3

MDRGAVVGFLLGVCVVSCLAGTPKTSWRRVSVGEDVSLLPAPGPTGRGPTQKLLWAVEPLDGCGPLHPSW

VSLMPPKQVPETVVDAACMRAPVPLAMAYAPPAPSATGGLRTDFVWQERAAVVNRSLVIHGVRETDSGLY

TLSVGDIKDPARQVASVVLVVQPAPVPTPPPTPADYDEDDNDEGEDESLAGTPASGTPRLPPPPAPPRSW

PSAPEVSHVRGVTVRMETPEAILFSPGETFSTNVSIHSIAHDDQTYSMDVVWLRFDVPTSCAEMRIYESC

LYHPQLPECLSPADAPCAASTWTSRLAVRSYAGCSRTNPPPRCSAEAHMEPVPGLAWQAASVNLEFRDAS

PQHSGLYLCVVYVNDHIHAWGHITISTAAQYRNAVVEQPLPQRGADLAEPTHPHVGAPPHAPPTHGALRL

GAVMGAALLLSALGLSVWACMTCWRRRAWRAVKSRASGKGPTYIRVADSELYADWSSDSEGERDQVPWLA

PPERPDSPSTNGSGFEILSPTAPSVYPRSDGHQSRRQLTTFGSGRPDRRYSQASDSSVFW

>HSV1GPE_4

MDRGAVVGFLLGVCVVSCLAGTPKTSWRRVSVGEDVSLLPAPGPTGRGPTQKLLWAVEPLDGCGPLHPSW

VSLMPPKQVPETVVDAACMRAPVPLAMAYAPPAPSATGGLRTDFVWQERAAVVNRSLVIHGVRETDSGLY

TLSMGDIKDPARQVASVVLVVQPAPVPTPPPTPADYDEDDNDEGEDESLAGTPASGTPRLPPPPAPPRSW

PSAPEVSHVRGVTVRMETPEAILFSPGETFSTNVSIHAIAHDDQTYSMDVVWLRFDVPTSCAEMRIYESC

LYHPQLPECLSPADAPCAASTWTSRLAVRSYAGCSRTNPPPRCSAEAHMEPVPGLAWQAASVNLEFRDAS

PQHSGLYLCVVYVNDHIHAWGHITISTAAQYRNAVVEQPLPQRGADLAEPTHPHVGAPPHAPPTHGALRL

GAVMGAALLLSALGLSVWACMTCWRRRAWRAVKSRASGKGPTYIRVADSELYADWSSDSEGERDQVPWLA

PPERPDSPSTNGSGFEILSPTAPSVYPRSDGHQSRRQLTTFGSGRPDRRYSQASDSSVFW

>HSV1GPE_5

MDRGAVVGFLLGVCVVSCLAGTPKTSWRRVSVGEDVSLLPAPGPTGRGPTQKLLWAVEPLDGCGPLHPSW

VSLMPPKQVPETVVDAACMRAPVPLAMAYAPPAPSATGGLRTDFVWQERAAVVNRSLVIHGVRETDSGLY

TLSVGDIKDPARQVASVVLVVQPAPVPTPPPTPADYDEDDNDEGEDESLAGTPASGTPRLPPPPAPPRSW

PSAPEVSHVRGVTVRMETPEAILFSPGETFSTNVSIHAIAHDDQTYSMDVVWLRFDVPTSCAEMRIYESC

LYHPQLPECLSPADAPCAASTWTSRLAVRSYAGCSRTNPQPRCSAEAHMEPVPGLAWQAASVNLEFRDAS

PQHSGLYLCVVYVNDHIHAWGHITISTAAQYRNAVVEQPLPQRGADLAEPTHPHVGAPPHAPPTHGALRL

GAVMGAALLLSALGLSVWACMTCWRRRAWRAVKSRASGKGPTYIRVADSELYADWSSDSEGERDQVPWLA

PPERPDSPSTNGSGFEILSPTAPSVYPRSDGHQSRRQLTTFGSGRPDRRYSQASDSSVFW

>HSV1GPE_6

MDRGAVVGFLLGVCVVSCLAGTPKTSWRRVSVGEDVSLLPAPGPTGRGPTQKLLWAVEPLDGCGPLHPSW

VSLMPPKQVPETVVDAACMRAPVPLAMAYAPPAPSATGGLRTDFVWQERAAVVNRSLVIYGVRETDSGLY

TLSVGDIKDPARQVASVVLVVQPAPVPTPPPTPADYDEDDNDEGEDESLAGTPASGTPRLPPPPAPPRSW

PSAPEVSHVRGVTVRMETPEAILFSPGETFSTNVSIHAIAHDDQTYAMDVVWLRFDVPTSCAEMRIYESC

LYHPQLPECLSPADAPCAASTWTSRLAVRSYAGCSRTNPPPRCSAEAHMEPVPGLAWQAASVNLEFRDAS

PQHSGLYLCVVYVNDHIHAWGHITISTAAQYRNAVVEQPLPQRGADLAEPTHPHVGAPPHAPPTHGALRL

GAVMGAALLLSALGLSVWACMTCWRRRAWRAVKSRASGKGPTYIRVADSELYADWSSDSEGERDQVPWLA

PPERPDSPSTNGSGFEILSPTAPSVYPRSDGHQSRRQLTTFGSGRPDRRYSQASDSSVFW

>HSV1GPE_7

MDRGAVVGFLLGVCVVSCLAGTPKTSWRRVSVGEDVSLLPAPGPTGRGPTQKLLWAVEPLDGCGPLHPSW

VSLMPPKQVPETVVDAACMRAPVPLAMAYAPPAPSATGGLRTDFVWQERAAVVNRSLVIHGVRETDSGLY

TLSVGDIKDPARQVASVVLVVQPVPVPTPPPTPADYDEDDNDEGEDESLAGTPASGTPRLPPPPAPPRSW

PSAPEVSHVRGVTVRMETPEAILFSPGETFSTNVSIHAIAHDDQTYAMDVVWLRFDVPTSCAEMRIYESC

LYHPQLPECLSPADAPCAASTWTSRLAVRSYAGCSRTNPPPRCSAEAHMEPVPGLAWQAASVNLEFRDAS

PQHSGLYLCVVYVNDHIHAWGHITISTAAQYRNAVVEQPLPQRGADLAEPTHPHVGAPPHAPPTHGALRL

GAVMGAALLLSALGLSVWACMTCWRRRAWRAVKSRASGKGPTYIRVADSELYADWSSDSEGERDQVPWLA

PPERPDSPSTNGSGFEILSPTAPSVYPRSDGHQSRRQLTTFGSGRPDRRYSQASDSSVFW

>HSV1GPE_8

MDRGAVVGFLLGVCVVSCLAGTPKTSWRRVSVGEDVSLLPAPGPTGRGPTQKLLWAVEPLDGCGPLHPSW

VSLMPPKQVPETVVDAACMRAPVPLAMAYAPPAPSATGGLRTDFVWQERAAVVNRSLVIHGVRETDSGLY

TLSVGDIKDPARQVASVVLVVQPAPVPTPPPTPADYDEEDNDEGEDESLAGTPASGTPRLPPPPAPPRSW

PSAPEVSHVRGVTVRMETPEAILFSPGETFSTNVSIHAIAHDDQTYAMDVVWLRFDVPTSCAEMRIYESC

LYHPQLPECLSPADAPCAASTWTSRLAVRSYAGCSRTNPPPRCSAEAHMEPVPGLAWQAASVNLEFRDAS

PQHSGLYLCVVYVNDHIHAWGHITISTAAQYRNAVVEQPLPQRGADLAEPTHPHVGAPPHAPPTHGALRL

GAVMGAALLLSALGLSVWACMTCWRRRAWRAVKSRASGKGPTYIRVADSELYADWSSDSEGERDQVPWLA

PPERPDSPSTNGSGFEILSPTAPSVYPRSDGHQSRRQLTTFGSGRPDRRYSQASDSSVFW

>HSV1GPE_9

MDRGAVVGFLLGVCVVSCLAGTPKTSWRRVSVGEDVSLLPAPGPTGRGPTQKLLWAVEPLDGCGPLHPSW

VSLMPPKQVPETVVDAACMRAPVPLAMAYAPPAPSATGGLRTDFVWQERAAVVNRSLVIYGVRETDSGLY

TLSVGDIKDPARQVASVVLVVQPAPVPTPPPTPADYDEDDNDEGEDESLAGTPASGTPRLPPPPAPPRSW

PSAPEVSHVRGVTVRMETPEAILFSPGETFSTNVSIHAIAHDDQTYAMDVVWLRFDVPTSCAEMRIYESC

LYHPQLPECLSPADAPCAASTWTSRLAVRSYAGCSKTNPPPRCSAEAHMEPVPGLAWQAASVNLEFRDAS

PQHSGLYLCVVYVNDHIHAWGHITISTAAQYRNAVVEQPLPQRGADLAEPTHPHVGAPPHAPPTHGALRL

GAVMGAALLLSALGLSVWACMTCWRRRAWRAVKSRASGKGPTYIRVADSELYADWSSDSEGERDQVPWLA

PPERPDSPSTNGSGFEILSPTAPSVYPRSDGHQSRRQLTTFGSGRPDRRYSQASDSSVFW

>HSV1GPE_10

MDRGAVVGFLLGVCVVSCLAGTPKTSWRRVSVGEDVSLLPAPGPTGRGPTQKLLWAVEPLDGCGPLHPSW

VSLMPPKQVPETVVDAACMRAPVPLAMAYAPPAPSATGGLPTDFVWQERAAVVNRSLVIHGVRETDSGLY

TLSVGDIKDPARQVASVVLVVQPAPVPTPPPTPADYDEDDNDEGEDESLAGTPASGTPRLPPPPAPPRSW

PSAPEVSHVRGVTVRMETPEAILFSPGETFSTNVSIHAIAHDDQTYSMDVVWLRFDVPTSCAEMRIYESC

LYHPQLPECLSPADAPCAASTWTSRLAVRSYAGCSRTNPPPRCSAEAHMEPVPGLAWQAASVNLEFRDAS

PQHSGLYLCVVYVNDHIHAWGHITISTAAQYRNAVVEQPLPQRGADLAEPTHPHVGAPPHAPPTHGALRL

GAVMGAALLLSALGLSVWACMTCWRRRAWRAVKSRASGKGPTYIRVADSELYADWSSDSEGERDQVPWLA

PPERPDSPSTNGSGFEILSPTAPSIYPRSDGHQSRRQLTTFGSGRPDRRYSQASDSSVFW

>HSV1GPE_11

MDRGAVVGFLLGVCVVSCLAGTPKTSWKRVSVGEDVSLLPAPGPTGRGPTQKLLWAVEPLDGCGPLHPSW

VSLMPPKQVPETVVDAACMRAPVPLAMAYAPPAPSATGGLRTDFVWQERAAVVNRSLVIYGVRETDSGLY

TLSVGDIKDPARQVASVVLVVQPAPVPTPPPTPADYDEDDNDEGEDESLAGTPASGTPRLPPPPAPPRSW

PSAPEVSHVRGVTVRMETPEAILFSPGETFSTNVSIHAIAHDDQTYAMDVVWLRFDVPTSCAEMRIYESC

LYHPQLPECLSPADAPCAASTWTSRLAVRSYAGCSRTNPPPRCSAEAHMEPVPGLAWQAASVNLEFRDAS

PQHSGLYLCVVYVNDHIHAWGHITISTAAQYRNAVVEQPLPQRGADLAEPTHPHVGAPPHAPPTHGALRL

GAVMGAALLLSALGLSVWACMTCWRRRAWRAVKSRASGKGPTYIRVADSELYADWSSDSEGERDQVPWLA

PPERPDSPSTNGSGFEILSPTAPSVYPRSDGHQSRRQLTTFGSGRPDRRYSQASDSSVFW

>HSV1GPE_12

MDRGAVVGFLLGVCVVSCLAGTPKTSWRRVSVGEDVSLLPAPGPTGRGPTQKLLWAVEPLDGCGPLHPSW

VSLMPPKQVPETVVDAACMRAPVPLAMAYAPPAPSATGGLPTDFVWQERAAVVNRSLVIHGVRETDSGLY

TLSVGDIKDPARQVASVVLVVQPAPVPTPPPTPADYDEDDNDEGEDESLAGTPASGTPRLPPPPAPPRSW

PSAPEVSHVRGVTVRMETPEAILFSPGETFSTNVSIHAIAHDDQTYAMDVVWLRFDVPTSCAEMRIYESC

LYHPQLPECLSPADAPCAASTWTSRLAVRSYAGCSRTNPPPRCSAEAHMEPVPGLAWQAASVNLEFRDAS

PQHSGLYLCVVYVNDHIHAWGHITISTAAQYRNAVVEQPLPQRGADLAEPTHPHIGAPPHAPPTHGALRL

GAVMGAALLLSALGLSVWACMTCWRRRAWRAVKSRASGKGPTYIRVADSELYADWSSDSEGERDQVPWLA

PPERPDSPSTNGSGFEILSPTAPSVYPRSDGHQSRRQLTTFGSGRPDRRYSQASDSSVFW

>HSV1GPE_13

MDRGAVVGFLLGVCVVSCLAGTPKTSWRRVSVGEDVSLLPAPGPTGRGPTQKLLWAVEPLDGCGPLHPSW

VSLMPPQQVPETVVDAACMRAPVPLAMAYAPPAPSATGGLRTDFVWQERAAVVNRSLVIHGVRETDSGLY

TLSVGDIKDPARQVASVVLVVQPAPVPTPPPTPADYDEEDNDEGEDESLAGTPASGTPRLPPPPAPPRSW

PSAPEVSHVRGVTVRMETPEAILFSPGETFSTNVSIHAIAHDDQTYAMDVVWLRFDVPTSCAEMRIYESC

LYHPQLPECLSPADAPCAASTWTSRLAVRSYAGCSRTNPPPRCSAEAHMEPVPGLAWQAASVNLEFRDAS

PQHSGLYLCVVYVNDHIHAWGHITISTAAQYRNAVVEQPLPQRGADLAEPTHPHVGAPPHAPPTHGALRL

GAVMGAALLLSALGLSVWACMTCWRRRAWRAVKSRASGKGPTYIRVADSELYADWSSDSEGERDQVPWLA

PPERPDSPSTNGSGFEILSPTAPSVYPRSDGHQSRRQLTTFGSGRPDRRYSQASDSSVFW

>HSV1GPE_14

MDRGAVVGFLLGVCVVSCLAGTPKTSWRRASVGEDVSLLPAPGPTGRGPTQKLLWAVEPLDGCGPLHPSW

VSLMPPKQVPETVVDAACMRAPVPLAMAYAPPAPSATGGLRTDFVWQERAAVVNRSLVIHGVRETDSGLY

TLSVGDIKDPARQVASVVLVVQPAPVPTPPPTPADYDEEDNDEGEDESLAGTPASGTPRLPPPPAPPRSW

PSAPEVSHVRGVTVRMETPEAILFSPGETFSTNVSIHAIAHDDQTYAMDVVWLRFDVPTSCAEMRIYESC

LYHPQLPECLSPADAPCAASTWTSRLAVRSYAGCSRTNPPPRCSAEAHMEPVPGLAWQAASVNLEFRDAS

PQHSGLYLCVVYVNDHIHAWGHITISTAAQYRNAVVEQPLPQRGADLAEPTHPHVGAPPHAPPTHGALRL

GAVMGAALLLSALGLSVWACMTCWRRRAWRAVKSRASGKGPTYIRVADSELYADWSSDSEGERDQVPWLA

PPERPDSPSTNGSGFEILSPTAPSVYPRSDGHQSRRQLTTFGSGRPDRRYSQASDSSVFW

>HSV1GPE_15

MDRGAVVGFLLGVCVVSCLAGTPKTSWRRVSVGEDVSLLPAPGPTGRGPTQKLLWAVEPLDGCGPLHPSW

VSLMPPKQVPETVVDAACMRAPVPLAMAYAPPAPSATGGLRTDFVWQERAAVVNRSLVIYGVRETDSGLY

TLSVGDIKDPARQVASVVLVVQPAPVPTPPPTPADYDEDDNDEGEDESLAGTPASGTPRLPPPPAPPRSW

PSAPEVSHVRGVTVRMETPEAILFSPGETFNTNVSIHAIAHDDQTYAMDVVWLRFDVPTSCAEMRIYESC

LYHPQLPECLSPADAPCAASTWTSRLAVRSYAGCSRTNPPPRCSAEAHMEPVPGLAWQAASVNLEFRDAS

PQHSGLYLCVVYVNDHIHAWGHITISTAAQYRNAVVEQPLPQRGADLAEPTHPHVGAPPHATPTHGALRL

GAVMGAALLLSALGLSVWACMTCWRRRAWRAVKSRASGKGPTYIRVADSELYADWSSDSEGERDQVPWLA

PPERPDSPSTNGSGFEILSPTAPSVYPRSDGHQSRRQLTTFGSGRPDRRYSQASDSSVFW

>HSV1GPE_16

MDRGAVVGFLLGVCVVSCLAGTPKTSWRRVSVGEDVSLLPAPGPTGRGPTQKLLWAVEPLDGCGPLHPSW

VSLMPPKQVPETVVDAACMRAPVPLAMAYAPPAPSATGGLRTDFVWQERTAVVNRSLVIYGVRETDSGLY

TLSVGDIKDPARQVASVVLVVQPAPVPTPPPTPADYDEDDNDEGEDESLAGTPASGTPRLPPPPAPPRSW

PSAPEVSHVRGVTVRMETPEAILFSPGEAFSTNVSIHAIAHDDQTYAMDVVWLRFDVPTSCAEMRIYESC

LYHPQLPECLSPADAPCAASTWTSRLAVRSYAGCSRTNPPPRCSAEAHMEPVPGLAWQAASVNLEFRDAS

PQHSGLYLCVVYVNDHIHAWGHITISTAAQYRNAVVEQPLPQRGADLAEPTHPHVGAPPHAPPTHGALRL

GAVMGAALLLSALGLSVWACMTCWRRRAWRAVKSRASGKGPTYIRVADSELYADWSSDSEGERDQVPWLA

PPERPDSPSTNGSGFEILSPTAPSVYPRSDGHQSRRQLTTFGSGRPDRRYSQASDSSVFW

>HSV1GPE_17

MDRGAVVGFLLGVCVVSCLAGTPKTSWRRVSVGEDVSLLPAPGPTGRGPTQKLLWAVEPLDGCGPLHPSW

VSLMPPKQVPETVVDAACMRAPVPLAMAYAPPAPSATGGLRTDFVWQERAAVVNRSLVIYGVRETDSGLY

TLSVGDIKDPARQVASVVLVVQPAPVPTPPPTPADYDEDDNDEGEDESLAGTPASGTPRLPPPPAPPRSW

PSAPEVSHVRGVTVRMETPEAILFSPGEAFSTNVSIHAIAHDDQTYTMDVVWLRFDVPTSCAEMRIYESC

LYHPQLPECLSPADAPCAASTWTSRLAVRSYAGCSRTNPPPRCSAEAHMEPFPGLAWQAASVNLEFRDAS

PQHSGLYLCVVYVNDHIHAWGHITISTAAQYRNAVVEQPLPQRGADLAEPTHPHVGAPPHAPPTHGALRL

GAVMGAALLLSALGLSVWACMTCWRRRAWRAVKSRASGKGPTYIRVADSELYADWSSDSEGERDQVPWLA

PPERPDSPSTNGSGFEILSPTAPSVYPRSDGHQSRRQLTTFGSGRPDRRYSQASDSSVFW

>HSV1GPE_18

MDRGAVVGFLLGVCVVSCLAGTPKTSWRRVSVGEDVSLLPAPGPTGRGPTQKLLWAVEPLDGCGPLHPSW

VSLMPPKQVPETVVDAACMRAPVPLAMAYAPPAPSATGGLRTDFVWQERAAVVNRSLVIYGVRETDSGLY

TLSVGDIKDPARQVASVVLVVQPAPVPTPPPTPADYDEDDNDEGEDESLAGTPASGTPRLPPPPAPPRSW

PSAPEVSHVRGVTVRMETPEAILFSPGEAFSTNVSIHAIAHDDQTYTMDVVWLRFDVPTSCAEMRIYESC

LYHPQLPECLSPADAPCAASTWTSRLAVRSYAGCSRTNPPPRCSAEAHMEPFPGLAWQAASVNLEFRDAS

PQHSGLYLCVVYVNDHIHAWGHITISTAAQYRNAVVEQPLPQRGADLAEPTHPHVGAPPHAPPTHGALRL

GAVMGAALLLSALGLSVWACMTCWRRRAWRAVKSRASGKGPTYIRVADSELYADWSSDSEGERDQVPWLA

PPERPDSPSTNGSGFKILSPTAPSVYPRSDGHQSRRQLTTFGSGRPDRRYSQASDSSVFW

>HSV1GPE_19

MDRGAVVGFLLGVCVVSCLAGTPKTSWRRVSVGEDVSLLPAPGPTGRGSTQKLLWAVEPLDGCGPLHPSW

VSLMPPKQVPETVVDAACMRAPVPLAMAYAPPAPSATGGLPTDFVWQERAAVVNRSLVIHGVRETDSGLY

TLSVGDIKDPARQVASVVLVVQPAPVPTPPPTPADYDEDDNDEGEDESLAGTPASGTPRLPPPPAPPRSW

PSAPEVSHVRGVTVRMETPEAILFSPGETFSTNVSIHAIAHDDQTYAMDVVWLRFDVPTSCAEMRIYESC

LYHPQLPECLSPADAPCAASTWTSRLAVRSYAGCSRTNPPPRCSAEAHMEPVPGLAWQAASVNLEFRDAS

PQHSGLYLCVVYVNDHIHAWGHITISTAAQYRNAVVEQPLPQRGADLAEPTHPHVGAPPHAPPTHGALRL

GAVMGAALLLSALGLSVWACMTCWRRRAWRAVKSRASGKGPTYIRVADSELYADWSSDSEGERDQVPWLA

PPERPDSPSTNGSGFEILSPTAPSVYPRSDGHQSRRQLTTFGSGRPDRRYSQASDSSVFW

>HSV1GPE_20

MDRGAVVGFLLGVCVVSCLAGTPKTSWRRVSVGEDVSLLPAPGPTGRGPTQKLLWAVEPLDGCGPLHPSW

VSLMPPKQVPETVVDAACMRAPVPLAMAYAPPAPSATGGLRTDFVWQERTAVVNRSLVIYGVRETDSGLY

TLSVGDIKDPARQVASVVLVVQPAPVPTPPPTPADYDEDDNDEGEDESLAGTPASGTPRLPPPPAPPRSW

PSAPEVSHVRGVTVRMETPEAILFSPGEAFSTNVSIHAIAHDDQTYAMDVVWLRFDVPTSCAEMRIYESC

LYHPQLPECLSPADAPCAASTWTSRLAVRSYAGCSRTNPPPRCSAEAHMEPVPGLAWQAASVNLEFRDAS

PQHSGLYLCVVYVNDHIHAWGHITISTAAQYRNAVVEQPLPQRGADLAEPTHPHVGAPPHAPPTHGALRL

GAVMGAALLLSVLGLSVWACMTCWRRRAWRAVKSRASGKGPTYIRVADSELYADWSSDSEGERDQVPWLA

PPERPDSPSTNGSGFEILSPTAPSVYPRSDGHQSRRQLTTFGSGRPDRRYSQASDSSVFW

>HSV2GPE_21

MARGAGLVFFVGVWVVSCLAAAPRTSWKRVTSGEDVVLLPAPAERTRAHKLLWAAEPLDACGPLRPSWVA

LWPPRRVLETVVDAACMRAPEPLAIAYSPPFPAGDEGLYSELAWRDRVAVVNESLVIYGALETDSGLYTL

SVVGLSDEARQVASVVLVVEPAPVPTPTPDDYDEEDDAGVTNARRSAFPPQPPPRRPPVAPPTHPRVIPE

VSHVRGVTVHMETLEAILFAPGETFGTNVSIHAIAHDDGPYAMDVVWMRFDVPSSCADMRIYEACLYHPQ

LPECLSPADAPCAVSSWAYRLAVRSYAGCSRTTPPPRCFAEARMEPVPGLAWLASTVNLEFQHASPQHAG

LYLCVVYVDDHIHAWGHMTISTAAQYRNAVVEQHLPQRQPEPVEPTRPHVRAPHPAPSARGPLRLGAVLG

AALLLAALGLSAWACMTCWRRRSWRAVKSRASATGPTYIRVADSELYADWSSDSEGERDGSLWQDPPERP

DSPSTNGSGFEILSPTAPSVYPHSEGRKSRRPLTTFGSGSPGRRHSQASYPSVLW

>HSV2GPE_22

MARGAGLVFFVGVWVVSCLAAAPRTSWKRVTSGEDVVLLPAPAERTRAHKLLWAAEPLDACGPLRPSWVA

LWPPRRVLETVVDAACMRAPEPLAIAYSPPFPAGDEGLYSELAWRDRVAVVNESLVIYGALETDSGLYTL

SVVGLSDEARQVASVVLVVEPAPVPTPTPDDYDEEDDAGVSERTPVSVPPPTPPRRPPVAPPTHPRVIPE

VSHVRGVTVHMETPEAILFAPGETFGTNVSIHAIAHDDGPYAMDVVWMRFDVPSSCAEMRIYEACLYHPQ

LPECLSPADAPCAVSSWAYRLAVRSYAGCSRTTPPPRCFAEARMEPVPGLAWLASTVNLEFQHASPQHAG

LYLCVVYVDDHIHAWGHMTISTAAQYRNAVVEQHLPQRQPEPVEPTRPHVRAPHPAPSARGPLRLGAVLG

AALLLAALGLSAWACMTCWRRRSWRAVKSRASATGPTYIRVADSELYADWSSDSEGERDGSLWQDPPERP

DSPSTNGSGFEILSPTAPSVYPHSEGRKSRRPLTTFGSGSPGRRHSQASYPSVLW

>HSV2GPE_23

MARGAGLVFFVGVWVVSCLAAAPRTSWKRVTSGEDVVLLPAPAERTRAHKLLWAAEPLDACGPLRPSWVA

LWPPRRVLETVVDAACMRAPEPLAIAYSPPFPAGDEGLYSELAWRDRVAVVNESLVIYGALETDSGLYTL

SVVGLSDEARQVASVVLVVEPAPVPTPTPDDYDEEDDAGVSERTPVSVPPPTPPRRPPVAPPTHPRVIPE

VSHVRGVTVHMETPEAILFAPGETFGTNVSIHAIAHDDGPYAMDVVWMRFDVPSSCAEMRIYEACLYHPQ

LPECLSPADAPCAVSSWAYRLAVRSYAGCFRTTPPPRCFAEARMEPVPGLAWLASTVNLEFQHASPQHAG

LYLCVVYVDDHIHAWGHMTISTAAQYRNAVVEQHLPQRQPEPVEPTRPHVRAPHPAPSARGPLRLGAVLG

AALLLAALGLSAWACMTCWRRRSWRAVKSRASATGPTYIRVADSELYADWSSDSEGERDGSLWQDPPERP

DSPSTNGSGFEILSPTAPSVYPHSEGRKSRRPLTTFGSGSPGRRHSQASYPSVLW

>HSV2GPE_24

MARGAGLVFFVGVWVVSCLAAAPRTSWKRVTSGEDVVLLPAPAGPEERTRAHKLLWAAEPLDACGPLRPS

WVALWPPRRVLETVVDAACMRAPEPLAIAYSPPFPAGDEGLYSELAWRDRVAVVNESLVIYGALETDSGL

YTLSVVGLSDEARQVASVVLVVEPAPVPTPTPDDYDEEDDAGVSERTPVSVPPPPPPRRPPVAPPTHPRV

IPEVSHVRGVTVHMETPEAILFAPGETFGTNVSIHAIAHDDGPYAMDVVWMRFDVPSSCAEMRIYEACLY

HPQLPECLSPADAPCAVSSWAYRLAVRSYAGCSRTTPPPRCFAEARMEPVPGLAWLASTVNLEFQHASPQ

HAGLYLCVVYVDDHIHAWGHMTISTAAQYRNAVVEQHLPQRQPEPVEPTRPHVRAPHPAPSARGPLRLGA

VLGAALLLAALGLSAWACMTCWRRRSWRAVKSRASATGPTYIRVADSELYADWSSDSEGERDGSLWQDPP

ERPDSPSTNGSGFEILSPTAPSVYPHSEGRKSRRPLTTFGSGSPGRRHSQASYSSVLW

>HSV2GPE_25

MARGAGLVFFVGVWVVSCLAAAPRTSWKRVTSGEDVVLLPAPAGPEERTRAHKLLWAAEPLDACGPLRPS

WVALWPPRRVLETVVDAACMRAPEPLAIAYSPPFPAGDEGLYSELAWRDRVAVVNESLVIYGALETDSGL

YTLSVVGLSDEARQVASVVLVVEPAPVPTPTPDDYDEEDDAGVSERTPVSVPPATPPRRPPVAPPTHPRV

IPEVSHVRGVTVHMETPEAILFAPGETFGTNVSIHAIAHDDGPYAMDVVWMRFDVPSSCAEMRIYEACLY

HPQLPECLSPADAPCAVSSWAYRLAVRSYAGCSRTTPPPRCFAEARMEPVPGLAWLASTVNLEFQHASPQ

HAGLYLCVVYVDDHIHAWGHMTISTAAQYRNAVVEQHLPQRQPEPVEPTRPHVRAPHPAPSARGPLRLGA

VLGAALLLAALGLSAWACMTCWRRRSWRAVKSRASATGPTYIRVADSELYADWSSDSEGERDGSLWQDPP

ERPDSPSTNGSGFEILSPTAPSVYPHSEGRKSRRPLTTFGSGSPGRRHSQASYSSVLW

>HSV2GPE_26

MARGAGLVFFVGVWVVSCLAAAPRTSWKRVTSGEDVVLLPAPAGPEERTRAHKLLWAAEPLDACGPLRPS

WVALWPPRRVLETVVDAACMRAPEPLAIAYSPPFPAGDEGLYSELAWRDRVAVVNESLVIYGALETDSGL

YTLSVVGLSDEARQVASVVLVVEPAPVPTPTPDDYDEEDDAGVSERTPVSVPPPTPPRRPPVAPPTHPRV

IPEVSHVRGVTVHMETPEAILFAPGETFGTNVSIHAIAHDDGPYAMDVVWMRFDVPSSCAEMRIYEACLY

HPQLPECLSPADAPCAVSSWAYRLAVRSYAGCSRTTPPPRCFAEARMEPVPGLAWLASTVNLEFQHASPQ

HAGLYLCVVYVDDHIHAWGHMTISTAAQYRNAVVEQHLPQRQPEPVEPTRPHVRAPHPAPSARGPLRLGA

VLGAALLLAALGLSAWACMTCWRRRSWRAVKSRASATGPTYIRVADSELYADWSSDSEGERDGSLWQDPP

ERPDSPSTNGSGFEILSPTAPSVYPHSEGRKSRRPLTTFGSGSPGRRHSQASYSSVLW

>HSV2GPE_27

MARGAGLVFFVGVWVVSCLAAAPRTSWKRVTSGEDVVLLPAPAGPEERTRAHKLLWAAEPLDACGPLRPS

WVALWPPRRVLETVVDAACMRAPEPLAIAYSPPFPAGDEGLYSELAWRDRVAVVNESLVIYGALETDSGL

YTLSVVGLSDEARQVASVVLVVEPAPVPTPTPDDYDEEDDAGVSERTPVSVPPPPPPRRPPVAPPTHPRV

IPEVSHVRGVTVHMETPEAILFAPGETFGTNVSIHAIAHDDGPYAMDVVWMRFDVPSSCAEMRIYEACLY

HPQLPECLSPADAPCAVSSWAYRLAVRSYAGCSRTTPPPRCFAEARMEPVPGLAWLASTVNLEFQHASPQ

HAGLYLCVVYVDDHIHAWGHMTISTAAQYRNAVVEQHLPQRQPEPVEPTRPHVRAPHPVPSARGPLRLGA

VLGAALLLAALGLSAWACMTCWRRRSWRAVKSRASATGPTYIRVADSELYADWSSDSEGERDGSLWQDPP

ERPDSPSTNGSGFEILSPTAPSVYPHSEGRKSRRPLTTFGSGSPGRRHSQASYSSVLW

>HSV2GPE_28

MARGAGLVFFVGVWVVSCLAAAPRTSWKRVTSGEDVVLLPAPAGPEERTRTHKLLWAAEPLDACGPLRPS

WVALWPPRRVLETVVDAACMRAPEPLAIAYSPPFPAGDEGLYSELAWRDRVAVVNESLVIYGALETDSGL

YTLSVVGLSDEARQVASVVLVVEPAPVPTPTPDDYDEEDDAGVSERTPVSVPPATPPRRPPVAPPTHPRV

IPEVSHVRGVTVHMETPEAILFAPGETFGTNVSIHAIAHDDGPYAMDVVWMRFDVPSSCAEMRIYEACLY

HPQLPECLSPADAPCAVSSWAYRLAVRSYAGCSRTTPPPRCFAEARMEPVPGLAWLASTVNLEFQHASPQ

HAGLYLCVVYVDDHIHAWGHMTISTAAQYRNAVVEQHLPQRQPEPVEPTRPHVRAPHPAPSARGPLRLGA

VLGAALLLAALGLSAWACMTCWRRRSWRAVKSRASATGPTYIRVADSELYADWSSDSEGERDGSLWQDPP

ERPDSPSTNGSGFEILSPTAPSIYPHSEGRKSRRPLTTFGSGSPGRRHSQASYSSVLW

>HSV2GPE_29

MARGAGLVFFVGVWVVSCLAAAPRTSWKRVTSGEDVVLLPAPAGPEERTRAHKLLWAAEPLDACGPLRPS

WVALWPPRRVLETVVDAACMRAPEPLAIAYSPPFPAGDEGLYSELAWRDRVAVVNESLVIYGALDTDSGL

YTLSVVGLSDEARQVASVVLVVEPAPVPTPTPDDYDEEDDAGVSERTPVSVPPPTPPRRPPVAPPTHPRV

IPEVSHVRGVTVHMETPEAILFAPGETFGTNVSIHAIAHDDGPYAMDVVWMRFDVPSSCAEMRIYEACLY

HPQLPECLSPADAPCAVSSWAYRLAVRSYAGCSRTTPPPRCFAEARMEPVPGLAWLASTVNLEFQHASPQ

HAGLYLCVVYVDDHIHAWGHMTISTAAQYRNAVVEQHLPQRQPEPVEPTRPHVRAPHPAPSARGPLRLGA

VLGAALLLAALGLSAWACMTCWRRRSWRAVKSRASATGPTYIRVADSELYADWSSDSEGERDGSLWQDPP

ERPDSPSTNGSGFEILSPTAPSVYPHSEGRKSRRPLTTFGSGSPGRRHSQASYSSVLW

>HSV2GPE_30 MARGAGLVFFVGVWVVSCLAAAPRTSWKRVTSGEDVVLLPAPAGPEERTRAHKLLWAAEPLDACGPLRPS

WVALWPPRRVLETVVDAACMRAPEPLAIAYSPPFPAGDEGLYSELAWRDRVAVVNESLVIYGARETDSGL

YTLSVVGLSDEARQVASVVLVVEPAPVPTPTPDDYDEEDDAGVSERTPVSVPPTTPPRRPPVAPPTHPRV

IPEVSHVRGVTVHMETPEAILFAPGETFGTNVSIHAIAHDDGPYAMDVVWMRFDVPSSCAEMRIYEACLY

HPQLPECLSPADAPCAVSSWAYRLAVRSYAGCSRTTPPPRCFAEARMEPVPGLAWLASTVNLEFQHASPQ

HAGLYLCVVYVDDHIHAWGHMTISTAAQYRNAVVEQHLPQRQPEPVEPTRPHVRAPHPAPSARGPLRLGA

VLGAALLLAALGLSAWACMTCWRRRSWRAVKSRASATGPTYIRVADSELYADWSSDSEGERDGSLWQDPP

ERPDSPSTNGSGFEILSPTAPSVYPHSEGRKSRRPLTTFGSGSPGRRHSQASYSSVLW

>HSV2GPE_31

MARGAGLVFFVGVWVVSCLGAAPRTSWKRVTSGEDVVLLPAPAGPEERTRAHKLLWAAEPLDACGPLRPS

WVALWPPRRVLETVVDAACMRAPEPLAIAYSPPFPAGDEGLYSELAWRDRVAVVNESLVIYGALETDSGL

YTLSVVGLSDEARQVASVVLVVEPAPVPTPTPDDYDEEDDAGVSERTPVSVPPPTPPRRPPVAPPTHPRV

IPEVSHVRGVTVHMETPEAILFAPGETFGTNVSIHAIAHDDGPYAMDVVWMRFDVPSSCAEMRIYEACLY

HPQLPECLSPADAPCAVSSWAYRLAVRSYAGCSRTTPPPRCFAEARMEPVPGLAWLASTVNLEFQHASPQ

HAGLYLCVVYVDDHIHAWGHMTISTAAQYRNAVVEQHLPQRQPEPVEPTRPHVRAPHPAPSARGPLRLGA

VLGAALLLAALGLSAWACMTCWRRRSWRAVKSRASATGPTYIRVADSELYADWSSDSEGERDGSLWQDPP

ERPDSPSTNGSGFEILSPTAPSVYPHSEGRKSRRPLTTFGSGSPGRRHSQASYSSVLW

>HSV2GPE_32 MARGAGLVFFVGVWVVSCLAAAPRTSWKRVTSGEDVVLLPAPVGPEERTRAHKLLWAAEPLDACGPLRPS

WVALWPPRRVLETVVDAACMRAPEPLAIAYSPPFPAGDEGLYSELAWRDRVAVVNESLVIYGALETDSGL

YTLSVVGLSDEARQVASVVLVVEPAPVPTPTPDDYDEEDDAGVSERTPVSVPPPTPPRRPPVAPPTHPRV

IPEVSHVRGVTVHMETPEAILFAPGETFGTNVSIHAIAHDDGPYAMDVVWMRFDVPSSCAEMRIYEACLY

HPQLPECLSPADAPCAVSSWAYRLAVRSYAGCSRTTPPPRCFAEARMEPVPGLAWLASTVNLEFQHASPQ

HAGLYLCVVYVDDHIHAWGHMTISTAAQYRNAVVEQHLPQRQPEPVEPTRPHVRAPHPAPSARGPLRLGA

VLGAALLLAALGLSAWACMTCWRRRSWRAVKSRASATGPTYIRVADSELYADWSSDSEGERDGSLWQDPP

ERPDSPSTNGSGFEILSPTAPSVYPHSEGRKSRRPLTTFGSGSPGRRHSQASYSSVLW

>HSV2GPE_33

MARGAGLVFFVGVWVVSCLAAAPRTSWKRVTSGEDVVLLPAPAGPEERTRAHKLLWAAEPLDACGPLRPS

WVALGPPRRVLETVVDAACMRAPEPLAIAYSPPFPAGDEGLYSELAWRDRVAVVNESLVIYGALETDSGL

YTLSVVGLSDEARQVASVVLVVEPAPVPTPTPDDYDEEDDAGVSERTPVSVPPTPPPRRPPVAPPTHPRV

IPEVSHVRGVTVHMETPEAILFAPGETFGTNVSIHAIAHDDGPYAMDVVWMRFDVPSSCAEMRIYEACLY

HPQLPECLSPADAPCAVSSWAYRLAVRSYAGCSRTTPPPRCFAEARMEPVQGLAWLASTVNLEFQHASPQ

HAGLYLCVVYVDDHIHAWGHMTISTAAQYRNAVVEQHLPQRQPEPVEPTRPHVRAPHPAPSARGPLRLGA

VLGAALLLAALGLSAWACMTCWRRRSWRAVKSRASATGPTYIRVADSELYADWSSDSEGERDGSLWQDPP

ERPDSPSTNGSGFEILSPTAPSVYPHSEGRKSRRPLTTFGSGSPGRRHSQASYSSVLW

>HSV2GPE_34 MARGAGLVFFVGVWVVSCLAAAPRTSWKRVTSGEDVVLLPAPAGPEERTRAHKLLWAAEPLDACGPLRPS

WVALWPPRRVLETVVDAACMRAPEPLAIAYSPPFPAGDEGLYSELAWRDRVAVVNESLVIYGALETDSGL

YTLSVVGLSDEARQVASVVLVVEPAPVPTPTPDDYDEEDDAGVSERTPVSVPPATPPRRPPVAPPTHPRV

IPEVSHVRGVTVHMETPEAILFAPGETFGTNVSIHAIAHDDGPYAMDVVWMRFDVPSSCAEMRIYEACLY

HPQLPECLSPADAPCAVSSWAYRLAVRSYAGCSRTTPPPRCFAEARMEPVPGLAWLASTVNLEFQHASPQ

HAGLYLCVVYVDDHIHAWGHMTISTAAQYRNAVVEQHLPQRQPEPVEPTRPHVRAPPPAPSARGPLRLGA

VLGAALLLAALGLSAWACMTCWRRRSWRAVKSRASATGPTYIRVADSELYADWSSDSEGERDGSLWQDPP

ERPDSPSTNGSGFEILSPTAPSVYPHSEGRKSRRPLTTFGSGSPGRRHSQASYSSVLW

>HSV2GPE_35

MARGAGLVFFVGVWVVSCLAAAPRTSWKRVTSGEDVVLLPAPAGPEERTRAHKLLWAAEPLDACGPLRPS

WVALWPPRRVLETVVDAACMRAPEPLAIAYSPPFPAGDEGLYSELAWRDRVAVVNESLVIYGALETDSGL

YTLSVVGLSDEARQVASVVLVVEPAPVPTPTPDDYDEEDDAGVSERTLVSVPPPTPPRRPPVAPPTHPRV

IPEVSHVRGVTVHMETPEAILFAPGETFGTNVSIHAIAHDDGPYAMDVVWMRFDVPSSCAEMRIYEACLY

HPQLPECLSPADAPCAVSSWAYRLAVRSYAGCSRTTPPPRCFAEARMEPVPGLAWLASTVNLEFQHASPQ

HAGLYLCVVYVDDHIHAWGHMTISTAAQYRNAVVEQHLPQRQPEPVEPTRPHVRAPPPAPSARGPLRLGA

VLGAALLLAALGLSAWACMTCWRRRSWRAVKSRASATGPTYIRVADSELYADWSSDSEGERDGSLWQDPP

ERPDSPSTNGSGFEILSPTAPSVYPHSEGRKSRRPLTTFGSGSPGRRHSQASYSSVLW

> HSV2GPE_36

MARGAGLVFFVGVWVVSCLAAAPRTSWKRVTSGEDVVLLPAPAGPEERTRAHKLLWAAEPLDACGPLRPS

WVALWPPRRVLETVVDAACMRAPEPLAIAYSPPFPAGDEGLYSELAWRDRVAVVNESLVIYGALETDSGL

YTLSVVGLSDEARQVASVVLVVEPAPVPTPTPDDYDEEDDAGVSERTPVSVPPPTPPRRPPVAPPTHPRV

IPEVSHVRGVTVHMETPEAILFAPGETFGTNVSIHAIAHDDGPYAMDVVWMRFDVPSSCAEMRIYEACLY

HPQLPECLSPADAPCAVSSWAYRLAVRSYAGCSRTTPPPRCFAEARMEPVPGLAWLASTVNLEFQHASPQ

HAGLYLCVVYVDDHIHAWGHMTISTAAQYRNAVVEQHLPQRQPEPVEPTRPHVRAPPPAPSARGPLRLGA

VLGAALLLAALGLSAWACMTCWRRRSWRAVKSRASATGPTYIRVADSELYADWSSDSEGERDGSLWQDPP

ERPDSPSTNGSGFEILSPTAPSVYPHSEGRKSRRPLTTFGSGSPGRRHSQASYSSVLW

>HSV2GPE_37

MARGAGLVFFVGVWVVSCLAAAPRTSWKRVTSGEDVVLLPAPAGPEERTRAHKLLWAAEPLDACGPLRPS

WVALWPPRRVLETVVDAACMRAPEPLAIAYSPPFPAGDEGLYSELAWRDRVAVVNESLVIYGARETDSGL

YTLSVVGLSDEARQVASVVLVVEPAPVPTPTPDDYDEEDDAGVSERTPVSVPPPTPPRRPPVAPQTHPRV

IPEVSHVRGVTVHMETPEAILFAPGETFGTNVSIHAIAHDDGPYAMDVVWMRFDVPSSCAEMRIYEACLY

HPQLPECLSPADAPCAVSSWAYRLAVRSYAGCSRTTPPPRCFAEARMEPVPGLAWLASTVNLEFQHASPQ

HAGLYLCVVYVDDHIHAWGHMTISTAAQYRNAVVEQHLPQRQPEPVEPTRPHVRAPHPAPSARGPLRLGA

VLGAALLLAALGLSAWACMTCWRRRSWRAVKSRASATGPTYIRVADSELYADWSSDSEGERDGSLWQDPP

ERPDSPSTNGSGFEILSPTAPSVYPHSEGRKSRRPLTTFGSGSPGRRHSQASYSSVLW

>HSV2GPE_38

MARGAGLVFFVGVWVVSCLAAAPRTSWKRVTSGEDVVLLPAPTGPEERTRAHKLLWAAEPLDACGPLRPS

WVALWPPRRVLETVVDAACMRAPEPLAIAYSPPFPAGDEGLYSELAWRDRVAVVNESLVIYGARETDSGL

YTLSVVGLSDEARQVASVVLVVEPAPVPTPTPDDYDEEDDAGVSERTPVSVPPPTPPRRPPVAPQTHPRV

IPEVSHVRGVTVHMETPEAILFAPGETFGTNVSIHAIAHDDGPYAMDVVWMRFDVPSSCAEMRIYEACLY

HPQLPECLSPADAPCAVSSWAYRLAVRSYAGCSRTTPPPRCFAEARMEPVPGLAWLASTVNLEFQHASPQ

HAGLYLCVVYVDDHIHAWGHMTISTAAQYRNAVVEQHLPQRQPEPVEPTRPHVRAPHPAPSARGPLRLGA

VLGAALLLAALGLSAWACMTCWRRRSWRAVKSRASATGPTYIRVADSELYADWSSDSEGERDGSLWQDPP

ERPDSPSTNGSGFEILSPTAPSVYPHSEGRKSRRPLTTFGSGSPGRRHSQASYSSVLW

>HSV2GPE_39

MARGAGLVFFVGVWVVSCLAAAPRTSWKRVTSGEDVVLLPAPAGPEERTRAHKLLWAAEPLDACGPLRPS

WVALWPPRRVLETVVDAACMRAPEPLAIAYSPPFPAGDEGLYSELAWRDRVAVVNESLVIYGALETDSGL

YTLSVVGLSDEARQVASVVLVVEPAPVPTPTPDDYDEEDDAGVSERTPVSVPPPTPPRRPPVAPPTHPRV

IPEVSHVRGVTVHMETPEAILFAPGETFGTNVSIHAIAHDDGPYAMDVVWMRFDVPSSCAEMRIYEACLY

HPQLPECLSPADAPCAVSSWAYRLAVRSYAGCSRTTPPPRCFAEARMEPVPGLAWLASTVNLEFQHASPQ

HAGLYLCVVYVDDHIHAWGHMTISTAAQYRNAVVEQHLPQRQPEPVEPTRPHVRAPHPAPSARGPLRLGA

VLGAALLLAALGLSAWACMTCWRRRSWRAVKSRASATGPTYIRVADSELYADWSSDSEGERDGSLWQDPP

ERPDSPSTNGSGFEILSPTAPSVYPHSEGRKSRRPLTTFGSGSPGRRHSQASY

>HSV2GPE_40 MARGAGLVFFVGVWVVSCLAAAPRTSWKRVTSGEDVVLLPAPAGPEERTRAHKLLWAAEPLDACGPLRPS

WVALWPPRRVLETVVDAACMRAPEPLAIAYSPPFPAGDEGLYSELAWRDRVAVVNESLVIYGALETDSGL

YTLSVVGLSDEARQVASVVLVVEPAPVPTPTPDDYDEEDDAGVSERTPVSVPPPTPPRRPPVAPPTHPRV

IPEVSHVRGVTVHMETPEAILFAPGETFGTNVSIHAIAHDDGPYAMDVVWMRFDVPSSCAEMRIYEACLY

HPQLPECLSPADAPCAVSSWAYRLAVRSYAGCSRTTPPPRCFAEARMEPVPGLAWLASTVNLEFQHASPQ

HAGLYLCVVYVDDHIHAWGHMTISTAAQYRNAVVEQHLPQRQPEPVEPTRPHVRAPPPAPSARGPLRLRA

VLGAALLLAALGLSAWACMTCWRRRSWRAVKSRASATGPTYIRVADSELYADWSSDSEGERDGSLWQDPP

ERPDSPSTNGSGFEILSPTAPSVYPHSEGRKSRRPLTTFGSGSPGRRHSQASYSSVLW
