## Supplementary file 5 for "Contriving a chimeric polyvalent vaccine to prevent infections caused by Herpes Simplex Virus (Type-1 and Type-2): an exploratory immunoinformatic approach"

**ENVELOPE GLYCOPRPOTEIN G**

>HSV1GPG_1

MSQGAMRAVVPIIPFLLVLVGVSGVPTNVSSTTQPQLQTTGRPSHEAPNMTQTGTTDSPTAISLTTPDHT

PPMPSIGLEEEEEEEGAGDGEHLEGGDGTRDTLPQSPGPAFPLAEDVEKDKPNRPVVPSPDPNNSPARPE

TSRPKTPPTIIGPLATRPTTRLTSKGRPLVPTPQHTPLFSFLTASPALDTLFVVSTVIHTLSFLCIGAMA

THLCGGWSRRGRRTHPSVRYVCLPSERG

>HSV1GPG_2

MSQGAMRAVVPIIPFLLVLVGVSGVPTNVSSTTQPQLQTTGRPSHEAPNMTQTGTTDSPTAISLTTPDHT

PPMPSIGLEEEEEEEGAGDGEHLEGGDGTRDTLPQSPGPAFPLAEDVEKDKPNRPVVPSPDPNNSPARPE

TNRPKTPPTIIGPLATRPTTRLTSKGRPLVPTPQHTPLFSFLTASPALDTLFVVSTVIHTLSFLCIGAMA

THLCGGWSRRGRRTHPSVRYVCLPSERG

>HSV1GPG_3

MSQGAMRAVVPIIPFLLVLVGVSGVPTNVSSTTQPQLQTTGRPSHEAPNMTQTGTTDSPTAISLTTPDHT

PPMPSIGLEEEEEEEGAGDGEHLEGGDGTRDTLPQSPGPAFPLAEDVEKDKPDRPVVPSPDPNNSPARPE

TSRPKTPPTIIGPLATRPTTRLTSKGRPLVPTPQHTPLFSFLTASPALDTLFVVSTVIHTLSFLCIGAMA

THLCGGWSRRGRRTHPSVRYVCLPSERG

>HSV1GPG_4 MSQGAMRAVVPIIPFLLVLVGVSGVPTNVSSTTQPQLQTTGRTSHEAPNMTQTGTTDSPTAISLTTPDHT

PPMPSIGLEEEEEEEGAGDGEHLEGGDGTRDTLPQSPGPAFPLAEDVEKDKPNRPVVPSPDPNNSPARPE

TSRPKTPPTIIGPLATRPTTRLTSKGRPLVPTPQHTPLFSFLTASPALDTLFVVSTVIHTLSFLCIGAMA

THLCGGWSRRGRRTHPSVRYVCLPSERG

>HSV1GPG_5

MSPGAMRAVVPIIPFLLVLVGVSGVPTNVSSTTQPQLQTTGRPSHEAPNMTQTGTTDSPTAISLTTPDHT

PPMPSIGLEEEEEEEGAGDGEHLEGGDGTRDTLPQSPGPAFPLAEDVEKDKPNRPVVPSPDPNNSPARPE

TSRPKTPPTIIGPLATRPTTRLTSKGRPLVPTPQHTPLFSFLTASPALDTLFVVSTVIHTLSFLCIGAMA

THLCGGWSRRGRRTHPSVRYVCLPSERG

>HSV1GPG_6

MSPGAMRAVVPIIPFLLVLVGVSGVPTNVSSTTQPQLQTTGRPSHEAPNMTQTGTTDSPTAISLTTPDHT

PPMPSIGLEEEEEEEGAGDGEHLEGGDGTRDTLPQSPGPAFPLAEDVEKDNPNRPVVPSPDPNNSPARPE

TSRPKTPPTIIGPLATRPTTRLTSKGRPLVPTPQHTPLFSFLTASPALDTLFVVSTVIHTLSFLCIGAMA

THLCGGWSRRGRRTHPSVRYVCLPSERG

>HSV2GPG_7

MHAIAPRLLLLFVLSGLPGTRGGSGVPGPINPPNSDVVFPGGSPVAQYCYAYPRLDDPGPLGSADAGRQD

LPRRVVRHEPLGRSFLTGGLVLLAPPVRGFGAPNATYAARVTYYRLTRACRQPILLRQYGGCRGGEPPSP

KTCGSYTYTYQGGGPPTRYALVNASLLVPIWDRAAETFEYQIELGGELHVGLLWVEVGGEGPGPTAPPQA

ARAEGGPCVPPVPAGRPWRSVPPVWYSAPNPGFRGLRFRERCLPPQTPAAPSDLPRVAFAPQSLLVGITG

RTFIRMARPTEDVGVLPPHWAPGALDDGPYAPFPPRPRFRRALRTDPEGVDPDVRAPRTGRRLMALTEDT

SSDSPTSAPEKTPLPVSATAMAPSVDPSAEPTAPATTTPPDEMATQAATVAVTPEETAVASPPATASVES

SPLPAAAAATPGAGHTNTSSASAAKTPPTTPAPTTPPPTSTHATPRPTTPGPQTTPPGPATPGPVGASAA

PTADSPLTASPPATAPGPSAANVSVAATTATPGTRGTARTPPTDPKTHPHGPADAPPGSPAPPPPEHRGG

PEEFEGAGDGEPPEDDDSATGLAFRTPNPNKPPPARPGPIRPTLPPGILGPLAPNTPRPPAQAPAKDMPS

GPTPQHIPLFWFLTASPALDILFIISTTIHTAAFVCLVALAAQLWRGRAGRRRYAHPSVRYVCLPPERD

>HSV2GPG_8

MHAIAPRLLLLFVLSGLPGTRGGSGVPGPINPPNSDVVFPGGSPVAQYCYAYPRLDDPGPLGSADAGRQD

LPRRVVRHEPLGRSFLTGGLVLLAPPVRGFGAPNATYAARVTYYRLTRACRQPILLRQYGGCRGGEPPSP

KTCGSYTYTYQGGGPPTRYALVNASLLVPIWDRAAETFEYQIELGGELHVGLLWVEVGGEGPGPTAPPQA

ARAEGGPCVPPVPAGRPWRSVPPVWYSAPNPGFRGLRFRERCLPPQTPAAPSDLPRVAFAPQSLLVGITG

RTFIRMARPTEDVGVLPPHWAPGALDDGPYAPFPPRPRFRRALRTDPEGVDPDVRAPRTGRRLMALTEDA

SSDSPTSAPEKTPLPVSATAMAPSVDPSAEPTAPATTTPPDEMATQAATVAVTPEETAVASPPATASVES

SPLPAAAAATPGAGHTNTSSASAAKTPPTTPAPTTPPPTSTHATPRPTTPGPQTTPPGPATPGPVGASAA

PTADSPLTASPPATAPGPSAANVSVAATTATPGTRGTARTPPTDPKTHPHGPADAPPGSPAPPPPEHRGG

PEEFEGAGDGEPPEDDDSATGLAFRTPNPNKPPPARPGPIRPTLPPGILGPLAPNTPRPPAQAPAKDMPS

GPTPQHIPLFWFLTASPALDILFIISTTIHTAAFVCLVALAAQLWRGRAGRRRYAHPSVRYVCLPPERD

>HSV2GPG_9

MHAIAPRLLLLFVLSGLPGTRGGSGVPGPINPPNSDVVFPGGSPVAQYCYAYPRLDDPGPLGSADAGRQD

LPRRVVRHEPLGRSFLTGGLVLLAPPVRGFGAPNATYAARVTYYRLTRACRQPILLRQYGGCRGGEPPSP

KTCGSYTYTYQGGGPPTRYALVNASLLVPIWDRAAETFEYQIELGGELHVGLLWVEVGGEGPGPTAPPQA

ARAEGGPCVPPVPAGRPWRSVPPVWYSAPNPGFRGLRFRERCLPPQTPAAPSDLPRVAFAPQSLLVGITG

RTFIRMARPTEDVGVLPPHWAPGALDDGPYAPFPPRPRFRRALRTDPEGVDPDVRAPRTGRRLMALTEDA

SSDSPTSAPEKTPLPVSATAMAPSVDPSAEPTAPATTTPPDEMAAQAATVAVTPEETAVASPPATASVES

SPLPAAAAATPGAGHTNTSSASAAKTPPTTPAPTTPPPTSTHATPRPTTPGPQTTPPGPATPGPVGASAA

PTADSPLTASPPATAPGPSAANVSVAATTATPGTRGTARTPPTDPKTHPHGPADAPPGSPAPPPPEHRGG

PEEFEGAGDGEPPEDDDSATGLAFRTPNPNKPPPARPGPIRPTLPPGILGPLAPNTPRPPAQAPAKDMPS

GPTPQHIPLFWFLTASPALDILFIISTTIHTAAFVCLVALAAQLWRGRAGRRRYAHPSVRYVCLPPERD

>HSV2GPG_10

MHAIAPRLLLLFVLSGLPGTRGGSGVPGPINPPNSDVVFPGGSPVAQYCYAYPRLDDPGPLGSADAGRQD

LPRRVVRHEPLGRSFLTGGLVLLAPPVRGFGAPNATYAARVTYYRLTRACRQPILLRQYGGCRGGEPPSP

KTCGSYTYTYQGGGPPTRYALVNASLLVPIWDRAAETFEYQIELGGELHVGLLWVEVGGEGPGPTAPPQA

ARAEGGPCVPPVPAGRPWRSVPPVWYSAPNPGFRGLRFRERCLPPQTPAAPSDLPRVAFAPQSLLVGITG

RTFIRMARPTEDVGVLPPHWAPGALDDGPYAPFPPRPRFRRALRTDPEGVDPDVRAPRTGRRLMALTEDA

SSDSPTSAPEKTPLPVSATAMAPSVDPSAEPTAPATTTPPDEMATQAATVAVTPEETAVASPPATASVES

SPLPAAAATPGAGHTNTSSASAAKTPPTTPAPTTPPPTSTHATPRPTTPGPQTTPPGPATPGPVGASAAP

TADSPLTASPPATAPGPSAANVSVAATTATPGTRGTARTPPTDPKTHPHGPADAPPGSPAPPPPEHRGGP

EEFEGAGDGEPPEDDDSATGLAFRTPNPNKPPPARPGPIRPTLPPGILGPLAPNTPRPPAQAPAKDMPSG

PTPQHIPLFWFLTASPALDILFIISTTIHTAAFVCLVALAAQLWRGRAGRRRYAHPSVRYVCLPPERD

>HSV2GPG_11 MHAIAPRLLLLFVLSGLPGTRGGSGVPGPINPPNNDVVFPGGSPVAQYCYAYPRLDDPGPLGSADAGRQD

LPRRVVRHEPLGRSFLTGGLVLLAPPVRGFGAPNATYAARVTYYRLTRACRQPILLRQYGGCRGGEPPSP

KTCGSYTYTYQGGGPPTRYALVNASLLVPIWDRAAETFEYQIELGGELHVGLLWVEVGGEGPGPTAPPQA

ARAEGGPCVPPVPAGRPWRSVPPVWYSAPNPGFRGLRFRERCLPPQTPAAPSDLPRVAFAPQSLLVGITG

RTFIRMARPTEDVGVLPPHWAPGALDDGPYAPFPPRPRFRRALRTDPEGVDPDVRAPRTGRRLMALTEDA

SSDSPTSAPEKTPLPVSATAMAPSVDPSAEPTAPATTTPPDEMATQAATVAVTPEETAVASPPATASVES

SPLPAAAATPGAGHTNTSSASAAKTPPTTPAPTTPPPTSTHATPRPTTPGPQTTPPGPATPGPVGASAAP

TADSPLTASPPATAPGPSAANVSVAATTATPGTRGTARTPPTDPKTHPHGPADAPPGSPAPPPPEHRGGP

EEFEGAGDGEPPEDDDSATGLAFRTPNPNKPPPARPGPIRPTLPPGILGPLAPNTPRPPAQAPAKDMPSG

PTPQHIPLFWFLTASPALDILFIISTTIHTAAFVCLVALAAQLWRGRAGRRRYAHPSVRYVCLPPERD

>HSV2GPG_12

MHAIAPRLLLLFVLSGLPGTRGGSGVPGPINPPNNDVVFPGGSPVAQYCYAYPRLDDPGPLGSADAGRQD

LPRRVVRHEPLGRSFLTGGLVLLAPPVRGFGAPNATYAARVTYYRLTRACRQPILLRQYGGCRGGEPPSP

KTCGSYTYTYQGGGPPTRYALVNASLLVPIWDRAAETFEYQIELGGELHVGLLWVEVGGEGPGPTAPPQA

ARAEGGPCVPPVPAGRPWRSVPPVWYSAPNPGFRGLRFRERCLPPQTPAAPSDLPRVAFAPQSLLVGITG

RTFIRMARPTEDVGVLPPHWAPGALDDGPYAPFPPRPRFRRALRTDPEGVDPDVRAPRTGRRLMALTENA

SSDSPTSAPEKTPLPVSATAMAPSVDPSAEPTAPATTTPPDEMATQAATVAVTPEETAVASPPATASVES

SPLPAAAATPGAGHTNTSSASAAKTPPTTPAPTTPPPTSTHATPRPTTPGPQTTPPGPATPGPVGASAAP

TADSPLTASPPATAPGPSAANVSVAATTATPGTRGTARTPPTDPKTHPHGPADAPPGSPAPPPPEHRGGP

EEFEGAGDGEPPEDDDSATGLAFRTPNPNKPPPARPGPIRPTLPPGILGPLAPNTPRPPAQAPAKDMPSG

PTPQHIPLFWFLTASPALDILFIISTTIHTAAFVCLVALAAQLWRGRAGRRRYAHPSVRYVCLPPERD

>HSV2GPG_13 MHAIAPRLLLLFVLSGLPGTRGGSGVPGPINPPNNDVVFPGGSPVAQYCYAYPRLDDPGPLGSADAGRQD

LPRRVVRHEPLGRSFLTGGLVLLAPPVRGFGAPNATYAARVTYYRLTRACRQPILLRQYGGCRGGEPPSP

KTCGSYTYTYQGGGPPTRYALVNASLLVPIWDRAAETFEYQIELGGELHVGLLWVEVGGEGPGPTAPPQA

ARAEGGPCVPPVPAGRPWRSVPPVWYSAPNPGFRGLRFRERCLPPQTPAAPSDLPRVAFAPQSLLVGITG

RTFIRMARPTEDVGVLPPHWAPGALDDGPYAPFPPRPRFRRALRTDPEGVDPDVRAPRTGRRLMALTEDA

SSDSPTSAPEKTPLPVSATAMAPSVDPSAEPTAPATTTPPDEMATQAATVAVTPDETAVASPPATASVES

SPLPAAAATPGAGHTNTSSASAAKTPPTTPAPTTPPPTSTHATPRPTTPGPQTTPPGPATPGPVGASAAP

TADSPLTASPPATAPGPSAANVSVAATTATPGTRGTARTPPTDPKTHPHGPADAPPGSPAPPPPEHRGGP

EEFEGAGDGEPPEDDDSATGLAFRTPNPNKPPPARPGPIRPTLPPGILGPLAPNTPRPPAQAPAKDMPSG

PTPQHIPLFWFLTASPALDILFIISTTIHTAAFVCLVALAAQLWRGRAGRRRYAHPSVRYVCLPPERD

>HSV2GPG_14

MHAIAPRLLLLFVLSGLPGTRGGSGVPGPINPPNSDVVFPGGAPVAQYCYAYPRLDDPGPLGSADAGRQD

LPRRVVRHEPLGRSFLTGGLVLLAPPVRGFGAPNATYAARVTYYRLTRACRQPILLRQYGGCRGGEPPSP

KTCGSYTYTYQGGGPPTRYALVNASLLVPIWDRAAETFEYQIELGGELHVGLLWVEVGGEGPGPTAPPQA

ARAEGGPCVPPVPAGRPWRSVPPVWYSAPNPGFRGLRFRERCLPPQTPAAPSDLPRVAFAPQSLLVGITG

RTFIRMARPTEDVGVLPPHWAPGALDDGPYAPFPPRPRFRRALRTDPEGVDPDVRAPRTGRRLMALTEDA

SSDSPTSAPEKTPLPVSATAMAPSVDPSAEPTAPATTTPPDEMATQAATVAVTPDETAVASPPATASVES

SPLPAAAATPGAGHTNTSSASAAKTPPTTPAPTTPPPTSTHATPRPTTPGPQTTPPGPATPGPVGASAAP

TADSPLTASPPATAPGPSAANVSVAATTATPGTRGTARTPPTDPKTHPHGPADAPPGSPAPPPPEHRGGP

EEFEGAGDGEPPEDDDSATGLAFRTPNPNKPPPARPGPIRPTLPPGILGPLAPNTPRPPAQAPAKDMPSG

PTPQHIPLFWFLTASPALDILFIISTTIHTAAFVCLVALAAQLWRGRAGRRRYAHPSVRYVCLPPERD

>HSV2GPG_15

MHAIAPRLLLLFVLSGLPGTRGGSGVPGPINPPNNDVVFPGGSPVAQYCYAYPRLDDPGPLGSADAGRQD

LPRRVVRHEPLGRSFLTGGLVLLAPPVRGFGAPNATYVARVTYYRLTRACRQPILLRQYGGCRGGEPPSP

KTCGSYTYTYQGGGPPTRYALVNASLLVPIWDRAAETFEYQIELGGELHVGLLWVEVGGEGPGPTAPPQA

ARAEGGPCVPPVPAGRPWRSVPPVWYSAPNPGFRGLRFRERCLPPQTPAAPSDLPRVAFAPQSLLVGITG

RTFIRMARPTEDVGVLPPHWAPGALDDGPYAPFPPRPRFRRALRTDPEGVDPDVRAPRTGRRLMALTENA

SSDSPTSAPEKTPLPVSATAMAPSVDPSAEPTAPATTTPPDEMATQAATVAVTPEETAVASPPATASVES

SPLPAAAATPGAGHTNTSSASAAKTPPTTPAPTTPPPTSTHATPRPTTPGPQTTPPGPATPGPVGASAAP

TADSPLTASPPATAPGPSAANVSVAATTATPGTRGTARTPPTDPKTHPHGPADAPPGSPAPPPPEHRGGP

EEFEGAGDGEPPEDDDSATGLAFRTPNPNKPPPARPGPIRPTLPPGILGPLAPNTPRPPAQAPAKDMPSG

PTPQHIPLFWFLTASPALDILFIISTTIHTAAFVCLVALAAQLWRGRAGRRRYAHPSVRYVCLPPERD

>HSV2GPG_16 MHAIAPRLLLLFVLSGLPGTRGGSGVPGPINPPNNDVVFPGGSPVAQYCYAYPRLDDPGPLGSADAGRQD

LPRRVVRHEPLGRSFLTGGLVLLAPPVRGFGAPNATYAARVTYYRLTRACRQPILLRQYGGCRGGEPPSP

KTCGSYTYTYQGGGPPTRYALVNASLLVPIWDRAAETFEYQIELGGELHVGLLWVEVGGEGPGPTAPPQA

ARAEGGPCVPPVPAGRPWRSVPPVWYSAPNPGFRGLRFRERCLPPQTPAAPSDLPRVAFAPQSLLVGITG

RTFIRMARPTEDVGVLPPHWAPGALDDGPYAPFPPRPRFRRALRTDPKGVDPDVRAPRTGRRLMALTEDA

SSDSPTSAPEKTPLPVSATAMAPSVDPSAEPTAPATTTPPDEMATQAATVAVTPEETAVASPPATASVES

SPLPAAAATPGAGHTNTSSASAAKTPPTTPAPTTPPPTSTHATPRPTTPGPQTTPPGPATPGPVGASAAP

TADSPLTASPPATAPGPSAANVSVAATTATPGTRGTARTPPTDPKTHPHGPADAPPGSPAPPPPEHRGGP

EEFEGAGDGEPPEDDDSATGLAFRTPNPNKPPPARPGPIRPTLPPGILGPLAPNTPRPPAQAPAKDMPSG

PTPQHIPLFWFLTASPALDILFIISTTIHTAAFVCLVALAAQLWRGRAGRRRYAHPSVRYVCLPPERD

>HSV2GPG_17

MHAIAPRLLLLFVLSGLPGTRGGSGVPGPINPPNNDVVFPGGSPVAQYCYAYPRLDDPGPLGSADAGRQD

LPRRVVRHEPLGRSFLTGGLVLLAPPVRGFGAPNATYAARVTYYRLTRACRQPILLRQYGGCRGGEPPSP

KTCGSYTYTYQGGGPPTRYALVNASLLVPIWDRAAETFEYQIELGGELHVGLLWVEVGGEGPGPTAPPQA

ARAEGGPCVPPVPAGRPWRSVPPVWYSAPNPGFRGLRFRERCLPPQTPAAPSDLPRVAFAPQSLLVGITG

RTFIRMARPTEDVGVLPPHWAPGALDDGPYAPFPPRPRFRRALRTDPEGVDPDVRAPRTGRRLMALTEDA

SSDSPTSAPEKTPLPVSATAMAPSVDPSAEPTAPATTTPPDEMATQAATVAVTPEETAVASPPATASVES

SPLPAAAATPGAGHTNTSSASAAKTPPTTPAPTTPPPTSTHATPRPTTPGPQTTPPGPATPGPVGASAAP

TADSPLTASPPATAPGPSAANVSVAATTATPGTRGTARTPPTDPKTHPHGPADAPPGSPAPPPPEHRGGP

EEFEGAGDGEPPEDDDSATGLAFRTPNPNKPPPARPGPIRPTLPPGILGPLAPNTPHPPAQAPAKDMPSG

PTPQHIPLFWFLTASPALDILFIISTTIHTAAFVCLVALAAQLWRGRAGRRRYAHPSVRYVCLPPERD

>HSV2GPG_18

MHAIAPRLLLLFVLSGLPGTRGGSGVPGPINPPNNDVVFPGGSPVAQYCYAYPRLDDPGPLGSADAGRQD

LPRRVVRHEPLGRSFLTGGLVLLAPPVRGFGAPNATYAARVTYYRLTRACRQPILLRQYGGCRGGEPPSP

KTCGSYTYTYQGGGPPTRYALVNASLLVPIWDRAAETFEYQIELGGELHVGLLWVEVGGEGPGPTAPPQA

ARAEGGPCVPPVPAGRPWRSVPPVWYSAPNPGFRGLRFRERCLPPQTPAAPSDLPRVAFAPQSLLVGITG

RTFIRMARPTEDVGVLPPHWAPGALDDGPYAPFPPRPRFRRALRTDPKGVDPDVRAPRTGRRLMALTEDA

SSDSPTSAPEKTPLPVSATAMAPSVDPSAEPTAPATTTPPDEMATQAATVAVTPEETAVASPPATASVES

SPLPAAAATPGAGHTNTSSASAAKTPPTTPAPTTPPPTSTHATPRPTTPGPQTTPPGPATPGPVGASAAP

TADSPLTASPPATAPGPSAANVSVAATTATPGTRGTARTPPTDPKTHPHGPADAPPGSPAPPPPEHRGGP

EEFEGAGDGEPPEDDDSATGLAFRTPNPNKPPPARPGPIRPTLPPGILGPLAPNTPRPPTQAPAKDMPSG

PTPQHIPLFWFLTASPALDILFIISTTIHTAAFVCLVALAAQLWRGRAGRRRYAHPSVRYVCLPPERD

>HSV2GPG_19

MHAIAPRLLLLFVLSGLPGTRGGSGVPGPINPPNNDVVFPGGSPVAQYCYAYPRLDDPGPLGSADAGRQD

LPRRVVRHEPLGRSFLTGGLVLLAPPVRGFGAPNATYAARVTYYRLTRACRQPILLRQYGGCRGGEPPSP

KTCGSYTYTYQGGGPPTRYALVNASLLVPIWDRAAETFEYQIELGGELHVGLLWVEVGGEGPGPTAPPQA

ARAEGGPCVPPVPAGRPWRSVPPVWYSAPNPGFRGLRFRERCLPPQTPAAPSDLPRVAFAPQSLLVGITG

RTFIRMARPTEDVGVLPPHWAPGALDDGPYAPFPPRPRFRRALRTDPEGVDPDVRAPRTGRRLMALTEDA

SSDSPTSAPEKTPLPVSATAMAPSVDPSAEPTAPATTTPPDEMATQAATVAVTPEETAVASPPATASVES

SPPPAAAATPGAGHTNTSSASAAKTPPTTPAPTTPPPTSTHATPRPTTPGPQTTPPGPATPGPVGASAAP

TADSPLTASPPATAPGPSAANVSVAATTATPGTRGTARTPPTDPKTHPHGPADAPPGSPAPPPPEHRGGP

EEFEGAGDGEPPEDDDSATGLAFRTPNPNKPPPARPGPIRPTLPPGILGPLAPNTPRPPAQAPAKDMPSG

PTPQHIPLFWFLTASPALDILFIISTTIHTAAFVCLVALAAQLWRGRAGRRRYAHPSVRYVCLPPERD

>HSV2GPG_20

MHAIAPRLLLLFVLSGLPGTRGGSGVPGPINPPNNDVVFPGGSPVAQYCYAYPRLDDPGPLGSADAGRQD

LPRRVVRHEPLGRSFLTGGLVLLAPPVRGFGAPNATYAARVTYYRLTRACRQPILLRQYGGCRGGEPPSP

KTCGSYTYTYQGGGPPTRYALVNASLLVPIWDRAAETFEYQIELDGELHVGLLWVEVGGEGPGPTAPPQA

ARAEGGPCVPPVPAGRPWRSVPPVWYSAPNPGFRGLRFRERCLPPQTPAAPSDLPRVAFAPQSLLVGITG

RTFIRMARPTEDVGVLPPHWAPGALDDGPYAPFPPRPRFRRALRTDPEGVDPDVRAPRTGRRLMALTEDA

SSDSPTSAPEKTPLPVSATAMAPSVDPSAEPTAPATTTPPDEMATQAATVAVTPEETAVASPPATASVES

SPLPAAAATPGAGHTNTSSASAAKTPPTTPAPTTPPPTSTHATPRPTTPGPQTTPPGPATPGPVGASAAP

TADSPLTASPPATAPGPSAANVSVAATTATPGTRGTARTPPTDPKTHPHGPADAPPGSPAPPPPEHRGGP

EEFEGAGDGEPPEDDDSATGLAFRTPNPNKPPPARPGPIRPTLPPGILGPLAPNTPRPPAQAPAKDMPSG

PTPQHIPLFWFLTASPALDILFIISTTIHTAAFVCLVALAAQLWRGRAGRRRYAHPSVRYVCLPPERD

>HSV2GPG_21 MHAIAPRLLLLFVLSGLPGTRGGSGVPGPINPPNNDVVFPGGSPVAQYCYAYPRLDDPGPLGSADAGRQD

LPRRVVRHEPLGRSFLTGGLVLLAPPVRGFGAPNATYAARVTYYRLTRACRQPILLRQYGGCRGGEPPSP

KTCGSYTYTYQGGGPPTRYALVNASLLVPIWDRAAETFEYQIELGGELHVGLLWVEVGGEGPGPTAPPQA

ARAEGGPCVPPVPAGRPWRSVPPVWYSAPNPGFRGLRFRERCLPPQTPAAPSDLPRVAFAPQSLLVGITG

RTFIRMARPTEDVGVLPPHWAPGALDDGPYAPFPPRPRFRRALRTDPEGVDPDVRAPLTGRRLMALTEDA

SSDSPTSAPEKTPLPVSATAMAPSVDPSAEPTAPATTTPPDEMATQAATVAVTPEETAVASPPATASVES

SPLPAAAATPGAGHTNTSSASAAKTPPTTPAPTTPPPTSTHATPRPTTPGPQTTPPGPATPGPVGASAAP

TADSPLTASPPATAPGPSAANVSVAATTATPGTRGTARTPPTDPKTHPHGPADAPPGSPAPPPPEHRGGP

EEFEGAGDGEPPEDDDSATGLAFRTPNPNKPPPARPGPIRPTLPPGILGPLAPNTPRPPAQAPAKDMPSG

PTPQHIPLFWFLTASPALDILFIISTTIHTAAFVCLVALAAQLWRGRAGRRRYAHPSVRYVCLPPERD

>HSV2GPG_22 MHAIAPRLLLLFVLSGLPGTRGGSGVPGPINPPNNDVVFPGGSPVAQYCYAYPRLDDPGPLGSADAGRQD

LPRRVVRHEPLGRSFLTGGLVLLAPPVRGFGAPNATYAARVTYYRLTRACRQPILLRQYGGCRGGEPPSP

KTCGSYTYTYQGGGPPTRYALVNASLLVPIWDRAAETFEYQIELGGELHVGLLWVEVGGEGPGPTAPPQA

AHAEGGPCVPPVPAGRPWRSVPPVWYSAPNPGFRGLRFRERCLPPQTPAAPSDLPRVAFAPQSLLVGITG

RTFIRMARPTEDVGVLPPHWAPGALDDGPYAPFPPRPRFRRALRTDPEGVDPDVRAPRTGRRLMALTENA

SSDSPTSAPEKTPLPVSATAMAPSVDPSAEPTAPATTTPPDEMATQAATVAVTPEETAVASPPATASVES

SPLPAAAATPGAGHTNTSSASAAKTPPTTPAPTTPPPTSTHATPRPTTPGPQTTPPGPATPGPVGASAAP

TADSPLTASPPATAPGPSAANVSVAATTATPGTRGTARTPPTDPKTHPHGPADAPPGSPAPPPPEHRGGP

EEFEGAGDGEPPEDDDSATGLAFRTPNPNKPPPARPGPIRPTLPPGILGPLAPNTPRPPAQAPAKDMPSG

PTPQHIPLFWFLTASPALDILFIISTTIHTAAFVCLVALAAQLWRGRAGRRRYAHPSVRYVCLPPERD

>HSV2GPG_23 MHAIAPRLLLLFVLSGLPGTRGGSGVPGPINPPNNDVVFPGGSPVAQYCYAYPRLDDPGPLGSADAGRQD

LPRRVVRHEPLGRSFLTGGLVLLAPPVRGFGAPNATYAARVTYYRLTRACRQPILLRQYGGCRGGEPPSP

KTCGSYTYTYQGGGPPTRYALVNASLLVPIWDRAAETFEYQIELGGELHVGLLWVEVGGEGPGPTAPPQA

ARAEGGPCVPPVPAGRPWRSVPPVWYSAPNPGFRGLRFRERCLPPQTPAAPSDLPRVAFAPQSLLVGITG

RTFIRMARPTGDVGVLPPHWAPGALDDGPYAPFPPRPRFRRALRTDPEGVDPDVRAPRTGRRLMALTEDA

SSDSPTSAPEKTPLPVSATAMAPSVDPSAEPTAPATTTPPDEMATQAATVAVTPEETAVASPPATASVES

SPLPAAAATPGAGHTNTSSASAAKTPPTTPAPTTPPPTSTHATPRPTTPGPQTTPPGPATPGPVGASAAP

TADSPLTASPPATAPGPSAANVSVAATTATPGTRGTARTPPTDPKTHPHGPADAPPGSPAPPPPEHRGGP

EEFEGAGDGEPPEDDDSATGLAFRTPNPNKPPPARPGPIRPTLPPGILGPLAPNTPRPPAQAPAKDMPSG

PTPQHIPLFWFLTASPALDILFIISTTIHTAAFVCLVALAAQLWRGRAGRRRYAHPSVRYVCLPPERD

>HSV2GPG_24

MHAIAPRLLLLFVLSGLPGTRGGSGVPGPINPPNNDVVFPGGSPVAQYCYAYPRLDDPGPLGSADAGRQD

LPRRVVRHEPLGRSFLTGGLVLLAPPVRGFGAPNATYAARVTYYRLTRACRQPILLRQYGGCRGGEPPSP

KTCGSYTYTYQGGGPPTRYALVNASLLVPIWDRAAETFEYQIELGGELHVGLLWVEVGGEGPGPTAPPQA

ARAEGGPCVPPVPAGRPWRSVPPVWYSAPNPGFRGLRFRERCLPPQTPAAPSDLPRVAFAPQSLLVGITG

RTFIRMARPTEDVGVLPPHWAPGALDDGPYAPFPPRPRFRRALRTDPEGVDPDVRAPLTGRRLMALTEDA

SSDSPTSAPEKTPLPVSATAMAPSVDPSAEPTAPTTTTPPDEMATQAATVAVTPEETAVASPPATASVES

SPLPAAAATPGAGHTNTSSASAAKTPPTTPAPTTPPPTSTHATPRPTTPGPQTTPPGPATPGPVGASAAP

TADSPLTASPPATAPGPSAANVSVAATTATPGTRGTARTPPTDPKTHPHGPADAPPGSPAPPPPEHRGGP

EEFEGAGDGEPPEDDDSATGLAFRTPNPNKPPPARPGPIRPTLPPGILGPLAPNTPRPPAQAPAKDMPSG

PTPQHIPLFWFLTASPALDILFIISTTIHTAAFVCLVALAAQLWRGRAGRRRYAHPSVRYVCLPPERD

>HSV2GPG_25

MHAIAPRLLLLFVLSGLPGTRGGSGVPGPINPPNNDVVFPGGSPVAQYCYAYPRLDDPGPLGSADAGRQD

LPRRVVRHEPLGRSFLTGGLVLLAPPVRGFGAPNATYAARVTYYRLTRACRQPILLRQYGGCRGGEPPSP

KTCGSYTYTYQGGGPPTRYALVNASLLVPIWDRAAETFEYQIELGGELHVGLLWVEVGGEGPGLTAPPQA

ARAEGGPCVPPVPAGRPWRSVPPVWYSAPNPGFRGLRFRERCLPPQTPAAPSDLPRVAFAPQSLLVGITG

RTFIRMARPTEDVGVLPPHWAPGALDDGPYAPFPPRPRFRRALRTDPEGVDPDVRAPRTGRRLMALTEDA

SSDSPTSAPEKTPLPVSATAMAPSVDPSAEPTAPATTTPPDEMATQAATVAVTPEETAVASPPATASVES

SPLPAAAATPGAGHTNTSSASAAKTPPTTPAPTTPPPTSTHATPRPTTPGPQTTPPGPATPGPVGASAAP

TADSPLTASPPATAPGPSAANVSVAATTATPGTRGTARTPPTDPKTHPHGPADAPPGSPAPPPPEHRGGP

EEFEGAGDGEPPEDDDSATGLAFRTPNPNKPPPARPGPIRPTLPPGILGPLAPNTPRPPAQAPAKDMPSG

PTPQHIPLFWFLTASPALDILFIISTTIHTAAFVCLVALAAQLWRGRAGRRRYAHPSVRYVCLPPERD

>HSV2GPG_26

MHAIAPRLLLLFVLSGLPGTRGGSGVPGPINPPNNDVVFPGGSPVAQYCYAYPRLDDPGPLGSADAGRQD

LPRRVVRHEPLGRSFLTGGLVLLAPPVRGFGAPNATYAARVTYYRLTRACRRPILLRQYGGCRGGEPPSP

KTCGSYTYTYQGGGPPTRYALVNASLLVPIWDRAAETFEYQIELGGELHVGLLWVEVGGEGPGPTAPPQA

ARAEGGPCVPPVPAGRPWRSVPPVWYSAPNPGFRGLRFRERCLPPQTPAAPSDLPRVAFAPQSLLVGITG

RTFIRMARPTEDVGVLPPHWAPGALNDGPYAPFPPRPRFRRALRTDPEGVDPDVRAPRTGRRLMALTEDA

SSDSPTSAPEKTPLPVSATAMAPSVDPSAEPTAPATTTPPDEMATQAATVAVTPEETAVASPPATASVES

SPLPAAAATPGAGHTNTSSASAAKTPPTTPAPTTPPPTSTHATPRPTTPGPQTTPPGPATPGPVGASAAP

TADSPLTASPPATAPGPSAANVSVAATTATPGTRGTARTPPTDPKTHPHGPADAPPGSPAPPPPEHRGGP

EEFEGAGDGEPPEDDDSATGLAFRTPNPNKPPPARPGPIRPTLPPGILGPLAPNTPRPPAQAPAKDMPSG

PTPQHIPLFWFLTASPALDILFIISTTIHTAAFVCLVALAAQLWRGRAGRRRYAHPSVRYVCLPPERD
