## Supplementary file 6 for "Contriving a chimeric polyvalent vaccine to prevent infections caused by Herpes Simplex Virus (Type-1 and Type-2): an exploratory immunoinformatic approach"

**Envelope protein UL45**

>HS1UL45_1 MPLRASEHAYRPLGPGTPPMRARLPAAAWVGVGTIIGGVVIIAALVLVPSRASWALSPCDSGWHEFNLGC

ISWDPTPMEHEQAVGGCSAPATLIPRAAAKQLAAVARVQSARSSGYWWVSGDGIRACLRLVDGVGGIDQF

CEEPALRICYYPRSPGGFVQFVTSTRNALGLP

>HS1UL45_2 MVDPERHTAWSKTPPPEPLSYQHTGMPLRASEHAYRPLGPGTPPMRARLPAAAWVGVGTIIGGVVIIAAL

VLVPSRASWALSPCDSGWHEFNLGCISWDPTPMEHEQAVGGCSAPATLIPRAAAKQLAAVARVQSARSSG

YWWVSGDGIRACLRLVDGVGGIDQFCEEPALRICYYPRSPGGFVQFVTSTRNALGLP

>HS1UL45_3 MPLRAAEHAYRPLGPGTPPMRARLPAAAWVGVGTIIGGVVIIAALVLVPSRASWALSPCDSGWHEFNLGC

ISWDPTPMEHEQAVGGCSAPATLIPRAAAKQLAAVARVQSARSSGYWWVSGDGIRACLRLVDGVGGIDQF

CEEPALRICYYPRSPGGFVQFVTSTRNALGLP

>HS1UL45_4 MPLRASEHAYRPLGPGTPPMRARLPAAAWVGVGTIIGGVVIIAALVLVPSRASWALSPCDSGWHEFNLGC

ISWDPTPMEHEQAVGGCSAPATLIPRAAAKQLAAVTRVQSARSSGYWWVSGDGIRACLRLVDGVGGIDQF

CEEPALRICYYPRSPGGFVQFVTSTRNALGLP

>HS1UL45_5 MPLRASEHAYRPLGPGTPPMRARLPAAVWVGVGTIIGGVVIIAALVLVPSRASWALSPCDSGWHEFNLGC

ISWDPTPMEHEQAVGGCSAPATLIPRAAAKQLAAVARVQSARSSGYWWVSGDGIRACLRLVDGVGGIDQF

CEEPALRICYYPRSPGGFVQFVTSTRNALGLP

>HS1UL45_6

MPLRASEHAYRPLGPGTPPVRARLPAAAWVGVGTIIGGVVIIAALVLVPSRASWALSPCDSGWHEFNLGC

ISWDPTPMEHEQAVGGCSAPATLIPRAAAKQLAAVARVQSARSSGYWWVSGDGIRACLRLVDGVGGIDQF

CEEPALRICYYPRSPGGFVQFVTSTRNALGLP

>HS1UL45_7 MPLRASEHAYRPLGPGTPPTRARLPAAAWVGVGTIIGGVVIIAALVLVPSRASWALSPCDSGWHEFNLGC

ISWDPTPMEHEQAVGGCSAPATLIPRAAAKQLAAVARVQSARSSGYWWVSGDGIRACLRLVDGVGGIDQF

CEEPALRICYYPRSPGGFVQFVTSTRNALGLP

>HS1UL45_8 MPLRASEHAYRPLGPGTPPMRARLPAAAWVGVGTIIGGVVIIAALVLVPSRASWALSPCDSGWHEFNLGC

ISWDPTPMEHEQAVGGCGAPATLIPRAAAKQLAAVARVQSARSSGYWWVSGDGIRACLRLVDGVGGIDQF

CEEPALRICYYPRSPGGFVQFVTSTRNALGLP

>HS1UL45_9 MPLRASEHAYRPLGPGPPPMRARLPAAAWVGVGTIIGGVVIIAALVLVPSRASWALSPCDSGWHEFNLGC

ISWDPTPMEHEQAVGGCSAPATLIPRAAAKQLAAVARVQSARSSGYWWVSGDGIRACLRLVDGVGGIDQF

CEEPALRICYYPRSPGGFVQFVTSTRNALGLP

>HS1UL45_10

MPLRASEHAYRPLGPGTPPMRARLPAAAWVGVGTIIGGVVIIAALVLVPSRASWALSPCDSGWHEFNLGC

ISWDPTPMEHEQAVGGCSAPATLIPRAAAKQLAAVARVQSARSSGYWWVSGDGIRARLRLVDGVGGIDQF

CEEPALRICYYPRSPGGFVQFVTSTRNALGLP

>HS1UL45_11 MPLRASEHAYRPLGPGTPPVRARLPAAAWVGVGTIIGGVVIIAALVLVPSRASWALSPCDSGWHEFNLGC

ISWDPTPMEHEQAVGGCSAPATLIPRAAAKQLAAVARVQSARSSGYWWVSGDGIRARLRLVDGVGGIDQF

CEEPALRICYYPRSPGGFVQFVTSTRNALGLP

>HS2UL45_12 MAFRASGPAYQPLAPAASPARARVPAVAWIGVGAIVGAFALVAALVLVPPRSSWGLSPCDSGWQEFNAGC

VAWDPTPVEHEQAVGGCSAPATLIPRAAAKHLAALTRVQAERSSGYWWVNGDGIRTCLRLVDSVSGIDEF

FEELAIRICYYPRSPGGFVRFVTSIRNALGLP

>HS2UL45_13

MAFRASGPAYQPLAPAASPARARVPAVAWIGVGAIVGAFALVAALVLVPPRSSWGLSPCDSGWQEFNAGC

VAWDPTPVEHEQAVGGCSAPATLIPRAAAKHLAALTRVQAERSSGYWWVNGDGIRTCLRLVDSVSGIDEF

CEELAIRICYYPRSPGGFVRFVTSIRNALGLP

>HS2UL45_14 MAFRASGPAYRPLAPAASSARAGLPAVAWIGVGTIVGAFALVAALVLVPPRSSWGLSPCASGWQEFNAGC

VSWDPTPVEHEQAVGGCSAPATLIPRAAAKHLAALARVQAERSSGYWWVSGDGIRACLRLVDSVSGIDQF

CEEPAIRICYYPRSPGGFVRFVTSIRNTLGLP

>HS2UL45_15 MAFRASGPAYQPLAPRPPPARARVPAVAWIGVGAIVGAFALVAALVLVPPRSSWGLCPCDSGWQEFNAGC

VAWDPTPVEHEQAVGGCSAPATLIPRAAAKHLAALTRVQAERSSGYWWVNGDGIRTCLRLVDSVSGIDEF

CEEL

>HS2UL45_16

MPLRASEHAYRPLGPGTPPMRARLPAAAWVGVGTIIGGVVIIAALVLVPSRASWALSPCDSGWHEFNLGC

ISWDPTPMEHEQAVGGCSAPATLIPRAAAKQLAAVTRVQSARSSGYWWVSGDGIRACLRLVDGVGGIDQF

CEEPALRICYYPRSPGGFVQFVTSTRNALGLP
