## Supplementary file 7 for "Contriving a chimeric polyvalent vaccine to prevent infections caused by Herpes Simplex Virus (Type-1 and Type-2): an exploratory immunoinformatic approach"

```
Alignment of GLYCOPROTEIN B


HS1GPB_3        MRQGAPARGRRWFVVWALLGLTLGVLVVSAAPSSPGTPGVAAATQAANGGPATPAPPAPG 60
HS1GPB_5        MRQGAPARGRRWFVVWALLGLTLGVLVASAAPSSPGTPGVAAATQAANGGPATPAPPAPG 60
HS1GPB_7        ------------------------------------------------------------
HS1GPB_9        MRQGAPARGRRWFVVWALLGLTLGVLVASAAPSSPGTPGVAAATQAANGGPATPAPPAPG 60
HS1GPB_4        ------------------------------------------------------------
HS1GPB_1        ------------------------------------------------------------
HS1GPB_2        ------------------------------------------------------------
HS1GPB_12       MRQGAPTRGCRWFVVWALLGLTLGVLVASAAPSSPGTPGVAAATQAANGGPATPAPPAPG 60
HS1GPB_13       MRQGAPARGCRWFVVWALLGLTLGVLVASAAPSSPGTPGVAAATQAANGGPATPAPPALG 60
HS1GPB_16       MRQGAPARGCRWFVVWVLLGLTLGVLVASAAPSSPGTPGVAAATQAANGGPATPAPPALG 60
HS1GPB_8        ------------------------------------------------------------
HS1GPB_6        MRQGAPARGRRWFVVWALLGLTLGVLVASAAPSSPGTPGVAAATQAANGGPATPAPPALG 60
HS1GPB_11       MRQGAPARGRRWFVVWALLGLTLGVLVASAAPSSPGTPGVAAATPAANGGPATPAPPALG 60
HS1GPB_15       MRQGAPARGRRWFVVWALLGLTLGVLVASAAPSSPGTPGGAAATQAADGGPATPAPPALG 60
HS1GPB_19       ------------------------------------------------------------
HS1GPB_14       ------------------------------------------------------------
HS1GPB_17       MRQGAPARGCRWFVVWALLGLTLGVLVASAAPSSPGTPGVAAATQAANGGPATPAPPAPG 60
HS1GPB_20       ------------------------------------------------------------
HS1GPB_10       MRQGAPARGRRWFVVWALLGLTLGVLVASAAPSSPGTPGVAAATQAANGGPATPAPPAPG 60
HS1GPB_18       MRQGAPARGCRWFVVWALLGLTLGVLVASAAPSSPGTPGVAAATQAANGGPATPAPPAPG 60
HS2GPB_21       -----MRGGGLICALVVGALVAAVASAAPAAPAAPRASGGVAATVAANGGPASRPPPVPS 55
HS2GPB_22       -----MRGGGLICALVVGALVAAVASAAPAAPAAPRASGGVAATVAANGGPASRPPPVPS 55
HS2GPB_27       -----MRGGGLICALVVGALVAAVASAAPAAPAAPRASGGVAATVAANGGPASRPPPVPS 55
HS2GPB_30       -----MRGGGLICALVVGALVAAVASAAPAAPAAPRASGGVAATVAANGGPASRPPPVPS 55
HS2GPB_31       -----MRGGGLICALVVGALVAAVASAAPAAPAAPRASGGVAATVAANGGPASRPPPVPS 55
HS2GPB_34       -----MRGGGLICALVVGALVAAVASAAPAAPAAPRASGGVAATVAAKGGPASRPPPVPS 55
HS2GPB_28       -----MRGGGLICALVVGALVAAVASAAPAAPAAPRASGGVAATVAANGGPASRPPPVPS 55
HS2GPB_24       -----MRGGGLICALVVGALVAAVASAAPAAPAAPRASGGVAATVAANGGPASRPPPVPS 55
HS2GPB_26       -----MRGGGLICALVVGALVAAVASAAPAAPAAPRASGGVAATVAANGGPASRPPPVPS 55
HS2GPB_35       -----MRGGGLICALVVGALVAAVASAAP---AAPRASGGVAATVAANGGPASRPPPVPS 52
HS2GPB_37       -----MRGGGLICALVVGALVAAVASAAP---AAPRASGGVAATVAANGGPASRPPPVPS 52
HS2GPB_36       -----MRGGGLICALVVGALVAAVASAAP---AAPRASGGVAATVAANGGPASRPPPVPS 52
HS2GPB_25       ------------------------------------------------------------
HS2GPB_23       ------------------------------------------------------------
HS2GPB_29       ------------------------------------------------------------
HS2GPB_32       ------------------------------------------------------------
HS2GPB_33       ------------------------------------------------------------
HS2GPB_38       ------------------------------------------------------------
HS2GPB_39       ------------------------------------------------------------


HS1GPB_3        APPTGDPKPKKNKKPKPPKPPRPAGDNATVAAGHATLREHLRDIKAENTDANFYVCPPPT 120
HS1GPB_5        APPTGDPKPKKNKKPKPPKPPRPAGDNATVAAGHATLREHLRDIKAENTDANFYVCPPPT 120
HS1GPB_7        ----------KNKKPKPPKLPRPAGDNATVAAGHATLREHLRDIKAENTDANFYVCPPPT 50
HS1GPB_9        APPTGDPKPKKNKKPKPPKLPRPAGDNATVAAGHATLREHLRDIKAENTDANFYVCPPPT 120
HS1GPB_4        ----------KNKKPKPPKPPRPAGDNATVAAGHATLREHLRDIKAENTDANFYVCPPPT 50
HS1GPB_1        ----------KNRKPKPPKPPRPAGDNATVAAGHATLREHLRDIKAENTDANFYVCPPPT 50
HS1GPB_2        ----------KNKKPKPPKPPRPAGDNATVAAGHATLREHLRDIKAENTDANFYVCPPPT 50
HS1GPB_12       PAPTGDPKPRKNKKPKPPTPPRPAGDNATVAAGHATLREHLRDIKAENTDANFYVCPPPT 120
HS1GPB_13       AAPTGDPKPKKNKKPKNPTPPRPAGDNATVAAGHATLREHLRDIKAENTDANFYVCPPPT 120
HS1GPB_16       AAPTGDPKPKKNKKPKNPTPPRPAGDNATVAAGHATLREHLRDIKAENTDANFYVCPPPT 120
HS1GPB_8        ----------KNKKPKNPTPPRPAGDNATVAAGHATLREHLRDIKAENTDANFYVCPPPT 50
HS1GPB_6        AAPTGDPKPKKNKKPKNPTPPRPAGDNATVAAGHATLREHLRDIKAENTDANFYVCPPPT 120
HS1GPB_11       AAPTGDPKPKKNKKPKNPTPPRPAGDNATVAAGHATLREHLRDIKAENTDANFYVCPPPT 120
HS1GPB_15       AAPTGDPKPKKNKKPKNPTPPRPAGDNATVAAGHATLREHLRDIKAENTDANFYVCPPPT 120
HS1GPB_19       ----------KNKKPKNPTPPRPAGDNATVAAGHATLREHLRDIKAENTDANFYVCPPPT 50
HS1GPB_14       ----------KNKKPKNPPPPRPAGDNATVAAGHATLREHLRDIKAENTDANFYVCPPPT 50
HS1GPB_17       PAPTGDTKPKKNKKPKNPPPPRPAGDNATVAAGHATLREHLRDIKAENTDANFYVCPPPT 120
HS1GPB_20       ----------KNKKPKNPPPPRPAGDNATVAAGHATLREHLRDIKAENTDANFYVCPPPT 50
HS1GPB_10       PAPTGDPKPRKNKKPKNPTPPRPAGDNATVAAGHATLREHLRDIKAENTDANFYVCPPPT 120
HS1GPB_18       PAPTGDTKPQKNKKPKPPXXPRPAGDNATVAAGHATLREHLRDIKAENTDANFYVCPPPT 120
HS2GPB_21       PATTKARKRKTKKPPKRPEATPPPDANATVAAGHATLRAHLREIKVENADAQFYVCPPPT 115
HS2GPB_22       PATTKARKRKTKKPPKRPEATPPPDANATVAAGHATLRAHLREIKVENADAQFYVCPPPT 115
HS2GPB_27       PATTKARKRKTKKPPKRPEATPPPDANATVAAGHATLRAHLREIKVENADAQFYVCPPPT 115
HS2GPB_30       PATTKARKRKTKKPPKRPEATPPPDANATVAAGHATLRAHLREIKVENADAQFYVCPPPT 115
HS2GPB_31       PATTKARKRKTKKPPKRPEATPPPDANATVAAGHATLRAHLREIKVENADAQFYVCPPPT 115
HS2GPB_34       PATTKARKRKTKKPPKRPEATPPPDANATVAAGHATLRAHLREIKVENADAQFYVCPPPT 115
HS2GPB_28       PATTKARKRKTKKPPERPEATPPPDANATVAAGHATLRAHLREIKVENADAQFYVCPPPT 115
HS2GPB_24       PATTKARKRKTKKPPKRPEATPPPDANATVAAGHATLRAHLREIKVENADAQFYVCPPPT 115
HS2GPB_26       PATTRARKRKTKKPPKRPEATPPPDANATVAAGHATLRAHLREIKVENADAQFYVCPPPT 115
HS2GPB_35       PATTRARKRKTKKPPKRPEATPPPDANATVAAGHATLRAHLREIKVENADAQFYVCPPPT 112
HS2GPB_37       PATTKARKRKTKKPPKRPEATPPPDANATVAAGHATLRAHLREIKVENADAQFYVCPPPT 112
HS2GPB_36       PATTKARKRKTKKPPERPEATPPPDANATVAAGHATLRAHLREIKVENADAQFYVCPPPT 112
HS2GPB_25       ---------------ERPEATPPPDANATVAAGHATLRAHLREIKVENADAQFYVCPPPT 45
HS2GPB_23       ---------------KRPEATPPPDANATVAAGHATLRAHLREIKVENADAQFYVCPPPT 45
HS2GPB_29       ---------------KRPEATPPPDANATVAAGHATLRAHLREIKVENADAQFYVCPPPT 45
HS2GPB_32       ---------------KRPEATPPPDANATVAAGHATLRAHLREIKVENADAQFYVCPPPT 45
HS2GPB_33       ------------------EATPPPDANATVAAGHATLRAHLREIKVENADAQFYVCPPPT 42
HS2GPB_38       ------------------EATPPPDANATVAAGHATLRAHLREIKVENADAQFYVCPPPT 42
HS2GPB_39       ------------------EATPPPDANATVAAGHATLRAHLREIKVENADAQFYVCPPPT 42
                                    . *.. ************ ***:**.**:**:********

HS1GPB_3        GATVVQFEQPRRCPTRPEGQNYTEGIAVVFKENIAPYKFKATMYYKDVTVSQVWFGHRYS 180
HS1GPB_5        GATVVQFEQPRRCPTRPEGQNYTEGIAVVFKENIAPYKFKATMYYKDVTVSQVWFGHRYS 180
HS1GPB_7        GATVVQFEQPRRCPTRPEGQNYTEGIAVVFKENIAPYKFKATMYYKDVTVSQVWFGHRYS 110
HS1GPB_9        GATVVQFEQPRRCPTRPEGQNYTEGIAVVFKENIAPYKFKATMYYKDVTVSQVWFGHRYS 180
HS1GPB_4        GATVVQFEQPRRCPTRPEGQNYTEGIAVVFKENIAPYKFKATMYYKDVTVSQVWFGHRYS 110
HS1GPB_1        GATVVQFEQPRRCPTRPEGQNYTEGIAVVFKENIAPYKFKATMYYKDVTVSQVWFGHRYS 110
HS1GPB_2        GATVVQFEQPRRCPTRPEGQNYTEGIAVVFKENIAPYKFKATMYYKDVTVSQVWFGHRYS 110
HS1GPB_12       GATVVQFEQPRRCPTRPEGQNYTEGIAVVFKENIAPYKFKATMYYKDVTVSQVWFGHRYS 180
HS1GPB_13       GATVVQFEQPRRCPTRPEGQNYTEGIAVVFKENIAPYKFKATMYYKDVTVSQVWFGHRYS 180
HS1GPB_16       GATVVQFEQPRRCPTRPEGQNYTEGIAVVFKENIAPYKFKATMYYKDVTVSQVWFGHRYS 180
HS1GPB_8        GATVVQFEQPRRCPTRPEGQNYTEGIAVVFKENIAPYKFKATMYYKDVTVSQVWFGHRYS 110
HS1GPB_6        GATVVQFEQPRRCPTRPEGQNYTEGIAVVFKENIAPYKFKATMYYKDVTVSQVWFGHRYS 180
HS1GPB_11       GATVVQFEQPRRCPTRPEGQNYTEGIAVVFKENIAPYKFKATMYYKDVTVSQVWFGHRYS 180
HS1GPB_15       GATVVQFEQPRRCPTRPEGQNYTEGIAVVFKENIAPYKFKATMYYKDVTVSQVWFGHRYS 180
HS1GPB_19       GATVVQFEQPRRCPTRPEGQNYTEGIAVVFKENIAPYKFKATMYYKDVTVSQVWFGHRYS 110
HS1GPB_14       GATVVQFEQPRRCPTRPEGQNYTEGIAVVFKENIAPYKFKATMYYKDVTVSQVWFGHRYS 110
HS1GPB_17       GATVVQFEQPRRCPTRPEGQNYTEGIAVVFKENIAPYKFKATMYYKDVTVSQVWFGHRYS 180
HS1GPB_20       GATVVQFEQPRRCPTRPEGQNYTEGIAVVFKENIAPYKFKATMYYKDVTVSQVWFGHRYS 110
HS1GPB_10       GATVVQFEQPRRCPTRPEGQNYTEGIAVVFKENIAPYKFKATMYYKDVTVSQVWFGHRYS 180
HS1GPB_18       GATVVQFEQPRRCPTRPEGQNYTEGIAVVFKENIAPYKFKATMYYKDVTVSQVWFGHRYS 180
HS2GPB_21       GATVVQFEQPRRCPTRPEGQNYTEGIAVVFKENIAPYKFKATMYYKDVTVSQVWFGHRYS 175
HS2GPB_22       GATVVQFEQPRRCPTRPEGQNYTEGIAVVFKENIAPYKFKATMYYKDVTVSQVWFGHRYS 175
HS2GPB_27       GATVVQFEQPRRCPTRPEGQNYTEGIAVVFKENIAPYKFKATMYYKDVTVSQVWFGHRYS 175
HS2GPB_30       GATVVQFEQPRRCPTRPEGQNYTEGIAVVFKENIAPYKFKATMYYKDVTVSQVWFGHRYS 175
HS2GPB_31       GATVVQFEQPRRCPTRPEGQNYTEGIAVVFKENIAPYKFKATMYYKDVTVSQVWFGHRYS 175
HS2GPB_34       GATVVQFEQPRRCPTRPEGQNYTEGIAVVFKENIAPYKFKATMYYKDVTVSQVWFGHRYS 175
HS2GPB_28       GATVVQFEQPRRCPTRPEGQNYTEGIAVVFKENIAPYKFKATMYYKDVTVSQVWFGHRYS 175
HS2GPB_24       GATVVQFEQPRRCPTRPEGQNYTEGIAVVFKENIAPYKFKATMYYKDVTVSQVWFGHRYS 175
HS2GPB_26       GATVVQFEQPRRCPTRPEGQNYTEGIAVVFKENIAPYKFKATMYYKDVTVSQVWFGHRYS 175
HS2GPB_35       GATVVQFEQPRRCPTRPEGQNYTEGIAVVFKENIAPYKFKATMYYKDVTVSQVWFGHRYS 172
HS2GPB_37       GATVVQFEQPRRCPTRPEGQNYTEGIAVVFKENIAPYKFKATMYYKDVTVSQVWFGHRYS 172
HS2GPB_36       GATVVQFEQPRRCPTRPEGQNYTEGIAVVFKENIAPYKFKATMYYKDVTVSQVWFGHRYS 172
HS2GPB_25       GATVVQFEQPRRCPTRPEGQNYTEGIAVVFKENIAPYKFKATMYYKDVTVSQVWFGHRYS 105
HS2GPB_23       GATVVQFEQPRRCPTRPEGQNYTEGIAVVFKENIAPYKFKATMYYKDVTVSQVWFGHRYS 105
HS2GPB_29       GATVVQFEQPRRCPTRPEGQNYTEGIAVVFKENIAPYKFKATMYYKDVTVSQVWFGHRYS 105
HS2GPB_32       GATVVQFEQPRRCPTRPEGQNYTEGIAVVFKENIAPYKFKATMYYKDVTVSQVWFGHRYS 105
HS2GPB_33       GATVVQFEQPRRCPTRPEGQNYTEGIAVVFKENIAPYKFKATMYYKDVTVSQVWFGHRYS 102
HS2GPB_38       GATVVQFEQPRRCPTRPEGQNYTEGIAVVFKENIAPYKFKATMYYKDVTVSQVWFGHRYS 102
HS2GPB_39       GATVVQFEQPRRCPTRPEGQNYTEGIAVVFKENIAPYKFKATMYYKDVTVSQVWFGHRYS 102
                ************************************************************

HS1GPB_3        QFMGIFEDRAPVPFEEVIDKINAKGVCRSTAKYVRNNLETTAFHRDDHETDMELKPANAA 240
HS1GPB_5        QFMGIFEDRAPVPFEEVIDKINAKGVCRSTAKYVRNNLETTAFHRDDHETDMELKPANAA 240
HS1GPB_7        QFMGIFEDRAPVPFEEVIDKINAKGVCRSTAKYVRNNLETTAFHRDDHETDMELKPANAA 170
HS1GPB_9        QFMGIFEDRAPVPFEEVIDKINAKGVCRSTAKYVRNNLETTAFHRDDHETDMELKPANAA 240
HS1GPB_4        QFMGIFEDRAPVPFEEVIDKINAKGVCRSTAKYVRNNLETTAFHRDDHETDMELKPANAA 170
HS1GPB_1        QFMGIFEDRAPVPFEEVIDKINAKGVCRSTAKYVRNNLETTAFHRDDHETDMELKPANAA 170
HS1GPB_2        QFMGIFEDRAPVPFEEVIDKINAKGVCRSTAKYVRNNLETTAFHRDDHETDMELKPANAA 170
HS1GPB_12       QFMGIFEDRAPVPFEEVIDKINAKGVCRSTAKYVRNNLETTAFHRDDHETDMELKPANAA 240
HS1GPB_13       QFMGIFEDRAPVPFEEVIDKINAKGVCRSTAKYVRNNLETTAFHRDDHETDMELKPANAA 240
HS1GPB_16       QFMGIFEDRAPVPFEEVIDKINAKGVCRSTAKYVRNNLETTAFHRDDHETDMELKPANAA 240
HS1GPB_8        QFMGIFEDRAPVPFEEVIDKINAKGVCRSTAKYVRNNLETTAFHRDDHETDMELKPANAA 170
HS1GPB_6        QFMGIFEDRAPVPFEEVIDKINAKGVCRSTAKYVRNNLETTAFHRDDHETDMELKPANAA 240
HS1GPB_11       QFMGIFEDRAPVPFEEVIDKINAKGVCRSTAKYVRNNLETTAFHRDDHETDMELKPANAA 240
HS1GPB_15       QFMGIFEDRAPVPFEEVIDKINAKGVCRSTAKYVRNNLETTAFHRDDHETDMELKPANAA 240
HS1GPB_19       QFMGIFEDRAPVPFEEVIDKINAKGVCRSTAKYVRNNLETTAFHRDDHETDMELKPANAA 170
HS1GPB_14       QFMGIFEDRAPVPFEEVIDKINAKGVCRSTAKYVRNNLETTAFHRDDHETDMELKPANAA 170
HS1GPB_17       QFMGIFEDRAPVPFEEVIDKINAKGVCRSTAKYVRNNLETTAFHRDDHETDMELKPANAA 240
HS1GPB_20       QFMGIFEDRAPVPFEEVIDKINAKGVCRSTAKYVRNNLETTAFHRDDHETDMELKPANAA 170
HS1GPB_10       QFMGIFEDRAPVPFEEVIDKINAKGVCRSTAKYVRNNLETTAFHRDDHETDMELKPANAA 240
HS1GPB_18       QFMGIFEDRAPVPFEEVIDKINAKGVCRSTAKYVRNNLETTAFHRDDHETDMELKPANAA 240
HS2GPB_21       QFMGIFEDRAPVPFEEVIDKINTKGVCRSTAKYVRNNMETTAFHRDDHETDMELKPAKVA 235
HS2GPB_22       QFMGIFEDRAPVPFEEVIDKINAKGVCRSTAKYVRNNMETTAFHRDDHETDMELKPAKVA 235
HS2GPB_27       QFMGIFEDRAPVPFEEVIDKINAKGVCRSTAKYVRNNMETTAFHRDDHETDMELKPAKVA 235
HS2GPB_30       QFMGIFEDRAPVPFEEVIDKINAKGVCRSTAKYVRNNMETTAFHRDDHETDMELKPAKVA 235
HS2GPB_31       QFMGIFEDRAPVPFEEVIDKINAKGVCRSTAKYVRNNMETTAFHRDDHETDMELKPAKVA 235
HS2GPB_34       QFMGIFEDRAPVPFEEVIDKINAKGVCRSTAKYVRNNMETTAFHRDDHETDMELKPAKVA 235
HS2GPB_28       QFMGIFEDRAPVPFEEVIDKINAKGVCRSTAKYVRNNMETTAFHRDDHETDMELKPAKVA 235
HS2GPB_24       QFMGIFEDRAPVPFEEVIDKINAKGVCRSTAKYVRNNMETTAFHRDDHETDMELKPAKVA 235
HS2GPB_26       QFMGIFEDRAPVPFEEVIDKINAKGVCRSTAKYVRNNMETTAFHRDDHETDMELKPAKVA 235
HS2GPB_35       QFMGIFEDRAPVPFEEVIDKINTKGVCRSTAKYVRNNMETTAFHRDDHETDMELKPAKVA 232
HS2GPB_37       QFMGIFEDRAPVPFEEVIDKINAKGVCRSTAKYVRNNMETTAFHRDDHETDMELKPAKVA 232
HS2GPB_36       QFMGIFEDRAPVPFEEVIDKINAKGVCRSTAKYVRNNMETTAFHRDDHETDMELKPAKVA 232
HS2GPB_25       QFMGIFEDRAPVPFEEVIDKINAKGVCRSTAKYVRNNMETTAFHRDDHETDMELKPAKVA 165
HS2GPB_23       QFMGIFEDRAPVPFEEVIDKINAKGVCRSTAKYVRNNMETTAFHRDDHETDMELKPAKVA 165
HS2GPB_29       QFMGIFEDRAPVPFEEVIDKINAKGVCRSTAKYVRNNMETTAFHRDDHETDMELKPAKVA 165
HS2GPB_32       QFMGIFEDRAPVPFEEVIDKINAKGVCRSTAKYVRNNMETTAFHRDDHETDMELKPAKVA 165
HS2GPB_33       QFMGIFEDRAPVPFEEVIDKINAKGVCRSTAKYVRNNMETTAFHRDDHETDMELKPAKVA 162
HS2GPB_38       QFMGIFEDRAPVPFEEVIDKINAKGVCRSTAKYVRNNMETTAFHRDDHETDMELKPAKVA 162
HS2GPB_39       QFMGIFEDRAPVPFEEVIDKINAKGVCRSTAKYVRNNMETTAFHRDDHETDMELKPAKVA 162
                **********************:**************:*******************:.*

HS1GPB_3        TRTSRGWHTTDLKYNPSRVEAFHRYGTTVNCIVEEVDARSVYPYDEFVLATGDFVYMSPF 300
HS1GPB_5        TRTSRGWHTTDLKYNPSRVEAFHRYGTTVNCIVEEVDARSVYPYNEFVLATGDFVYMSPF 300
HS1GPB_7        TRTSRGWHTTDLKYNPSRVEAFHRYGTTVNCIVEEVDARSVYPYDEFVLATGDFVYMSPF 230
HS1GPB_9        TRTSRGWHTTDLKYNPSRVEAFHRYGTTVNCIVEEVDARSVYPYDEFVLATGDFVYMSPF 300
HS1GPB_4        TRTSRGWHTTDLKYNPSRVEAFHRYGTTVNCIVEEVDARSVYPYDEFVLATGDFVYMSPF 230
HS1GPB_1        TRTSRGWHTTDLKYNPSRVEAFHRYGTTVNCIVEEVDARSVYPYDEFVLATGDFVYMSPF 230
HS1GPB_2        TRTSRGWHTTDLKYNPSRVEAFHRYGTTVNCIVEEVDARSVYPYDEFVLATGDFVYMSPF 230
HS1GPB_12       TRTSRGWHTTDLKYNPSRVEAFHRYGTTVNCIVEEVDARSVYPYDEFVLATGDFVYMSPF 300
HS1GPB_13       TRTSRGWHTTDLKYNPSRVEAFHRYGTTVNCIVEEVDARSVYPYDEFVLATGDFVYMSPF 300
HS1GPB_16       TRTSRGWHTTDLKYNPSRVEAFHRYGTTVNCIVEEVDARSVYPYDEFVLATGDFVYMSPF 300
HS1GPB_8        TRTSRGWHTTDLKYNPSRVEAFHRYGTTVNCIVEEVDARSVYPYDEFVLATGDFVYMSPF 230
HS1GPB_6        TRTSRGWHTTDLKYNPSRVEAFHRYGTTVNCIVEEVDARSVYPYDEFVLATGDFVYMSPF 300
HS1GPB_11       TRTSRGWHTTDLKYNPSRVEAFHRYGTTVNCIVEEVDARSVYPYDEFVLATGDFVYMSPF 300
HS1GPB_15       TRTSRGWHTTDLKYNPSRVEAFHRYGTTVNCIVEEVDARSVYPYDEFVLATGDFVYMSPF 300
HS1GPB_19       TRTSRGWHTTDLKYNPSRVEAFHRYGTTVNCIVEEVDARSVYPYDEFVLATGDFVYMSPF 230
HS1GPB_14       TRTSRGWHTTDLKYNPSRVEAFHRYGTTVNCIVEEVDARSVYPYDEFVLATGDFVYMSPF 230
HS1GPB_17       TRTSRGWHTTDLKYNPSRVEAFHRYGTTVNCIVEEVDARSVYPYDEFVLATGDFVYMSPF 300
HS1GPB_20       TRTSRGWHTTDLKYNPSRVEAFHRYGTTVNCIVEEVDARSVYPYDEFVLATGDFVYMSPF 230
HS1GPB_10       TRTSRGWHTTDLKYNPSRVEAFHRYGTTVNCIVEEVDARSVYPYDEFVLATGDFVYMSPF 300
HS1GPB_18       TRTSRGWHTTDLKYNPSRVEAFHRYGTTVNCIVEEVDARSVYPYDEFVLATGDFVYMSPF 300
HS2GPB_21       TRTSRGWHTTDLKYNPSRVEAFHRYGTTVNCIVEEVDARSVYPYDEFVLATGDFVYMSPF 295
HS2GPB_22       TRTSRGWHTTDLKYNPSRVEAFHRYGTTVNCIVEEVDARSVYPYDEFVLATGDFVYMSPF 295
HS2GPB_27       TRTSRGWHTTDLKYNPSRVEAFHRYGTTVNCIVEEVDARSVYPYDEFVLATGDFVYMSPF 295
HS2GPB_30       TRTSRGWHTTDLKYNPSRVEAFHRYGTTVTCIVEEVDARSVYPYDEFVLATGDFVYMSPF 295
HS2GPB_31       TRTSRGWHTTDLKYNPSRVEAFHRYGTTVNCIVEEVDARSVYPYDEFVLATGDFVYMSPF 295
HS2GPB_34       TRTSRGWHTTDLKYNPSRVEAFHRYGTTVNCIVEEVDARSVYPYDEFVLATGDFVYMSPF 295
HS2GPB_28       TRTSRGWHTTDLKYNPSRVEAFHRYGTTVNCIVEEVDARSVYPYDEFVLATGDFVYMSPF 295
HS2GPB_24       TRTSRGWHTTDLKYNPSRVEAFHRYGTTVNCIVEEVDARSVYPYDEFVLATGDFVYMSPF 295
HS2GPB_26       TRTSRGWHTTDLKYNPSRVEAFHRYGTTVNCIVEEVDARSVYPYDEFVLATGDFVYMSPF 295
HS2GPB_35       TRTSRGWHTTDLKYNPSRVEAFHRYGTTVNCIVEEVDARSVYPYDEFVLATGDFVYMSPF 292
HS2GPB_37       TRTSRGWHTTDLKYNPSRVEAFHRYGTTVNCIVEEVDARSVYPYDEFVLATGDFVYMSPF 292
HS2GPB_36       TRTSRGWHTTDLKYNPSRVEAFHRYGTTVNCIVEEVDARSVYPYDEFVLATGDFVYMSPF 292
HS2GPB_25       TRTSRGWHTTDLKYNPSRVEAFHRYGTTVNCIVEEVDARSVYPYDEFVLATGDFVYMSPF 225
HS2GPB_23       TRTSRGWHTTDLKYNPSRVEAFHRYGTTVNCIVEEVDARSVYPYDEFVLATGDFVYMSPF 225
HS2GPB_29       TRTSRGWHTTDLKYNPSRVEAFHRYGTTVNCIVEEVDARSVYPYDEFVLATGDFVYMSPF 225
HS2GPB_32       TRTSRGWHTTDLKYNPSRVEAFHRYGTTVNCIVEEVDARSVYPYDEFVLATGDFVYMSPF 225
HS2GPB_33       TRTSRGWHTTDLKYNPSRVEAFHRYGTTVNCIVEEVDARSVYPYDEFVLATGDFVYMSPF 222
HS2GPB_38       TRTSRGWHTTDLKYNPSRVEAFHRYGTTVNCIVEEVDARSVYPYDEFVLATGDFVYMSPF 222
HS2GPB_39       TRTSRGWHTTDLKYNPSRVEAFHRYGTTVNCIVEEVDARSVYPYDEFVLATGDFVYMSPF 222
                *****************************.**************:***************

HS1GPB_3        YGYREGSHTEHTSYAADRFKQVDGFYARDLTTKARATAPTTRNLLTTPKFTVAWDWVPKR 360
HS1GPB_5        YGYREGSHTEHTSYAADRFKQVDGFYARDLTTKARATAPTTRNLLTTPKFTVAWDWVPKR 360
HS1GPB_7        YGYREGSHTEHTSYAADRFKQVDGFYARDLTTKARATAPTTRNLLTTPKFTVAWDWVPKR 290
HS1GPB_9        YGYREGSHTEHTSYAADRFKQVDGFYARDLTTKARATAPTTRNLLTTPKFTVAWDWVPKR 360
HS1GPB_4        YGYREGSHTEHTSYAADRFKQVDGFYARDLTTKARATAPTTRNLLTTPKFTVAWDWVPKR 290
HS1GPB_1        YGYREGSHTEHTSYAADRFKQVDGFYARDLTTKARATAPTTRNLLTTPKFTVAWDWVPKR 290
HS1GPB_2        YGYREGSHTEHTSYAADRFKQVDGFYARDLTTKARATAPTTRNLLTTPKFTVAWDWVPKR 290
HS1GPB_12       YGYREGSHTEHTSYAADRFKQVDGFYARDLTTKARATAPTTRNLLTTPKFTVAWDWVPKR 360
HS1GPB_13       YGYREGSHTEHTSYAADRFKQVDGFYARDLTTKARATAPTTRNLLTTPKFTVAWDWVPKR 360
HS1GPB_16       YGYREGSHTEHTSYAADRFKQVDGFYARDLTTKARATAPTTRNLLTTPKFTVAWDWVPKR 360
HS1GPB_8        YGYREGSHTEHTSYAADRFKQVDGFYARDLTTKARATAPTTRNLLTTPKFTVAWDWVPKR 290
HS1GPB_6        YGYREGSHTEHTSYAADRFKQVDGFYARDLTTKARATAPTTRNLLTTPKFTVAWDWVPKR 360
HS1GPB_11       YGYREGSHTEHTSYAADRFKQVDGFYARDLTTKARATAPTTRNLLTTPKFTVAWDWVPKR 360
HS1GPB_15       YGYREGSHTEHTSYAADRFKQVDGFYARDLTTKARATAPTTRNLLTTPKFTVAWDWVPKR 360
HS1GPB_19       YGYREGSHTEHTSYAADRFKQVDGFYARDLTTKARATAPTTRNLLTTPKFTVAWDWVPKR 290
HS1GPB_14       YGYREGSHTEHTSYAADRFKQVDGFYARDLTTKARATAPTTRNLLTTPKFTVAWDWVPKR 290
HS1GPB_17       YGYREGSHTEHTSYAADRFKQVDGFYARDLTTKARATAPTTRNLLTTPKFTVAWDWVPKR 360
HS1GPB_20       YGYREGSHTEHTSYAADRFKQVDGFYARDLTTKARATAPTTRNLLTTPKFTVAWDWVPKR 290
HS1GPB_10       YGYREGSHTEHTSYAADRFKQVDGFYARDLTTKARATAPTTRNLLTTPKFTVAWDWVPKR 360
HS1GPB_18       YGYREGSHTEHTSYAADRFKQVDGFYARDLTTKARATAPTTRNLLTTPKFTVAWDWVPKR 360
HS2GPB_21       YGYREGSHTEHTSYAADRFKQVDGFYARDLTTKARATSPTTRNLLTTPKFTVAWDWVPKR 355
HS2GPB_22       YGYREGSHTEHTSYAADRFKQVDGFYARDLTTKARATSPTTRNLLTTPKFTVAWDWVPKR 355
HS2GPB_27       YGYREGSHTEHTSYAADRFKQVDGFYARDLTTKARATSPTTRNLLTTPKFTVAWDWVPKR 355
HS2GPB_30       YGYREGSHTEHTSYAADRFKQVDGFYARDLTTKARATSPTTRNLLTTPKFTVAWDWVPKR 355
HS2GPB_31       YGYREGSHTEHTSYAADRFKQVDGFYARDLTTKARATSPTTRNLLTTPKFTVAWDWVPKR 355
HS2GPB_34       YGYREGSHTEHTSYAADRFKQVDGFYARDLTTKARATSPTTRNLLTTPKFTVAWDWVPKR 355
HS2GPB_28       YGYREGSHTEHTSYAADRFKQVDGFYARDLTTKARATSPTTRNLLTTPKFTVAWDWVPKR 355
HS2GPB_24       YGYREGSHTEHTSYAADRFKQVDGFYARDLTTKARATSPTTRNLLTTPKFTVAWDWVPKR 355
HS2GPB_26       YGYREGSHTEHTSYAADRFKQVDGFYARDLTTKAQATSPTTRNLLTTPKFTVAWDWVPKR 355
HS2GPB_35       YGYREGSHTEHTSYAADRFKQVDGFYARDLTTKARATSPTTRNLLTTPKFTVAWDWVPKR 352
HS2GPB_37       YGYREGSHTEHTSYAADRFKQVDGFYARDLTTKARATSPTTRNLLTTPKFTVAWDWVPKR 352
HS2GPB_36       YGYREGSHTEHTSYAADRFKQVDGFYARDLTTKARATSPTTRNLLTTPKFTVAWDWVPKR 352
HS2GPB_25       YGYREGSHTEHTSYAADRFKQVDGFYARDLTTKARATSPTTRNLLTTPKFTVAWDWVPKR 285
HS2GPB_23       YGYREGSHTEHTSYAADRFKQVDGFYARDLTTKARATSPTTRNLLTTPKFTVAWDWVPKR 285
HS2GPB_29       YGYREGSHTEHTSYAADRFKQVDGFYARDLTTKARATSPTTRNLLTTPKFTVAWDWVPKR 285
HS2GPB_32       YGYREGSHTEHTSYAADRFKQVDGFYARDLTTKARATSPTTRNLLTTPKFTVAWDWVPKR 285
HS2GPB_33       YGYREGSHTEHTSYAADRFKQVDGFYARDLTTKARATSPTTRNLLTTPKFTVAWDWVPKR 282
HS2GPB_38       YGYREGSHTEHTSYAADRFKQVDGFYARDLTTKARATSPTTRNLLTTPKFTVAWDWVPKR 282
HS2GPB_39       YGYREGSHTEHTSYAADRFKQVDGFYARDLTTKARATSPTTRNLLTTPKFTVAWDWVPKR 282
                **********************************:**:**********************

HS1GPB_3        PSVCTMTKWQEVDEMLRSEYGGSFRFSSDAISTTFTTNLTEYPLSRVDLGDCIGKDARDA 420
HS1GPB_5        PSVCTMTKWQEVDEMLRSEYGGSFRFSSDAISTTFTTNLTEYPLSRVDLGDCIGKDARDA 420
HS1GPB_7        PSVCTMTKWQEVDEMLRSEYGGSFRFSSDAISTTFTTNLTEYPLSRVDLGDCIGKDARDA 350
HS1GPB_9        PSVCTMTKWQEVDEMLRSEYGGSFRFSSDAISTTFTTNLTEYPLSRVDLGDCIGKDARDA 420
HS1GPB_4        PSVCTMTKWQEVDEMLRSEYGGSFRFSSDAISTTFTTNLTEYPLSRVDLGDCIGKDARDA 350
HS1GPB_1        PSVCTMTKWQEVDEMLRSEYGGSFRFSSDAISTTFTTNLTEYPLSRVDLGDCIGKDARDA 350
HS1GPB_2        PSVCTMTKWQEVDEMLRSEYGGSFRFSSDAISTTFTTNLTEYPLSRVDLGDCIGKDARDA 350
HS1GPB_12       PSVCTMTKWQEVDEMLRSEYGGSFRFSSDAISTTFTTNLTEYPLSRVDLGDCIGKDARDA 420
HS1GPB_13       PSVCTMTKWQEVDEMLRSEYGGSFRFSSDAISTTFTTNLTEYPLSRVDLGDCIGKDARDA 420
HS1GPB_16       PSVCTMTKWQEVDEMLRSEYGGSFRFSSDAISTTFTTNLTEYPLSRVDLGDCIGKDARDA 420
HS1GPB_8        PSVCTMTKWQEVDEMLRSEYGGSFRFSSDAISTTFTTNLTEYPLSRVDLGDCIGKDARDA 350
HS1GPB_6        PSVCTMTKWQEVDEMLRSEYGGSFRFSSDAISTTFTTNLTEYPLSRVDLGDCIGKDARDA 420
HS1GPB_11       PSVCTMTKWQEVDEMLRSEYGGSFRFSSDAISTTFTTNLTEYPLSRVDLGDCIGKDARDA 420
HS1GPB_15       PSVCTMTKWQEVDEMLRSEYGGSFRFSSDAISTTFTTNLTEYPLSRVDLGDCIGKDARDA 420
HS1GPB_19       PSVCTMTKWQEVDEMLRSEYGGSFRFSSDAISTTFTTNLTEYPLSRVDLGDCIGKDARDA 350
HS1GPB_14       PSVCTMTKWQEVDEMLRSEYGGSFRFSSDAISTTFTTNLTEYPLSRVDLGDCIGKDARDA 350
HS1GPB_17       PSVCTMTKWQEVDEMLRSEYGGSFRFSSDAISTTFTTNLTEYPLSRVDLGDCIGKDARDA 420
HS1GPB_20       PSVCTMTKWQEVDEMLRSEYGGSFRFSSDAISTTFTTNLTEYPLSRVDLGDCIGKDARDA 350
HS1GPB_10       PSVCTMTKWQEVDEMLRSEYGGSFRFSSDAISTTFTTNLTEYPLSRVDLGDCIGKDARDA 420
HS1GPB_18       PSVCTMTKWQEVDEMLRSEYGGSFRFSSDAISTTFTTNLTEYPLSRVDLGDCIGKDARDA 420
HS2GPB_21       PAVCTMTKWQEVDEMLRAEYGGSFRFSSDAISTTFTTNLTEYSLSRVDLGDCIGRDAREA 415
HS2GPB_22       PAVCTMTKWQEVDEMLRAEYGGSFRFSSDAISTTFTTNLTEYSLSRVDLGDCIGRDAREA 415
HS2GPB_27       PAVCTMTKWQEVDEMLRAEYGGSFRFSSDAISTTFTTNLTQYSLSRVDLGDCIGRDAREA 415
HS2GPB_30       PAVCTMTKWQEVDEMLRAEYGGSFRFSSDAISTTFTTNLTQYSLSRVDLGDCIGRDAREA 415
HS2GPB_31       PAVCTMTKWQEVDEMLRAEYGGSFRFSSDAISTTFTTNLTQYSLSRVDLGDCIGRDAREA 415
HS2GPB_34       PAVCTMTKWQEVDEMLRAEYGGSFRFSSDAISTTFTTNLTQYSLSRVDLGDCIGRDAREA 415
HS2GPB_28       PAVCTMTKWQEVDEMLRAEYGGSFRFSSDAISTTFTTNLTQYSLSRVDLGDCIGRDAREA 415
HS2GPB_24       PVVCTMTKWQEVDEMLRAEYGGSFRFSSDAISTTFTTNLTQYSLSRVDLGDCIGRDAREA 415
HS2GPB_26       PAVCTMTKWQEVDEMLRAEYGGSFRFSSDAISTTFTTNLTQYSLSRVDLGDCIGRDAREA 415
HS2GPB_35       PAVCTMTKWQEVDEMLRAEYGGSFRFSSDAISTTFTTNLTEYSLSRVDLGDCIGRDAREA 412
HS2GPB_37       PAVCTMTKWQEVDEMLRAEYGGSFRFSSDAISTTFTTNLTEYSLSRVDLGDCIGRDAREA 412
HS2GPB_36       PAVCTMTKWQEVDEMLRAEYGGSFRFSSDAISTTFTTNLTEYSLSRVDLGDCIGRDAREA 412
HS2GPB_25       PAVCTMTKWQEVDEMLRAEYGGSFRFSSDAISTTFTTNLTQYSLSRVDLGDCIGRDAREA 345
HS2GPB_23       PAVCTMTKWQEVDEMLRAEYGGSFRFSSDAISTTFTTNLTQYSLSRVDLGDCIGRDAREA 345
HS2GPB_29       PAVCTMTKWQEVDEMLRAEYGGSFRFSSDAISTTFTTNLTQYSLSRVDLGDCIGRDAREA 345
HS2GPB_32       PAVCTMTKWQEVDEMLRAEHGGSFRFSSDAISTTFTTNLTQYSLSRVDLGDCIGRDAREA 345
HS2GPB_33       PAVCTMTKWQEVDEMLRAEYGGSFRFSSDAISTTFTTNLTEYSLSRVDLGDCIGRDAREA 342
HS2GPB_38       PAVCTMTKWQEVDEMLRAEYGGSFRFSSDAISTTFTTNLTEYSLSRVDLGDCIGRDAREA 342
HS2GPB_39       PAVCTMTKWQEVDEMLRAEYGGSFRFSSDAISTTFTTNLTEYSLSRVDLGDCIGRDAREA 342
                * ***************:*:********************:*.***********:***:*

HS1GPB_3        MDRIFARRYNATHIKVGQPQYYLANGGFLIAYQPLLSNTLAELYVREHLREQSRKPPNPT 480
HS1GPB_5        MDRIFARRYNATHIKVGQPQYYLANGGFLIAYQPLLSNTLAELYVREHLREQSRKPPNPT 480
HS1GPB_7        MDRIFARRYNATHIKVGQPQYYLANGGFLIAYQPLLSNALAELYVREHLREQSRKPPNPT 410
HS1GPB_9        MDRIFARRYNATHIKVGQPQYYLANGGFLIAYQPLLSNTLAELYVREHLREQSRKPPNPT 480
HS1GPB_4        MDRIFARRYNATHIKVGQPQYYLANGGFLIAYQPLLSNTLAELYVREHLREQSRKPPNPT 410
HS1GPB_1        MDRIFARRYNATHIKVGQPQYYLANGGFLIAYQPLLSNTLAELYVREHLREQSRKPPNPT 410
HS1GPB_2        MDRIFARRYNATHIKVGQPQYYLANGGFLIAYQPLLSNTLAELYVREHLREQSRKPPNPT 410
HS1GPB_12       MDRIFARRYNATHIKVGQPQYYLANGGFLIAYQPLLSNTLAELYVREHLREQSRKPPNPT 480
HS1GPB_13       MDRIFARRYNATHIKVGQPQYYLANGGFLIAYQPLLSNTLAELYVREHLREQSRKPPNPT 480
HS1GPB_16       MDRIFARRYNATHIKVGQPQYYLANGGFLIAYQPLLSNTLAELYVREHLREQSRKPPNPT 480
HS1GPB_8        MDRIFARRYNATHIKVGQPQYYLANGGFLIAYQPLLSNTLAELYVREHLREQSRKPPNPT 410
HS1GPB_6        MDRIFARRYNATHIKVGQPQYYLANGGFLIAYQPLLSNTLAELYVREHLREQSRKPPNPT 480
HS1GPB_11       MDRIFARRYNATHIKVGQPQYYLANGGFLIAYQPLLSNTLAELYVREHLREQSRKPPNPT 480
HS1GPB_15       MDRIFARRYNATHIKVGQPQYYLANGGFLIAYQPLLSNTLAELYVREHLREQSRKPPNPT 480
HS1GPB_19       MDRIFARRYNATHIKVGQPQYYLANGGFLIAYQPLLSNTLAELYVREHLREQSRKPPNPT 410
HS1GPB_14       MDRIFARRYNATHIKVGQPQYYLANGGFLIAYQPLLSNTLAELYVREHLREQSRKPPNPT 410
HS1GPB_17       MDRIFARRYNATHIKVGQPQYYLANGGFLIAYQPLLSNTLAELYVREHLREQSRKPPNPT 480
HS1GPB_20       MDRIFARRYNATHIKVGQPQYYLANGGFLIAYQPLLSNTLAELYVREHLREQSRKPPNPT 410
HS1GPB_10       MDRIFARRYNATHIKVGQPQYYLANGGFLIAYQPLLSNTLAELYVREHLREQSRKPPNPT 480
HS1GPB_18       MDRIFARRYNATHIKVGQPQYYLANGGFLIAYQPLLSNTLAELYVREHLREQSRKPPNPT 480
HS2GPB_21       IDRMFARKYNATHIKVGQPQYYLATGGFLIAYQPLLSNTLAELYVREYMREQDRKPRNAT 475
HS2GPB_22       IDRMFARKYNATHIKVGQPQYYLATGGFLIAYQPLLSNTLAELYVREYMREQDRKPRNAT 475
HS2GPB_27       IDRMFARKYNATHIKVGQPQYYLATGGFLIAYQPLLSNTLAELYVREYMREQDRKPRNAT 475
HS2GPB_30       IDRMFARKYNATHIKVGQPQYYLATGGFLIAYQPLLSNTLAELYVREYMREQDRKPRNAT 475
HS2GPB_31       IDRMFARKYNATHIKVGQPQYYLATGGFLIAYQPLLSNTLAELYVREYMREQDRKPRNAT 475
HS2GPB_34       IDRMFARKYNATHIKVGQPQYYLATGGFLIAYQPLLSNTLAELYVREYMREQDRKPRNAT 475
HS2GPB_28       IDRMFARKYNATHIKVGQPQYYLATGGFLIAYQPLLSNTLAELYVREYMREQDRKPRNAT 475
HS2GPB_24       IDRMFARKYNATHIKVGQPQYYLATGGFLIAYQPLLSNTLAELYVREYMREQDRKPRNAT 475
HS2GPB_26       IDRMFARKYNATHIKVGQPQYYLATGGFLIAYQPLLSNTLAELYVREYMREQDRKPRNAT 475
HS2GPB_35       IDRMFARKYNATHIKVGQPQYYLATGGFLIAYQPLLSNTLAELYVREYMREQDRKPRNAT 472
HS2GPB_37       IDRMFARKYNATHIKVGQPQYYLATGGFLIAYQPLLSNTLAELYVREYMREQDRKPRNAT 472
HS2GPB_36       IDRMFARKYNATHIKVGQPQYYLATGGFLIAYQPLLSNTLAELYVREYMREQDRKPRNAT 472
HS2GPB_25       IDRMFARKYNATHIKVGQPQYYLATGGFLIAYQPLLSNTLAELYVREYMREQDRKPRNAT 405
HS2GPB_23       IDRMFARKYNATHIKVGQPQYYLATGGFLIAYQPLLSNTLAELYVREYMREQDRKPRNAT 405
HS2GPB_29       IDRMFARKYNATHIKVGQPQYYLATGGFLIAYQPLLSNTLAELYVREYMREQDRKPRNAT 405
HS2GPB_32       IDRMFARKYNATHIKVGQPQYYLATGGFLIAYQPLLSNTLAELYVREYMREQDRKPRNAT 405
HS2GPB_33       IDRMFARKYNATHIKVGQPQYYLATGGFLIAYQPLLSNTLAELYVREYMREQDRKPRNAT 402
HS2GPB_38       IDRMFARKYNATHIKVGQPQYYLATGGFLIAYQPLLSNTLAELYVREYMREQDRKPRNAT 402
HS2GPB_39       IDRMFARKYNATHIKVGQPQYYLATGGFLIAYQPLLSNTLAELYVREYMREQDRKPRNAT 402
                :**:***:****************.*************:********::***.*** *.*

HS1GPB_3        PPPP--GASANASVERIKTTSSIEFARLQFTYNHIQRHVNDMLGRVAIAWCELQNHELTL 538
HS1GPB_5        PPPP--GASANASVERIKTTSSIEFARLQFTYNHIQRHVNDMLGRVAIAWCELQNHELTL 538
HS1GPB_7        PPPP--GASANASVERIKTTSSIEFARLQFTYNHIQRHVNDMLGRVAIAWCELQNHELTL 468
HS1GPB_9        PPPP--GASANASVERIKTTSSIEFARLQFTYNHIQRHVNDMLGRVAIAWCELQNHELTL 538
HS1GPB_4        PPPP--GASANASVERIKTTSSIEFARLQFTYNHIQRHVNDMLGRVAIAWCELQNHELTL 468
HS1GPB_1        PPPP--GASANASVERIKTTSSIEFARLQFTYNHIQRHVNDMLGRVAIAWCELQNHELTL 468
HS1GPB_2        PPPP--GASANASVERIKTTSSIEFARLQFTYNHIQRHVNDMLGRVAIAWCELQNHELTL 468
HS1GPB_12       PPPP--GASANASVERIKTTSSIEFARLQFTYNHIQRHVNDMLGRVAIAWCELQNHELTL 538
HS1GPB_13       PPPP--GASANASVERIKTTSSIEFARLQFTYNHIQRHVNDMLGRVAIAWCELQNHELTL 538
HS1GPB_16       PPPP--GASANASVERIKTTSSIEFARLQFTYNHIQRHVNDMLGRVAIAWCELQNHELTL 538
HS1GPB_8        PPPP--GASANASVERIKTTSSIEFARLQFTYNHIQRHVNDMLGRVAIAWCELQNHELTL 468
HS1GPB_6        PPPP--GASANASVERIKTTSSIEFARLQFTYNHIQRHVNDMLGRVAIAWCELQNHELTL 538
HS1GPB_11       PPPP--GASANASVERIKTTSSIEFARLQFTYNHIQRHVNDMLGRVAIAWCELQNHELTL 538
HS1GPB_15       PPPP--GASANASVERIKTTSSIEFARLQFTYNHIQRHVNDMLGRVAIAWCELQNHELTL 538
HS1GPB_19       PPPP--GASANASVERIKTTSSIEFARLQFTYNHIQRHVNDMLGRVAIAWCELQNHELTL 468
HS1GPB_14       PPPP--GASANASVERIKTTSSIEFARLQFTYNHIQRHVNDMLGRVAIAWCELQNHELTL 468
HS1GPB_17       PPPP--GASANASVERIKTTSSIEFARLQFTYNHIQRHVNDMLGRVAIAWCELQNHELTL 538
HS1GPB_20       PPPP--GASANASVERIKTTSSIEFARLQFTYNHIQRHVNDMLGRVAIAWCELQNHELTL 468
HS1GPB_10       PPPP--GASANASVERIKTTSSIEFARLQFTYNHIQRHVNDMLGRVAIAWCELQNHELTL 538
HS1GPB_18       PPPP--GASANASVERIKTTSSIEFARLQFTYNHIQRHVNDMLGRVAIAWCELQNHELTL 538
HS2GPB_21       PAPLREAPSANASVERIKTTSSIEFARLQFTYNHIQRHVNDMLGRIAVAWCELQNHELTL 535
HS2GPB_22       PAPLREAPSANASVERIKTTSSIEFARLQFTYNHIQRHVNDMLGRIAVAWCELQNHELTL 535
HS2GPB_27       PAPLREAPSANASVERIKTTSSIEFARLQFTYNHIQRHVNDMLGRIAVAWCELQNHELTL 535
HS2GPB_30       PAPLREAPSANASVERIKTTSSIEFARLQFTYNHIQRHVNDMLGRIAVAWCELQNHELTL 535
HS2GPB_31       PAPLREAPSANASVERIKTTSSIEFARLQFTYNHIQRHVNDMLGRIAVAWCELQNHELTL 535
HS2GPB_34       PAPLREAPSANASVERIKTTSSIEFARLQFTYNHIQRHVNDMLGRIAVAWCELQNHELTL 535
HS2GPB_28       PAPLREAPSANASVERIKTTSSIEFARLQFTYNHIQRHVNDMLGRIAVAWCELQNHELTL 535
HS2GPB_24       PAPLREAPSANASVERIKTTSSIEFARLQFTYNHIQRHVNDMLGRIAVAWCELQNHELTL 535
HS2GPB_26       PAPLREAPSANASVERIKTTSSIEFARLQFTYNHIQRHVNDMLGRIAVAWCELQNHELTL 535
HS2GPB_35       PAPLREAPSANASVERIKTTSSIEFARLQFTYNHIQRHVNDMLGRIAVAWCELQNHELTL 532
HS2GPB_37       PAPLREAPSANASVERIKTTSSIEFARLQFTYNHIQRHVNDMLGRIAVAWCELQNHELTL 532
HS2GPB_36       PAPLREAPSANASVERIKTTSSIEFARLQFTYNHIQRHVNDMLGRIAVAWCELQNHELTL 532
HS2GPB_25       PAPLREAPSANASVERIKTTSSIEFARLQFTYNHIQRHVNDMLGRIAVAWCELQNHELTL 465
HS2GPB_23       PAPLREAPSANASVERIKTTSSIEFARLQFTYNHIQRHVNDMLGRIAVAWCELQNHELTL 465
HS2GPB_29       PAPLREAPSANASVERIKTTSSIEFARLQFTYNHIQRHVNDMLGRIAVAWCELQNHELTL 465
HS2GPB_32       PAPLREAPSANASVERIKTTSSIEFARLQFTYNHIQRHVNDMLGRIAVAWCELQNHELTL 465
HS2GPB_33       PAPLREAPSANASVERIKTTSSIEFARLQFTYNHIQRHVNDMLGRIAVAWCELQNHELTL 462
HS2GPB_38       PAPLREAPSANASVERIKTTSSIEFARLQFTYNHIQRHVNDMLGRIAVAWCELQNHELTL 462
HS2GPB_39       PAPLREAPSANASVERIKTTSSIEFARLQFTYNHIQRHVNDMLGRIAVAWCELQNHELTL 462
                *.*   ..*************************************:*:************

HS1GPB_3        WNEARKLNPNAIASATVGRRVSARMLGDVMAVSTCVPVAADNVIVQNSMRISSRPGACYS 598
HS1GPB_5        WNEARKLNPNAIASATVGRRVSARMLGDVMAVSTCVPVAADNVIVQNSMRISSRPGACYS 598
HS1GPB_7        WNEARKLNPNAIASATMGRRVSARMLGDVMAVSTCVPVAADNVIVQNSMRISSRPGACYS 528
HS1GPB_9        WNEARKLNPNAIASATMGRRVSARMLGDVMAVSMCVPVAADNVIVQNSMRISSRPGACYS 598
HS1GPB_4        WNEARKLNPNAIASATVGRRVSARMLGDVMAVSTCVPVAADNVIVQNSMRISSRPGACYS 528
HS1GPB_1        WNEARKLNPNAIASATVGRRVSARMLGDVMAVSTCVPVAADNVIVQNSMRISSRPGACYS 528
HS1GPB_2        WNEARKLNPNAIASATVGRRVSARMLGDVMAVSTCVPVAADNVIVQNSMRISSRPGACYS 528
HS1GPB_12       WNEARKLNPNAIASATVGRRVSARMLGDVMAVSTCVPVAADNVIVQNSMRISSRPGACYS 598
HS1GPB_13       WNEARKLNPNAIASATVGRRVSARMLGDVMAVSTCVPVAADNVIVQNSMRISSRPGACYS 598
HS1GPB_16       WNEARKLNPNAIASATVGRRVSARMLGDVMAVSTCVPVAADNVIVQNSMRISSRPGACYS 598
HS1GPB_8        WNEARKLNPNAIASATVGRRVSARMLGDVMAVSTCVPVAADNVIVQNSMRISSRPGACYS 528
HS1GPB_6        WNEARKLNPNAIASATVGRRVSARMLGDVMAVSTCVPVAADNVIVQNSMRISSRPGACYS 598
HS1GPB_11       WNEARKLNPNAIASATVGRRVSARMLGDVMAVSTCVPVAADNVIVQNSMRISSRPGACYS 598
HS1GPB_15       WNEARKLNPNAIASATVGRRVSARMLGDVMAVSTCVPVAADNVIVQNSMRISSRPGACYS 598
HS1GPB_19       WNEARKLNPNAIASATVGRRVSARMLGDVMAVSTCVPVAADNVIVQNSMRISSRPGACYS 528
HS1GPB_14       WNEARKLNPNAIASATVGRRVSARMLGDVMAISTCVPVAADNVIVQNSMRISSRPGACYS 528
HS1GPB_17       WNEARKLNPNAIASATVGRRVSARMLGDVMAVSTCVPVAADNVIVQNSMRISSRPGACYS 598
HS1GPB_20       WNEARKLNPNAIASATVGRRVSARMLGDVMAVSTCVPVAADNVIVQNSMRISSRPGACYS 528
HS1GPB_10       WNEARKLNPNAIASATVGRRVSARMLGDVMAVSTCVPVAADNVIVQNSMRISSRPGACYS 598
HS1GPB_18       WNEARKLNPNAIASATVGRRVSARMLGDVMAVSTCVPVAADNVIVQNSMRISSRPGACYS 598
HS2GPB_21       WNEARKLNPNAIASATVGRRVSARMLGDVMAVSTCVPVAPDNVIVQNSMRVSSRPGTCYS 595
HS2GPB_22       WNEARKLNPNAIASATVGRRVSARMLGDVMAVSTCVPVAPDNVIVQNSMRVSSRPGTCYS 595
HS2GPB_27       WNEARKLNPNAIASATVGRRVSARMLGDVMAVSTCVPVAPDNVIVQNSMRVSSRPGTCYS 595
HS2GPB_30       WNEARKLNPNAIASATVGRRVSARMLGDVMAVSTCVPVAPDNVIVQNSMRVSSRPGTCYS 595
HS2GPB_31       WNEARKLNPNAIASATVGRRVSARMLGDVMAVSTCVPVAPDNVIVQNSMRVSSRPGTCYS 595
HS2GPB_34       WNEARKLNPNTIASATVGRRVSARMLGDVMAVSTCVPVAPDNVIVQNSMRVSSRPGTCYS 595
HS2GPB_28       WNEARKLNPNAIASATVGRRVSARMLGDVMAVSTCVPVAPDNVIVQNSMRVSSRPGTCYS 595
HS2GPB_24       WNEARKLNPNAIASATVGRRVSARMLGDVMAVSTCVPVAPDNVIVQNSMRVSSRPGTCYS 595
HS2GPB_26       WNEARKLNPNAIASATVGRRVSARMLGDVMAVSTCVPVAPDNVIVQNSMRVSSRPGTCYS 595
HS2GPB_35       WNEARKLNPNAIASATVGRRVSARMLGDVMAVSTCVPVAPDNVIVQNSMRVSSRPGTCYS 592
HS2GPB_37       WNEARKLNPNAIASATVGRRVSARMLGDVMAVSTCVPVAPDNVIVQNSMRVSSRPGTCYS 592
HS2GPB_36       WNEARKLNPNAIASATVGRRVSARMLGDVMAVSTCVPVAPDNVIVQNSMRVSSRPGTCYS 592
HS2GPB_25       WNEARKLNPNAIASATVGRRVSARMLGDVMAVSTCVPVAPDNVIVQNSMRVSSRPGTCYS 525
HS2GPB_23       WNEARKLNPNAIASATVGRRVSARMLGDVMAVSTCVPVAPDNVIVQNSMRVSSRPGTCYS 525
HS2GPB_29       WNEARKLNPNAIASATVGRRVSARMLGDVMAVSTCVPVAPDNVIVQNSMRVSSRPGTCYS 525
HS2GPB_32       WNEARKLNPNAIASATVGRRVSARMLGDVMAVSTCVPVAPDNVIVQNSMRVSSRPGTCYS 525
HS2GPB_33       WNEARKLNPNAIASATVGRRVSARMLGDVMAVSTCVPVAPDNVIVQNSMRVSSRPGTCYS 522
HS2GPB_38       WNEARKLNPNAIASATVGRRVSARMLGDVMAVSTCVPVAPDNVIVQNSMRVSSRPGTCYS 522
HS2GPB_39       WNEARKLNPNAIASATVGRRVSARMLGDVMAVSTCVPVAPDNVIVQNSMRVSSRPGTCYS 522
                **********:*****:**************:* *****.**********:*****:***

HS1GPB_3        RPLVSFRYEDQGPLVEGQLGENNELRLTRDAIEPCTVGHRRYFTFGGGYVYFEEYAYSHQ 658
HS1GPB_5        RPLVSFRYEDQGPLVEGQLGENNELRLTRDAIEPCTVGHRRYFTFGGGYVYFEEYAYSHQ 658
HS1GPB_7        RPLVSFRYEDQGPLVEGQLGENNELRLTRDAIEPCTVGHRRYFTFGGGYVYFEEYAYSHQ 588
HS1GPB_9        RPLVSFRYEDQGPLVEGQLGENNELRLTRDAIEPCTVGHRRYFTFGGGYVYFEEYAYSHQ 658
HS1GPB_4        RPLVSFRYEDQGPLVEGQLGENNELRLTRDAIEPCTVGHRRYFTFGGGYVYFEEYAYSHQ 588
HS1GPB_1        RPLVSFRYEDQGPLVEGQLGENNELRLTRDAIEPCTVGHRRYFTFGGGYVYFEEYAYSHQ 588
HS1GPB_2        RPLVSFRYEDQGPLVEGQLGENNELRLTRDAIEPCTVGHRRYFTFGGGYVYFEEYAYSHQ 588
HS1GPB_12       RPLVSFRYEDQGPLVEGQLGENNELRLTRDAIEPCTVGHRRYFTFGGGYVYFEEYAYSHQ 658
HS1GPB_13       RPLVSFRYEDQGPLVEGQLGENNELRLTRDAIEPCTVGHRRYFTFGGGYVYFEEYAYSHQ 658
HS1GPB_16       RPLVSFRYEDQGPLVEGQLGENNELRLTRDAIEPCTVGHRRYFTFGGGYVYFEEYAYSHQ 658
HS1GPB_8        RPLVSFRYEDQGPLVEGQLGENNELRLTRDAIEPCTVGHRRYFTFGGGYVYFEEYAYSHQ 588
HS1GPB_6        RPLVSFRYEDQGPLVEGQLGENNELRLTRDAIEPCTVGHRRYFTFGGGYVYFEEYAYSHQ 658
HS1GPB_11       RPLVSFRYEDQGPLVEGQLGENNELRLTRDAIEPCTVGHRRYFTFGGGYVYFEEYAYSHQ 658
HS1GPB_15       RPLVSFRYEDQGPLVEGQLGENNELRLTRDAIEPCTVGHRRYFTFGGGYVYFEEYAYSHQ 658
HS1GPB_19       RPLVSFRYEDQGPLVEGQLGENNELRLTRDAIEPCTVGHRRYFTFGGGYVYFEESAYSHQ 588
HS1GPB_14       RPLVSFRYEDQGPLVEGQLGENNELRLTRDAIEPCTVGHRRYFTFGGGYVYFEEYAYSHQ 588
HS1GPB_17       RPLVSFRYEDQGPLVEGQLGENNELRLTRDAIEPCTVGHRRYFTFGGGYVYFEEYAYSHQ 658
HS1GPB_20       RPLVSFRYEDQGPLVEGQLGENNELRLTRDAIEPCTVGHRRYFTFGGGYVYFEEYAYSHQ 588
HS1GPB_10       RPLVSFRYEDQGPLVEGQLGENNELRLTRDAIEPCTVGHRRYFTFGGGYVYFEEYAYSHQ 658
HS1GPB_18       RPLVSFRYEDQGPLVEGQLGENNELRLTRDAIEPCTVGHRRYFTFGGGYVYFEEYAYSHQ 658
HS2GPB_21       RPLVSFRYEDQGPLIEGQLGENNELRLTRDALEPCTVGHRRYFIFGGGYVYFEEYAYSHQ 655
HS2GPB_22       RPLVSFRYEDQGPLIEGQLGENNELRLTRDALEPCTVGHRRYFIFGGGYVYFEEYAYSHQ 655
HS2GPB_27       RPLVSFRYEDQGPLIEGQLGENNELRLTRDALEPCTVGHRRYFIFGGGYVYFEEYAYSHQ 655
HS2GPB_30       RPLVSFRYEDQGPLIEGQLGENNELRLTRDALEPCTVGHRRYFIFGGGYVYFEEYAYSHQ 655
HS2GPB_31       RPLVSFRYEDQGPLIEGQLGENNELRLTRDALEPCTVGHRRYFIFGGGYVYFEEYAYSHQ 655
HS2GPB_34       RPLVSFRYEDQGPLIEGQLGENNELRLTRDALEPCTVGHRRYFIFGGGYVYFEEYAYSHQ 655
HS2GPB_28       RPLVSFRYEDQGPLIEGQLGENNELRLTRDALEPCTVGHRRYFIFGGGYVYFEEYAYSHQ 655
HS2GPB_24       RPLVSFRYEDQGPLIEGQLGENNELRLTRDALEPCTVGHRRYFIFGGGYVYFEEYAYSHQ 655
HS2GPB_26       RPLVSFRYEDQGPLIEGQLGENNELRLTRDALEPCTVGHRRYFIFGGGYVYFEEYAYSHQ 655
HS2GPB_35       RPLVSFRYEDQGPLIEGQLGENNELRLTRDALEPCTVGHRRYFIFGGGYVYFEEYAYSHQ 652
HS2GPB_37       RPLVSFRYEDQGPLIEGQLGENNELRLTRDALEPCTVGHRRYFIFGGGYVYFEEYAYSHQ 652
HS2GPB_36       RPLVSFRYEDQGPLIEGQLGENNELRLTRDALEPCTVGHRRYFIFGGGYVYFEEYAYSHQ 652
HS2GPB_25       RPLVSFRYEDQGPLIEGQLGENNELRLTRDALEPCTVGHRRYFIFGGGYVYFEEYAYSHQ 585
HS2GPB_23       RPLVSFRYEDQGPLIEGQLGENNELRLTRDALEPCTVGHRRYFIFGGGYVYFEEYAYSHQ 585
HS2GPB_29       RPLVSFRYEDQGPLIEGQLGENNELRLTRDALEPCTVGHRRYFIFGGGYVYFEEYAYSHQ 585
HS2GPB_32       RPLVSFRYEDQGPLIEGQLGENNELRLTRDALEPCTVGHRRYFIFGGGYVYFEEYAYSHQ 585
HS2GPB_33       RPLVSFRYEDQGPLIEGQLGENNELRLTRDALEPCTVGHRRYFIFGGGYVYFEEYAYSHQ 582
HS2GPB_38       RPLVSFRYEDQGPLIEGQLGENNELRLTRDALEPCTVGHRRYFIFGGGYVYFEEYAYSHQ 582
HS2GPB_39       RPLVSFRYEDQGPLIEGQLGENNELRLTRDALEPCTVGHRRYFIFGGGYVYFEEYAYSHQ 582
                **************:****************:*********** ********** *****

HS1GPB_3        LSRADITTVSTFIDLNITMLEDHEFVPLEVYTRHEIKDSGLLDYTEVQRRNQLHDLRFAD 718
HS1GPB_5        LSRADITTVSTFIDLNITMLEDHEFVPLEVYTRHEIKDSGLLDYTEVQRRNQLHDLRFAD 718
HS1GPB_7        LSRADITTVNTFIDLNITMLEDHEFVPLEVYTRHEIKDSGLLDYTEVQRRNQLHDLRFAD 648
HS1GPB_9        LSRADITTVNTFIDLNITMLEDHEFVPLEVYTRHEIKDSGLLDYTEVQRRNQLHDLRFAD 718
HS1GPB_4        LSRADITTVSTFIDLNITMLEDHEFVPLEVYTRHEIKDSGLLDYTEVQRRNQLHDLRFAD 648
HS1GPB_1        LSRADITTVSTFIDLNITMLEDHEFVPLEVYTRHEIKDSGLLDYTEVQRRNQLHDLRFAD 648
HS1GPB_2        LSRADITTVSTFIDLNITMLEDHEFVPLEVYTRHEIKDSGLLDYTEVQRRNQLHDLRFAD 648
HS1GPB_12       LSRADITTVSTFIDLNITMLEDHEFVPLEVYTRHEIKDSGLLDYTEVQRRNQLHDLRFAD 718
HS1GPB_13       LSRADITTVSTFIDLNITMLEDHEFVPLEVYTRHEIKDSGLLDYTEVQRRNQLHDLRFAD 718
HS1GPB_16       LSRADITTVSTFIDLNITMLEDHEFVPLEVYTRHEIKDSGLLDYTEVQRRNQLHDLRFAD 718
HS1GPB_8        LSRADITTVSTFIDLNITMLEDHEFVPLEVYTRHEIKDSGLLDYTEVQRRNQLHDLRFAD 648
HS1GPB_6        LSRADITTVSTFIDLNITMLEDHEFVPLEVYTRHEIKDSGLLDYTEVQRRNQLHDLRFAD 718
HS1GPB_11       LSRADITTVSTFIDLNITMLEDHEFVPLEVYTRHEIKDSGLLDYTEVQRRNQLHDLRFAD 718
HS1GPB_15       LSRADITTVSTFIDLNITMLEDHEFVPLEVYTRHEIKDSGLLDYTEVQRRNQLHDLRFAD 718
HS1GPB_19       LSRADITTVSTFIDLNITMLEDHEFVPLEVYTRHEIKDSGLLDYTEVQRRNQLHDLRFAD 648
HS1GPB_14       LSRADITTVSTFIDLNITMLEDHEFVPLEVYTRHEIKDSGLLDYTEVQRRNQLHDLRFAD 648
HS1GPB_17       LSRADITTVSTFIDLNITMLEDHEFVPLEVYTRHEIKDSGLLDYTEVQRRNQLHDLRFAD 718
HS1GPB_20       LSRADITTVSTFIDLNITMLEDHEFVPLEVYTRHEIKDSGLLDYTEVQRRNQLHDLRFAD 648
HS1GPB_10       LSRADITTVSTFIDLNITMLEDHEFVPLEVYTRHEIKDSGLLDYTEVQRRNQLHDLRFAD 718
HS1GPB_18       LSRADITTVSTFIDLNITMLEDHEFVPLEVYTRHEIKDSGLLDYTEVQRRNQLHDLRFAD 718
HS2GPB_21       LSRADVTTVSTFIDLNITMLEDHEFVPLEVYTRHEIKDSGLLDYTEVQRRNQLHDLRFAD 715
HS2GPB_22       LSRADVTTVSTFIDLNITMLEDHEFVPLEVYTRHEIKDSGLLDYTEVQRRNQLHDLRFAD 715
HS2GPB_27       LSRADVTTVSTFIDLNITMLEDHEFVPLGVYTRHEIKDSGLLDYTEVQRRNQLHDLRFAD 715
HS2GPB_30       LSRADVTTVSTFIDLNITMLEDHEFVPLGVYTRHEIKDSGLLDYTEVQRRNQLHDLRFAD 715
HS2GPB_31       LSRADVTTVSTFIDLNITMLEDHEFVPLGVYTRHEIKDSGLLDYTEVQRRNQLHDLRFAD 715
HS2GPB_34       LSRADVTTVSTFIDLNITMLEDHEFVPLGVYTRHEIKDSGLLDYTEVQRRNQLHDLRFAD 715
HS2GPB_28       LSRADVTTVSTFIDLNITMLEDHEIVPLEVYTRHEIKDSGLLDYTEVQRRNQLHDLRFAD 715
HS2GPB_24       LSRADVTTVSTFIDLNITMLEDHEFVPLEVYTRHEIKDSGLLDYTEVQRRNQLHDLRFAD 715
HS2GPB_26       LSRADVTTVSTFIDLNITMLEDHEFVPLEVYTRHEIKDSGLLDYTEVQRRNQLHDLRFAD 715
HS2GPB_35       LSRADVTTVSTFIDLNITMLEDHEFVPLEVYTRHEIKDSGLLDYTEVQRRNQLHDLRFAD 712
HS2GPB_37       LSRADVTTVSTFIDLNITMLEDHEFVPLEVYTRHEIKDSGLLDYTEVQRRNQLHDLRFAD 712
HS2GPB_36       LSRADVTTVSTFIDLNITMLEDHEFVPLEVYTRHEIKDSGLLDYTEVQRRNQLHDLRFAD 712
HS2GPB_25       LSRADVTTVSTFIDLNITMLEDHEFVPLEVYTRHEIKDSGLLDYTEVQRRNQLHDLRFAD 645
HS2GPB_23       LSRADVTTVSTFIDLNITMLEDHEFVPLEVYTRHEIKDSGLLDYTEVQRRNQLHDLRFAD 645
HS2GPB_29       LSRADVTTVSTFIDLNITMLEDHEFVPLGVYTRHEIKDSGLLDYTEVQRRNQLHDLRFAD 645
HS2GPB_32       LSRADVTTVSTFIDLNITMLEDHEFVPLGVYTRHEIKDSGLLDYTEVQRRNQLHDLRFAD 645
HS2GPB_33       LSRADVTTVSTFIDLNITMLEDHEFVPLEVYTRHEIKDSGLLDYTEVQRRNQLHDLRFAD 642
HS2GPB_38       LSRADVTTVSTFIDLNITMLEDHEFVPLEVYTRHEIKDSGLLDYTEVQRRNQLHDLRFAD 642
HS2GPB_39       LSRADVTTVSTFIDLNITMLEDHEFVPLEVYTRHEIKDSGLLDYTEVQRRNQLHDLRFAD 642
                *****:***.**************:*** *******************************

HS1GPB_3        IDTVIHADANAAMFAGLGAFFEGMGDLGRAVGKVVMGIVGGVVSAVSGVSSFMSNPFGAL 778
HS1GPB_5        IDTVIHADANAAMFAGLGAFFEGMGDLGRAVGKVVMGIVGGVVSAVSGVSSFMSNPFGAL 778
HS1GPB_7        IDTVIHADANAAMFAGLGAFFEGMGDLGRAVGKVVMGIVGGVVSAVSGVSSFMSNPFGAL 708
HS1GPB_9        IDTVIHADANAAMFAGLGAFFEGMGDLGRAVGKVVMGIVGGVVSAVSGVSSFMSNPFGAL 778
HS1GPB_4        IDTVIHADANAAMFAGLGAFFEGMGDLGRAVGKVVMGIVGGVVSAVSGVSSFISNPFGAL 708
HS1GPB_1        IDTVIHADANAAMFAGLGAFFEGMGDLGRAVGKVVMGIVGGVVSAVSGVSSFMSNPFGAL 708
HS1GPB_2        IDTVIHADANAAMFAGLGAFFEGMGDLGRAVGKVVMGIVGGVVSAVSGVSSFMSNPFGAL 708
HS1GPB_12       IDTVIHADANAAMFAGLGAFFEGMGDLGRAVGKVVMGIVGGVVSAVSGVSSFMSNPFGAL 778
HS1GPB_13       IDTVIHADANAAMFAGLGAFFEGMGDLGRAVGKVVMGIVGGVVSAVSGVSSFMSNPFGAL 778
HS1GPB_16       IDTVIHADANAAMFAGLGAFFEGMGDLGRAVGKVVMGIVGGVVSAVSGVSSFMSNPFGAL 778
HS1GPB_8        IDTVIHADANAAMFAGLGAFFEGMGDLGRAVGKVVMGIVGGVVSAVSGVSSFMSNPFGAL 708
HS1GPB_6        IDTVIHADANAAMFAGLGAFFEGMGDLGRAVGKVVMGIVGGVVSAVSGVSSFMSNPFGAL 778
HS1GPB_11       IDTVIHADANAAMFAGLGAFFEGMGDLGRAVGKVVMGIVGGVVSAVSGVSSFMSNPFGAL 778
HS1GPB_15       IDTVIHADANAAMFAGLGAFFEGMGDLGRAVGKVVMGIVGGVVSAVSGVSSFMSNPFGAL 778
HS1GPB_19       IDTVIHADANAAMFAGLGAFFEGMGDLGRAVGKVVMGIVGGVVSAVSGVSSFMSNPFGAL 708
HS1GPB_14       IDTVIHADANAAMFAGLGAFFEGMGDLGRAVGKVVMGIVGGVVSAVSGVSSFMSNPFGAL 708
HS1GPB_17       IDTVIHADANAAMFAGLGAFFEGMGDLGRAVGKVVMGIVGGVVSAVSGVSSFMSNPFGAL 778
HS1GPB_20       IDTVIHADANAAMFAGLGAFFEGMGDLGRAVGKVVMGIVGGVVSAVSGVSSFMSNPFGAL 708
HS1GPB_10       IDTVIHADANAAMFAGLGAFFEGMGDLGRAVGKVVMGIVGGVVSAVSGVSSFMSNPFGAL 778
HS1GPB_18       IDTVIHADANAAMFAGLGAFFEGMGDLGRAVGKVVMGIVGGVVSAVSGVSSFMSNPFGAL 778
HS2GPB_21       IDTVIRADANAAMFAGLCAFFEGMGDLGRAVGKVVMGVVGGVVSAVSGVSSFMSNPFGAL 775
HS2GPB_22       IDTVIRADANAAMFAGLCAFFEGMGDLGRAVGKVVMGVVGGVVSAVSGVSSFMSNPFGAL 775
HS2GPB_27       IDTVIRADANAAMFAGLCAFFEGMGDLGRAVGKVVMGVVGGVVSAVSGVSSFMSNPFGAL 775
HS2GPB_30       IDTVIRADANAAMFAGLCAFFEGMGDLGRAVGKVVMGVVGGVVSAVSGVSSFMSNPFGAL 775
HS2GPB_31       IDTVIRADANAAMFAGLCAFFEGMGDLGRAVGKVVMGVVGGVVSAVSGVSSFMSNPFGAL 775
HS2GPB_34       IDTVIRADANAAMFAGLCAFFEGMGDLGRAVGKVVMGVVGGVVSAVSGVSSFMSNPFGAL 775
HS2GPB_28       IDTVIRADANAAMFAGLCAFFEGMGDLGRAVGKVVMGVVGGVVSAVSGVSSFMSNPFGAL 775
HS2GPB_24       IDTVIRADANAAMFAGLCAFFEGMGDLGRAVGKVVMGVVGGVVSAVSGVSSFMSNPFGAL 775
HS2GPB_26       IDTVIRADANAAMFAGLCAFFEGMGDLGRAVGKVVMGVVGGVVSAVSGVSSFMSNPFGAL 775
HS2GPB_35       IDTVIRADANAAMFAGLCAFFEGMGDLGRAVGKVVMGVVGGVVSAVSGVSSFMSNPFGAL 772
HS2GPB_37       IDTVIRADANAAMFAGLCAFFEGMGDLGRAVGKVVMGVVGGVVSAVSGVSSFMSNPFGAL 772
HS2GPB_36       IDTVIRADANAAMFAGLCAFFEGMGDLGRAVGKVVMGVVGGVVSAVSGVSSFMSNPFGAL 772
HS2GPB_25       IDTVIRADANAAMFAGLCAFFEGMGDLGRAVGKVVMGVVGGVVSAVSGVSSFMSNPFGAL 705
HS2GPB_23       IDTVIRADANAAMFAGLCAFFEGMGDLGRAVGKVVMGVVGGVVSAVSGVSSFMSNPFGAL 705
HS2GPB_29       IDTVIRADANAAMFAGLCAFFEGMGDLGRAVGKVVMGVVGGVVSAVSGVSSFMSNPFGAL 705
HS2GPB_32       IDTVIRADANAAMFAGLCAFFEGMGDLGRAVGKVVMGVVGGVVSAVSGVSSFMSNPFGAL 705
HS2GPB_33       IDTVIRADANAAMFAGLCAFFEGMGDLGRAVGKVVMGVVGGVVSAVSGVSSFMSNPFGAL 702
HS2GPB_38       IDTVIRADANAAMFAGLCAFFEGMGDLGRAVGKVVMGVVGGVVSAVSGVSSFMSNPFGAL 702
HS2GPB_39       IDTVIRADANAAMFAGLCAFFEGMGDLGRAVGKVVMGVVGGVVSAVSGVSSFMSNPFGAL 702
                *****:*********** *******************:**************:*******

HS1GPB_3        AVGLLVLAGLAAAFFAFRYVMRLQSNPMKALYPLTTKELKNPTNPDASGEGEE---GGDF 835
HS1GPB_5        AVGLLVLAGLAAAFFAFRYVMRLQSNPMKALYPLTTKELKNPTNPDASGEGEE---GGDF 835
HS1GPB_7        AVGLLVLAGLAAAFFAFRYVMRLQSNPMKALYPLTTKEPKNPTNPDASGEGEE---GGDF 765
HS1GPB_9        AVGLLVLAGLAAAFFAFRYVMRLQSNPMKALYPLTTKEPKNPTNPDASGEGEE---GGDF 835
HS1GPB_4        AVGLLVLAGLAAAFFAFRYVMRLQSNPMKALYPLTTKELKNPTNPDASGEGEE---GGDF 765
HS1GPB_1        AVGLLVLAGLAAAFFAFRYVMRLQSNPMKALYPLTTKELKNPTNPDASGEGEE---GGDF 765
HS1GPB_2        AVGLLVLAGLAAAFFAFRYVMRLQSNPMKALYPLTTKELKNPTNPDASGEGEE---GGDF 765
HS1GPB_12       AVGLLVLAGLAAAFFAFRYVMRLQSNPMKALYPLTTKELKNPTNPDASGEGEE---GGDF 835
HS1GPB_13       AVGLLVLAGLAAAFFAFRYVMRLQSNPMKALYPLTTKELKNPTNPDASGEGEE---GGDF 835
HS1GPB_16       AVGLLVLAGLAAAFFAFRYVMRLQSNPMKALYPLTTKELKNPTNPDASGEGEE---GGDF 835
HS1GPB_8        AVGLLVLAGLAAAFFAFRYVMRLQSNPMKALYPLTTKELKNPTNPDASGEGEE---GGDF 765
HS1GPB_6        AVGLLVLAGLAAAFFAFRYVMRLQSNPMKALYPLTTKELKNPTNPDASGEGEE---GGDF 835
HS1GPB_11       AVGLLVLAGLAAAFFAFRYVMRLQSNPMKALYPLTTKELKNPTNPDASGEGEE---GGDF 835
HS1GPB_15       AVGLLVLAGLAAAFFAFRYVMRLQSNPMKALYPLTTKELKNPTNPDASGEGEE---GGDF 835
HS1GPB_19       AVGLLVLAGLAAAFFAFRYVMRLQSNPMKALYPLTTKELKNPTNPDASGEGEE---GGDF 765
HS1GPB_14       AVGLLVLAGLAAAFFAFRYVMRLQSNPMKALYPLTTKELKNPTNPDASGEGEE---GGDF 765
HS1GPB_17       AVGLLVLAGLAAAFFAFRYVMRLQSNPMKALYPLTTKELKNPTNPDASGEGEE---GGDF 835
HS1GPB_20       AVGLLVLAGLAAAFFAFRYVMRLQSNPMKALYPLTTKELKNPTNPDASGEGEE---GGDF 765
HS1GPB_10       AVGLLVLAGLAAAFFAFRYVMRLQSNPMKALYPLTTKELKNPTNPDASGEGEE---GGDF 835
HS1GPB_18       AVGLLVLAGLAAAFFAFRYVMRLQSNPMKALYPLTTKELKNPTNPDASGEGEE---GGDF 835
HS2GPB_21       AVGLLVLAGLVAAFFAFRYVLQLQRNPMKALYPLTTKELKTSDPGGVGGEGEEGAEGGGF 835
HS2GPB_22       AVGLLVLAGLVAAFFAFRYVLQLQRNPMKALYPLTTKELKTSDPGGVGGEGEEGAEGGGF 835
HS2GPB_27       AVGLLVLAGLVAAFFAFRYVLQLQRNPMKALYPLTTKELKTSDPGGVGGEGEEGAEGGGF 835
HS2GPB_30       AVGLLVLAGLVAAFFAFRYVLQLQRNPMKALYPLTTKELKTSDPGGVGGEGEEGAEGGGF 835
HS2GPB_31       AVGLLVLAGLVAAFFAFRYVLQLQRNPMKALYPLTTKEPKTSDPGGVGGEGEEGAEGGGF 835
HS2GPB_34       AVGLLVLAGLVAAFFAFRYVLQLQRNPMKALYPLTTKEPKTSDPGGVGGEGEEGAEGGGF 835
HS2GPB_28       AVGLLVLAGLVAAFFAFRYVLQLQRNPMKALYPLTTKELKTSDPGGVGGEGEEGAEGGGF 835
HS2GPB_24       AVGLLVLAGLVAAFFAFRYVLQLQRNPMKALYPLTTKELKTSDPGGVGGEGEEGAEGGGF 835
HS2GPB_26       AVGLLVLAGLVAAFFAFRYVLQLQRNPMKALYPLTTKELKTSDPGGVGGEGEEGAEGGGF 835
HS2GPB_35       AVGLLVLAGLVAAFFAFRYVLQLQRNPMKALYPLTTKELKTSDPGGVGGEGEEGAEGGGF 832
HS2GPB_37       AVGLLVLAGLVAAFFAFRYVLQLQRNPMKALYPLTTKELKTSDPGGVGGEGEEGAEGGGF 832
HS2GPB_36       AVGLLVLAGLVAAFFAFRYVLQLQRNPMKALYPLTTKELKTSDPGGVGGEGEEGAEGGGF 832
HS2GPB_25       AVGLLVLAGLVAAFFAFRYVLQLQRNPMKALYPLTTKELKTSDPGGVGGEGEEGAEGGGF 765
HS2GPB_23       AVGLLVLAGLVAAFFAFRYVLQLQRNPMKALYPLTTKELKTSDPGGVGGEGEEGAEGGGF 765
HS2GPB_29       AVGLLVLAGLVAAFFAFRYVLQLQRNPMKALYPLTTKELKTSDPGGVGGEGEEGAEGGGF 765
HS2GPB_32       AVGLLVLAGLVAAFFAFRYVLQLQRNPMKALYPLTTKEPKTSDPGGVGGEGEEGAEGGGF 765
HS2GPB_33       AVGLLVLAGLVAAFFAFRYVLQLQRNPMKALYPLTTKELKTSDPGGVGGEGEEGAEGGGF 762
HS2GPB_38       AVGLLVLAGLVAAFFAFRYVLQLQRNPMKALYPLTTKELKTSDPGGVGGEGEEGAEGGGF 762
HS2GPB_39       AVGLLVLAGLVAAFFAFRYVLQLQRNPMKALYPLTTKELKTSDPGGVGGEGEEGAEGGGF 762
                **********.*********::** ************* *..   ...*****   **.*

HS1GPB_3        DEAKLAEAREMIRYMALVSAMERTEHKAKKKGTSALLSAKVTDMVMRKRRNTNYTQVPNK 895
HS1GPB_5        DEAKLAEAREMIRYMALVSVMERTEHKAKKKGTSALLSAKVTDMVMRKRRNTNYTQVPNK 895
HS1GPB_7        DEAKLAEAREMIRYMALVSAMERTEHKAKKKGTSALLSAKVTDMVMRKRRNTNYTQVPNK 825
HS1GPB_9        DEAKLAEAREMIRYMALVSAMERTEHKAKKKGTSALLSAKVTDMVMRKRRNTNYTQVPNK 895
HS1GPB_4        DEAKLAEAREMIRYMALVSAMERTEHKAKKKGTSALLSAKVTDMVMRKRRNTNYTQVPNK 825
HS1GPB_1        DEAKLAEAREMIRYMALVSAMERTEHKAKKKGTSALLSAKVTDMVMRKRRNTNYTQVPNK 825
HS1GPB_2        DEAKLAEAREMIRYMALVSAMERTEHKAKKKGTSALLSAKVTDMVMRKRRNTNYTQVPNK 825
HS1GPB_12       DEAKLAEAREMIRYMALVSAMERTEHKAKKKGTSALLSAKVTDMVMRKRRNTNYTQVPNK 895
HS1GPB_13       DEAKLAEAREMIRYMALVSAMERTEHKAKKKGTSALLSAKVTDMVMRKRRNTNYTQVPNK 895
HS1GPB_16       DEAKLAEAREMIRYMALVSAMERTEHKAKKKGTSALLSAKVTDMVMRKRRNTNYTQVPNK 895
HS1GPB_8        DEAKLAEAREMIRYMALVSAMERTEHKAKKKGTSALLSAKVTDMVMRKRRNTNYTQVPNK 825
HS1GPB_6        DEAKLAEAREMIRYMALVSAMERTEHKAKKKGTSALLSAKVTDMVMRKRRNTNYTQVPNK 895
HS1GPB_11       DEAKLAEAREMIRYMALVSAMERTEHKAKKKGTSALLSAKVTDMVMRKRRNTNYTQVPNK 895
HS1GPB_15       DEAKLAEAREMIRYMALVSAMERTEHKAKKKGTSALLSAKVTDMVMRKRRNTNYTQVPNK 895
HS1GPB_19       DEAKLAEAREMIRYMALVSAMERTEHKAKKKGTSALLSAKVTDMVMRKRRNTNYTQVPNK 825
HS1GPB_14       DEAKLAEAREMIRYMALVSAMERTEHKAKKKGTSALLSAKVTDMVMRKRRNTNYTQVPNK 825
HS1GPB_17       DEAKLAEAREMIRYMALVSAMERTEHKAKKKGTSALLSAKVTDMVMRKRRNTNYTQVPNK 895
HS1GPB_20       DEAKLAEAREMIRYMALVSAMERTEHKAKKKGTSALLSAKVTDMVMRKRRNTNYTQVPNK 825
HS1GPB_10       DEAKLAEAREMIRYMALVSAMERTEHKAKKKGTSALLSAKVTDMVMRKRRNTNYTQVPNK 895
HS1GPB_18       DEAKLAEAREMIRYMALVSAMERTEHKAKKKGTSALLSAKVTDMVMRKRRNTNYTQVPNK 895
HS2GPB_21       DEAKLAEAREMIRYMALVSAMERTEHKARKKGTSALLSSKVTNMVLRKRNKARYSPLHNE 895
HS2GPB_22       DEAKLAEAREMIRYMALVSAMERTEHKARKKGTSALLSSKVTNMVLRKRNKARYSPLHNE 895
HS2GPB_27       DEAKLAEAREMIRYMALVSAMERTEHKARKKGTSALLSSKVTNMVLRKRNKARYSPLHNE 895
HS2GPB_30       DEAKLAEAREMIRYMALVSAMERTEHKARKKGTSALLSSKVTNMVLRKRNKARYSPLHNE 895
HS2GPB_31       DEAKLAEAREMIRYMALVSAMERTEHKARKKGTSALLSSKVTNMVLRKRNKARYSPLHNE 895
HS2GPB_34       DEAKLAEAREMIRYMALVSAMERTEHKARKKGTSALLSSKVTNMVLRKRNKARYSPLHNE 895
HS2GPB_28       DEAKLAEAREMIRYMALVSAMERTEHKARKKGTSALLSSKVTNMVLRKRNKARYSPLHNE 895
HS2GPB_24       DEAKLAEAREMIRYMALVSAMERTEHKARKKGTSALLSSKVTNMVLRKRNKARYSPLHNE 895
HS2GPB_26       DEAKLAEAREMIRYMALVSAMERTEHKARKKGTSALLSSKVTNMVLRKRNKARYSPLHNE 895
HS2GPB_35       DEAKLAEARQMIRYMALVSAMERTEHKARKKGTSALLSSKVTNMVLRKRNKARYSPLHNE 892
HS2GPB_37       DEAKLAEARQMIRYMALVSAMERTEHKARKKGTSALLSSKVTNMVLRKRNKARYSPLHNE 892
HS2GPB_36       DEAKLAEAREMIRYMALVSAMERTEHKARKKGTSALLSSKVTNMVLRKRNKARYSPLHNE 892
HS2GPB_25       DEAKLAEAREMIRYMALVSAMERTEHKARKKGTSALLSSKVTNMVLRKRNKARYSPLHNE 825
HS2GPB_23       DEAKLAEAREMIRYMALVSAMERTEHKARKKGTSALLSSKVTNMVLRKRNKARYSPLHNE 825
HS2GPB_29       DEAKLAEAQEMIRYMALVSAMERTEHKARKKGTSALLSSKVTNMVLRKRNKARYSPLHNE 825
HS2GPB_32       DEAKLAEAREMIRYMALVSAMERTEHKARKKGTSALLSSKVTNMVLRKRNKARYSPLHNE 825
HS2GPB_33       DEAKLAEAREMIRYMALVSAMERTEHKARKKGTSALLSSKVTNMVLRKRNKARYSPLHNE 822
HS2GPB_38       DEAKLAEAREMIRYMALVSAMERTEHKARKKGTSALLSSKVTNMVLRKRNKARYSPLHNE 822
HS2GPB_39       DEAKLAEAREMIRYMALVSAMERTEHKARKKGTSALLSSKVTNMVLRKRNKARYSPLHNE 822
                ********::*********.********:*********:***:**:***.::.*: : *:

HS1GPB_3        DGDADEDDL 904
HS1GPB_5        DGDADEDDL 904
HS1GPB_7        DGDADEDDL 834
HS1GPB_9        DGDADEDDL 904
HS1GPB_4        DGDADEDDL 834
HS1GPB_1        DGDADEDDL 834
HS1GPB_2        DGDADEDDL 834
HS1GPB_12       DGDADEDDL 904
HS1GPB_13       DGDADEDDL 904
HS1GPB_16       DGDADEDDL 904
HS1GPB_8        DGDADEDDL 834
HS1GPB_6        DGDADEDDL 904
HS1GPB_11       DGDADEDDL 904
HS1GPB_15       DGDADEDDL 904
HS1GPB_19       DGDADEDDL 834
HS1GPB_14       DGDADEDDL 834
HS1GPB_17       DGDADEDDL 904
HS1GPB_20       DGDADEDNL 834
HS1GPB_10       DGDADEDDL 904
HS1GPB_18       DGDADEDDL 904
HS2GPB_21       DEAGDEDEL 904
HS2GPB_22       DEAGDEDEL 904
HS2GPB_27       DEAGDEDEL 904
HS2GPB_30       DEAGDEDEL 904
HS2GPB_31       DEAGDEDEL 904
HS2GPB_34       DEAGDEDEL 904
HS2GPB_28       DEAGDEDEL 904
HS2GPB_24       DEAGDEDEL 904
HS2GPB_26       DEAGDEDEL 904
HS2GPB_35       DEAGDEDEL 901
HS2GPB_37       DEAGDEDEL 901
HS2GPB_36       DEAGDEDEL 901
HS2GPB_25       DEAGDEDEL 834
HS2GPB_23       DEAGDEDEL 834
HS2GPB_29       DEAGDEDEL 834
HS2GPB_32       DEAGDEDEL 834
HS2GPB_33       DEAGDEDEL 831
HS2GPB_38       DEAGDEDEL 831
HS2GPB_39       DEAGDEDEL 831
                *  .***:*
```
