## Supplementary file 8 for "Contriving a chimeric polyvalent vaccine to prevent infections caused by Herpes Simplex Virus (Type-1 and Type-2): an exploratory immunoinformatic approach"

```
Alignment of Glycoprotein C

HSV1GPC_10      MAPGRVGLAVVLWSLLWLGAGVAGGSETASTGPTITAGAVTNASEAPTSGSPGSAASPEV 60
HSV1GPC_20      MAPGRVGLAVVLWSLLWLGAGVAGGSETASTGPTITAGAVTNASEAPTSGSPGSAASPEV 60
HSV1GPC_8       MAPGRVGLAVVLWSLLWLGAGVAGGSETASTGPTITAGAVTNASEAPTSGSPGSAASPEV 60
HSV1GPC_14      ------------------------------------------------------------
HSV1GPC_6       ------------------------------------------------------------
HSV1GPC_15      ------------------------------------------------------------
HSV1GPC_1       ------------------------------------------------------------
HSV1GPC_12      MAPGRVGLAVVLWSLVWLGAGVSGGSETASTGPTITAGAVMNASEAPTSGSPGSAASPEV 60
HSV1GPC_7       MAPGRVGLAVVLWSLLWLGAGVSGGSETASTGPTITAGAVTNASEAPTSGSPGSAASPEV 60
HSV1GPC_19      MAPGRVGLAVVLWSLLWLGAGVSGGSETASTGPTITAGAVTNASEAPTSGSPGSAASPEV 60
HSV1GPC_5       MAPGRVGLAVVLWSLLWLGAGVSGGSETASTGPTITAGAVTNASEAPTSGSPGSAASPEV 60
HSV1GPC_11      ------------------------------------------------------------
HSV1GPC_17      ------------------------------------------------------------
HSV1GPC_18      ------------------------------------------------------------
HSV1GPC_4       MAPGRVGLAVVLWSLLWLGAGVSGGSETASTGPTITAGAVTNASEAPTSGSPGSAASPEV 60
HSV1GPC_13      MAPGRVGLAVVLWSLLWLGAGVSGGSETASTGPTITAGAVTNASEAPTSGSPGSAASPEV 60
HSV1GPC_9       MAPGRVGLAVVLWSLLWLGAGVSGDSETASTGPTITAGAVTNASEAPTSGSPGSAASPEV 60
HSV1GPC_2       MAPGRVGLAVVLWSLLWLGAGVSGGSETASTGPTITAGAVTNASEAPTSGSPGSAASPEV 60
HSV1GPC_3       MAPGRVGLAVVLWSLLWLGAGVSGGSETASTGPTITAGAVTNASEAPTSGSPGSAANPEV 60
HSV1GPC_16      MAPGRVGLAVVLWSLLWLGAGVAGGSETASTGPTITAGAVTNASEAPTSGSPGSAASPEV 60
HSV2GPC_25      MALGRVGLAVGLWGLLWVGVVVVLAN--ASPGRTITVGPRGNASN--------------- 43
HSV2GPC_27      MALGRVGLTVGLWGLLWVGVVVVLAN--ASPGRTITVGPRGNASN--------------- 43
HSV2GPC_26      MALGRVGLAVGLWGLLWVGVVVVLAN--ASPGRTITVGPRGNASN--------------- 43
HSV2GPC_24      MALGRVGLAVGLWGLLWVGVVVVLAN--ASPGRTITVGPRGNASN--------------- 43
HSV2GPC_21      MALGRVGLAVGLWGLLWVGVVVVLAN--ASPGRTITVGPRGNASN--------------- 43
HSV2GPC_23      MALGRVGLAVGLWGLLWVGVVVVLAN--ASPGRTITVGPRGNASN--------------- 43
HSV2GPC_22      ------------------------------------------------------------


HSV1GPC_10      TPTSTPNPNNVTQNKTTPTEPASPPTTPKPTSTPKSPPTSTPDPKPKNNTTPAKSGRPTK 120
HSV1GPC_20      TPTSTPNPNNVTQNKTTPTEPASPPTTPKPTSTPKSPPTSTPDPKPKNNTTPAKSGRPTK 120
HSV1GPC_8       TPTSTPTPNNVTQNKTTPTEPASPPTTPKPTSTPKSPPTSTPDPKPKNNTTPAKSGRPTK 120
HSV1GPC_14      ----------VTQNKTTPTEPASPPTTPKPTSTPKSPPTSTPDPKPKNNTTPAKSGRPTK 50
HSV1GPC_6       ----------VTQNKTTPTEPASPPTTPKPTSTPKSPPTSTPDPKPKNNTTPAKSGRPTK 50
HSV1GPC_15      ----------VTQNKTTPTEPASPPTTPKPTSTPKSPPTSTPDPKPKNNTTPAKSGRPTK 50
HSV1GPC_1       ----------VTQNKTTPTEPASPPTTPKPTSTPKSPPTSTPDPKPKNNTTPAKSGRPTK 50
HSV1GPC_12      TPTSTPNPNNVTQNQTTPTEPASPPTTPKPTSTPKSPPTSTPDPKPKNNTTPAKSGRPTK 120
HSV1GPC_7       TPTSTPNPNNVTQNKTTPTEPASPPTTPKPTSTPKSPPTSTPDPKPKNNTTPAKSGRPTK 120
HSV1GPC_19      TPTSTPNPNNVTQNKTTPTEPASPPTTPKPTSTPKSPPTSTPDPKPKNNTTPAKSGRPTN 120
HSV1GPC_5       TPTSTPNPNNVTQNKTTPTEPASPPTTPKPTSTPKSPPTSTPDPKPKNNTTPAKSGRPTK 120
HSV1GPC_11      ----------VTQNKTTPTEPASPPTTPKPTSTPKSPPTSTPDPKPKNNTTPAKSGRPTK 50
HSV1GPC_17      ----------VTQNKTTPTEPASPPTTPKPTSTPKSPPTSTPDPKPKNNTTPAKSGRPTK 50
HSV1GPC_18      ----------VTQNKTTPTEPASPPTTPKPTSTPKSPPTSTPDPKPKNNTTPAKSGRPTK 50
HSV1GPC_4       TPTSTPNPNNVTQNKTTPTEPASPPTTPKPTSTPKSPPTSTPDPKPKNNTTPAKSGRPTK 120
HSV1GPC_13      TPTSTPNPNNVTQNKTIPTEPASPPTTPKPTSTPKSPPTSTPDPKPKNNTTPAKSGRPTK 120
HSV1GPC_9       TPTSTPNPNNVTQNKTIPTEPASPPTTPKPTSTPKSPPTSTPDPKPKNNTTPAKSGRPTK 120
HSV1GPC_2       TPTSTPNPNNVTQNKTTPTEPASPPTTPKPTSTPKSPPTSTPDPKPKNNTTPAKSGRPTK 120
HSV1GPC_3       TPTSTPNPNNVTQNKTTPTEPASPPTTPKPTSTPKSPPTSTPDPKPKNNTTPAKSGRPTK 120
HSV1GPC_16      TPTSTPNPNNVTQNKTTPTEPASPPTTPKPTSTPKSPPTSTPDPKPKNNTTPAKSGRPTK 120
HSV2GPC_25      -----------AAPSASPRNASAPRTTPTPPQPRKATKSKASTAKP---APPPKTGPPKT 89
HSV2GPC_27      -----------AAPSV-PRNRSAPRTTPTPPQPRKATKSKASTAKP---APPPKTGPPKT 88
HSV2GPC_26      -----------AAPSASPRNASAPRTTPTPPQPRKATKSKASTAKP---APPPKTGPPKT 89
HSV2GPC_24      -----------AAPSASPRNASAPRTTPTPPQPRKATKSKASTAKP---APPPKTGPPKT 89
HSV2GPC_21      -----------AAPSASPRNASAPRTTPTPPQPRKATKSKASTAKP---APPPKTGPPKT 89
HSV2GPC_23      -----------AAPSASPRNASAPRTTPTPPQPRKATKSKASTAKP---APPPKTGPPKT 89
HSV2GPC_22      --------------------------------------SKASTAKP---APPPKTGPPKT 19
                                                      :.:. .**   :.*.*:* *..

HSV1GPC_10      PPGPVWCDRRDPLARYGSRVQIRCRFRNSTRMEFRLQIWRYSMGPSPPIAPAPDLEEVLT 180
HSV1GPC_20      PPGPVWCDRRDPLARYGSRVQIRCRFRNSTRMEFRLQIWRYSMGPSPPIAPAPDLEEVLT 180
HSV1GPC_8       PPGPVWCDRRDPLARYGSRVQIRCRFRNSTRMEFRLQIWRYSMGPSPPIAPAPDLEEVLT 180
HSV1GPC_14      PPGPVWCDRRDPLARYGSRVQIRCRFRNSTRMEFRLQIWRYSMGPSPPIAPAPDLEEVLT 110
HSV1GPC_6       PPGPVWCDRRDPLARYGSRVQIRCRFRNSTRMEFRLQIWRYSMGPSPPIAPAPDLEEVLT 110
HSV1GPC_15      PPGPVWCDRRDPLARYGSRVQIRCRFRNSTRMEFRLQIWRYSMGPSPPIAPAPDLEEVLT 110
HSV1GPC_1       PPGPVWCDRRDPLARYGSRVQIRCRFRNSTRMEFRLQIWRYSMGPSPPIAPAPDLEEVLT 110
HSV1GPC_12      PPGPVWCDRRDPLARYGSRVQIRCRFRNSTRMEFRLQIWRYSMGPSPPIAPAPDLEEVLT 180
HSV1GPC_7       PPGPVWCDRRDPLARYGSRGQIRCRFRNSTRMEFRLQIWRYSMGPSPPIAPAPDLEEVLT 180
HSV1GPC_19      PPGPVWCDRRDPLARYGSRVQIRCRFRNSTRMEFRLQIWRYSMGPSPPIAPAPDLEEVLT 180
HSV1GPC_5       PPGPVWCDRRDPLARYGSRVQIRCRFRNSTRMEFRLQIWRYSMGPSPPIAPAPDLEEVLT 180
HSV1GPC_11      PPGPVWCDRRDPLARYGSRVQIRCRFRNSTRMEFRLQIWRYSMGPSPPIAPAPDLEEVLT 110
HSV1GPC_17      PPGPVWCDRRDPLARYGSRVQIRCRFRNSTRMEFRLQIWRYSMGPSPPIAPAPDLEEVLT 110
HSV1GPC_18      PPGPVWCDRRDPLARYGSRVQIRCRFRNSTRMEFRLQIWRYSTGPSPPIAPAPDLEEVLT 110
HSV1GPC_4       PPGPVWCDRRDPLARYGSRVQIRCRFRNSTRMEFRLQIWRYSMGPSPPIAPAPDLEEVLT 180
HSV1GPC_13      PPGPVWCDRRDPLARYGSRVQIRCRFRNSTRMEFRLQIWRYSMGPSPPIAPAPDLEEVLT 180
HSV1GPC_9       PPGPVWCDRRDPLARYGSRVQIRCRFRNSTRMEFRLQIWRYSMGPSPPIAPAPDLEEVLT 180
HSV1GPC_2       PPGPVWCDRRDPLARYGSRVQIRCRFRNSTRMEFRLQIWRYSMGPSPPIAPAPDLEEVLT 180
HSV1GPC_3       PPGPVWCDRRDPLARYGSRVQIRCRFRNSTRMEFRLQIWRYSMGPSPPIAPAPDLEEVLT 180
HSV1GPC_16      PPGPVWCDRRDPLARYGSRVQIRCRFRNSTRMEFRLQIWRYSMGPSPPIAPAPDLEEVLT 180
HSV2GPC_25      SSEPVRCNRHDPLARYGSRVQIRCRFPNSTRTESRLQIWRYATATDAEIGTAPSLEEVMV 149
HSV2GPC_27      SSEPVRCNRHDPLARYGSRVQIRCRFPNSTRTESRLQIWRYATATDAEIGTAPSLEEVMV 148
HSV2GPC_26      SSEPVRCNRHDPLARYGSRVQIRCRFPNSTRTESRLQIWRYATATDAEIGTAPSLEEVMV 149
HSV2GPC_24      SSEPVRCNRHDPLARYGSRVQIRCRFPNSTRTESRLQIWRYATATDAEIGTAPSLEEVMV 149
HSV2GPC_21      SSEPVRCNRHDPLARYGSRVQIRCRFPNSTRTEFRLQIWRYATATDAEIGTAPSLEEVMV 149
HSV2GPC_23      SSEPVRCNRHDPLARYGSRVQIRCRFPNSTRTEFRLQIWRYATATDAEIGTAPSLEEVMV 149
HSV2GPC_22      SSEPVRCNRHDPLARYGSRVQIRCRFPNSTRTEFRLQIWRYATATDAEIGTAPSLEEVMV 79
                .. ** *:*:********* ****** **** * *******: .... *..**.****:.

HSV1GPC_10      NITAPPGGLLVYDSAPNLTDPHVLWAEGAGPGADPPLYSVTGPLPTQRLIIGEVTPATQG 240
HSV1GPC_20      NITAPPGGLLVYDSAPNLTDPHVLWAEGAGPGADPPLYSVTGPLPTQRLIIGEVTPATQG 240
HSV1GPC_8       NITAPPGGLLVYDSAPNLTDPHVLWAEGAGPGADPPLYSVTGPLPTQRLIIGEVTPATQG 240
HSV1GPC_14      NITAPPGGLLVYDSAPNLTDPHVLWAEGAGPGADPPLYSVTGPLPTQRLIIGEVTPATQG 170
HSV1GPC_6       NITAPPGGLLVYDSAPNLTDPHVLWAEGAGPGADPPLYSVTGPLPTQRLIIGEVTPATQG 170
HSV1GPC_15      NITAPPGGLLVYDSAPNLTDPHVLWAEGAGPGADPPLYSVTGPLPTQRLIIGEVTPATQG 170
HSV1GPC_1       NITAPPGGLLVYDSAPNLTDPHVLWAEGAGPGADPPLYSVTGPLPTQRLIIGEVTPATQG 170
HSV1GPC_12      NITAPPGGLLVYDSAPNLTDPHVLWAEGAGPGADPPLYSVTGPLPTQRLIIGEVTPATQG 240
HSV1GPC_7       NITAPPGGLLVYDSAPNLTDPHVLWAEGAGPGADPPLYSVTGPLPTQRLIIGEVTPATQG 240
HSV1GPC_19      NITAPPGGLLVYDSAPNLTDPHVLWAEGAGPGADPPLYSVTGPLPTQRLIIGEVTPATQG 240
HSV1GPC_5       NITAPPGGLLVYDSAPNLTDPHVLWAEGAGPGADPPLYSVTGPLPTQRLIIGEVTPATQG 240
HSV1GPC_11      NITAPPGGLLVYDSAPNLTDPHVLWAEGAGPGADPPLYSVTGPLPTQRLIIGEVTPATQG 170
HSV1GPC_17      NIPAPPGGLLVYDSAPNLTDPHVLWAEGAGPGADPPLYSVTGPLPTQRLIIGEVTPATQG 170
HSV1GPC_18      NITAPPGGLLVYDSAPNLTDPHVLWAEGAGPGADPPLYSVTGPLPTQRLIIGEVTPATQG 170
HSV1GPC_4       NITAPPGGLLVYDSAPNLTDPHVLWAEGAGPGADPPLYSVTGPLPTQRLIIGEVTPATQG 240
HSV1GPC_13      NITAPPGGLLVYDSAPNLTDPHVLWAEGAGPGADPPLYSVTGPLPTQRLIIGEVTPATQG 240
HSV1GPC_9       NITAPPGGLLVYDSAPNLTDPHVLWAEGAGPGADPPLYSVTGPLPTQRLIIGEVTPATQG 240
HSV1GPC_2       NITAPPGGLLVYDSAPNLTDPHVLWAEGAGPGADPPLYSVTGPLPTQRLIIGEVTPATQG 240
HSV1GPC_3       NITAPPGGLLVYDSAPNLTDPHVLWAEGAGPGADPPLYSVTGPLPTQRLIIGEVTPATQG 240
HSV1GPC_16      NITAPPGGLLVYDSAPNLTDPHVLWAEGAGPGADPPLYSVTGPLPTQRLIIGEVTPATQG 240
HSV2GPC_25      NVSAPPGGQLVYDSAPNRTDPHVIWAEGAGPGASPRLYSVVGPLGRQRPIIEELTLETQG 209
HSV2GPC_27      NVSAPPGGQLVYDSAPNRTDPHVIWAEGAGPGASPRLYSVVGPLGRQRLIIEELTLETQG 208
HSV2GPC_26      NVSAPPGGQLVYDSPPNRTDPHVIWAEGAGPGASPRLYSVVGPLGRQRLIIEELTLETQG 209
HSV2GPC_24      NVSAPPGGQLVYDSAPNRTDPHVIWAEGAGPGASPRLYSVVGPLGRQRLIIEELTLETQG 209
HSV2GPC_21      NVSAPPGGQLVYDSAPNRTDPHVIWAEGAGPGASPRLYSVVGPLGRQRLIIEELTLETQG 209
HSV2GPC_23      NVSAPPGGQLVYDSAPNRTDPHVIWAEGAGPGASPRLYSVVGPLGRQRLIIEELTLETQG 209
HSV2GPC_22      NVSAPPGGQLVYDSAPNRTDPHVIWAEGAGPGASPRLYSVVGPLGRQRLIIEELTLETQG 139
                *:.***** *****.** *****:*********.* ****.***  ** ** *:*  ***

HSV1GPC_10      MYYLAWGRMDSPHEYGTWVRVRMFRPPSLTLQPHAVMEGQPFKATCTAAAYYPRNPVEFV 300
HSV1GPC_20      MYYLAWGRMDNPHEYGTWVRVRMFRPPSLTLQPHAVMEGQPFKATCTAAAYYPRNPVEFV 300
HSV1GPC_8       MYYLAWGRMDSPHEYGTWVRVRMFRPPSLTLQPHAVMEGQPFKATCTAAAYYPRNPVEFV 300
HSV1GPC_14      MYYLAWGRMDSPHEYGTWVRVRMFRPPSLTLQPHAVMEGQPFKATCTAAAYYPRNPVEFV 230
HSV1GPC_6       MYYLAWGRMDSPHEYGTWVRVRMFRPPSLTLQPHAVMEGQPFKATCTAAAYYPRNPVEFV 230
HSV1GPC_15      MYYLAWGRIDSPHEYGTWVRVRMFRPPSLTLQPHAVMEGQPFKATCTAAAYYPRNPVEFV 230
HSV1GPC_1       MYYLAWGRMDSPHEYGTWVRVRMFRPPSLTLQPHAVMEGQPFKATCTAAAYYPRNPVEFV 230
HSV1GPC_12      MYYLAWGRMDSPHEYGTWVRVRMFRPPSLTLQPHAVMEGQPFKATCTAAAYYPRNPVEFV 300
HSV1GPC_7       MYYLAWGRMDSPHEYGTWVRVRMFRPPSLTLQPHAVMEGQPFKATCTAAAYYPRNPVEFV 300
HSV1GPC_19      MYYLAWGRMDSPHEYGTWVRVRMFRPPSLTLQPHAVMEGQPFKATCTATAYYPRNPVEFV 300
HSV1GPC_5       MYYLAWGRMDSPHEYGTWVRVRMFRPPSLTLQPHAVMEGQPFKATCTAAAYYPRNPVEFV 300
HSV1GPC_11      MYYLAWGRMDSPHEYGTWVRVRMFRPPSLTLQPHAVMEGQPFKATCTAAAYYPRNPVEFV 230
HSV1GPC_17      MYYLAWGRMDSPHEYGTWVRVRMFRPPSLTLQPHAVMEGQPFKATCTAAAYYPRNPVEFV 230
HSV1GPC_18      MYYLAWGRMDSPHEYGTWVRVRMFRPPSLTLQPHAVMEGQPFKATCTAAAYYPRNPVEFV 230
HSV1GPC_4       MYYLAWGRMDSPHEYGTWVRVRMFRPPSLTLQPHAVMEGQPFKATCTADAYYPRNPVEFV 300
HSV1GPC_13      MYYLAWGRMDSPHEYGTWVRVRMFRPPSLTLQPHAVMEGQPFKATCTADAYYPRNPVEFV 300
HSV1GPC_9       MYYLAWGRMDSPHEYGTWVRVRMFRPPSLTLQPHAVMEGQPFKATCTAAAYYPRNPVEFV 300
HSV1GPC_2       MYYLAWGRMDSPHEYGTWVRVRMFRPPSLTLQPHAVMEGQPFKATCTAAAYYPRNPVEFV 300
HSV1GPC_3       MYYLAWGRMDSPHEYGTWVRVRMFRPPSLTLQPHAVMEGQPFKATCTAAAYYPRNPVEFV 300
HSV1GPC_16      MYYLAWGRMDSPHEYGTWVRVRMFRPPSLTLQPHAVMEGQPFKATCTAAAYYPRNPVEFD 300
HSV2GPC_25      MYYWVWGRTDRPSAYGTWVRVRVFRPPSLTIHPHAVLEGQPFKATCTAATYYPGNRAEFV 269
HSV2GPC_27      MYYWVWGRTDRPSAYGTWVRVRVFRPPSLTIHPHAVLEGQPFKATCTAATYYPGNRAEFV 268
HSV2GPC_26      MYYWVWGRTDRPSAYGTWVRVRVFRPPSLTIHPHAVLEGQPFKATCTAATYYPGNRAEFV 269
HSV2GPC_24      MYYWVWGRTDRPSAYGTWVRVRVFRPPSLTIHPHAVLEGQPFKATCTAATYYPGNRAEFV 269
HSV2GPC_21      MYYWVWGRTDRPSAYGTWVRVRVFRPPSLTIHPHAVLEGQPFKATCTAATYYPGNRAEFV 269
HSV2GPC_23      MYYWVWGRTDRPSAYGTWVRVRVFRPPSLTIHPHAVLEGQPFKATCTAATYYPGNRAEFV 269
HSV2GPC_22      MYYWVWGRTDRPSAYGTWVRVRVFRPPSLTIHPHAVLEGQPFKATCTAATYYPGNRAEFV 199
                *** .*** * *  ********:*******::****:*********** :*** * .**

HSV1GPC_10      WFEDDRQVFNPGQIDTQTHEHPDGFTTVSTVTSEAVGGQVPPRTFTCQMTWHRDSVMFSR 360
HSV1GPC_20      WFEDDRQVFNPGQIDTQTHEHPDGFTTVSTVTSEAVGGQVPPRTFTCQMTWHRDSVMFSR 360
HSV1GPC_8       WFEDDRQVFNPGQIDTQTHEHPDGFTTVSTVTSEAVGGQVPPRTFTCQMTWHRDSVTFSR 360
HSV1GPC_14      WFEDDRQVFNXGQIDTQTHEHPDGFTTVSTVTSEAVGGQVPPRTFTCQMTWHRDSVTFSR 290
HSV1GPC_6       WFEDDRQVFNPGQIDTQTHEHPDGFTTVSTVTSEAVGGQVPPRTFTCQMTWHRDSVTFSR 290
HSV1GPC_15      WFEDDRQVFNPGQIDTQTHEHPDGFTTVSTVTSEAVGGQVPPRTFTCQMTWHRDSVTFSR 290
HSV1GPC_1       WFEDDHQVFNPGQIDTQTHEHPDGFTTVSTVTSEAVGGQVPPRTFTCQMTWHRDSVTFSR 290
HSV1GPC_12      WFEDDHQVFNPVQIDTQTHEHPDGFTTVSTVTSEAVGGQVPPRTFTCQMTWHRDSVTFSR 360
HSV1GPC_7       WFEDDHQVFNPGQIDTQTHEHPDGFTTVSTVTSEAVGGQVPPRTFTCQMTWHRDSVTFSR 360
HSV1GPC_19      WFEDDHQVFNPGQIDTQTHEHPDGFTTVSTVTSEAVGGQVPPRTFTCQMTWHRDSVTFSR 360
HSV1GPC_5       WFEDDHQVFNPGQIDTQTHEHPDGFTTVSTVTSEAVGGQVPPRTFTCQMTWHRDSVTFSR 360
HSV1GPC_11      WFEDDHQVFNPGQIDTQTHEHPDGFTTVSTVTSEAVGGQVPPRTFTCQMTWHRDSVTFSR 290
HSV1GPC_17      WFEDDHQVFNPGQIDTQTHEHPDGFTTVSTVTSEAVGGQVPPRTFTCQMTWHRDSVTFSR 290
HSV1GPC_18      WFEDDHQVFNPGQIDTQTHEHPDGFTTVSTVTSEAVGGQVPPRTFTCQMTWHRDSVTFSR 290
HSV1GPC_4       WFEDDHQVFNPGQIDTQTHEHPDGFTTVSTVTSEAVGGQVPPRTFTCQMTWHRDSVTFSR 360
HSV1GPC_13      WFEDDHQVFNPGQIDTQTHEHPDGFTTVSTVTSEAVGGQVPPRTFTCQMTWHRDSVTFSR 360
HSV1GPC_9       WFEDDHQVFNPGQIDTQTHEHPDGFTTVSTVTSEAVGGQVPPRTFTCQITWHRDSVTFSR 360
HSV1GPC_2       WFEDDHQVFNPGQIDTQTHEHPDGFTTVSTVTSEAVGGQVPPRTFTCQMTWHRDSVTFSR 360
HSV1GPC_3       WFEDDHQVFNPGQIDTQTHEHPDGFTTVSTVTSEAVGGQVPPRTFTCQMTWHRDSVTFSR 360
HSV1GPC_16      WFEDDRQVFNPGQIDTQTHEHPDGFTTVSTVTSEAVGGQVPPRTFTCQMTWHRDSVTFSR 360
HSV2GPC_25      WFEDGRRVFDPAQIHTQTQENPDGFSTVSTVTSAAVGGQGPPRTFTCQLTWHRDSVSFSR 329
HSV2GPC_27      WFEDGRRVFDPAQIHTQTQENPDGFSTVSTVTSAAVGGQGPPRTFTCQLTWHRDSVSFSR 328
HSV2GPC_26      WFEDGRRVFDPAQIHTQTQENPDGFSTVSTVTSAAVGGQGPPRTFTCQLTWHRDSVSFSR 329
HSV2GPC_24      WFEDGRRVFDPAQIHTQTQENPDGFSTVSTVTSAAVGGQGPPRTFTCQLTWHRDSVSFSR 329
HSV2GPC_21      WFEDGRRVFDPAQIHTQTQENPDGFSTVSTVTSAAVGGQGPPRTFTCQLTWHRDSVSFSR 329
HSV2GPC_23      WFEDGRRVFDPAQIHTQTQENPDGFSTVSTVTSAAVGGQGPPRTFTCQLTWHRDSVSFSR 329
HSV2GPC_22      WFEDGRRVFDPAQIHTQTQENPDGFSTVSTVTSAAVGGQGPPRTFTCQLTWHRDSVSFSR 259
                ****.::**:  **.***:*:****:******* ***** ********:******* ***

HSV1GPC_10      RNATGLALVLPRPTITMEFGVRHVVCTAGCVPEGVTFAWFLGDDPSPAAKSAVTAQESCD 420
HSV1GPC_20      RNATGLALVLPRPTITMEFGVRHVVCTAGCVPEGVTFAWFLGDDPSPAAKSAVTAQESCD 420
HSV1GPC_8       RNATGLALVLPRPTITMEFGVRHVVCTAGCVPEGVTFAWFLGDDPSPAAKSAVTAQESCD 420
HSV1GPC_14      RNATGLALVLPRPTITMEFGVRHVVCTAGCVPEGVTFAWFLGDDPSPAAKSAVTAQESCD 350
HSV1GPC_6       RNATGLALVLPRPTITMEFGVRHVVCTAGCVPEGVTFAWFLGDDPSPAAKSAVTAQESCD 350
HSV1GPC_15      RNATGLALVLPRPTITMEFGVRHVVCTAGCVPEGVTFAWFLGDDPSPAAKSAVTAQESCD 350
HSV1GPC_1       RNATGLALVLPRPTITMEFGVRIVVCTAGCVPEGVTFAWFLGDDPSPAAKSAVTAQESCD 350
HSV1GPC_12      RNATGLALVLPRPTITMEFGVRHVVCTAGCVPEGVTFAWFLGDDPSPAAKSAVTAQESCD 420
HSV1GPC_7       RNATGLALVLPRPTITMEFGVRHVVCTAGCVPEGVTFAWFLGDDPSPAAKSAVTAQESCD 420
HSV1GPC_19      RNATGLALVLPRPTITMEFGVRHVVCTAGCVPEGVTFAWFLGDDPSPAAKSAVTAQESCD 420
HSV1GPC_5       RNATGLALVLPRPTITMEFGVRHVVCTAGCVPEGVTFAWFLGDDPSPAAKSAVTAQESCD 420
HSV1GPC_11      RNATGLALVLPRPTITMEFGVRHVVCTAGCVPEGVTFAWFLGDDPSPAAKSAVTAQESCD 350
HSV1GPC_17      RNATGLALVLPRPTITMEFGVRHVVCTAGCVPEGVTFAWFLGDDPSPAAKSAVTAQESCD 350
HSV1GPC_18      RNATGLALVLPRPTITMEFGVRHVVCTAGCVPEGVTFAWFLGDDPSPAAKSAVTAQESCD 350
HSV1GPC_4       RNATGLALVLPRPTITMEFGVRHVVCTAGCVPEGVTFAWFLGDDPSPAAKSAVTAQESCD 420
HSV1GPC_13      RNATGLALVLPRPTITMEFGVRHVVCTAGCVPEGVTFAWFLGDDPSPAAKSAVTAQESCD 420
HSV1GPC_9       RNATGLALVLPRPTITMEFGVRHVVCTAGCVPEGVTFAWFLGDDPSPAAKSAVTAQESCD 420
HSV1GPC_2       RNATGLALVLPRPTITMEFGVRHVVCTAGCVPEGVTFAWFLGDDPSPAAKSAVTAQESCD 420
HSV1GPC_3       RNATGLALVLPRPTITMEFGVRHVVCTAGCVPEGVTFAWFLGDDPSPAAKSAVTAQESCD 420
HSV1GPC_16      RNATGLALVLPRPTITMEFGVRHVVCTAGCVPEGVTFAWFLGDDPSPAAKSAVTAQESCD 420
HSV2GPC_25      RNASGTASVLPRPTITMEFTGDHAVCTAGCVPEGVTFAWFLGDDSSPAEKVAVASQTSCG 389
HSV2GPC_27      RNASGTASVLPRPTITMEFTGDHAVCTAGCVPEGVTFAWFLGDDSSPAEKVAVASQTSCG 388
HSV2GPC_26      RNASGTASVLPRPTITMEFTGDHAVCTAGCVPEGVTFAWFLGDDSSPAEKVAVASQTSCG 389
HSV2GPC_24      RNASGTASVLPRPTITMEFTGDHAVCTAGCVPEGVTFAWFLGDDSSPAEKVAVASQTSCG 389
HSV2GPC_21      RNASGTASVLPRPTITMEFTGDHAVCTAGCVPEGVTFAWFLGDDSSPAEKVAVASQTSCG 389
HSV2GPC_23      RNASGTASVLPRPTITMEFTGDHAVCTAGCVPEGVTFAWFLGDDSSPAEKVAVASQTSCG 389
HSV2GPC_22      RNASGTASVLPRPTITMEFTGDHAVCTAGCVPEGVTFAWFLGDDSSPAEKVAVASQTSCG 319
                ***:* * ***********    .********************.*** * **::* **.

HSV1GPC_10      HPGLATVRSTLPISYDYSEYICRLTGYPAGIPVLEHHGSHQPPPRDPTERQVIEAIEWVG 480
HSV1GPC_20      HPGLATVRSTLPISYDYSEYICRLTGYPAGIPVLEHHGSHQPPPRDPTERQVIEAIEWVG 480
HSV1GPC_8       HPGLATVRSTLPISYDYSEYICRLTGYPAGIPVLEHHGSHQPPPRDPTERQVIEAIEWVG 480
HSV1GPC_14      HPGLATVRSTLPISYDYSEYICRLTGYPAGIPVLEHHGSHQPPPRDPTERQVIEAIEWVG 410
HSV1GPC_6       HPGLATVRSTLPISYDYSEYICRLTGYPAGIPVLEHHGSHQPPPRDPTERQVIEAIEWVG 410
HSV1GPC_15      HPGLATVRSTLPISYDYSEYICRLTGYPAGIPVLEHHGSHQPPPRDPTERQVIEAIEWVG 410
HSV1GPC_1       HPGLATVRSTLPISYDYSEYICRLTGYPAGIPVLEHHGSHQPPPRDPTERQVIEAIEWVG 410
HSV1GPC_12      HPGLATVRSTLPISYDYSEYICRLTGYPAGIPVLEHHGSHQPPPRDPTERQVIEAIEWVG 480
HSV1GPC_7       RPGLATVRSTLPISYDYSEYICRLTGYPAGIPVLEHHGSHQPPPRDPTERQVIEAIEWVG 480
HSV1GPC_19      RPGLATVRSTLPISYDYSEYICRLTGYPAGIPVLEHHGSHQPPPRDPTERQVIEAIEWVG 480
HSV1GPC_5       RPGLATVRSTLPISYDYSEYICRLTGYPAGIPVLEHHGSHQPPPRDPTERQVIEAIEWVG 480
HSV1GPC_11      RPGLATVRSTLPISYDYSEYICRLTGYPAGIPVLEHHGSHQPPPRDPTERQVIEAIECVG 410
HSV1GPC_17      RPGLATVRSTLPISYDYSEYICRLTGYPAGIPVLEHHGSHQPPPRDPTERQVIEAIEWVG 410
HSV1GPC_18      RPGLATVRSTLPISYDYSEYICRLTGYPAGIPVLEHHGSHQPPPRDPTERQVIEAIEWVG 410
HSV1GPC_4       HPGLATVRSTLPISYDYSEYICRLTGYPAGIPVLEHHGSHQPPPRDPTERQVIEAIEWVG 480
HSV1GPC_13      HPGLATVRSTLPISYDYSEYICRLTGYPAGIPVLEHHGSHQPPPRDPTERKVIEAIEWVG 480
HSV1GPC_9       HPGLATVRSTLPISYDYSEYICRLTGYPAGIPVLEHHGSHQPPPRDPTERQVIEAIEWVG 480
HSV1GPC_2       HPGLATVRSTLPISYDYSEYICRLTGYPAGIPVLEHHGSHQPPPRDPTERQVIEAIEWVG 480
HSV1GPC_3       HPGLATVRSTLPISYDYSEYICRLTGYPAGIPVLEHHGSHQPPPRDPTERQVIEAIEWVG 480
HSV1GPC_16      HPGLATVRSTLPISYDYSEYICRLTGYPAGIPVLEHHGSHQPPPRDPTERQVIEAIEWVG 480
HSV2GPC_25      RPGTATIRSTLPVSYEQTEYICRLAGYPDGIPVLEHHGSHQPPPRDPTERQVIRAVEGAG 449
HSV2GPC_27      RPGTATIRSTLPVSYEQTEYICRLAGYPDGIPVLEHHGSHQPPPRDPTERQVIRAVEGAG 448
HSV2GPC_26      RPGTATIRSTLPVSYEQTEYICRLAGYPDGIPVLEHHGSHQPPPRDPTERQVIRAVEGAG 449
HSV2GPC_24      RPGTATIRSTLPVSYEQTEYICRLAGYPDGIPVLEHHGSHQPPPRDPTERQVIRAVEGAG 449
HSV2GPC_21      RPGTATIRSTLPVSYEQTEYICRLAGYPDGIPVLEHHGSHQPPPRDPTERQVIRAVEGAG 449
HSV2GPC_23      RPGTATIRSTLPVSYEQTEYICRLAGYPHGIPVLEHHGSHQPPPRDPTERQVIRAVEGAG 449
HSV2GPC_22      RPGTATIRSTLPVSYEQTEYICRLAGYPDGIPVLEHHGSHQPPPRDPTKRQVIRAVEGAG 379
                :** **:*****:**: :******:*** *******************:*:**.*:* .*

HSV1GPC_10      IGIGVLAAGVLVVTAIVYVVRTSQSRQRHRR 511
HSV1GPC_20      IGIGVLAAGVLVVTAIVYVVRTSQSRQRHRR 511
HSV1GPC_8       IGIGVLAAGVLVVTAIVYVVRTSQSRQRHRR 511
HSV1GPC_14      IGIGVLAAGVLVVTAIVYVVRTSQSRQRHRR 441
HSV1GPC_6       IGIGVLAAGVLVVTAIVYVVRTSQSRQRHRR 441
HSV1GPC_15      IGIGVLAAGVLVVTAIVYVVRTSQSRQRHRR 441
HSV1GPC_1       IGIGVLAAGVLVVTAIVYVVRTSQSRQRHRR 441
HSV1GPC_12      IGIGVLAAGVLVVTAIVYVVRTSQSRQRHRR 511
HSV1GPC_7       IGIGVLAAGVLVVTAIVYVVRTSQSRQRHRR 511
HSV1GPC_19      IGIGVLAAGVLVVTAIVYVVRTSQSRQRHRR 511
HSV1GPC_5       IGIGVLAAGVLVVTAIVYVVRTSQSRQRHRR 511
HSV1GPC_11      IGIGVLAAGVLVVTAIVYVVRTSQSRQRHRR 441
HSV1GPC_17      IGIGVLAAGVLVATAIVYVVRTSQSRQRHRR 441
HSV1GPC_18      IGIGVLAAGVLVVTAIVYVVRTSQSRQRHRR 441
HSV1GPC_4       IGIGVLAAGVLVVTAIVYVVRTSQSRQRHRR 511
HSV1GPC_13      IGIGVLAAGVLVVTAIVYVVRTSQSRQRHRR 511
HSV1GPC_9       IGIGVLAAGVLVVTAIVYVVRTSQSRQRHRR 511
HSV1GPC_2       IGIGVLAAGVLVVTAIVYVVRTSQSRQRHRR 511
HSV1GPC_3       IGIGVLAAGVLVVTAIVYVVRTSQSRQRHRR 511
HSV1GPC_16      IGIGVLAAGVLVVTAIVYVVRTSQSRQRHRR 511
HSV2GPC_25      IGVAVLVAVVLAGTAVVYLTHASSVRYRRLR 480
HSV2GPC_27      IGVAVLVAVVLAGTAVVYLTHASSVRYRRLR 479
HSV2GPC_26      IGVAVLVAVVLAGTAVVYLTHASSVRYRRLR 480
HSV2GPC_24      IGVAVLVAVVLAGTAVVYLTHASSVRYRRLR 480
HSV2GPC_21      IGVAVLVAVVLAGTAVVYLTHASSVRYRRLR 480
HSV2GPC_23      IGVAVLVAVVLAGTAVVYLTHASSVRYRRLR 480
HSV2GPC_22      IGVAVLVAVVLAGTAVVYLTHASSVRYRRLR 410
                **:.**.* **. **:**:.::*. * *: *
```
