## Supplementary file 9 for "Contriving a chimeric polyvalent vaccine to prevent infections caused by Herpes Simplex Virus (Type-1 and Type-2): an exploratory immunoinformatic approach"

```
Alignment  GLYCOPROTEIN D

HSV1GPD_1       MGGAAARLGAVILFVVIVGLHGVRSKYALVDASLKMADPNRFRGKDLPVLDQLTDPPGVR 60
HSV1GPD_2       MGGAAARLGAVILFVVIVGLHGVRGKYALVDASLKMADPNRFRGKDLPVLDQLTDPPGVR 60
HSV1GPD_4       MGGAAARLGAVILFVVIVGLHGVRGKYALADASLKMADPNRFRGKDLPVLDQLTDPPGVR 60
HSV1GPD_8       MGGAAARLGAVILFVVLVGLHGVRGKYALADASLKMADPNRFRGKDLPVLDQLTDPPGVR 60
HSV1GPD_10      MGGAAARLGAVILFVVIVGLHGVRGKYALADASLKMADPNRFRGKDLPVLDQLTDPPGVR 60
HSV1GPD_16      MGGAAARLGAVILFVVIVGLHGVRGKYALADASLKMADPNRFRGKDLPVLDQLTDPPGVR 60
HSV1GPD_14      MGGTAARLGAVILFVVIVGLHGVRGKYALADASLKMADPNRFRGKDLPVLDQLTDPPGVR 60
HSV1GPD_13      MGGAAARLGAVILFVVIVGLHGVRGKYALADASLKMADPNRFRGKDLPVLDRLTDPPGVR 60
HSV1GPD_18      MGGAAARLGAVILFVVIVGLHGVRGKYALADASLKMADPNRFRGKDLPVLDPLTDPPGVR 60
HSV1GPD_17      MGGAAARLGAVILFVVIVGLHGVRGKYALADASLKMADPNRFRGKDLPVPDQLTDPPGVR 60
HSV1GPD_20      MGGAAARLGAVILFVVIVGLHGVRGKYALADASLKMADPNRFRGKDLPVPDRLTDPPGVR 60
HSV1GPD_5       MGGAAARLGAVILFVVIVGLHGVRGKYALVDASLKMADPNRFRGKDLPVLDQLTDPPGVR 60
HSV1GPD_11      MGGAAARLGAVILFVVIVGLHGVRGKYALADASLKMADPNRFRGKDLPVLDQLTDPPGVR 60
HSV1GPD_12      MGGAAARLGAVILFVVIVGLHGVRGKYALADASLKMADPNRFRGKDLPVLDQLTDPPGVR 60
HSV1GPD_6       MGGAAARLGAVILFVVIVGLHGVRGKYALADASLKMADPNRFRGKDLPVLDQLTDPPGVR 60
HSV1GPD_7       MGGAAARLGAVILFVVIVGLHGVRGKYALADASLKMADPNRFRGKDHPVLDQLTDPPGVR 60
HSV1GPD_9       MGGAAARLGAVILFVVIVGLHGVRGKYALADASLKMADPNRFRGKDLPVLDQLTDPPGVR 60
HSV1GPD_19      MGGAAARLGAVILFVVIVGLHGVRGKYALADASLKMADPNRFRGKDLPVPDRLTDPPGVR 60
HSV1GPD_15      MGGAAARLGAVILFVVIVGLHGVRGKYALADASLKLADPNRFRRKDLPVLDQLTDPPGVR 60
HSV1GPD_3       ------------------------------------------------------------
HSV2GPD_22      MGRLTSGVGTAALLVVAVGLRVVCAKYALADPSLKMADPNRFRGKNLPVLDQLTDPPGVK 60
HSV2GPD_24      MGRLTSGVGTAALLVVAVGLRVVCAKYALADPSLKMADPNRFRGKNLPVLDQLTDPPGVK 60
HSV2GPD_26      MGRLTSGVGTAALLVVAVGLRVVCAKYALADPSLKMADPNRFRGKNLPVLDQLTDPPGVK 60
HSV2GPD_21      MGRLTSGVGTAALLVVAVGLRVVCAKYALADPSLKMADPNRFRGKNLPVLDQLTDPPGVK 60
HSV2GPD_25      MGRLTSGVGTAALLVVAVGLRVVCAKYALADPSLKMADPNRFRGKNLPVLDRLTDPPGVK 60
HSV2GPD_23      MGRLTSGVGTAALLVVAVGLRVVCAKYALADPSLKMADPNRFRGKNLPVLDQLTDPPGVK 60
HSV2GPD_27      MGRLTSGVGTAALLVVAVGLRVVCAKYALADPSLKMADPNRFRGKNLPVLDQLTDPPGVK 60
HSV2GPD_28      MGRLTSGVGTAALLVVAVGLRVVCAKYALADPSLKMADPNRFRGKNLPVLDQLTDPPGVK 60


HSV1GPD_1       RVYHIQAGLPDPFQPPSLPITVYYAVLERACRSVLLNAPSEAPQIVRGASEDVRKQPYNL 120
HSV1GPD_2       RVYHIQAGLPDPFQPPSLPITVYYAVLERACRSVLLNAPSEAPQIVRGASEDVRKQPYNL 120
HSV1GPD_4       RVYHIQAGLPDPFQPPSLPITVYYAVLERACRSVLLNAPSEAPQIVRGASEDVRKQPYNL 120
HSV1GPD_8       RVYHIQAGLPDPFQPPSLPITVYYAVLERACRSVLLNAPSEAPQIVRGASEDVRKQPYNL 120
HSV1GPD_10      RVYHIQAGLPDPFQPPSLPITVYYAVLERACRSVLLNAPSEAPQIVRGASEDVRKQPYNL 120
HSV1GPD_16      RVYHIQAGLPDPFQPPSLPITVYYAVLERACRSVLLNAPSEAPQIVRGASEDVRKQPYNL 120
HSV1GPD_14      RVYHIQAGLPDPFQPPSLPITVYYAVLERACRSVLLNAPSEAPQIVRGASEDVRKQPYNL 120
HSV1GPD_13      RVYHIQAGLPDPFQPPSLPITVYYAVLERACRSVLLNAPSEAPQIVRGASEDVRKQPYNL 120
HSV1GPD_18      RVYHIQAGLPDPFQPPSLPITVYYAVLERACRSVLLNAPSEAPQIVRGASEDVRKQPYNL 120
HSV1GPD_17      RVYHIQAGLPDPFQPPSLPITVYYAVLERACRSVLLNAPSEAPQIVRGASEDVRKQPYNL 120
HSV1GPD_20      RVYHIQAGLPDPFQPPSLPITVYYAVLERACRSVLLNAPSEAPQIVRGASEDVRKQPYNL 120
HSV1GPD_5       RVYHIQAGLPDPFQPPSLPITVYYAVLERACRSVLLNAPSEAPQIVRGASEDVRKQPYNL 120
HSV1GPD_11      RVYHIQAGLPDPFQPPSLPITVYYAVLERACRSVLLNAPSEAPQIVRGASEDVRKQLYNL 120
HSV1GPD_12      RVYHIQAGLPDPFQPPSLPITVYYAVLERACRSVLLNAPSEAPQIVRGASEDVRKQPYNL 120
HSV1GPD_6       RVYHIQAGLPDPFQPPSLPITVYYAVLERACRSVLLNAPSEAPQIVRGASEDVRKQPYNL 120
HSV1GPD_7       RVYHIQAGLPDPFQPPSLPITVYYAVLERACRSVLLNAPSEAPQIVRGASEDVRKQPYNL 120
HSV1GPD_9       RVYHIQAGLPDPFQPPSLPITVYYAVLERACRSVLLNAPSEAPQIVRGASEDVRKQPYNL 120
HSV1GPD_19      RVYHIQAGLPDPFQPPSLPITVYYAVLERACRSVLLNAPSEAPQIVRGASEDVRKQPYNL 120
HSV1GPD_15      RVYHIQAGLPDPFQPPSLPITVYYAVLERACRSVLLNAPSEAPQIVRGASEDVRKQPYNL 120
HSV1GPD_3       ----------DPFQPPSLPITVYYAVLERACRSVLLNAPSEAPQIVRGASEDVRKQPYNL 50
HSV2GPD_22      RVYHIQPSLEDPFQPPSIPITVYYAVLERACRSVLLHAPSEAPQIVRGASDEARKHTYNL 120
HSV2GPD_24      RVYHIQPSLEDPFQPPSIPITVYYAVLERACRSVLLHAPSEAPQIVRGASDEARKHTYNL 120
HSV2GPD_26      RVYHIQPSLEDPFQPPSIPITVYYAVLERACRSVLLHAPSEAPQIVRGASDEARKHTYNL 120
HSV2GPD_21      RVYHIQPSLEDPFQPPSIPITVYYAVLERACRSVLLHAPSEAPQIVRGASDEARKHTYNL 120
HSV2GPD_25      RVYHIQPSLEDPFQPPSIPITVYYAVLERACRSVLLHAPSEAPQIVRGASDEARKHTYNL 120
HSV2GPD_23      RVYHIQPSLEDPFQPPSIPITVYYAVLERACRSVLLHAPSEAPQIVRGASDEARKHTYNL 120
HSV2GPD_27      RVYHIQPSLEDPFQPPSIPITVYYAVLERACRSVLLHAPSEAPQIVRGASDEARKHTYNL 120
HSV2GPD_28      RVYHIQPSLEDPFQPPSIPITVYYAVLERACRSVLLHAPSEAPQIVRGASDEARKHTYNL 120
                          *******:******************:*************::.**: ***

HSV1GPD_1       TIAWFRMGGNCAIPITVMEYTECSYNKSLGACPIRTQPRWNYYDSFSAVSEDNLGFLMHA 180
HSV1GPD_2       TIAWFRMGGNCAIPITVMEYTECSYNKSLGACPIRTQPRWNYYDSFSAVSEDNLGFLMHA 180
HSV1GPD_4       TIAWFRMGGNCAIPITVMEYTECSYNKSLGACPIRTQPRWNYYDSFSAVSEDNLGFLMHA 180
HSV1GPD_8       TIAWFRMGGNCAIPITVMEYTECSYNKSLGACPIRTQPRWNYYDSFSAVSEDNLGFLMHA 180
HSV1GPD_10      TIAWFRMGGNCAIPITVMEYTECSYNKSLGACPIRTQPRWNYYDSFSAVSEDNLGFLMHA 180
HSV1GPD_16      TIAWFRMGGNCAIPITVMEYTECSYNKSLGACPIRTQPRWNYYDSFSAVSEDNLGFLMHA 180
HSV1GPD_14      TIAWFRMGGNCAIPITVMEYTECSYNKSLGACPIRTQPRWNYYDSFSAVSEDNLGFLMHA 180
HSV1GPD_13      TIAWFRMGGNCAIPITVMEYTECSYNKSLGACPIRTQPRWNYYDSFSAVSEDNLGFLMHA 180
HSV1GPD_18      TIAWFRMGGNCAIPITVMEYTECSYNKSLGACPIRTQPRWNYYDSFSAVSEDNLGFLMHA 180
HSV1GPD_17      TIAWFRMGGNCAIPITVMEYTECSYNKSLGACPIRTQPRWNYYDSFSAVSEDNLGFLMHA 180
HSV1GPD_20      TIAWFRMGGNCAIPITVMEYTECSYNKSLGACPIRTQPRWNYYDSFSAVSEDNLGFLMHA 180
HSV1GPD_5       TIAWFRMGGNCAIPITVMEYTECSYNKSLGACPIRTQPRWNYYDSFSAVSEDNLGFLMHA 180
HSV1GPD_11      TIAWFRMGGNCAIPITVMEYTDCSYNKSLGACPIRTQPRWNYYDSFSAVSEDNLGFLMHA 180
HSV1GPD_12      TIAWFRMGGNCAIPITVMEYTECSYNKSLGACPIRTQPRWNYYDSFSAVSEDNLGFLMHA 180
HSV1GPD_6       TIAWFRMGGNCAIPITVMEYTECSYNKSLGACPIRTQPRWNYYDSFSAVSEDNLGFLMHA 180
HSV1GPD_7       TIAWFRMGGNCAIPITVMEYTECSYNKSLGACPIRTQPRWNYYDSFSAVSEDNLGFLMHA 180
HSV1GPD_9       TIAWFRMGGNCAIPITVMEYTECSYNKSLGACPIRTQPRWNYYDSFSAVSEDNLGFLMHA 180
HSV1GPD_19      TIAWFRMGGNCAIPITVMEYTECSYNKSLGACPIRTQPRWNYYDSFSAVSEDNLGFLMHA 180
HSV1GPD_15      TIAWFRMGGNCAIPITVMEYTECSYNKSLGACPIRTQPRWNYYDSFSAVSEDNLGFLMHA 180
HSV1GPD_3       TIAWFRMGGNCAIPITVMEYTECSYNKSLGACPIRTQPRWNYYDSFSAVSEDNLGFLMHA 110
HSV2GPD_22      TIAWYRMGDNCAIPITVMEYTECPYNKSLGVCPIRTQPRWSYYDSFSAVSEDNLGFLIHA 180
HSV2GPD_24      TIAWYRMGDNCAIPITVMEYTECPYNKSLGVCPIRTQPRWSYYDSFSAASEDNLGFLMHA 180
HSV2GPD_26      TIAWYRMGDNCAIPITVMEYTECPYNKSLGVCPIRTQPRWSYYDSFSAVSEDTLGFLMHA 180
HSV2GPD_21      TIAWYRMGDNCAIPITVMEYTECPYNKSLGVCPIRTQPRWSYYDSFSAVSEDNLGFLMHA 180
HSV2GPD_25      TIAWYRMGDNCAIPITVMEYTECPYNKSLGVCPIRTQPRWSYYDSFSAVSEDNLGFLMHA 180
HSV2GPD_23      TIAWYRMGDNCAIPITVMEYTECPYNKSLGVCPIRTQPRWSYYDSFSAVSEDNLGFLMHA 180
HSV2GPD_27      TIAWYRMGDNCAIPITVMEYTECPYNKSLGVCPIRTQPRWSYYDSFSAVSEDNLGFLMHA 180
HSV2GPD_28      TIAWYRMGDNCAIPITVMEYTECPYNKSLGVCPIRTQPRWSYYDSFSAVSEDNLGFLMHA 180
                ****:***.************:*.******.*********.*******.***.****:**

HSV1GPD_1       PAFETAGTYLRLVKINDWTEITQFILEHRAKGSCKYALPLRIPPSACLSPQAYQQGVTVD 240
HSV1GPD_2       PAFETAGTYLRLVKINDWTEITQFILEHRAKGSCKYALPLRIPPSACLSPQAYQQGVTVD 240
HSV1GPD_4       PAFETAGTYLRLVKINDWTEITQFILEHRAKGSCKYALPLRIPPSACLSPQAYQQGVTVD 240
HSV1GPD_8       PAFETAGTYLRLVKINDWTEITQFILEHRAKGSCKYALPLRIPPSACLSPQAYQQGVTVD 240
HSV1GPD_10      PAFETAGTYLRLVKINDWTEITQFILEHRAKGSCKYALPLRIPPSACLSPQAYQQGVTVD 240
HSV1GPD_16      PAFETAGTYLRLVKINDWTEITQFILEHRAKGSCKYALPLRIPPSACLSPQAYQQGVTVD 240
HSV1GPD_14      PAFETAGTYLRLVKINDWTEITQFILEHRAKGSCKYALPLRIPPSACLSPQAYQQGVTVD 240
HSV1GPD_13      PAFETAGTYLRLVKINDWTEITQFILEHRAKGSCKYALPLRIPPSACLSPQAYQQGVTVD 240
HSV1GPD_18      PAFETAGTYLRLVKINDWTEITQFILEHRAKGSCKYALPLRIPPSACLSPQAYQQGVTVD 240
HSV1GPD_17      PAFETAGTYLRLVKINDWTEITQFILEHRAKGSCKYALPLRIPPSACLSPQAYQQGVTVD 240
HSV1GPD_20      PAFETAGTYLRLVKINDWTEITQFILEHRAKGSCKYALPLRIPPSACLSPQAYQQGVTVD 240
HSV1GPD_5       PAFETAGTYLRLVKINDWTEITQFILEHRAKGSCKYALPLRIPPSACLSPQAYQQGVTVD 240
HSV1GPD_11      PAFETAGTYLRLVKINDWTEITQFILEHRAKGSCKYALPLRIPPSACLSPQAYQQGVTVD 240
HSV1GPD_12      PAFETAGTYLRLVKINDWTEITQFILEHRAKGSCKYALPLRIPPSACLSPQAYQQGVTVD 240
HSV1GPD_6       PAFETAGTYLRLVKINDWTEITQFILEHRAKGSCKYALPLRIPPSACLSPQAYQQGVTVD 240
HSV1GPD_7       PAFETAGTYLRLVKINDWTEITQFILEHRAKGSCKYALPLRIPPSACLSPQAYQQGVTVD 240
HSV1GPD_9       PAFETAGTYLRLVKINDWTEITQFILEHRAKGSCKYAIPLRIPPSACLSPQAYQQGVTVD 240
HSV1GPD_19      PAFETAGTYLRLVKINDWTEITQFILEHRAKGSCKYALPLRIPPSACLSPQAYQQGVTVD 240
HSV1GPD_15      PAFETAGTYLRLVKINDWTEITQFILEHRAKGSCKYALPLRIPPSACLSPQAYQQGVTVD 240
HSV1GPD_3       PAFETAGTYLRLVKINDWTEITQFILEHRAKGSCKYALPLRIPPSACLSPQAYQQGVTVD 170
HSV2GPD_22      PAFETAGTYLRLVKINDWTEITQFILEHRARASCKYALPLRIPPAACLTSKAYQQGVTVD 240
HSV2GPD_24      PAFETAGTYLRLVKINDWTEITQFILEHRARASCKYALPLRIPPAACLTSKAYQQGVTVD 240
HSV2GPD_26      PAFETAGTYLRLVKINDWTEITQFILEHRARASCKYALPLRIPPAACLTSKAYQQGVTVD 240
HSV2GPD_21      PAFETAGTYLRLVKINDWTEITQFILEHRARASCKYALPLRIPPAACLTSKAYQQGVTVD 240
HSV2GPD_25      PAFETAGTYLRLVKINDWTEITQFILEHRARASCKYALPLRIPPAACLTSKAYQQGVTVD 240
HSV2GPD_23      PAFETAGTYLRLVKINDWTEITQFILEHRARASCKYALPLRIPPAACLTSKAYQQGVTVD 240
HSV2GPD_27      PAFETAGTYLRLVKINDWTEITQFILEHRARASCKYALPLRIPPAACLTSKAYQQGVTVD 240
HSV2GPD_28      PAFETAGTYLRLVKINDWTEITQFILEHRARASCKYALPLRIPPAACLTSKAYQQGVTVD 240
                ******************************:.*****:******:***:.:*********

HSV1GPD_1       SIGMLPRFIPENQRTVAVYSLKIAGWHGPKAPYTSTLLPPELSETPNATQPELAPEDPED 300
HSV1GPD_2       SIGMLPRFIPENQRTVAVYSLKIAGWHGPKAPYTSTLLPPELSETPNATQPELAPEDPED 300
HSV1GPD_4       SIGMLPRFIPENQRTVAVYSLKIAGWHGPKAPYTSTLLPPELSETPNATQPELAPEDPED 300
HSV1GPD_8       SIGMLPRFIPENQRTVAVYSLKIAGWHGPKAPYTSTLLPPELSETPNATQPELAPEDPED 300
HSV1GPD_10      SIGMLPRFIPENQRTVAVYSLKIAGWHGPKAPYTSTLLPPELSETPNATQPELAPEDPED 300
HSV1GPD_16      SIGMLPRFIPENQRTVAVYSLKIAGWHGPKAPYTSTLLPPELSETPNATQPELAPEDPED 300
HSV1GPD_14      SIGMLPRFIPENQRTVAVYSLKIAGWHGPKAPYTSTLLPPELSETPNATQPELAPEDPED 300
HSV1GPD_13      SIGMLPRFIPENQRTVAVYSLKIAGWHGPKAPYTSTLLPPELSETPNATQPELAPEDPED 300
HSV1GPD_18      SIGMLPRFIPENQRTVAVYSLKIAGWHGPKAPYTSTLLPPELSETPNATQPELAPEDPED 300
HSV1GPD_17      SIGMLPRFIPENQRTVAVYSLKIAGWHGPKAPYTSTLLPPELSETPNATQPELAPEDPED 300
HSV1GPD_20      SIGMLPRFIPENQRTVAVYSLKIAGWHGPKAPYTSTLLPPELSETPNATQPELAPEDPED 300
HSV1GPD_5       SIGMLPRFIPENQRTVAVYSLKIAGWHGPKAPYTSTLLPPELSETPNATQPELAPEDPED 300
HSV1GPD_11      SIGMLPRFIPENQRTVAVYSLKIAGWHGPKAPYTSTLLPPELSETPNATQPELAPEDPED 300
HSV1GPD_12      SIGMLPRFIPENQRTVAVYSLKIAGWHGPKAPYTSTLLPPELSETPNATQPELAPEDPED 300
HSV1GPD_6       SIGMLPRFIPENQRTVAVYSLKIAGWHGPKAPYTSTLLPPELSETPNATQPELTPEDPED 300
HSV1GPD_7       SIGMLPRFIPENQRTVAVYSLKIAGWHGPKAPYTSTLLPPELSETPNATQPELAPEDPED 300
HSV1GPD_9       SIGMLPRFIPENQRTVAVYSLKIAGWHGPKAPYTSTLLPPELSETPNATQPELAPEDPED 300
HSV1GPD_19      SIGMLPRFIPENQRTVAVYSLKIAGWHGPKAPYTSTLLPPELSETPNATQPELAPEDPED 300
HSV1GPD_15      SIGMLPRFIPENQRTVAVYSLKIAGWHGPKAPYTSTLLPPELSETPNATQPELAPEAPED 300
HSV1GPD_3       SIGMLPRFIPENQRTVAVYSLKIAGWHGPKAPYTSTLLPPELSETPNATQPELAPEDPED 230
HSV2GPD_22      SIGMLPRFIPENQRTVALYSLKIAGWHGPKPPYTSTLLPPELSDTTNATQPELVPEDPED 300
HSV2GPD_24      SIGMLPRFIPENQRTVALYSLKIAGWHGPKPPYTSTLLPPELSDTTNATQPELVPEDPED 300
HSV2GPD_26      SIGMLPRFIPENQRTVALYSLKIAGWHGPKPPYTSTLLPPELSDTTNATQPELVPEDPED 300
HSV2GPD_21      SIGMLPRFIPENQRTVALYSLKIAGWHGPKPPYTSTLLPPELSDTTNATQPELVPEDPED 300
HSV2GPD_25      SIGMLPRFIPENQRTVALYSLKIAGWHGPKPPYTSTLLPPELSDTTNATQPELVPEDPED 300
HSV2GPD_23      SIGMLPRFIPENQRTVALYSLKIAGWHGPKPPYTSTLLPPELSDTTNATQPELVPEDPED 300
HSV2GPD_27      SIGMLPRFIPENQRTVALYSLKIAGWHGPKPPYTSTLLPPELSDTTNATQPELVPEDPED 300
HSV2GPD_28      SIGMLPRFTPENQRTVALYSLKIAGWHGPKPPYTSTLLPPELSDTTNATQPELVPEDPED 300
                ******** ********:************.************:*.*******.** ***

HSV1GPD_1       SALLEDPVGTVAPQIPPNWHIPSIQDAATPYHPPATPNNMGLIAGAVGGSLLAALVICGI 360
HSV1GPD_2       SALLEDPVGTVAPQIPPNWHIPSIQDAATPYHPPATPNNMGLIAGAVGGSLLAALVICGI 360
HSV1GPD_4       SALLEDPVGTVAPQIPPNWHIPSIQDAATPYHPPATPNNMGLIAGAVGGSLLAALVICGI 360
HSV1GPD_8       SALLEDPVGTVAPQIPPNWHIPSIQDAATPYHPPATPNNMGLIAGAVGGSLLAALVICGI 360
HSV1GPD_10      SALLEDPVGTVAPQIPPNWHIPSIQDAATPYHPPATPNNMGLIAGAVGGSLLAALVICGI 360
HSV1GPD_16      SALLEDPVGTVALQIPPNWHIPSIQDAATPYHPPATPNNMGLIAGAVGGSLLAALVICGI 360
HSV1GPD_14      SALLEDPVGTVAPQIPPNWHIPSIQDAATPYHPPATPNNMGLIAGAVGGSLLAALVICGI 360
HSV1GPD_13      SALLEDPVGTVAPQIPPNWHIPSIQDAATPYHPPATPNNMGLIAGAVGGSLLAALVICGI 360
HSV1GPD_18      SALLEDPVGTVAPQIPPNWHIPSIQDAATPYHPPATPNNMGLIAGAVGGSLLAALVICGI 360
HSV1GPD_17      SALLEDPVGTVAPQIPPNWHIPSIQDAATPYHPPATPNNMGLIAGAVGGSLLAALVICGI 360
HSV1GPD_20      SALLEDPVGTVAPQIPPNWHIPSIQDAATPYHPPATPNNMGLIAGAVGGSLLAALVICGI 360
HSV1GPD_5       SALLEDPVGTVAPQIPPNWHIPSIQDAATPYHPPATPNNMGLIAGAVGGSLLAALVICGI 360
HSV1GPD_11      SALLEDPVGTVAPQIPPNWHIPSIQDAATPYHPPATPNNMGLIAGAVGGSLLAALVICGI 360
HSV1GPD_12      SALLEDPVGTVAPQIPPNWHIPSIQDAATPYHPPATPNNMGLIAGAVGGSLLAALVICGI 360
HSV1GPD_6       SALLEDPVGTVAPQIPPNWHIPSIQDAATPYHPPATPNNMGLIAGAVGGSLLAALVICGI 360
HSV1GPD_7       SALLEDPVGTVAPQIPPNWHIPSIQDAATPYHPPATPNNMGLIAGAVGGSLLAALVICGI 360
HSV1GPD_9       SALLEDPVGTVAPQIPPNWHIPSIQDAATPYHPPATPNNMGLIAGAVGGSLLVALVICGI 360
HSV1GPD_19      SALLEDPVGTVVPQIPPNWHIPSIQDAATPYHPPATPNNMGLIAGAVGGSLLVALVICGI 360
HSV1GPD_15      SALLEDPVGTVAPQIPPNWHIPSIQDAATPYHPPATPNNMGLIAGAVGGSLLAALVICGI 360
HSV1GPD_3       SALLEDPVGTVAPQIPPNWHIPSIQDAATPYHPPATPNNMGLIAGAVGGSLLAALVICGI 290
HSV2GPD_22      SALLEDPAGTVSSQIPPNWHIPSIQDVA-PHHAPAAPSNPGLIIGALAGSTLAVLVIGGI 359
HSV2GPD_24      SALLEDPAGTVSSQIPPNWHIPSIQDVA-PHHAPAAPSNPGLIIGALAGSTLAVLVIGGI 359
HSV2GPD_26      SALLEDPAGTVSSQIPPNWHIPSIQDVA-PHHAPAAPSNPGLIIGALAGSTLAVLVIGGI 359
HSV2GPD_21      SALLEDPAGTVSSQIPPNWHIPSIQDVA-PHHAPAAPSNPGLIIGALAGSTLAVLVIGGI 359
HSV2GPD_25      SALLEDPAGTVSSQIPPNWHIPSIQDVA-PHHAPAAPSNPGLIIGALAGSTLAVLVIGGI 359
HSV2GPD_23      SALLEDPAGTVSSQIPPNWHIPSIQDVA-PHHAPAAPSNPGLIIGALAGSTLAALVIGGI 359
HSV2GPD_27      SALLEDPAGTVSSQIPPNWHIPSIQDVA-PHHAPAAPSNPGLIIGALAGSTLAALVIGGI 359
HSV2GPD_28      SALLEDPAGTVSSQIPPNWHIPSIQDVA-PHHAPAAPANPGLIIGALAGSTLAALVIGGI 359
                *******.***  *************.* *:*.**:* * *** **:.** *..*** **

HSV1GPD_1       VYWMRRHTQKAPKRIRLPHIREDDQPSSHQPLFY 394
HSV1GPD_2       VYWMRRHTQKAPKRIRLPHIREDDQPSSHQPLFY 394
HSV1GPD_4       VYWMRRRTRKAPKRIRLPHIREDDQPSSHQPLFY 394
HSV1GPD_8       VYWMRRRTRKAPKRIRLPHIREDDQPSSHQPLFY 394
HSV1GPD_10      VYWMHRRTRKAPKRIRLPHIREDDQPSSHQPLFY 394
HSV1GPD_16      VYWMHRRTRKAPKRIRLPHIREDDQPSSHQPLFY 394
HSV1GPD_14      VYWMHRRTRKAPKRIRLPHIREDDQPSSHQPLFY 394
HSV1GPD_13      VYWMHRRTRKAPKRIRLPHIREDDQPSSHQPLFY 394
HSV1GPD_18      VYWMHRRTRKAPKRIRLPHIREDDQPSSHQPLFY 394
HSV1GPD_17      VYWMHRRTRKAPKRIRLPHIREDDQPSSHQPLFY 394
HSV1GPD_20      VYWMHRRTRKAPKRIRLPHIREDDQPSSHQPLFY 394
HSV1GPD_5       VYWMHRRTRKAPKRIRLPHIREDDQPSSHQPLFY 394
HSV1GPD_11      VYWMRRRTQKAPKRIRLPHIREDDQPSSHQPLFY 394
HSV1GPD_12      VYRMRRRTQKAPKRIRLPHIREDDQPSSHQPLFY 394
HSV1GPD_6       VYWMRRRTQKAPKRIRLPHIREDDQPSSHQPLFY 394
HSV1GPD_7       VYWMRRRTQKAPKRIRLPHIREDDQPSSHQPLFY 394
HSV1GPD_9       VYWMRRRTQKAPKRIRLPHIREDDQPSSHQPLFY 394
HSV1GPD_19      VYWMRRRTQKAPKRIRLPHIREDDQPSSHQPLFY 394
HSV1GPD_15      VYWMRRRTQKAPKRIRLPHIREDDQPSSHQPLFY 394
HSV1GPD_3       VYWMRRRTQKAPKRIRLPHIREDDQPSSHQPLFY 324
HSV2GPD_22      AFWVRRRAQMAPKRLRLPHIRDDDAPPSHQPLFY 393
HSV2GPD_24      AFWVRRRAQMAPKRLRLPHIRDDDAPPSHQPLFY 393
HSV2GPD_26      AFWVRRRAQMAPKRLRLPHIRDDDAPPSHQPLFY 393
HSV2GPD_21      AFWVRRRAQMAPKRLRLPHIRDDDAPPSHQPLFY 393
HSV2GPD_25      AFWVRRRAQMAPKRLRLPHIRDDDAPPSHQPLFY 393
HSV2GPD_23      AFWVRRRAQMAPKRLRLPHIRDDDAPPSHQPLFY 393
HSV2GPD_27      AFWVRRRAQMAPKRPRLPHIRDDDAPPSHQPLFY 393
HSV2GPD_28      AFWVRRRRSVAPKRLRLPHIRDDDAPPSHQPLFY 393
                .: ::*:   **** ******:** *.*******
```
