## Supplementary file 10 for "Contriving a chimeric polyvalent vaccine to prevent infections caused by Herpes Simplex Virus (Type-1 and Type-2): an exploratory immunoinformatic approach"

```
Alignment Glycoprotein E

HSV2GPE_29      MARGAGLVFFVGVWVVSCLAAAPRTSWKRVTSGEDVVLLPAPAGPEERTRAHKLLWAAEP 60
HSV2GPE_39      MARGAGLVFFVGVWVVSCLAAAPRTSWKRVTSGEDVVLLPAPAGPEERTRAHKLLWAAEP 60
HSV2GPE_26      MARGAGLVFFVGVWVVSCLAAAPRTSWKRVTSGEDVVLLPAPAGPEERTRAHKLLWAAEP 60
HSV2GPE_37      MARGAGLVFFVGVWVVSCLAAAPRTSWKRVTSGEDVVLLPAPAGPEERTRAHKLLWAAEP 60
HSV2GPE_38      MARGAGLVFFVGVWVVSCLAAAPRTSWKRVTSGEDVVLLPAPTGPEERTRAHKLLWAAEP 60
HSV2GPE_28      MARGAGLVFFVGVWVVSCLAAAPRTSWKRVTSGEDVVLLPAPAGPEERTRTHKLLWAAEP 60
HSV2GPE_34      ------------------------------------------------------------
HSV2GPE_25      MARGAGLVFFVGVWVVSCLAAAPRTSWKRVTSGEDVVLLPAPAGPEERTRAHKLLWAAEP 60
HSV2GPE_36      MARGAGLVFFVGVWVVSCLAAAPRTSWKRVTSGEDVVLLPAPAGPEERTRAHKLLWAAEP 60
HSV2GPE_40      ------------------------------------------------------------
HSV2GPE_35      MARGAGLVFFVGVWVVSCLAAAPRTSWKRVTSGEDVVLLPAPAGPEERTRAHKLLWAAEP 60
HSV2GPE_31      MARGAGLVFFVGVWVVSCLGAAPRTSWKRVTSGEDVVLLPAPAGPEERTRAHKLLWAAEP 60
HSV2GPE_32      ------------------------------------------------------------
HSV2GPE_22      MARGAGLVFFVGVWVVSCLAAAPRTSWKRVTSGEDVVLLPAPA---ERTRAHKLLWAAEP 57
HSV2GPE_23      MARGAGLVFFVGVWVVSCLAAAPRTSWKRVTSGEDVVLLPAPA---ERTRAHKLLWAAEP 57
HSV2GPE_24      MARGAGLVFFVGVWVVSCLAAAPRTSWKRVTSGEDVVLLPAPAGPEERTRAHKLLWAAEP 60
HSV2GPE_27      MARGAGLVFFVGVWVVSCLAAAPRTSWKRVTSGEDVVLLPAPAGPEERTRAHKLLWAAEP 60
HSV2GPE_33      MARGAGLVFFVGVWVVSCLAAAPRTSWKRVTSGEDVVLLPAPAGPEERTRAHKLLWAAEP 60
HSV2GPE_21      MARGAGLVFFVGVWVVSCLAAAPRTSWKRVTSGEDVVLLPAPA---ERTRAHKLLWAAEP 57
HSV2GPE_30      ------------------------------------------------------------
HSV1GPE_3       MDRGAVVGFLLGVCVVSCLAGTPKTSWRRVSVGEDVSLLPAPG-PTGRGPTQKLLWAVEP 59
HSV1GPE_4       MDRGAVVGFLLGVCVVSCLAGTPKTSWRRVSVGEDVSLLPAPG-PTGRGPTQKLLWAVEP 59
HSV1GPE_5       MDRGAVVGFLLGVCVVSCLAGTPKTSWRRVSVGEDVSLLPAPG-PTGRGPTQKLLWAVEP 59
HSV1GPE_10      MDRGAVVGFLLGVCVVSCLAGTPKTSWRRVSVGEDVSLLPAPG-PTGRGPTQKLLWAVEP 59
HSV1GPE_1       ------------------------------------------------------------
HSV1GPE_2       MDRGAVVGFLLGVCVVSCLAGTPKTSWRRVSVGEDVSLLPAPG-PTGRGPTQKLLWAVEP 59
HSV1GPE_7       MDRGAVVGFLLGVCVVSCLAGTPKTSWRRVSVGEDVSLLPAPG-PTGRGPTQKLLWAVEP 59
HSV1GPE_12      MDRGAVVGFLLGVCVVSCLAGTPKTSWRRVSVGEDVSLLPAPG-PTGRGPTQKLLWAVEP 59
HSV1GPE_19      MDRGAVVGFLLGVCVVSCLAGTPKTSWRRVSVGEDVSLLPAPG-PTGRGSTQKLLWAVEP 59
HSV1GPE_8       MDRGAVVGFLLGVCVVSCLAGTPKTSWRRVSVGEDVSLLPAPG-PTGRGPTQKLLWAVEP 59
HSV1GPE_14      MDRGAVVGFLLGVCVVSCLAGTPKTSWRRASVGEDVSLLPAPG-PTGRGPTQKLLWAVEP 59
HSV1GPE_13      MDRGAVVGFLLGVCVVSCLAGTPKTSWRRVSVGEDVSLLPAPG-PTGRGPTQKLLWAVEP 59
HSV1GPE_6       MDRGAVVGFLLGVCVVSCLAGTPKTSWRRVSVGEDVSLLPAPG-PTGRGPTQKLLWAVEP 59
HSV1GPE_9       MDRGAVVGFLLGVCVVSCLAGTPKTSWRRVSVGEDVSLLPAPG-PTGRGPTQKLLWAVEP 59
HSV1GPE_16      MDRGAVVGFLLGVCVVSCLAGTPKTSWRRVSVGEDVSLLPAPG-PTGRGPTQKLLWAVEP 59
HSV1GPE_20      MDRGAVVGFLLGVCVVSCLAGTPKTSWRRVSVGEDVSLLPAPG-PTGRGPTQKLLWAVEP 59
HSV1GPE_17      MDRGAVVGFLLGVCVVSCLAGTPKTSWRRVSVGEDVSLLPAPG-PTGRGPTQKLLWAVEP 59
HSV1GPE_18      MDRGAVVGFLLGVCVVSCLAGTPKTSWRRVSVGEDVSLLPAPG-PTGRGPTQKLLWAVEP 59
HSV1GPE_15      MDRGAVVGFLLGVCVVSCLAGTPKTSWRRVSVGEDVSLLPAPG-PTGRGPTQKLLWAVEP 59
HSV1GPE_11      MDRGAVVGFLLGVCVVSCLAGTPKTSWKRVSVGEDVSLLPAPG-PTGRGPTQKLLWAVEP 59


HSV2GPE_29      LDACGPLRPSWVALWPPRRVLETVVDAACMRAPEPLAIAYSPPFPAGDEGLYSELAWRDR 120
HSV2GPE_39      LDACGPLRPSWVALWPPRRVLETVVDAACMRAPEPLAIAYSPPFPAGDEGLYSELAWRDR 120
HSV2GPE_26      LDACGPLRPSWVALWPPRRVLETVVDAACMRAPEPLAIAYSPPFPAGDEGLYSELAWRDR 120
HSV2GPE_37      LDACGPLRPSWVALWPPRRVLETVVDAACMRAPEPLAIAYSPPFPAGDEGLYSELAWRDR 120
HSV2GPE_38      LDACGPLRPSWVALWPPRRVLETVVDAACMRAPEPLAIAYSPPFPAGDEGLYSELAWRDR 120
HSV2GPE_28      LDACGPLRPSWVALWPPRRVLETVVDAACMRAPEPLAIAYSPPFPAGDEGLYSELAWRDR 120
HSV2GPE_34      ----------WVALWPPRRVLETVVDAACMRAPEPLAIAYSPPFPAGDEGLYSELAWRDR 50
HSV2GPE_25      LDACGPLRPSWVALWPPRRVLETVVDAACMRAPEPLAIAYSPPFPAGDEGLYSELAWRDR 120
HSV2GPE_36      LDACGPLRPSWVALWPPRRVLETVVDAACMRAPEPLAIAYSPPFPAGDEGLYSELAWRDR 120
HSV2GPE_40      ----------WVALWPPRRVLETVVDAACMRAPEPLAIAYSPPFPAGDEGLYSELAWRDR 50
HSV2GPE_35      LDACGPLRPSWVALWPPRRVLETVVDAACMRAPEPLAIAYSPPFPAGDEGLYSELAWRDR 120
HSV2GPE_31      LDACGPLRPSWVALWPPRRVLETVVDAACMRAPEPLAIAYSPPFPAGDEGLYSELAWRDR 120
HSV2GPE_32      ----------WVALWPPRRVLETVVDAACMRAPEPLAIAYSPPFPAGDEGLYSELAWRDR 50
HSV2GPE_22      LDACGPLRPSWVALWPPRRVLETVVDAACMRAPEPLAIAYSPPFPAGDEGLYSELAWRDR 117
HSV2GPE_23      LDACGPLRPSWVALWPPRRVLETVVDAACMRAPEPLAIAYSPPFPAGDEGLYSELAWRDR 117
HSV2GPE_24      LDACGPLRPSWVALWPPRRVLETVVDAACMRAPEPLAIAYSPPFPAGDEGLYSELAWRDR 120
HSV2GPE_27      LDACGPLRPSWVALWPPRRVLETVVDAACMRAPEPLAIAYSPPFPAGDEGLYSELAWRDR 120
HSV2GPE_33      LDACGPLRPSWVALGPPRRVLETVVDAACMRAPEPLAIAYSPPFPAGDEGLYSELAWRDR 120
HSV2GPE_21      LDACGPLRPSWVALWPPRRVLETVVDAACMRAPEPLAIAYSPPFPAGDEGLYSELAWRDR 117
HSV2GPE_30      ----------WVALWPPRRVLETVVDAACMRAPEPLAIAYSPPFPAGDEGLYSELAWRDR 50
HSV1GPE_3       LDGCGPLHPSWVSLMPPKQVPETVVDAACMRAPVPLAMAYAPPAPSATGGLRTDFVWQER 119
HSV1GPE_4       LDGCGPLHPSWVSLMPPKQVPETVVDAACMRAPVPLAMAYAPPAPSATGGLRTDFVWQER 119
HSV1GPE_5       LDGCGPLHPSWVSLMPPKQVPETVVDAACMRAPVPLAMAYAPPAPSATGGLRTDFVWQER 119
HSV1GPE_10      LDGCGPLHPSWVSLMPPKQVPETVVDAACMRAPVPLAMAYAPPAPSATGGLPTDFVWQER 119
HSV1GPE_1       -----------VSLMPPKQVPETVVDAACMRAPVPLAMAYAPPAPSATGGLRTDFVWQER 49
HSV1GPE_2       LDGCGPLHPSWVSLMPPKQVPETVVDAACMRAPVPLAMAYAPPAPSATGGLRTDFVWQER 119
HSV1GPE_7       LDGCGPLHPSWVSLMPPKQVPETVVDAACMRAPVPLAMAYAPPAPSATGGLRTDFVWQER 119
HSV1GPE_12      LDGCGPLHPSWVSLMPPKQVPETVVDAACMRAPVPLAMAYAPPAPSATGGLPTDFVWQER 119
HSV1GPE_19      LDGCGPLHPSWVSLMPPKQVPETVVDAACMRAPVPLAMAYAPPAPSATGGLPTDFVWQER 119
HSV1GPE_8       LDGCGPLHPSWVSLMPPKQVPETVVDAACMRAPVPLAMAYAPPAPSATGGLRTDFVWQER 119
HSV1GPE_14      LDGCGPLHPSWVSLMPPKQVPETVVDAACMRAPVPLAMAYAPPAPSATGGLRTDFVWQER 119
HSV1GPE_13      LDGCGPLHPSWVSLMPPQQVPETVVDAACMRAPVPLAMAYAPPAPSATGGLRTDFVWQER 119
HSV1GPE_6       LDGCGPLHPSWVSLMPPKQVPETVVDAACMRAPVPLAMAYAPPAPSATGGLRTDFVWQER 119
HSV1GPE_9       LDGCGPLHPSWVSLMPPKQVPETVVDAACMRAPVPLAMAYAPPAPSATGGLRTDFVWQER 119
HSV1GPE_16      LDGCGPLHPSWVSLMPPKQVPETVVDAACMRAPVPLAMAYAPPAPSATGGLRTDFVWQER 119
HSV1GPE_20      LDGCGPLHPSWVSLMPPKQVPETVVDAACMRAPVPLAMAYAPPAPSATGGLRTDFVWQER 119
HSV1GPE_17      LDGCGPLHPSWVSLMPPKQVPETVVDAACMRAPVPLAMAYAPPAPSATGGLRTDFVWQER 119
HSV1GPE_18      LDGCGPLHPSWVSLMPPKQVPETVVDAACMRAPVPLAMAYAPPAPSATGGLRTDFVWQER 119
HSV1GPE_15      LDGCGPLHPSWVSLMPPKQVPETVVDAACMRAPVPLAMAYAPPAPSATGGLRTDFVWQER 119
HSV1GPE_11      LDGCGPLHPSWVSLMPPKQVPETVVDAACMRAPVPLAMAYAPPAPSATGGLRTDFVWQER 119
                           *:* **::* ************ ***:**:** *:.  ** :::.*::*

HSV2GPE_29      VAVVNESLVIYGALDTDSGLYTLSVVGLSDEARQVASVVLVVEPAPVPTP--TPDDYDEE 178
HSV2GPE_39      VAVVNESLVIYGALETDSGLYTLSVVGLSDEARQVASVVLVVEPAPVPTP--TPDDYDEE 178
HSV2GPE_26      VAVVNESLVIYGALETDSGLYTLSVVGLSDEARQVASVVLVVEPAPVPTP--TPDDYDEE 178
HSV2GPE_37      VAVVNESLVIYGARETDSGLYTLSVVGLSDEARQVASVVLVVEPAPVPTP--TPDDYDEE 178
HSV2GPE_38      VAVVNESLVIYGARETDSGLYTLSVVGLSDEARQVASVVLVVEPAPVPTP--TPDDYDEE 178
HSV2GPE_28      VAVVNESLVIYGALETDSGLYTLSVVGLSDEARQVASVVLVVEPAPVPTP--TPDDYDEE 178
HSV2GPE_34      VAVVNESLVIYGALETDSGLYTLSVVGLSDEARQVASVVLVVEPAPVPTP--TPDDYDEE 108
HSV2GPE_25      VAVVNESLVIYGALETDSGLYTLSVVGLSDEARQVASVVLVVEPAPVPTP--TPDDYDEE 178
HSV2GPE_36      VAVVNESLVIYGALETDSGLYTLSVVGLSDEARQVASVVLVVEPAPVPTP--TPDDYDEE 178
HSV2GPE_40      VAVVNESLVIYGALETDSGLYTLSVVGLSDEARQVASVVLVVEPAPVPTP--TPDDYDEE 108
HSV2GPE_35      VAVVNESLVIYGALETDSGLYTLSVVGLSDEARQVASVVLVVEPAPVPTP--TPDDYDEE 178
HSV2GPE_31      VAVVNESLVIYGALETDSGLYTLSVVGLSDEARQVASVVLVVEPAPVPTP--TPDDYDEE 178
HSV2GPE_32      VAVVNESLVIYGALETDSGLYTLSVVGLSDEARQVASVVLVVEPAPVPTP--TPDDYDEE 108
HSV2GPE_22      VAVVNESLVIYGALETDSGLYTLSVVGLSDEARQVASVVLVVEPAPVPTP--TPDDYDEE 175
HSV2GPE_23      VAVVNESLVIYGALETDSGLYTLSVVGLSDEARQVASVVLVVEPAPVPTP--TPDDYDEE 175
HSV2GPE_24      VAVVNESLVIYGALETDSGLYTLSVVGLSDEARQVASVVLVVEPAPVPTP--TPDDYDEE 178
HSV2GPE_27      VAVVNESLVIYGALETDSGLYTLSVVGLSDEARQVASVVLVVEPAPVPTP--TPDDYDEE 178
HSV2GPE_33      VAVVNESLVIYGALETDSGLYTLSVVGLSDEARQVASVVLVVEPAPVPTP--TPDDYDEE 178
HSV2GPE_21      VAVVNESLVIYGALETDSGLYTLSVVGLSDEARQVASVVLVVEPAPVPTP--TPDDYDEE 175
HSV2GPE_30      VAVVNESLVIYGARETDSGLYTLSVVGLSDEARQVASVVLVVEPAPVPTP--TPDDYDEE 108
HSV1GPE_3       AAVVNRSLVIHGVRETDSGLYTLSVGDIKDPARQVASVVLVVQPAPVPTPPPTPADYDED 179
HSV1GPE_4       AAVVNRSLVIHGVRETDSGLYTLSMGDIKDPARQVASVVLVVQPAPVPTPPPTPADYDED 179
HSV1GPE_5       AAVVNRSLVIHGVRETDSGLYTLSVGDIKDPARQVASVVLVVQPAPVPTPPPTPADYDED 179
HSV1GPE_10      AAVVNRSLVIHGVRETDSGLYTLSVGDIKDPARQVASVVLVVQPAPVPTPPPTPADYDED 179
HSV1GPE_1       AAVVNRSLVIHGVRETDSGLYTLSVGDIKDPARQVASVVLVVQPAPVPTPPPTPADYDED 109
HSV1GPE_2       AAVVNRSLVIHGVRETDSGLYTLSVGDIKDPARQVASVVLVVQPAPVPTPPPTPADYDED 179
HSV1GPE_7       AAVVNRSLVIHGVRETDSGLYTLSVGDIKDPARQVASVVLVVQPVPVPTPPPTPADYDED 179
HSV1GPE_12      AAVVNRSLVIHGVRETDSGLYTLSVGDIKDPARQVASVVLVVQPAPVPTPPPTPADYDED 179
HSV1GPE_19      AAVVNRSLVIHGVRETDSGLYTLSVGDIKDPARQVASVVLVVQPAPVPTPPPTPADYDED 179
HSV1GPE_8       AAVVNRSLVIHGVRETDSGLYTLSVGDIKDPARQVASVVLVVQPAPVPTPPPTPADYDEE 179
HSV1GPE_14      AAVVNRSLVIHGVRETDSGLYTLSVGDIKDPARQVASVVLVVQPAPVPTPPPTPADYDEE 179
HSV1GPE_13      AAVVNRSLVIHGVRETDSGLYTLSVGDIKDPARQVASVVLVVQPAPVPTPPPTPADYDEE 179
HSV1GPE_6       AAVVNRSLVIYGVRETDSGLYTLSVGDIKDPARQVASVVLVVQPAPVPTPPPTPADYDED 179
HSV1GPE_9       AAVVNRSLVIYGVRETDSGLYTLSVGDIKDPARQVASVVLVVQPAPVPTPPPTPADYDED 179
HSV1GPE_16      TAVVNRSLVIYGVRETDSGLYTLSVGDIKDPARQVASVVLVVQPAPVPTPPPTPADYDED 179
HSV1GPE_20      TAVVNRSLVIYGVRETDSGLYTLSVGDIKDPARQVASVVLVVQPAPVPTPPPTPADYDED 179
HSV1GPE_17      AAVVNRSLVIYGVRETDSGLYTLSVGDIKDPARQVASVVLVVQPAPVPTPPPTPADYDED 179
HSV1GPE_18      AAVVNRSLVIYGVRETDSGLYTLSVGDIKDPARQVASVVLVVQPAPVPTPPPTPADYDED 179
HSV1GPE_15      AAVVNRSLVIYGVRETDSGLYTLSVGDIKDPARQVASVVLVVQPAPVPTPPPTPADYDED 179
HSV1GPE_11      AAVVNRSLVIYGVRETDSGLYTLSVGDIKDPARQVASVVLVVQPAPVPTPPPTPADYDED 179
                .****.****:*. :*********: .:.* ***********:*.*****  ** ****:

HSV2GPE_29      DDAGVSERTPVSVPPPTPPRRPP-VAPPTHPRVIPEVSHVRGVTVHMETPEAILFAPGET 237
HSV2GPE_39      DDAGVSERTPVSVPPPTPPRRPP-VAPPTHPRVIPEVSHVRGVTVHMETPEAILFAPGET 237
HSV2GPE_26      DDAGVSERTPVSVPPPTPPRRPP-VAPPTHPRVIPEVSHVRGVTVHMETPEAILFAPGET 237
HSV2GPE_37      DDAGVSERTPVSVPPPTPPRRPP-VAPQTHPRVIPEVSHVRGVTVHMETPEAILFAPGET 237
HSV2GPE_38      DDAGVSERTPVSVPPPTPPRRPP-VAPQTHPRVIPEVSHVRGVTVHMETPEAILFAPGET 237
HSV2GPE_28      DDAGVSERTPVSVPPATPPRRPP-VAPPTHPRVIPEVSHVRGVTVHMETPEAILFAPGET 237
HSV2GPE_34      DDAGVSERTPVSVPPATPPRRPP-VAPPTHPRVIPEVSHVRGVTVHMETPEAILFAPGET 167
HSV2GPE_25      DDAGVSERTPVSVPPATPPRRPP-VAPPTHPRVIPEVSHVRGVTVHMETPEAILFAPGET 237
HSV2GPE_36      DDAGVSERTPVSVPPPTPPRRPP-VAPPTHPRVIPEVSHVRGVTVHMETPEAILFAPGET 237
HSV2GPE_40      DDAGVSERTPVSVPPPTPPRRPP-VAPPTHPRVIPEVSHVRGVTVHMETPEAILFAPGET 167
HSV2GPE_35      DDAGVSERTLVSVPPPTPPRRPP-VAPPTHPRVIPEVSHVRGVTVHMETPEAILFAPGET 237
HSV2GPE_31      DDAGVSERTPVSVPPPTPPRRPP-VAPPTHPRVIPEVSHVRGVTVHMETPEAILFAPGET 237
HSV2GPE_32      DDAGVSERTPVSVPPPTPPRRPP-VAPPTHPRVIPEVSHVRGVTVHMETPEAILFAPGET 167
HSV2GPE_22      DDAGVSERTPVSVPPPTPPRRPP-VAPPTHPRVIPEVSHVRGVTVHMETPEAILFAPGET 234
HSV2GPE_23      DDAGVSERTPVSVPPPTPPRRPP-VAPPTHPRVIPEVSHVRGVTVHMETPEAILFAPGET 234
HSV2GPE_24      DDAGVSERTPVSVPPPPPPRRPP-VAPPTHPRVIPEVSHVRGVTVHMETPEAILFAPGET 237
HSV2GPE_27      DDAGVSERTPVSVPPPPPPRRPP-VAPPTHPRVIPEVSHVRGVTVHMETPEAILFAPGET 237
HSV2GPE_33      DDAGVSERTPVSVPPTPPPRRPP-VAPPTHPRVIPEVSHVRGVTVHMETPEAILFAPGET 237
HSV2GPE_21      DDAGVTNARRSAFPPQPPPRRPP-VAPPTHPRVIPEVSHVRGVTVHMETLEAILFAPGET 234
HSV2GPE_30      DDAGVSERTPVSVPPTTPPRRPP-VAPPTHPRVIPEVSHVRGVTVHMETPEAILFAPGET 167
HSV1GPE_3       DNDEGEDESLAGTPASGTPRLPPPPAPPRSWPSAPEVSHVRGVTVRMETPEAILFSPGET 239
HSV1GPE_4       DNDEGEDESLAGTPASGTPRLPPPPAPPRSWPSAPEVSHVRGVTVRMETPEAILFSPGET 239
HSV1GPE_5       DNDEGEDESLAGTPASGTPRLPPPPAPPRSWPSAPEVSHVRGVTVRMETPEAILFSPGET 239
HSV1GPE_10      DNDEGEDESLAGTPASGTPRLPPPPAPPRSWPSAPEVSHVRGVTVRMETPEAILFSPGET 239
HSV1GPE_1       DNDEGEDESLAGTPASGTPRLPPPPAPPRSWPSAPEVSHVRGVTVRMETPEAILFSPGET 169
HSV1GPE_2       DNDEGEDESLAGTPASGTPRLPPPPAPPRSWPSAPEVSHVRGVTVRMETPEAILFSPGET 239
HSV1GPE_7       DNDEGEDESLAGTPASGTPRLPPPPAPPRSWPSAPEVSHVRGVTVRMETPEAILFSPGET 239
HSV1GPE_12      DNDEGEDESLAGTPASGTPRLPPPPAPPRSWPSAPEVSHVRGVTVRMETPEAILFSPGET 239
HSV1GPE_19      DNDEGEDESLAGTPASGTPRLPPPPAPPRSWPSAPEVSHVRGVTVRMETPEAILFSPGET 239
HSV1GPE_8       DNDEGEDESLAGTPASGTPRLPPPPAPPRSWPSAPEVSHVRGVTVRMETPEAILFSPGET 239
HSV1GPE_14      DNDEGEDESLAGTPASGTPRLPPPPAPPRSWPSAPEVSHVRGVTVRMETPEAILFSPGET 239
HSV1GPE_13      DNDEGEDESLAGTPASGTPRLPPPPAPPRSWPSAPEVSHVRGVTVRMETPEAILFSPGET 239
HSV1GPE_6       DNDEGEDESLAGTPASGTPRLPPPPAPPRSWPSAPEVSHVRGVTVRMETPEAILFSPGET 239
HSV1GPE_9       DNDEGEDESLAGTPASGTPRLPPPPAPPRSWPSAPEVSHVRGVTVRMETPEAILFSPGET 239
HSV1GPE_16      DNDEGEDESLAGTPASGTPRLPPPPAPPRSWPSAPEVSHVRGVTVRMETPEAILFSPGEA 239
HSV1GPE_20      DNDEGEDESLAGTPASGTPRLPPPPAPPRSWPSAPEVSHVRGVTVRMETPEAILFSPGEA 239
HSV1GPE_17      DNDEGEDESLAGTPASGTPRLPPPPAPPRSWPSAPEVSHVRGVTVRMETPEAILFSPGEA 239
HSV1GPE_18      DNDEGEDESLAGTPASGTPRLPPPPAPPRSWPSAPEVSHVRGVTVRMETPEAILFSPGEA 239
HSV1GPE_15      DNDEGEDESLAGTPASGTPRLPPPPAPPRSWPSAPEVSHVRGVTVRMETPEAILFSPGET 239
HSV1GPE_11      DNDEGEDESLAGTPASGTPRLPPPPAPPRSWPSAPEVSHVRGVTVRMETPEAILFSPGET 239
                *:    :    . *.  .** **  **       ***********:*** *****:***:

HSV2GPE_29      FGTNVSIHAIAHDDGPYAMDVVWMRFDVPSSCAEMRIYEACLYHPQLPECLSPADAPCAV 297
HSV2GPE_39      FGTNVSIHAIAHDDGPYAMDVVWMRFDVPSSCAEMRIYEACLYHPQLPECLSPADAPCAV 297
HSV2GPE_26      FGTNVSIHAIAHDDGPYAMDVVWMRFDVPSSCAEMRIYEACLYHPQLPECLSPADAPCAV 297
HSV2GPE_37      FGTNVSIHAIAHDDGPYAMDVVWMRFDVPSSCAEMRIYEACLYHPQLPECLSPADAPCAV 297
HSV2GPE_38      FGTNVSIHAIAHDDGPYAMDVVWMRFDVPSSCAEMRIYEACLYHPQLPECLSPADAPCAV 297
HSV2GPE_28      FGTNVSIHAIAHDDGPYAMDVVWMRFDVPSSCAEMRIYEACLYHPQLPECLSPADAPCAV 297
HSV2GPE_34      FGTNVSIHAIAHDDGPYAMDVVWMRFDVPSSCAEMRIYEACLYHPQLPECLSPADAPCAV 227
HSV2GPE_25      FGTNVSIHAIAHDDGPYAMDVVWMRFDVPSSCAEMRIYEACLYHPQLPECLSPADAPCAV 297
HSV2GPE_36      FGTNVSIHAIAHDDGPYAMDVVWMRFDVPSSCAEMRIYEACLYHPQLPECLSPADAPCAV 297
HSV2GPE_40      FGTNVSIHAIAHDDGPYAMDVVWMRFDVPSSCAEMRIYEACLYHPQLPECLSPADAPCAV 227
HSV2GPE_35      FGTNVSIHAIAHDDGPYAMDVVWMRFDVPSSCAEMRIYEACLYHPQLPECLSPADAPCAV 297
HSV2GPE_31      FGTNVSIHAIAHDDGPYAMDVVWMRFDVPSSCAEMRIYEACLYHPQLPECLSPADAPCAV 297
HSV2GPE_32      FGTNVSIHAIAHDDGPYAMDVVWMRFDVPSSCAEMRIYEACLYHPQLPECLSPADAPCAV 227
HSV2GPE_22      FGTNVSIHAIAHDDGPYAMDVVWMRFDVPSSCAEMRIYEACLYHPQLPECLSPADAPCAV 294
HSV2GPE_23      FGTNVSIHAIAHDDGPYAMDVVWMRFDVPSSCAEMRIYEACLYHPQLPECLSPADAPCAV 294
HSV2GPE_24      FGTNVSIHAIAHDDGPYAMDVVWMRFDVPSSCAEMRIYEACLYHPQLPECLSPADAPCAV 297
HSV2GPE_27      FGTNVSIHAIAHDDGPYAMDVVWMRFDVPSSCAEMRIYEACLYHPQLPECLSPADAPCAV 297
HSV2GPE_33      FGTNVSIHAIAHDDGPYAMDVVWMRFDVPSSCAEMRIYEACLYHPQLPECLSPADAPCAV 297
HSV2GPE_21      FGTNVSIHAIAHDDGPYAMDVVWMRFDVPSSCADMRIYEACLYHPQLPECLSPADAPCAV 294
HSV2GPE_30      FGTNVSIHAIAHDDGPYAMDVVWMRFDVPSSCAEMRIYEACLYHPQLPECLSPADAPCAV 227
HSV1GPE_3       FSTNVSIHSIAHDDQTYSMDVVWLRFDVPTSCAEMRIYESCLYHPQLPECLSPADAPCAA 299
HSV1GPE_4       FSTNVSIHAIAHDDQTYSMDVVWLRFDVPTSCAEMRIYESCLYHPQLPECLSPADAPCAA 299
HSV1GPE_5       FSTNVSIHAIAHDDQTYSMDVVWLRFDVPTSCAEMRIYESCLYHPQLPECLSPADAPCAA 299
HSV1GPE_10      FSTNVSIHAIAHDDQTYSMDVVWLRFDVPTSCAEMRIYESCLYHPQLPECLSPADAPCAA 299
HSV1GPE_1       FSTNVSIHAIAHDDQTYSMDVVWLRFDVPTSCAEMRIYESCLYHPQLPECLSPADAPCAA 229
HSV1GPE_2       FSTNVSIHAIAHDDQTYAMDVVWLRFDVPTSCAEMRIYESCLYHPQLPECLSPADAPCAA 299
HSV1GPE_7       FSTNVSIHAIAHDDQTYAMDVVWLRFDVPTSCAEMRIYESCLYHPQLPECLSPADAPCAA 299
HSV1GPE_12      FSTNVSIHAIAHDDQTYAMDVVWLRFDVPTSCAEMRIYESCLYHPQLPECLSPADAPCAA 299
HSV1GPE_19      FSTNVSIHAIAHDDQTYAMDVVWLRFDVPTSCAEMRIYESCLYHPQLPECLSPADAPCAA 299
HSV1GPE_8       FSTNVSIHAIAHDDQTYAMDVVWLRFDVPTSCAEMRIYESCLYHPQLPECLSPADAPCAA 299
HSV1GPE_14      FSTNVSIHAIAHDDQTYAMDVVWLRFDVPTSCAEMRIYESCLYHPQLPECLSPADAPCAA 299
HSV1GPE_13      FSTNVSIHAIAHDDQTYAMDVVWLRFDVPTSCAEMRIYESCLYHPQLPECLSPADAPCAA 299
HSV1GPE_6       FSTNVSIHAIAHDDQTYAMDVVWLRFDVPTSCAEMRIYESCLYHPQLPECLSPADAPCAA 299
HSV1GPE_9       FSTNVSIHAIAHDDQTYAMDVVWLRFDVPTSCAEMRIYESCLYHPQLPECLSPADAPCAA 299
HSV1GPE_16      FSTNVSIHAIAHDDQTYAMDVVWLRFDVPTSCAEMRIYESCLYHPQLPECLSPADAPCAA 299
HSV1GPE_20      FSTNVSIHAIAHDDQTYAMDVVWLRFDVPTSCAEMRIYESCLYHPQLPECLSPADAPCAA 299
HSV1GPE_17      FSTNVSIHAIAHDDQTYTMDVVWLRFDVPTSCAEMRIYESCLYHPQLPECLSPADAPCAA 299
HSV1GPE_18      FSTNVSIHAIAHDDQTYTMDVVWLRFDVPTSCAEMRIYESCLYHPQLPECLSPADAPCAA 299
HSV1GPE_15      FNTNVSIHAIAHDDQTYAMDVVWLRFDVPTSCAEMRIYESCLYHPQLPECLSPADAPCAA 299
HSV1GPE_11      FSTNVSIHAIAHDDQTYAMDVVWLRFDVPTSCAEMRIYESCLYHPQLPECLSPADAPCAA 299
                *.******:***** .*:*****:*****:***:*****:*******************.

HSV2GPE_29      SSWAYRLAVRSYAGCSRTTPPPRCFAEARMEPVPGLAWLASTVNLEFQHASPQHAGLYLC 357
HSV2GPE_39      SSWAYRLAVRSYAGCSRTTPPPRCFAEARMEPVPGLAWLASTVNLEFQHASPQHAGLYLC 357
HSV2GPE_26      SSWAYRLAVRSYAGCSRTTPPPRCFAEARMEPVPGLAWLASTVNLEFQHASPQHAGLYLC 357
HSV2GPE_37      SSWAYRLAVRSYAGCSRTTPPPRCFAEARMEPVPGLAWLASTVNLEFQHASPQHAGLYLC 357
HSV2GPE_38      SSWAYRLAVRSYAGCSRTTPPPRCFAEARMEPVPGLAWLASTVNLEFQHASPQHAGLYLC 357
HSV2GPE_28      SSWAYRLAVRSYAGCSRTTPPPRCFAEARMEPVPGLAWLASTVNLEFQHASPQHAGLYLC 357
HSV2GPE_34      SSWAYRLAVRSYAGCSRTTPPPRCFAEARMEPVPGLAWLASTVNLEFQHASPQHAGLYLC 287
HSV2GPE_25      SSWAYRLAVRSYAGCSRTTPPPRCFAEARMEPVPGLAWLASTVNLEFQHASPQHAGLYLC 357
HSV2GPE_36      SSWAYRLAVRSYAGCSRTTPPPRCFAEARMEPVPGLAWLASTVNLEFQHASPQHAGLYLC 357
HSV2GPE_40      SSWAYRLAVRSYAGCSRTTPPPRCFAEARMEPVPGLAWLASTVNLEFQHASPQHAGLYLC 287
HSV2GPE_35      SSWAYRLAVRSYAGCSRTTPPPRCFAEARMEPVPGLAWLASTVNLEFQHASPQHAGLYLC 357
HSV2GPE_31      SSWAYRLAVRSYAGCSRTTPPPRCFAEARMEPVPGLAWLASTVNLEFQHASPQHAGLYLC 357
HSV2GPE_32      SSWAYRLAVRSYAGCSRTTPPPRCFAEARMEPVPGLAWLASTVNLEFQHASPQHAGLYLC 287
HSV2GPE_22      SSWAYRLAVRSYAGCSRTTPPPRCFAEARMEPVPGLAWLASTVNLEFQHASPQHAGLYLC 354
HSV2GPE_23      SSWAYRLAVRSYAGCFRTTPPPRCFAEARMEPVPGLAWLASTVNLEFQHASPQHAGLYLC 354
HSV2GPE_24      SSWAYRLAVRSYAGCSRTTPPPRCFAEARMEPVPGLAWLASTVNLEFQHASPQHAGLYLC 357
HSV2GPE_27      SSWAYRLAVRSYAGCSRTTPPPRCFAEARMEPVPGLAWLASTVNLEFQHASPQHAGLYLC 357
HSV2GPE_33      SSWAYRLAVRSYAGCSRTTPPPRCFAEARMEPVQGLAWLASTVNLEFQHASPQHAGLYLC 357
HSV2GPE_21      SSWAYRLAVRSYAGCSRTTPPPRCFAEARMEPVPGLAWLASTVNLEFQHASPQHAGLYLC 354
HSV2GPE_30      SSWAYRLAVRSYAGCSRTTPPPRCFAEARMEPVPGLAWLASTVNLEFQHASPQHAGLYLC 287
HSV1GPE_3       STWTSRLAVRSYAGCSRTNPPPRCSAEAHMEPVPGLAWQAASVNLEFRDASPQHSGLYLC 359
HSV1GPE_4       STWTSRLAVRSYAGCSRTNPPPRCSAEAHMEPVPGLAWQAASVNLEFRDASPQHSGLYLC 359
HSV1GPE_5       STWTSRLAVRSYAGCSRTNPQPRCSAEAHMEPVPGLAWQAASVNLEFRDASPQHSGLYLC 359
HSV1GPE_10      STWTSRLAVRSYAGCSRTNPPPRCSAEAHMEPVPGLAWQAASVNLEFRDASPQHSGLYLC 359
HSV1GPE_1       STWTSRLAVRSYAGCSRTNPPPRCSAEAHMEPVPGLAWQAASVNLEFRDASPQHSGLYLC 289
HSV1GPE_2       STWTSRLAVRSYAGCSRTNPPPRCSAEAHMEPVPGLAWQAASVNLEFRDASPQHSGLYLC 359
HSV1GPE_7       STWTSRLAVRSYAGCSRTNPPPRCSAEAHMEPVPGLAWQAASVNLEFRDASPQHSGLYLC 359
HSV1GPE_12      STWTSRLAVRSYAGCSRTNPPPRCSAEAHMEPVPGLAWQAASVNLEFRDASPQHSGLYLC 359
HSV1GPE_19      STWTSRLAVRSYAGCSRTNPPPRCSAEAHMEPVPGLAWQAASVNLEFRDASPQHSGLYLC 359
HSV1GPE_8       STWTSRLAVRSYAGCSRTNPPPRCSAEAHMEPVPGLAWQAASVNLEFRDASPQHSGLYLC 359
HSV1GPE_14      STWTSRLAVRSYAGCSRTNPPPRCSAEAHMEPVPGLAWQAASVNLEFRDASPQHSGLYLC 359
HSV1GPE_13      STWTSRLAVRSYAGCSRTNPPPRCSAEAHMEPVPGLAWQAASVNLEFRDASPQHSGLYLC 359
HSV1GPE_6       STWTSRLAVRSYAGCSRTNPPPRCSAEAHMEPVPGLAWQAASVNLEFRDASPQHSGLYLC 359
HSV1GPE_9       STWTSRLAVRSYAGCSKTNPPPRCSAEAHMEPVPGLAWQAASVNLEFRDASPQHSGLYLC 359
HSV1GPE_16      STWTSRLAVRSYAGCSRTNPPPRCSAEAHMEPVPGLAWQAASVNLEFRDASPQHSGLYLC 359
HSV1GPE_20      STWTSRLAVRSYAGCSRTNPPPRCSAEAHMEPVPGLAWQAASVNLEFRDASPQHSGLYLC 359
HSV1GPE_17      STWTSRLAVRSYAGCSRTNPPPRCSAEAHMEPFPGLAWQAASVNLEFRDASPQHSGLYLC 359
HSV1GPE_18      STWTSRLAVRSYAGCSRTNPPPRCSAEAHMEPFPGLAWQAASVNLEFRDASPQHSGLYLC 359
HSV1GPE_15      STWTSRLAVRSYAGCSRTNPPPRCSAEAHMEPVPGLAWQAASVNLEFRDASPQHSGLYLC 359
HSV1GPE_11      STWTSRLAVRSYAGCSRTNPPPRCSAEAHMEPVPGLAWQAASVNLEFRDASPQHSGLYLC 359
                *:*: ********** :*.* *** ***:***. **** *::*****:.*****:*****

HSV2GPE_29      VVYVDDHIHAWGHMTISTAAQYRNAVVEQHLPQRQPEPVEPTRPHVRAPHPAPSARGPLR 417
HSV2GPE_39      VVYVDDHIHAWGHMTISTAAQYRNAVVEQHLPQRQPEPVEPTRPHVRAPHPAPSARGPLR 417
HSV2GPE_26      VVYVDDHIHAWGHMTISTAAQYRNAVVEQHLPQRQPEPVEPTRPHVRAPHPAPSARGPLR 417
HSV2GPE_37      VVYVDDHIHAWGHMTISTAAQYRNAVVEQHLPQRQPEPVEPTRPHVRAPHPAPSARGPLR 417
HSV2GPE_38      VVYVDDHIHAWGHMTISTAAQYRNAVVEQHLPQRQPEPVEPTRPHVRAPHPAPSARGPLR 417
HSV2GPE_28      VVYVDDHIHAWGHMTISTAAQYRNAVVEQHLPQRQPEPVEPTRPHVRAPHPAPSARGPLR 417
HSV2GPE_34      VVYVDDHIHAWGHMTISTAAQYRNAVVEQHLPQRQPEPVEPTRPHVRAPPPAPSARGPLR 347
HSV2GPE_25      VVYVDDHIHAWGHMTISTAAQYRNAVVEQHLPQRQPEPVEPTRPHVRAPHPAPSARGPLR 417
HSV2GPE_36      VVYVDDHIHAWGHMTISTAAQYRNAVVEQHLPQRQPEPVEPTRPHVRAPPPAPSARGPLR 417
HSV2GPE_40      VVYVDDHIHAWGHMTISTAAQYRNAVVEQHLPQRQPEPVEPTRPHVRAPPPAPSARGPLR 347
HSV2GPE_35      VVYVDDHIHAWGHMTISTAAQYRNAVVEQHLPQRQPEPVEPTRPHVRAPPPAPSARGPLR 417
HSV2GPE_31      VVYVDDHIHAWGHMTISTAAQYRNAVVEQHLPQRQPEPVEPTRPHVRAPHPAPSARGPLR 417
HSV2GPE_32      VVYVDDHIHAWGHMTISTAAQYRNAVVEQHLPQRQPEPVEPTRPHVRAPHPAPSARGPLR 347
HSV2GPE_22      VVYVDDHIHAWGHMTISTAAQYRNAVVEQHLPQRQPEPVEPTRPHVRAPHPAPSARGPLR 414
HSV2GPE_23      VVYVDDHIHAWGHMTISTAAQYRNAVVEQHLPQRQPEPVEPTRPHVRAPHPAPSARGPLR 414
HSV2GPE_24      VVYVDDHIHAWGHMTISTAAQYRNAVVEQHLPQRQPEPVEPTRPHVRAPHPAPSARGPLR 417
HSV2GPE_27      VVYVDDHIHAWGHMTISTAAQYRNAVVEQHLPQRQPEPVEPTRPHVRAPHPVPSARGPLR 417
HSV2GPE_33      VVYVDDHIHAWGHMTISTAAQYRNAVVEQHLPQRQPEPVEPTRPHVRAPHPAPSARGPLR 417
HSV2GPE_21      VVYVDDHIHAWGHMTISTAAQYRNAVVEQHLPQRQPEPVEPTRPHVRAPHPAPSARGPLR 414
HSV2GPE_30      VVYVDDHIHAWGHMTISTAAQYRNAVVEQHLPQRQPEPVEPTRPHVRAPHPAPSARGPLR 347
HSV1GPE_3       VVYVNDHIHAWGHITISTAAQYRNAVVEQPLPQRGADLAEPTHPHVGAPPHAPPTHGALR 419
HSV1GPE_4       VVYVNDHIHAWGHITISTAAQYRNAVVEQPLPQRGADLAEPTHPHVGAPPHAPPTHGALR 419
HSV1GPE_5       VVYVNDHIHAWGHITISTAAQYRNAVVEQPLPQRGADLAEPTHPHVGAPPHAPPTHGALR 419
HSV1GPE_10      VVYVNDHIHAWGHITISTAAQYRNAVVEQPLPQRGADLAEPTHPHVGAPPHAPPTHGALR 419
HSV1GPE_1       VVYVNDHIHAWGHITISTAAQYRNAVVEQPLPQRGADLAEPTHPHVGAPPHAPPTHGALR 349
HSV1GPE_2       VVYVNDHIHAWGHITISTAAQYRNAVVEQPLPQRGADLAEPTHPHVGAPPHAPPTHGALR 419
HSV1GPE_7       VVYVNDHIHAWGHITISTAAQYRNAVVEQPLPQRGADLAEPTHPHVGAPPHAPPTHGALR 419
HSV1GPE_12      VVYVNDHIHAWGHITISTAAQYRNAVVEQPLPQRGADLAEPTHPHIGAPPHAPPTHGALR 419
HSV1GPE_19      VVYVNDHIHAWGHITISTAAQYRNAVVEQPLPQRGADLAEPTHPHVGAPPHAPPTHGALR 419
HSV1GPE_8       VVYVNDHIHAWGHITISTAAQYRNAVVEQPLPQRGADLAEPTHPHVGAPPHAPPTHGALR 419
HSV1GPE_14      VVYVNDHIHAWGHITISTAAQYRNAVVEQPLPQRGADLAEPTHPHVGAPPHAPPTHGALR 419
HSV1GPE_13      VVYVNDHIHAWGHITISTAAQYRNAVVEQPLPQRGADLAEPTHPHVGAPPHAPPTHGALR 419
HSV1GPE_6       VVYVNDHIHAWGHITISTAAQYRNAVVEQPLPQRGADLAEPTHPHVGAPPHAPPTHGALR 419
HSV1GPE_9       VVYVNDHIHAWGHITISTAAQYRNAVVEQPLPQRGADLAEPTHPHVGAPPHAPPTHGALR 419
HSV1GPE_16      VVYVNDHIHAWGHITISTAAQYRNAVVEQPLPQRGADLAEPTHPHVGAPPHAPPTHGALR 419
HSV1GPE_20      VVYVNDHIHAWGHITISTAAQYRNAVVEQPLPQRGADLAEPTHPHVGAPPHAPPTHGALR 419
HSV1GPE_17      VVYVNDHIHAWGHITISTAAQYRNAVVEQPLPQRGADLAEPTHPHVGAPPHAPPTHGALR 419
HSV1GPE_18      VVYVNDHIHAWGHITISTAAQYRNAVVEQPLPQRGADLAEPTHPHVGAPPHAPPTHGALR 419
HSV1GPE_15      VVYVNDHIHAWGHITISTAAQYRNAVVEQPLPQRGADLAEPTHPHVGAPPHATPTHGALR 419
HSV1GPE_11      VVYVNDHIHAWGHITISTAAQYRNAVVEQPLPQRGADLAEPTHPHVGAPPHAPPTHGALR 419
                ****:********:*************** **** .: .***:**: **  ...::*.**

HSV2GPE_29      LGAVLGAALLLAALGLSAWACMTCWRRRSWRAVKSRASATGPTYIRVADSELYADWSSDS 477
HSV2GPE_39      LGAVLGAALLLAALGLSAWACMTCWRRRSWRAVKSRASATGPTYIRVADSELYADWSSDS 477
HSV2GPE_26      LGAVLGAALLLAALGLSAWACMTCWRRRSWRAVKSRASATGPTYIRVADSELYADWSSDS 477
HSV2GPE_37      LGAVLGAALLLAALGLSAWACMTCWRRRSWRAVKSRASATGPTYIRVADSELYADWSSDS 477
HSV2GPE_38      LGAVLGAALLLAALGLSAWACMTCWRRRSWRAVKSRASATGPTYIRVADSELYADWSSDS 477
HSV2GPE_28      LGAVLGAALLLAALGLSAWACMTCWRRRSWRAVKSRASATGPTYIRVADSELYADWSSDS 477
HSV2GPE_34      LGAVLGAALLLAALGLSAWACMTCWRRRSWRAVKSRASATGPTYIRVADSELYADWSSDS 407
HSV2GPE_25      LGAVLGAALLLAALGLSAWACMTCWRRRSWRAVKSRASATGPTYIRVADSELYADWSSDS 477
HSV2GPE_36      LGAVLGAALLLAALGLSAWACMTCWRRRSWRAVKSRASATGPTYIRVADSELYADWSSDS 477
HSV2GPE_40      LRAVLGAALLLAALGLSAWACMTCWRRRSWRAVKSRASATGPTYIRVADSELYADWSSDS 407
HSV2GPE_35      LGAVLGAALLLAALGLSAWACMTCWRRRSWRAVKSRASATGPTYIRVADSELYADWSSDS 477
HSV2GPE_31      LGAVLGAALLLAALGLSAWACMTCWRRRSWRAVKSRASATGPTYIRVADSELYADWSSDS 477
HSV2GPE_32      LGAVLGAALLLAALGLSAWACMTCWRRRSWRAVKSRASATGPTYIRVADSELYADWSSDS 407
HSV2GPE_22      LGAVLGAALLLAALGLSAWACMTCWRRRSWRAVKSRASATGPTYIRVADSELYADWSSDS 474
HSV2GPE_23      LGAVLGAALLLAALGLSAWACMTCWRRRSWRAVKSRASATGPTYIRVADSELYADWSSDS 474
HSV2GPE_24      LGAVLGAALLLAALGLSAWACMTCWRRRSWRAVKSRASATGPTYIRVADSELYADWSSDS 477
HSV2GPE_27      LGAVLGAALLLAALGLSAWACMTCWRRRSWRAVKSRASATGPTYIRVADSELYADWSSDS 477
HSV2GPE_33      LGAVLGAALLLAALGLSAWACMTCWRRRSWRAVKSRASATGPTYIRVADSELYADWSSDS 477
HSV2GPE_21      LGAVLGAALLLAALGLSAWACMTCWRRRSWRAVKSRASATGPTYIRVADSELYADWSSDS 474
HSV2GPE_30      LGAVLGAALLLAALGLSAWACMTCWRRRSWRAVKSRASATGPTYIRVADSELYADWSSDS 407
HSV1GPE_3       LGAVMGAALLLSALGLSVWACMTCWRRRAWRAVKSRASGKGPTYIRVADSELYADWSSDS 479
HSV1GPE_4       LGAVMGAALLLSALGLSVWACMTCWRRRAWRAVKSRASGKGPTYIRVADSELYADWSSDS 479
HSV1GPE_5       LGAVMGAALLLSALGLSVWACMTCWRRRAWRAVKSRASGKGPTYIRVADSELYADWSSDS 479
HSV1GPE_10      LGAVMGAALLLSALGLSVWACMTCWRRRAWRAVKSRASGKGPTYIRVADSELYADWSSDS 479
HSV1GPE_1       LGAVMGAALLLSALGLSVWACMTCWRRRAWRAVKSRASGKGPTYIRVADSELYADWSSDS 409
HSV1GPE_2       LGAVMGAALLLSALGLSVWACMTCWRRRAWRAVKSRASGKGPTYIRVADSELYADWSSDS 479
HSV1GPE_7       LGAVMGAALLLSALGLSVWACMTCWRRRAWRAVKSRASGKGPTYIRVADSELYADWSSDS 479
HSV1GPE_12      LGAVMGAALLLSALGLSVWACMTCWRRRAWRAVKSRASGKGPTYIRVADSELYADWSSDS 479
HSV1GPE_19      LGAVMGAALLLSALGLSVWACMTCWRRRAWRAVKSRASGKGPTYIRVADSELYADWSSDS 479
HSV1GPE_8       LGAVMGAALLLSALGLSVWACMTCWRRRAWRAVKSRASGKGPTYIRVADSELYADWSSDS 479
HSV1GPE_14      LGAVMGAALLLSALGLSVWACMTCWRRRAWRAVKSRASGKGPTYIRVADSELYADWSSDS 479
HSV1GPE_13      LGAVMGAALLLSALGLSVWACMTCWRRRAWRAVKSRASGKGPTYIRVADSELYADWSSDS 479
HSV1GPE_6       LGAVMGAALLLSALGLSVWACMTCWRRRAWRAVKSRASGKGPTYIRVADSELYADWSSDS 479
HSV1GPE_9       LGAVMGAALLLSALGLSVWACMTCWRRRAWRAVKSRASGKGPTYIRVADSELYADWSSDS 479
HSV1GPE_16      LGAVMGAALLLSALGLSVWACMTCWRRRAWRAVKSRASGKGPTYIRVADSELYADWSSDS 479
HSV1GPE_20      LGAVMGAALLLSVLGLSVWACMTCWRRRAWRAVKSRASGKGPTYIRVADSELYADWSSDS 479
HSV1GPE_17      LGAVMGAALLLSALGLSVWACMTCWRRRAWRAVKSRASGKGPTYIRVADSELYADWSSDS 479
HSV1GPE_18      LGAVMGAALLLSALGLSVWACMTCWRRRAWRAVKSRASGKGPTYIRVADSELYADWSSDS 479
HSV1GPE_15      LGAVMGAALLLSALGLSVWACMTCWRRRAWRAVKSRASGKGPTYIRVADSELYADWSSDS 479
HSV1GPE_11      LGAVMGAALLLSALGLSVWACMTCWRRRAWRAVKSRASGKGPTYIRVADSELYADWSSDS 479
                * **:******:.****.**********:*********..********************

HSV2GPE_29      EGERDGSLWQDPPERPDSPSTNGSGFEILSPTAPSVYPHSEGRKSRRPLTTFGSGSPGRR 537
HSV2GPE_39      EGERDGSLWQDPPERPDSPSTNGSGFEILSPTAPSVYPHSEGRKSRRPLTTFGSGSPGRR 537
HSV2GPE_26      EGERDGSLWQDPPERPDSPSTNGSGFEILSPTAPSVYPHSEGRKSRRPLTTFGSGSPGRR 537
HSV2GPE_37      EGERDGSLWQDPPERPDSPSTNGSGFEILSPTAPSVYPHSEGRKSRRPLTTFGSGSPGRR 537
HSV2GPE_38      EGERDGSLWQDPPERPDSPSTNGSGFEILSPTAPSVYPHSEGRKSRRPLTTFGSGSPGRR 537
HSV2GPE_28      EGERDGSLWQDPPERPDSPSTNGSGFEILSPTAPSIYPHSEGRKSRRPLTTFGSGSPGRR 537
HSV2GPE_34      EGERDGSLWQDPPERPDSPSTNGSGFEILSPTAPSVYPHSEGRKSRRPLTTFGSGSPGRR 467
HSV2GPE_25      EGERDGSLWQDPPERPDSPSTNGSGFEILSPTAPSVYPHSEGRKSRRPLTTFGSGSPGRR 537
HSV2GPE_36      EGERDGSLWQDPPERPDSPSTNGSGFEILSPTAPSVYPHSEGRKSRRPLTTFGSGSPGRR 537
HSV2GPE_40      EGERDGSLWQDPPERPDSPSTNGSGFEILSPTAPSVYPHSEGRKSRRPLTTFGSGSPGRR 467
HSV2GPE_35      EGERDGSLWQDPPERPDSPSTNGSGFEILSPTAPSVYPHSEGRKSRRPLTTFGSGSPGRR 537
HSV2GPE_31      EGERDGSLWQDPPERPDSPSTNGSGFEILSPTAPSVYPHSEGRKSRRPLTTFGSGSPGRR 537
HSV2GPE_32      EGERDGSLWQDPPERPDSPSTNGSGFEILSPTAPSVYPHSEGRKSRRPLTTFGSGSPGRR 467
HSV2GPE_22      EGERDGSLWQDPPERPDSPSTNGSGFEILSPTAPSVYPHSEGRKSRRPLTTFGSGSPGRR 534
HSV2GPE_23      EGERDGSLWQDPPERPDSPSTNGSGFEILSPTAPSVYPHSEGRKSRRPLTTFGSGSPGRR 534
HSV2GPE_24      EGERDGSLWQDPPERPDSPSTNGSGFEILSPTAPSVYPHSEGRKSRRPLTTFGSGSPGRR 537
HSV2GPE_27      EGERDGSLWQDPPERPDSPSTNGSGFEILSPTAPSVYPHSEGRKSRRPLTTFGSGSPGRR 537
HSV2GPE_33      EGERDGSLWQDPPERPDSPSTNGSGFEILSPTAPSVYPHSEGRKSRRPLTTFGSGSPGRR 537
HSV2GPE_21      EGERDGSLWQDPPERPDSPSTNGSGFEILSPTAPSVYPHSEGRKSRRPLTTFGSGSPGRR 534
HSV2GPE_30      EGERDGSLWQDPPERPDSPSTNGSGFEILSPTAPSVYPHSEGRKSRRPLTTFGSGSPGRR 467
HSV1GPE_3       EGERDQVPWLAPPERPDSPSTNGSGFEILSPTAPSVYPRSDGHQSRRQLTTFGSGRPDRR 539
HSV1GPE_4       EGERDQVPWLAPPERPDSPSTNGSGFEILSPTAPSVYPRSDGHQSRRQLTTFGSGRPDRR 539
HSV1GPE_5       EGERDQVPWLAPPERPDSPSTNGSGFEILSPTAPSVYPRSDGHQSRRQLTTFGSGRPDRR 539
HSV1GPE_10      EGERDQVPWLAPPERPDSPSTNGSGFEILSPTAPSIYPRSDGHQSRRQLTTFGSGRPDRR 539
HSV1GPE_1       EGERDQVPWLAPPERPDSPSTNGSGFEILSPTAPSVYPRSDGHQSRRQLTTFGSGRPDRR 469
HSV1GPE_2       EGERDQVPWLAPPERPDSPSTNGSGFEILSPTAPSVYPRSDGHQSRRQLTTFGSGRPDRR 539
HSV1GPE_7       EGERDQVPWLAPPERPDSPSTNGSGFEILSPTAPSVYPRSDGHQSRRQLTTFGSGRPDRR 539
HSV1GPE_12      EGERDQVPWLAPPERPDSPSTNGSGFEILSPTAPSVYPRSDGHQSRRQLTTFGSGRPDRR 539
HSV1GPE_19      EGERDQVPWLAPPERPDSPSTNGSGFEILSPTAPSVYPRSDGHQSRRQLTTFGSGRPDRR 539
HSV1GPE_8       EGERDQVPWLAPPERPDSPSTNGSGFEILSPTAPSVYPRSDGHQSRRQLTTFGSGRPDRR 539
HSV1GPE_14      EGERDQVPWLAPPERPDSPSTNGSGFEILSPTAPSVYPRSDGHQSRRQLTTFGSGRPDRR 539
HSV1GPE_13      EGERDQVPWLAPPERPDSPSTNGSGFEILSPTAPSVYPRSDGHQSRRQLTTFGSGRPDRR 539
HSV1GPE_6       EGERDQVPWLAPPERPDSPSTNGSGFEILSPTAPSVYPRSDGHQSRRQLTTFGSGRPDRR 539
HSV1GPE_9       EGERDQVPWLAPPERPDSPSTNGSGFEILSPTAPSVYPRSDGHQSRRQLTTFGSGRPDRR 539
HSV1GPE_16      EGERDQVPWLAPPERPDSPSTNGSGFEILSPTAPSVYPRSDGHQSRRQLTTFGSGRPDRR 539
HSV1GPE_20      EGERDQVPWLAPPERPDSPSTNGSGFEILSPTAPSVYPRSDGHQSRRQLTTFGSGRPDRR 539
HSV1GPE_17      EGERDQVPWLAPPERPDSPSTNGSGFEILSPTAPSVYPRSDGHQSRRQLTTFGSGRPDRR 539
HSV1GPE_18      EGERDQVPWLAPPERPDSPSTNGSGFKILSPTAPSVYPRSDGHQSRRQLTTFGSGRPDRR 539
HSV1GPE_15      EGERDQVPWLAPPERPDSPSTNGSGFEILSPTAPSVYPRSDGHQSRRQLTTFGSGRPDRR 539
HSV1GPE_11      EGERDQVPWLAPPERPDSPSTNGSGFEILSPTAPSVYPRSDGHQSRRQLTTFGSGRPDRR 539
                *****   *  ***************:********:**:*:*::*** ******* *.**

HSV2GPE_29      HSQASYSSVLW 548
HSV2GPE_39      HSQASY----- 543
HSV2GPE_26      HSQASYSSVLW 548
HSV2GPE_37      HSQASYSSVLW 548
HSV2GPE_38      HSQASYSSVLW 548
HSV2GPE_28      HSQASYSSVLW 548
HSV2GPE_34      HSQASYSSVLW 478
HSV2GPE_25      HSQASYSSVLW 548
HSV2GPE_36      HSQASYSSVLW 548
HSV2GPE_40      HSQASYSSVLW 478
HSV2GPE_35      HSQASYSSVLW 548
HSV2GPE_31      HSQASYSSVLW 548
HSV2GPE_32      HSQASYSSVLW 478
HSV2GPE_22      HSQASYPSVLW 545
HSV2GPE_23      HSQASYPSVLW 545
HSV2GPE_24      HSQASYSSVLW 548
HSV2GPE_27      HSQASYSSVLW 548
HSV2GPE_33      HSQASYSSVLW 548
HSV2GPE_21      HSQASYPSVLW 545
HSV2GPE_30      HSQASYSSVLW 478
HSV1GPE_3       YSQASDSSVFW 550
HSV1GPE_4       YSQASDSSVFW 550
HSV1GPE_5       YSQASDSSVFW 550
HSV1GPE_10      YSQASDSSVFW 550
HSV1GPE_1       YSQASDSSVFW 480
HSV1GPE_2       YSQASDSSVFW 550
HSV1GPE_7       YSQASDSSVFW 550
HSV1GPE_12      YSQASDSSVFW 550
HSV1GPE_19      YSQASDSSVFW 550
HSV1GPE_8       YSQASDSSVFW 550
HSV1GPE_14      YSQASDSSVFW 550
HSV1GPE_13      YSQASDSSVFW 550
HSV1GPE_6       YSQASDSSVFW 550
HSV1GPE_9       YSQASDSSVFW 550
HSV1GPE_16      YSQASDSSVFW 550
HSV1GPE_20      YSQASDSSVFW 550
HSV1GPE_17      YSQASDSSVFW 550
HSV1GPE_18      YSQASDSSVFW 550
HSV1GPE_15      YSQASDSSVFW 550
HSV1GPE_11      YSQASDSSVFW 550
                :****
```
