## Supplementary file 11 for "Contriving a chimeric polyvalent vaccine to prevent infections caused by Herpes Simplex Virus (Type-1 and Type-2): an exploratory immunoinformatic approach"

```
HSV2GPG_7       MHAIAPRLLLLFVLSGLPGTRGGSGVPGPINPPNSDVVFPGGSPVAQYCYAYPRLDDPGP 60
HSV2GPG_9       MHAIAPRLLLLFVLSGLPGTRGGSGVPGPINPPNSDVVFPGGSPVAQYCYAYPRLDDPGP 60
HSV2GPG_8       MHAIAPRLLLLFVLSGLPGTRGGSGVPGPINPPNSDVVFPGGSPVAQYCYAYPRLDDPGP 60
HSV2GPG_10      MHAIAPRLLLLFVLSGLPGTRGGSGVPGPINPPNSDVVFPGGSPVAQYCYAYPRLDDPGP 60
HSV2GPG_13      ------------------------------------------------------------
HSV2GPG_14      MHAIAPRLLLLFVLSGLPGTRGGSGVPGPINPPNSDVVFPGGAPVAQYCYAYPRLDDPGP 60
HSV2GPG_23      ------------------------------------------------------------
HSV2GPG_11      ------------------------------------------------------------
HSV2GPG_25      MHAIAPRLLLLFVLSGLPGTRGGSGVPGPINPPNNDVVFPGGSPVAQYCYAYPRLDDPGP 60
HSV2GPG_26      MHAIAPRLLLLFVLSGLPGTRGGSGVPGPINPPNNDVVFPGGSPVAQYCYAYPRLDDPGP 60
HSV2GPG_20      MHAIAPRLLLLFVLSGLPGTRGGSGVPGPINPPNNDVVFPGGSPVAQYCYAYPRLDDPGP 60
HSV2GPG_19      MHAIAPRLLLLFVLSGLPGTRGGSGVPGPINPPNNDVVFPGGSPVAQYCYAYPRLDDPGP 60
HSV2GPG_17      MHAIAPRLLLLFVLSGLPGTRGGSGVPGPINPPNNDVVFPGGSPVAQYCYAYPRLDDPGP 60
HSV2GPG_21      ------------------------------------------------------------
HSV2GPG_24      MHAIAPRLLLLFVLSGLPGTRGGSGVPGPINPPNNDVVFPGGSPVAQYCYAYPRLDDPGP 60
HSV2GPG_16      ------------------------------------------------------------
HSV2GPG_18      MHAIAPRLLLLFVLSGLPGTRGGSGVPGPINPPNNDVVFPGGSPVAQYCYAYPRLDDPGP 60
HSV2GPG_12      MHAIAPRLLLLFVLSGLPGTRGGSGVPGPINPPNNDVVFPGGSPVAQYCYAYPRLDDPGP 60
HSV2GPG_15      MHAIAPRLLLLFVLSGLPGTRGGSGVPGPINPPNNDVVFPGGSPVAQYCYAYPRLDDPGP 60
HSV2GPG_22      ------------------------------------------------------------


HSV2GPG_7       LGSADAGRQDLPRRVVRHEPLGRSFLTGGLVLLAPPVRGFGAPNATYAARVTYYRLTRAC 120
HSV2GPG_9       LGSADAGRQDLPRRVVRHEPLGRSFLTGGLVLLAPPVRGFGAPNATYAARVTYYRLTRAC 120
HSV2GPG_8       LGSADAGRQDLPRRVVRHEPLGRSFLTGGLVLLAPPVRGFGAPNATYAARVTYYRLTRAC 120
HSV2GPG_10      LGSADAGRQDLPRRVVRHEPLGRSFLTGGLVLLAPPVRGFGAPNATYAARVTYYRLTRAC 120
HSV2GPG_13      ----------LPRRVVRHEPLGRSFLTGGLVLLAPPVRGFGAPNATYAARVTYYRLTRAC 50
HSV2GPG_14      LGSADAGRQDLPRRVVRHEPLGRSFLTGGLVLLAPPVRGFGAPNATYAARVTYYRLTRAC 120
HSV2GPG_23      ----------LPRRVVRHEPLGRSFLTGGLVLLAPPVRGFGAPNATYAARVTYYRLTRAC 50
HSV2GPG_11      ----------LPRRVVRHEPLGRSFLTGGLVLLAPPVRGFGAPNATYAARVTYYRLTRAC 50
HSV2GPG_25      LGSADAGRQDLPRRVVRHEPLGRSFLTGGLVLLAPPVRGFGAPNATYAARVTYYRLTRAC 120
HSV2GPG_26      LGSADAGRQDLPRRVVRHEPLGRSFLTGGLVLLAPPVRGFGAPNATYAARVTYYRLTRAC 120
HSV2GPG_20      LGSADAGRQDLPRRVVRHEPLGRSFLTGGLVLLAPPVRGFGAPNATYAARVTYYRLTRAC 120
HSV2GPG_19      LGSADAGRQDLPRRVVRHEPLGRSFLTGGLVLLAPPVRGFGAPNATYAARVTYYRLTRAC 120
HSV2GPG_17      LGSADAGRQDLPRRVVRHEPLGRSFLTGGLVLLAPPVRGFGAPNATYAARVTYYRLTRAC 120
HSV2GPG_21      ----------LPRRVVRHEPLGRSFLTGGLVLLAPPVRGFGAPNATYAARVTYYRLTRAC 50
HSV2GPG_24      LGSADAGRQDLPRRVVRHEPLGRSFLTGGLVLLAPPVRGFGAPNATYAARVTYYRLTRAC 120
HSV2GPG_16      ----------LPRRVVRHEPLGRSFLTGGLVLLAPPVRGFGAPNATYAARVTYYRLTRAC 50
HSV2GPG_18      LGSADAGRQDLPRRVVRHEPLGRSFLTGGLVLLAPPVRGFGAPNATYAARVTYYRLTRAC 120
HSV2GPG_12      LGSADAGRQDLPRRVVRHEPLGRSFLTGGLVLLAPPVRGFGAPNATYAARVTYYRLTRAC 120
HSV2GPG_15      LGSADAGRQDLPRRVVRHEPLGRSFLTGGLVLLAPPVRGFGAPNATYVARVTYYRLTRAC 120
HSV2GPG_22      ----------LPRRVVRHEPLGRSFLTGGLVLLAPPVRGFGAPNATYAARVTYYRLTRAC 50
                          *************************************.************

HSV2GPG_7       RQPILLRQYGGCRGGEPPSPKTCGSYTYTYQGGGPPTRYALVNASLLVPIWDRAAETFEY 180
HSV2GPG_9       RQPILLRQYGGCRGGEPPSPKTCGSYTYTYQGGGPPTRYALVNASLLVPIWDRAAETFEY 180
HSV2GPG_8       RQPILLRQYGGCRGGEPPSPKTCGSYTYTYQGGGPPTRYALVNASLLVPIWDRAAETFEY 180
HSV2GPG_10      RQPILLRQYGGCRGGEPPSPKTCGSYTYTYQGGGPPTRYALVNASLLVPIWDRAAETFEY 180
HSV2GPG_13      RQPILLRQYGGCRGGEPPSPKTCGSYTYTYQGGGPPTRYALVNASLLVPIWDRAAETFEY 110
HSV2GPG_14      RQPILLRQYGGCRGGEPPSPKTCGSYTYTYQGGGPPTRYALVNASLLVPIWDRAAETFEY 180
HSV2GPG_23      RQPILLRQYGGCRGGEPPSPKTCGSYTYTYQGGGPPTRYALVNASLLVPIWDRAAETFEY 110
HSV2GPG_11      RQPILLRQYGGCRGGEPPSPKTCGSYTYTYQGGGPPTRYALVNASLLVPIWDRAAETFEY 110
HSV2GPG_25      RQPILLRQYGGCRGGEPPSPKTCGSYTYTYQGGGPPTRYALVNASLLVPIWDRAAETFEY 180
HSV2GPG_26      RRPILLRQYGGCRGGEPPSPKTCGSYTYTYQGGGPPTRYALVNASLLVPIWDRAAETFEY 180
HSV2GPG_20      RQPILLRQYGGCRGGEPPSPKTCGSYTYTYQGGGPPTRYALVNASLLVPIWDRAAETFEY 180
HSV2GPG_19      RQPILLRQYGGCRGGEPPSPKTCGSYTYTYQGGGPPTRYALVNASLLVPIWDRAAETFEY 180
HSV2GPG_17      RQPILLRQYGGCRGGEPPSPKTCGSYTYTYQGGGPPTRYALVNASLLVPIWDRAAETFEY 180
HSV2GPG_21      RQPILLRQYGGCRGGEPPSPKTCGSYTYTYQGGGPPTRYALVNASLLVPIWDRAAETFEY 110
HSV2GPG_24      RQPILLRQYGGCRGGEPPSPKTCGSYTYTYQGGGPPTRYALVNASLLVPIWDRAAETFEY 180
HSV2GPG_16      RQPILLRQYGGCRGGEPPSPKTCGSYTYTYQGGGPPTRYALVNASLLVPIWDRAAETFEY 110
HSV2GPG_18      RQPILLRQYGGCRGGEPPSPKTCGSYTYTYQGGGPPTRYALVNASLLVPIWDRAAETFEY 180
HSV2GPG_12      RQPILLRQYGGCRGGEPPSPKTCGSYTYTYQGGGPPTRYALVNASLLVPIWDRAAETFEY 180
HSV2GPG_15      RQPILLRQYGGCRGGEPPSPKTCGSYTYTYQGGGPPTRYALVNASLLVPIWDRAAETFEY 180
HSV2GPG_22      RQPILLRQYGGCRGGEPPSPKTCGSYTYTYQGGGPPTRYALVNASLLVPIWDRAAETFEY 110
                *:**********************************************************

HSV2GPG_7       QIELGGELHVGLLWVEVGGEGPGPTAPPQAARAEGGPCVPPVPAGRPWRSVPPVWYSAPN 240
HSV2GPG_9       QIELGGELHVGLLWVEVGGEGPGPTAPPQAARAEGGPCVPPVPAGRPWRSVPPVWYSAPN 240
HSV2GPG_8       QIELGGELHVGLLWVEVGGEGPGPTAPPQAARAEGGPCVPPVPAGRPWRSVPPVWYSAPN 240
HSV2GPG_10      QIELGGELHVGLLWVEVGGEGPGPTAPPQAARAEGGPCVPPVPAGRPWRSVPPVWYSAPN 240
HSV2GPG_13      QIELGGELHVGLLWVEVGGEGPGPTAPPQAARAEGGPCVPPVPAGRPWRSVPPVWYSAPN 170
HSV2GPG_14      QIELGGELHVGLLWVEVGGEGPGPTAPPQAARAEGGPCVPPVPAGRPWRSVPPVWYSAPN 240
HSV2GPG_23      QIELGGELHVGLLWVEVGGEGPGPTAPPQAARAEGGPCVPPVPAGRPWRSVPPVWYSAPN 170
HSV2GPG_11      QIELGGELHVGLLWVEVGGEGPGPTAPPQAARAEGGPCVPPVPAGRPWRSVPPVWYSAPN 170
HSV2GPG_25      QIELGGELHVGLLWVEVGGEGPGLTAPPQAARAEGGPCVPPVPAGRPWRSVPPVWYSAPN 240
HSV2GPG_26      QIELGGELHVGLLWVEVGGEGPGPTAPPQAARAEGGPCVPPVPAGRPWRSVPPVWYSAPN 240
HSV2GPG_20      QIELDGELHVGLLWVEVGGEGPGPTAPPQAARAEGGPCVPPVPAGRPWRSVPPVWYSAPN 240
HSV2GPG_19      QIELGGELHVGLLWVEVGGEGPGPTAPPQAARAEGGPCVPPVPAGRPWRSVPPVWYSAPN 240
HSV2GPG_17      QIELGGELHVGLLWVEVGGEGPGPTAPPQAARAEGGPCVPPVPAGRPWRSVPPVWYSAPN 240
HSV2GPG_21      QIELGGELHVGLLWVEVGGEGPGPTAPPQAARAEGGPCVPPVPAGRPWRSVPPVWYSAPN 170
HSV2GPG_24      QIELGGELHVGLLWVEVGGEGPGPTAPPQAARAEGGPCVPPVPAGRPWRSVPPVWYSAPN 240
HSV2GPG_16      QIELGGELHVGLLWVEVGGEGPGPTAPPQAARAEGGPCVPPVPAGRPWRSVPPVWYSAPN 170
HSV2GPG_18      QIELGGELHVGLLWVEVGGEGPGPTAPPQAARAEGGPCVPPVPAGRPWRSVPPVWYSAPN 240
HSV2GPG_12      QIELGGELHVGLLWVEVGGEGPGPTAPPQAARAEGGPCVPPVPAGRPWRSVPPVWYSAPN 240
HSV2GPG_15      QIELGGELHVGLLWVEVGGEGPGPTAPPQAARAEGGPCVPPVPAGRPWRSVPPVWYSAPN 240
HSV2GPG_22      QIELGGELHVGLLWVEVGGEGPGPTAPPQAAHAEGGPCVPPVPAGRPWRSVPPVWYSAPN 170
                ****.****************** *******:****************************

HSV2GPG_7       PGFRGLRFRERCLPPQTPAAPSDLPRVAFAPQSLLVGITGRTFIRMARPTEDVGVLPPHW 300
HSV2GPG_9       PGFRGLRFRERCLPPQTPAAPSDLPRVAFAPQSLLVGITGRTFIRMARPTEDVGVLPPHW 300
HSV2GPG_8       PGFRGLRFRERCLPPQTPAAPSDLPRVAFAPQSLLVGITGRTFIRMARPTEDVGVLPPHW 300
HSV2GPG_10      PGFRGLRFRERCLPPQTPAAPSDLPRVAFAPQSLLVGITGRTFIRMARPTEDVGVLPPHW 300
HSV2GPG_13      PGFRGLRFRERCLPPQTPAAPSDLPRVAFAPQSLLVGITGRTFIRMARPTEDVGVLPPHW 230
HSV2GPG_14      PGFRGLRFRERCLPPQTPAAPSDLPRVAFAPQSLLVGITGRTFIRMARPTEDVGVLPPHW 300
HSV2GPG_23      PGFRGLRFRERCLPPQTPAAPSDLPRVAFAPQSLLVGITGRTFIRMARPTGDVGVLPPHW 230
HSV2GPG_11      PGFRGLRFRERCLPPQTPAAPSDLPRVAFAPQSLLVGITGRTFIRMARPTEDVGVLPPHW 230
HSV2GPG_25      PGFRGLRFRERCLPPQTPAAPSDLPRVAFAPQSLLVGITGRTFIRMARPTEDVGVLPPHW 300
HSV2GPG_26      PGFRGLRFRERCLPPQTPAAPSDLPRVAFAPQSLLVGITGRTFIRMARPTEDVGVLPPHW 300
HSV2GPG_20      PGFRGLRFRERCLPPQTPAAPSDLPRVAFAPQSLLVGITGRTFIRMARPTEDVGVLPPHW 300
HSV2GPG_19      PGFRGLRFRERCLPPQTPAAPSDLPRVAFAPQSLLVGITGRTFIRMARPTEDVGVLPPHW 300
HSV2GPG_17      PGFRGLRFRERCLPPQTPAAPSDLPRVAFAPQSLLVGITGRTFIRMARPTEDVGVLPPHW 300
HSV2GPG_21      PGFRGLRFRERCLPPQTPAAPSDLPRVAFAPQSLLVGITGRTFIRMARPTEDVGVLPPHW 230
HSV2GPG_24      PGFRGLRFRERCLPPQTPAAPSDLPRVAFAPQSLLVGITGRTFIRMARPTEDVGVLPPHW 300
HSV2GPG_16      PGFRGLRFRERCLPPQTPAAPSDLPRVAFAPQSLLVGITGRTFIRMARPTEDVGVLPPHW 230
HSV2GPG_18      PGFRGLRFRERCLPPQTPAAPSDLPRVAFAPQSLLVGITGRTFIRMARPTEDVGVLPPHW 300
HSV2GPG_12      PGFRGLRFRERCLPPQTPAAPSDLPRVAFAPQSLLVGITGRTFIRMARPTEDVGVLPPHW 300
HSV2GPG_15      PGFRGLRFRERCLPPQTPAAPSDLPRVAFAPQSLLVGITGRTFIRMARPTEDVGVLPPHW 300
HSV2GPG_22      PGFRGLRFRERCLPPQTPAAPSDLPRVAFAPQSLLVGITGRTFIRMARPTEDVGVLPPHW 230
                ************************************************** *********

HSV2GPG_7       APGALDDGPYAPFPPRPRFRRALRTDPEGVDPDVRAPRTGRRLMALTEDTSSDSPTSAPE 360
HSV2GPG_9       APGALDDGPYAPFPPRPRFRRALRTDPEGVDPDVRAPRTGRRLMALTEDASSDSPTSAPE 360
HSV2GPG_8       APGALDDGPYAPFPPRPRFRRALRTDPEGVDPDVRAPRTGRRLMALTEDASSDSPTSAPE 360
HSV2GPG_10      APGALDDGPYAPFPPRPRFRRALRTDPEGVDPDVRAPRTGRRLMALTEDASSDSPTSAPE 360
HSV2GPG_13      APGALDDGPYAPFPPRPRFRRALRTDPEGVDPDVRAPRTGRRLMALTEDASSDSPTSAPE 290
HSV2GPG_14      APGALDDGPYAPFPPRPRFRRALRTDPEGVDPDVRAPRTGRRLMALTEDASSDSPTSAPE 360
HSV2GPG_23      APGALDDGPYAPFPPRPRFRRALRTDPEGVDPDVRAPRTGRRLMALTEDASSDSPTSAPE 290
HSV2GPG_11      APGALDDGPYAPFPPRPRFRRALRTDPEGVDPDVRAPRTGRRLMALTEDASSDSPTSAPE 290
HSV2GPG_25      APGALDDGPYAPFPPRPRFRRALRTDPEGVDPDVRAPRTGRRLMALTEDASSDSPTSAPE 360
HSV2GPG_26      APGALNDGPYAPFPPRPRFRRALRTDPEGVDPDVRAPRTGRRLMALTEDASSDSPTSAPE 360
HSV2GPG_20      APGALDDGPYAPFPPRPRFRRALRTDPEGVDPDVRAPRTGRRLMALTEDASSDSPTSAPE 360
HSV2GPG_19      APGALDDGPYAPFPPRPRFRRALRTDPEGVDPDVRAPRTGRRLMALTEDASSDSPTSAPE 360
HSV2GPG_17      APGALDDGPYAPFPPRPRFRRALRTDPEGVDPDVRAPRTGRRLMALTEDASSDSPTSAPE 360
HSV2GPG_21      APGALDDGPYAPFPPRPRFRRALRTDPEGVDPDVRAPLTGRRLMALTEDASSDSPTSAPE 290
HSV2GPG_24      APGALDDGPYAPFPPRPRFRRALRTDPEGVDPDVRAPLTGRRLMALTEDASSDSPTSAPE 360
HSV2GPG_16      APGALDDGPYAPFPPRPRFRRALRTDPKGVDPDVRAPRTGRRLMALTEDASSDSPTSAPE 290
HSV2GPG_18      APGALDDGPYAPFPPRPRFRRALRTDPKGVDPDVRAPRTGRRLMALTEDASSDSPTSAPE 360
HSV2GPG_12      APGALDDGPYAPFPPRPRFRRALRTDPEGVDPDVRAPRTGRRLMALTENASSDSPTSAPE 360
HSV2GPG_15      APGALDDGPYAPFPPRPRFRRALRTDPEGVDPDVRAPRTGRRLMALTENASSDSPTSAPE 360
HSV2GPG_22      APGALDDGPYAPFPPRPRFRRALRTDPEGVDPDVRAPRTGRRLMALTENASSDSPTSAPE 290
                *****:*********************:********* **********::**********

HSV2GPG_7       KTPLPVSATAMAPSVDPSAEPTAPATTTPPDEMATQAATVAVTPEETAVASPPATASVES 420
HSV2GPG_9       KTPLPVSATAMAPSVDPSAEPTAPATTTPPDEMAAQAATVAVTPEETAVASPPATASVES 420
HSV2GPG_8       KTPLPVSATAMAPSVDPSAEPTAPATTTPPDEMATQAATVAVTPEETAVASPPATASVES 420
HSV2GPG_10      KTPLPVSATAMAPSVDPSAEPTAPATTTPPDEMATQAATVAVTPEETAVASPPATASVES 420
HSV2GPG_13      KTPLPVSATAMAPSVDPSAEPTAPATTTPPDEMATQAATVAVTPDETAVASPPATASVES 350
HSV2GPG_14      KTPLPVSATAMAPSVDPSAEPTAPATTTPPDEMATQAATVAVTPDETAVASPPATASVES 420
HSV2GPG_23      KTPLPVSATAMAPSVDPSAEPTAPATTTPPDEMATQAATVAVTPEETAVASPPATASVES 350
HSV2GPG_11      KTPLPVSATAMAPSVDPSAEPTAPATTTPPDEMATQAATVAVTPEETAVASPPATASVES 350
HSV2GPG_25      KTPLPVSATAMAPSVDPSAEPTAPATTTPPDEMATQAATVAVTPEETAVASPPATASVES 420
HSV2GPG_26      KTPLPVSATAMAPSVDPSAEPTAPATTTPPDEMATQAATVAVTPEETAVASPPATASVES 420
HSV2GPG_20      KTPLPVSATAMAPSVDPSAEPTAPATTTPPDEMATQAATVAVTPEETAVASPPATASVES 420
HSV2GPG_19      KTPLPVSATAMAPSVDPSAEPTAPATTTPPDEMATQAATVAVTPEETAVASPPATASVES 420
HSV2GPG_17      KTPLPVSATAMAPSVDPSAEPTAPATTTPPDEMATQAATVAVTPEETAVASPPATASVES 420
HSV2GPG_21      KTPLPVSATAMAPSVDPSAEPTAPATTTPPDEMATQAATVAVTPEETAVASPPATASVES 350
HSV2GPG_24      KTPLPVSATAMAPSVDPSAEPTAPTTTTPPDEMATQAATVAVTPEETAVASPPATASVES 420
HSV2GPG_16      KTPLPVSATAMAPSVDPSAEPTAPATTTPPDEMATQAATVAVTPEETAVASPPATASVES 350
HSV2GPG_18      KTPLPVSATAMAPSVDPSAEPTAPATTTPPDEMATQAATVAVTPEETAVASPPATASVES 420
HSV2GPG_12      KTPLPVSATAMAPSVDPSAEPTAPATTTPPDEMATQAATVAVTPEETAVASPPATASVES 420
HSV2GPG_15      KTPLPVSATAMAPSVDPSAEPTAPATTTPPDEMATQAATVAVTPEETAVASPPATASVES 420
HSV2GPG_22      KTPLPVSATAMAPSVDPSAEPTAPATTTPPDEMATQAATVAVTPEETAVASPPATASVES 350
                ************************:*********:*********:***************

HSV2GPG_7       SPLPAAAAATPGAGHTNTSSASAAKTPPTTPAPTTPPPTSTHATPRPTTPGPQTTPPGPA 480
HSV2GPG_9       SPLPAAAAATPGAGHTNTSSASAAKTPPTTPAPTTPPPTSTHATPRPTTPGPQTTPPGPA 480
HSV2GPG_8       SPLPAAAAATPGAGHTNTSSASAAKTPPTTPAPTTPPPTSTHATPRPTTPGPQTTPPGPA 480
HSV2GPG_10      SPLP-AAAATPGAGHTNTSSASAAKTPPTTPAPTTPPPTSTHATPRPTTPGPQTTPPGPA 479
HSV2GPG_13      SPLP-AAAATPGAGHTNTSSASAAKTPPTTPAPTTPPPTSTHATPRPTTPGPQTTPPGPA 409
HSV2GPG_14      SPLP-AAAATPGAGHTNTSSASAAKTPPTTPAPTTPPPTSTHATPRPTTPGPQTTPPGPA 479
HSV2GPG_23      SPLP-AAAATPGAGHTNTSSASAAKTPPTTPAPTTPPPTSTHATPRPTTPGPQTTPPGPA 409
HSV2GPG_11      SPLP-AAAATPGAGHTNTSSASAAKTPPTTPAPTTPPPTSTHATPRPTTPGPQTTPPGPA 409
HSV2GPG_25      SPLP-AAAATPGAGHTNTSSASAAKTPPTTPAPTTPPPTSTHATPRPTTPGPQTTPPGPA 479
HSV2GPG_26      SPLP-AAAATPGAGHTNTSSASAAKTPPTTPAPTTPPPTSTHATPRPTTPGPQTTPPGPA 479
HSV2GPG_20      SPLP-AAAATPGAGHTNTSSASAAKTPPTTPAPTTPPPTSTHATPRPTTPGPQTTPPGPA 479
HSV2GPG_19      SPPP-AAAATPGAGHTNTSSASAAKTPPTTPAPTTPPPTSTHATPRPTTPGPQTTPPGPA 479
HSV2GPG_17      SPLP-AAAATPGAGHTNTSSASAAKTPPTTPAPTTPPPTSTHATPRPTTPGPQTTPPGPA 479
HSV2GPG_21      SPLP-AAAATPGAGHTNTSSASAAKTPPTTPAPTTPPPTSTHATPRPTTPGPQTTPPGPA 409
HSV2GPG_24      SPLP-AAAATPGAGHTNTSSASAAKTPPTTPAPTTPPPTSTHATPRPTTPGPQTTPPGPA 479
HSV2GPG_16      SPLP-AAAATPGAGHTNTSSASAAKTPPTTPAPTTPPPTSTHATPRPTTPGPQTTPPGPA 409
HSV2GPG_18      SPLP-AAAATPGAGHTNTSSASAAKTPPTTPAPTTPPPTSTHATPRPTTPGPQTTPPGPA 479
HSV2GPG_12      SPLP-AAAATPGAGHTNTSSASAAKTPPTTPAPTTPPPTSTHATPRPTTPGPQTTPPGPA 479
HSV2GPG_15      SPLP-AAAATPGAGHTNTSSASAAKTPPTTPAPTTPPPTSTHATPRPTTPGPQTTPPGPA 479
HSV2GPG_22      SPLP-AAAATPGAGHTNTSSASAAKTPPTTPAPTTPPPTSTHATPRPTTPGPQTTPPGPA 409
                ** * *******************************************************

HSV2GPG_7       TPGPVGASAAPTADSPLTASPPATAPGPSAANVSVAATTATPGTRGTARTPPTDPKTHPH 540
HSV2GPG_9       TPGPVGASAAPTADSPLTASPPATAPGPSAANVSVAATTATPGTRGTARTPPTDPKTHPH 540
HSV2GPG_8       TPGPVGASAAPTADSPLTASPPATAPGPSAANVSVAATTATPGTRGTARTPPTDPKTHPH 540
HSV2GPG_10      TPGPVGASAAPTADSPLTASPPATAPGPSAANVSVAATTATPGTRGTARTPPTDPKTHPH 539
HSV2GPG_13      TPGPVGASAAPTADSPLTASPPATAPGPSAANVSVAATTATPGTRGTARTPPTDPKTHPH 469
HSV2GPG_14      TPGPVGASAAPTADSPLTASPPATAPGPSAANVSVAATTATPGTRGTARTPPTDPKTHPH 539
HSV2GPG_23      TPGPVGASAAPTADSPLTASPPATAPGPSAANVSVAATTATPGTRGTARTPPTDPKTHPH 469
HSV2GPG_11      TPGPVGASAAPTADSPLTASPPATAPGPSAANVSVAATTATPGTRGTARTPPTDPKTHPH 469
HSV2GPG_25      TPGPVGASAAPTADSPLTASPPATAPGPSAANVSVAATTATPGTRGTARTPPTDPKTHPH 539
HSV2GPG_26      TPGPVGASAAPTADSPLTASPPATAPGPSAANVSVAATTATPGTRGTARTPPTDPKTHPH 539
HSV2GPG_20      TPGPVGASAAPTADSPLTASPPATAPGPSAANVSVAATTATPGTRGTARTPPTDPKTHPH 539
HSV2GPG_19      TPGPVGASAAPTADSPLTASPPATAPGPSAANVSVAATTATPGTRGTARTPPTDPKTHPH 539
HSV2GPG_17      TPGPVGASAAPTADSPLTASPPATAPGPSAANVSVAATTATPGTRGTARTPPTDPKTHPH 539
HSV2GPG_21      TPGPVGASAAPTADSPLTASPPATAPGPSAANVSVAATTATPGTRGTARTPPTDPKTHPH 469
HSV2GPG_24      TPGPVGASAAPTADSPLTASPPATAPGPSAANVSVAATTATPGTRGTARTPPTDPKTHPH 539
HSV2GPG_16      TPGPVGASAAPTADSPLTASPPATAPGPSAANVSVAATTATPGTRGTARTPPTDPKTHPH 469
HSV2GPG_18      TPGPVGASAAPTADSPLTASPPATAPGPSAANVSVAATTATPGTRGTARTPPTDPKTHPH 539
HSV2GPG_12      TPGPVGASAAPTADSPLTASPPATAPGPSAANVSVAATTATPGTRGTARTPPTDPKTHPH 539
HSV2GPG_15      TPGPVGASAAPTADSPLTASPPATAPGPSAANVSVAATTATPGTRGTARTPPTDPKTHPH 539
HSV2GPG_22      TPGPVGASAAPTADSPLTASPPATAPGPSAANVSVAATTATPGTRGTARTPPTDPKTHPH 469
                ************************************************************

HSV2GPG_7       GPADAPPGSPAPPPPEHRGGPEEFEGAGDGEPPEDDDSATGLAFRTPNPNKPPPARPGPI 600
HSV2GPG_9       GPADAPPGSPAPPPPEHRGGPEEFEGAGDGEPPEDDDSATGLAFRTPNPNKPPPARPGPI 600
HSV2GPG_8       GPADAPPGSPAPPPPEHRGGPEEFEGAGDGEPPEDDDSATGLAFRTPNPNKPPPARPGPI 600
HSV2GPG_10      GPADAPPGSPAPPPPEHRGGPEEFEGAGDGEPPEDDDSATGLAFRTPNPNKPPPARPGPI 599
HSV2GPG_13      GPADAPPGSPAPPPPEHRGGPEEFEGAGDGEPPEDDDSATGLAFRTPNPNKPPPARPGPI 529
HSV2GPG_14      GPADAPPGSPAPPPPEHRGGPEEFEGAGDGEPPEDDDSATGLAFRTPNPNKPPPARPGPI 599
HSV2GPG_23      GPADAPPGSPAPPPPEHRGGPEEFEGAGDGEPPEDDDSATGLAFRTPNPNKPPPARPGPI 529
HSV2GPG_11      GPADAPPGSPAPPPPEHRGGPEEFEGAGDGEPPEDDDSATGLAFRTPNPNKPPPARPGPI 529
HSV2GPG_25      GPADAPPGSPAPPPPEHRGGPEEFEGAGDGEPPEDDDSATGLAFRTPNPNKPPPARPGPI 599
HSV2GPG_26      GPADAPPGSPAPPPPEHRGGPEEFEGAGDGEPPEDDDSATGLAFRTPNPNKPPPARPGPI 599
HSV2GPG_20      GPADAPPGSPAPPPPEHRGGPEEFEGAGDGEPPEDDDSATGLAFRTPNPNKPPPARPGPI 599
HSV2GPG_19      GPADAPPGSPAPPPPEHRGGPEEFEGAGDGEPPEDDDSATGLAFRTPNPNKPPPARPGPI 599
HSV2GPG_17      GPADAPPGSPAPPPPEHRGGPEEFEGAGDGEPPEDDDSATGLAFRTPNPNKPPPARPGPI 599
HSV2GPG_21      GPADAPPGSPAPPPPEHRGGPEEFEGAGDGEPPEDDDSATGLAFRTPNPNKPPPARPGPI 529
HSV2GPG_24      GPADAPPGSPAPPPPEHRGGPEEFEGAGDGEPPEDDDSATGLAFRTPNPNKPPPARPGPI 599
HSV2GPG_16      GPADAPPGSPAPPPPEHRGGPEEFEGAGDGEPPEDDDSATGLAFRTPNPNKPPPARPGPI 529
HSV2GPG_18      GPADAPPGSPAPPPPEHRGGPEEFEGAGDGEPPEDDDSATGLAFRTPNPNKPPPARPGPI 599
HSV2GPG_12      GPADAPPGSPAPPPPEHRGGPEEFEGAGDGEPPEDDDSATGLAFRTPNPNKPPPARPGPI 599
HSV2GPG_15      GPADAPPGSPAPPPPEHRGGPEEFEGAGDGEPPEDDDSATGLAFRTPNPNKPPPARPGPI 599
HSV2GPG_22      GPADAPPGSPAPPPPEHRGGPEEFEGAGDGEPPEDDDSATGLAFRTPNPNKPPPARPGPI 529
                ************************************************************

HSV2GPG_7       RPTLPPGILGPLAPNTPRPPAQAPAKDMPSGPTPQHIPLFWFLTASPALDILFIISTTIH 660
HSV2GPG_9       RPTLPPGILGPLAPNTPRPPAQAPAKDMPSGPTPQHIPLFWFLTASPALDILFIISTTIH 660
HSV2GPG_8       RPTLPPGILGPLAPNTPRPPAQAPAKDMPSGPTPQHIPLFWFLTASPALDILFIISTTIH 660
HSV2GPG_10      RPTLPPGILGPLAPNTPRPPAQAPAKDMPSGPTPQHIPLFWFLTASPALDILFIISTTIH 659
HSV2GPG_13      RPTLPPGILGPLAPNTPRPPAQAPAKDMPSGPTPQHIPLFWFLTASPALDILFIISTTIH 589
HSV2GPG_14      RPTLPPGILGPLAPNTPRPPAQAPAKDMPSGPTPQHIPLFWFLTASPALDILFIISTTIH 659
HSV2GPG_23      RPTLPPGILGPLAPNTPRPPAQAPAKDMPSGPTPQHIPLFWFLTASPALDILFIISTTIH 589
HSV2GPG_11      RPTLPPGILGPLAPNTPRPPAQAPAKDMPSGPTPQHIPLFWFLTASPALDILFIISTTIH 589
HSV2GPG_25      RPTLPPGILGPLAPNTPRPPAQAPAKDMPSGPTPQHIPLFWFLTASPALDILFIISTTIH 659
HSV2GPG_26      RPTLPPGILGPLAPNTPRPPAQAPAKDMPSGPTPQHIPLFWFLTASPALDILFIISTTIH 659
HSV2GPG_20      RPTLPPGILGPLAPNTPRPPAQAPAKDMPSGPTPQHIPLFWFLTASPALDILFIISTTIH 659
HSV2GPG_19      RPTLPPGILGPLAPNTPRPPAQAPAKDMPSGPTPQHIPLFWFLTASPALDILFIISTTIH 659
HSV2GPG_17      RPTLPPGILGPLAPNTPHPPAQAPAKDMPSGPTPQHIPLFWFLTASPALDILFIISTTIH 659
HSV2GPG_21      RPTLPPGILGPLAPNTPRPPAQAPAKDMPSGPTPQHIPLFWFLTASPALDILFIISTTIH 589
HSV2GPG_24      RPTLPPGILGPLAPNTPRPPAQAPAKDMPSGPTPQHIPLFWFLTASPALDILFIISTTIH 659
HSV2GPG_16      RPTLPPGILGPLAPNTPRPPAQAPAKDMPSGPTPQHIPLFWFLTASPALDILFIISTTIH 589
HSV2GPG_18      RPTLPPGILGPLAPNTPRPPTQAPAKDMPSGPTPQHIPLFWFLTASPALDILFIISTTIH 659
HSV2GPG_12      RPTLPPGILGPLAPNTPRPPAQAPAKDMPSGPTPQHIPLFWFLTASPALDILFIISTTIH 659
HSV2GPG_15      RPTLPPGILGPLAPNTPRPPAQAPAKDMPSGPTPQHIPLFWFLTASPALDILFIISTTIH 659
HSV2GPG_22      RPTLPPGILGPLAPNTPRPPAQAPAKDMPSGPTPQHIPLFWFLTASPALDILFIISTTIH 589
                *****************:**:***************************************

HSV2GPG_7       TAAFVCLVALAAQLWRGRAGRRRYAHPSVRYVCLPPERD 699
HSV2GPG_9       TAAFVCLVALAAQLWRGRAGRRRYAHPSVRYVCLPPERD 699
HSV2GPG_8       TAAFVCLVALAAQLWRGRAGRRRYAHPSVRYVCLPPERD 699
HSV2GPG_10      TAAFVCLVALAAQLWRGRAGRRRYAHPSVRYVCLPPERD 698
HSV2GPG_13      TAAFVCLVALAAQLWRGRAGRRRYAHPSVRYVCLPPERD 628
HSV2GPG_14      TAAFVCLVALAAQLWRGRAGRRRYAHPSVRYVCLPPERD 698
HSV2GPG_23      TAAFVCLVALAAQLWRGRAGRRRYAHPSVRYVCLPPERD 628
HSV2GPG_11      TAAFVCLVALAAQLWRGRAGRRRYAHPSVRYVCLPPERD 628
HSV2GPG_25      TAAFVCLVALAAQLWRGRAGRRRYAHPSVRYVCLPPERD 698
HSV2GPG_26      TAAFVCLVALAAQLWRGRAGRRRYAHPSVRYVCLPPERD 698
HSV2GPG_20      TAAFVCLVALAAQLWRGRAGRRRYAHPSVRYVCLPPERD 698
HSV2GPG_19      TAAFVCLVALAAQLWRGRAGRRRYAHPSVRYVCLPPERD 698
HSV2GPG_17      TAAFVCLVALAAQLWRGRAGRRRYAHPSVRYVCLPPERD 698
HSV2GPG_21      TAAFVCLVALAAQLWRGRAGRRRYAHPSVRYVCLPPERD 628
HSV2GPG_24      TAAFVCLVALAAQLWRGRAGRRRYAHPSVRYVCLPPERD 698
HSV2GPG_16      TAAFVCLVALAAQLWRGRAGRRRYAHPSVRYVCLPPERD 628
HSV2GPG_18      TAAFVCLVALAAQLWRGRAGRRRYAHPSVRYVCLPPERD 698
HSV2GPG_12      TAAFVCLVALAAQLWRGRAGRRRYAHPSVRYVCLPPERD 698
HSV2GPG_15      TAAFVCLVALAAQLWRGRAGRRRYAHPSVRYVCLPPERD 698
HSV2GPG_22      TAAFVCLVALAAQLWRGRAGRRRYAHPSVRYVCLPPERD 628
                ***************************************
```
