## Supplementary file 12 for "Contriving a chimeric polyvalent vaccine to prevent infections caused by Herpes Simplex Virus (Type-1 and Type-2): an exploratory immunoinformatic approach"

```
Alignment GLP UL45

HS1UL45_4       ------------------------------------------------------------
HS2UL45_16      MPLRASEHAYRPLGPGTPPMRARLPAAAWVGVGTIIGGVVIIAALVLVPSRASWALSPCD 60
HS1UL45_6       MPLRASEHAYRPLGPGTPPVRARLPAAAWVGVGTIIGGVVIIAALVLVPSRASWALSPCD 60
HS1UL45_8       ------------------------------------------------------------
HS1UL45_5       ------------------------------------------------------------
HS1UL45_9       ------------------------------------------------------------
HS1UL45_10      MPLRASEHAYRPLGPGTPPMRARLPAAAWVGVGTIIGGVVIIAALVLVPSRASWALSPCD 60
HS1UL45_11      ------------------------------------------------------------
HS1UL45_2       ---------------------------------------------VLVPSRASWALSPCD 15
HS1UL45_7       ------------------------------------------------------------
HS1UL45_3       ------------------------------------------------------------
HS1UL45_1       ------------------------------------------------------------
HS2UL45_13      MAFRASGPAYQPLAPAASPARARVPAVAWIGVGAIVGAFALVAALVLVPPRSSWGLSPCD 60
HS2UL45_15      ------------------------------------------------------------
HS2UL45_12      ------------------------------------------------------------
HS2UL45_14      ------------------------------------------------------------


HS1UL45_4       ----------ISWDPTPMEHEQAVGGCSAPATLIPRAAAKQLAAVTRVQSARSSGYWWVS 50
HS2UL45_16      SGWHEFNLGCISWDPTPMEHEQAVGGCSAPATLIPRAAAKQLAAVTRVQSARSSGYWWVS 120
HS1UL45_6       SGWHEFNLGCISWDPTPMEHEQAVGGCSAPATLIPRAAAKQLAAVARVQSARSSGYWWVS 120
HS1UL45_8       ----------ISWDPTPMEHEQAVGGCGAPATLIPRAAAKQLAAVARVQSARSSGYWWVS 50
HS1UL45_5       ----------ISWDPTPMEHEQAVGGCSAPATLIPRAAAKQLAAVARVQSARSSGYWWVS 50
HS1UL45_9       ----------ISWDPTPMEHEQAVGGCSAPATLIPRAAAKQLAAVARVQSARSSGYWWVS 50
HS1UL45_10      SGWHEFNLGCISWDPTPMEHEQAVGGCSAPATLIPRAAAKQLAAVARVQSARSSGYWWVS 120
HS1UL45_11      ----------ISWDPTPMEHEQAVGGCSAPATLIPRAAAKQLAAVARVQSARSSGYWWVS 50
HS1UL45_2       SGWHEFNLGCISWDPTPMEHEQAVGGCSAPATLIPRAAAKQLAAVARVQSARSSGYWWVS 75
HS1UL45_7       ----------ISWDPTPMEHEQAVGGCSAPATLIPRAAAKQLAAVARVQSARSSGYWWVS 50
HS1UL45_3       ----------ISWDPTPMEHEQAVGGCSAPATLIPRAAAKQLAAVARVQSARSSGYWWVS 50
HS1UL45_1       ----------ISWDPTPMEHEQAVGGCSAPATLIPRAAAKQLAAVARVQSARSSGYWWVS 50
HS2UL45_13      SGWQEFNAGCVAWDPTPVEHEQAVGGCSAPATLIPRAAAKHLAALTRVQAERSSGYWWVN 120
HS2UL45_15      ----------VAWDPTPVEHEQAVGGCSAPATLIPRAAAKHLAALTRVQAERSSGYWWVN 50
HS2UL45_12      ----------VAWDPTPVEHEQAVGGCSAPATLIPRAAAKHLAALTRVQAERSSGYWWVN 50
HS2UL45_14      ----------VSWDPTPVEHEQAVGGCSAPATLIPRAAAKHLAALARVQAERSSGYWWVS 50
                          ::*****:*********.************:***::***: ********.

HS1UL45_4       GDGIRACLRLVDGVGGIDQFCEEPALRICYYPRSPGGFVQFVTSTRNALGLP 102
HS2UL45_16      GDGIRACLRLVDGVGGIDQFCEEPALRICYYPRSPGGFVQFVTSTRNALGLP 172
HS1UL45_6       GDGIRACLRLVDGVGGIDQFCEEPALRICYYPRSPGGFVQFVTSTRNALGLP 172
HS1UL45_8       GDGIRACLRLVDGVGGIDQFCEEPALRICYYPRSPGGFVQFVTSTRNALGLP 102
HS1UL45_5       GDGIRACLRLVDGVGGIDQFCEEPALRICYYPRSPGGFVQFVTSTRNALGLP 102
HS1UL45_9       GDGIRACLRLVDGVGGIDQFCEEPALRICYYPRSPGGFVQFVTSTRNALGLP 102
HS1UL45_10      GDGIRARLRLVDGVGGIDQFCEEPALRICYYPRSPGGFVQFVTSTRNALGLP 172
HS1UL45_11      GDGIRARLRLVDGVGGIDQFCEEPALRICYYPRSPGGFVQFVTSTRNALGLP 102
HS1UL45_2       GDGIRACLRLVDGVGGIDQFCEEPALRICYYPRSPGGFVQFVTSTRNALGLP 127
HS1UL45_7       GDGIRACLRLVDGVGGIDQFCEEPALRICYYPRSPGGFVQFVTSTRNALGLP 102
HS1UL45_3       GDGIRACLRLVDGVGGIDQFCEEPALRICYYPRSPGGFVQFVTSTRNALGLP 102
HS1UL45_1       GDGIRACLRLVDGVGGIDQFCEEPALRICYYPRSPGGFVQFVTSTRNALGLP 102
HS2UL45_13      GDGIRTCLRLVDSVSGIDEFCEELAIRICYYPRSPGGFVRFVTSIRNALGLP 172
HS2UL45_15      GDGIRTCLRLVDSVSGIDEFCEEL---------------------------- 74
HS2UL45_12      GDGIRTCLRLVDSVSGIDEFFEELAIRICYYPRSPGGFVRFVTSIRNALGLP 102
HS2UL45_14      GDGIRACLRLVDSVSGIDQFCEEPAIRICYYPRSPGGFVRFVTSIRNTLGLP 102
                *****: *****.*.***:* **
```
